# Atomistic TCR-ligand interactions shape memory T-cell differentiation

**DOI:** 10.1101/2025.11.05.686789

**Authors:** Aoi Akitsu, Kemin Tan, Robert J. Mallis, Matthew A. Booker, Jonathan S. Duke-Cohan, Kristine N. Brazin, Evan H. Kirkpatrick, Andrew N. Parkins, Amelia G. Seabury, Sarita Aryal, Vincenzo Cinella, Jonathan J. Lee, Kaveri I. Uberoy, Jenna K. Koenig, Matthew Biddle, Cameron M. Messier, Patrick H. Lizotte, Michael Y. Tolstorukov, Wonmuk Hwang, Matthew J. Lang, Ellis L. Reinherz

## Abstract

Memory CD8⁺ T cells provide durable protection against recurrent infection and cancer, but how distinct memory fates are specified remains unclear. Here, we define the functional landscape of 242 murine CD8⁺ TCRαβ clonotypes specific for influenza NP_366–374_/H-2Dᵇ ligand (pMHC) by integrating single-cell transcriptomics and TCR sequencing with force-dependent biophysics, structural analysis, and in vivo investigation of memory development. Fate essentially tracks with TCR mechanotransduction: T_CM_-associated clonotypes preferentially engage pMHC through TCRβ, whereas T_EM_-associated clonotypes preferentially engage through TCRα. Clonotypes with balanced, bipolar signaling generate both memory subsets, optimize expansion, and display broad heterosubtypic crossreactivity but with exhaustion susceptibility. In contrast, diverse T_CM_ clonotypes are largely clonal singlets with considerable mutant-epitope recognition in aggregate and stemness preservation, revealing complementary strategies for immunodefense coverage. Atomistically, subtle pMHC contact differences and load transmission through the TCR holoreceptor appear to tune phosphorylation, memory differentiation, and functional durability, features applicable to adoptive T-cell immunotherapies.

## Introduction

Immunological memory is a hallmark of the adaptive immune system, enabling rapid and robust targeted responses upon re-exposure to pathogens or tumors^1–4^. CD8⁺ T cells play a central role in this process by first recognizing threats through their clone-specific (clonotypic) TCRs and then eliminating infected or malignant cells by elaborating cytotoxic molecules and cytokines. Following acute viral infection, a small population of effector CD8⁺ T cells that survive the predominant contraction phase differentiates into long-lived memory cells to provide durable protection ^1,2^. Memory CD8⁺ T cells are functionally and phenotypically heterogeneous but broadly classified into two subsets of circulating lymphocytes, termed central memory (T_CM_) and effector memory (T_EM_) cells ^3^. A non-circulating subset of tissue-resident memory T cells (T_RM_) also forms ^4^. T_CM_ and T_EM_ cells differ in surface markers, migratory behavior, functional potential, and transcriptional and epigenetic landscapes ^4^. T_CM_ cells express lymphoid-homing receptors such as CD62L and CCR7, enabling their lodging in secondary lymphoid organs. They also express transcription factors including Tcf1 ^5^, Lef1 ^6^, and Id3 ^7^ which support T_CM_-like properties such as self-renewal, long-term persistence, and a robust proliferative response upon antigen re-encounter ^8,9^. In contrast, T_EM_ cells lack CD62L and CCR7, allowing them to circulate through the blood and access both lymphoid and non-lymphoid tissues. They preferentially express Id2 ^10,11^, which promotes effector differentiation, and are enriched for cytotoxic mediators such as granzyme B (Gzmb) and granzyme K (Gzmk), positioning them as a frontline defense capable of exerting immediate effector function ^1,3^.

The generation of memory subsets during viral infection is believed to be shaped by both extrinsic factors such as antigen abundance, inflammatory cytokines, and the tissue microenvironment as well as intrinsic transcriptional regulators and metabolic programs ^1,12,13^. How memory fate is ultimately determined, however, remains unresolved. Notwithstanding, three conceptual models have been proposed ^1^. The first, the intrinsic memory T cell fate model, suggests that fate is imprinted in naïve T cells prior to activation, i.e. the TCR *per se* substantially directs memory T cell development. The second, the linear differentiation model, suggests that memory precursor cells (T_MP_) give rise to memory subsets sequentially from T effector cells (T_EFF_) to T_EM_ followed by T_CM_, with the latter especially pronounced in the context of high precursor frequencies ^8,14–16^. The third, so-called signal strength model, alternatively posits that the intensity of TCR and co-stimulatory signaling during priming governs fate decisions, with stronger signals promoting T_EM_ differentiation and weaker signals favoring T_CM_ ^17–19^. These models are largely based on several TCR transgenic systems, with or without altered peptide ligands (APLs) utilized to tune TCR signal strength. While elegant, these systems cannot recapitulate the diversity of TCRs and T-cell competition operative in the mammalian repertoire.

As recent studies have shown that TCR ligand recognition specificity and sensitivity are regulated by T-cell sensing of physical load placed on a TCR-pMHC bond during immune surveillance ^20–31^, we examined the biophysical and structural contributions of this mechanosensing to intrinsic CD8 memory fate within a single TCR repertoire. Our results indicate that memory T_EM_ and T_CM_ differentiation is unrelated to TCR signaling threshold performance metrics such as bond lifetime *per se*. Moreover, although T_CM_ favor slip bonds rather than catch bonds, denoting linkages weakening versus strengthening under load, respectively, this is not a defining characteristic. Instead, differentiation is dependent upon disparate signaling linked to the structural features of the TCR-pMHC interaction. Key factors include interface surface topology and mechanochemistry that facilitate physical load transfer involving dynamic TCR behaviors under force that likely impact holoreceptor mechanotransduction. These attributes suggest an atomistic basis for how subtle differences in structures and dynamic interactions among nearly identical αβTCRs with a single pMHC ligand orchestrate different memory fates of the corresponding T cells. Our collective data strongly support memory fate model 1.

## Results

### TCR clonotypes of varying T-cell burst sizes and polarities

To investigate whether αβTCRs *per se* impact memory T cell subset fate, we performed single-cell RNA Sequence (scRNA-Seq) on NP_366-374_/D^b^-specific CD8 T cells isolated from pooled mediastinal lymph nodes (mLNs) of 10 B6 mice at 34 days post-IAV infection (dpi 34) using the mouse-adapted A/Puerto Rico/8/1934 (PR8) strain (Figure S1A). A total of 5,429 CD8 T cells comprising 242 clonotypes with paired TCRα and TCRβ V domain sequences were identified. Uniform Manifold Approximation and Projection (UMAP) analysis revealed two major clusters corresponding to central memory CD8 T cells (T_CM_) and effector memory CD8 T cells (T_EM_), appearing as left- and right-sided lobulated aggregates, respectively, (Figure 1A). Furthermore, differentially expressed genes (DEGs) and hierarchical clustering (Data S1) discriminated three distinct T_CM_ clusters and three T_EM_ clusters. Of the former, T_CM-CONV_ (T_CMC_) expressed conventional T_CM_ genes (*Klf2, Id3, Tcf7, Ccr7*), T_CM-XCL1_ (T_CMX_) specifically expressed *Xcl1, Cxcl10, and Mif*, and T_CM-PRO_ (T_CMP_) was enriched for proliferative genes (histone-related genes, *Stmn1, Pclaf, Birc5, and Ube2x*). With respect to the T_EM_, T_EM-CONV_ (T_EMC_) cluster expressed conventional T_EM_ genes (*Gzmb, Gzmk, Id2, Ccl5*), T_EM-KLR_ (T_EMK_) was enriched for KLR genes (*Klrd1 and Klrc1*), and T_EM-EXH_ (T_EME_) expressed exhaustion-associated genes (*Pdcd1*, *Lag3*, *Havcr2*). Selected top genes are shown in Figure 1B. Independent gene signatures derived from T_CM_ and T_EM_ clusters, identified by separate scRNASeq analysis on CD8β^+^CD44^+^ memory cells isolated by FACS from pooled mLNs of four B6 mice at dpi 33 (Methods), validated the transcriptional identity of these subsets specifically in the IAV context (Figure 1C and Figure S1B). Interestingly, *Sell*-encoded CD62L, a canonical T_CM_ cell surface marker, was weakly expressed even within the T_CM_ cluster, as reflected by the low frequency on NP_366-374_/D^b^-specific memory CD8^+^ T cells (Figures S1C and S1D), compared to the level on PA_224-233_/D^b^-, and PB1₇₀₃-₇₁₁/K^b^-CD8⁺ T cells directed at other individual IAV specificities. Hence, the expression of classical memory subset markers appears to be modulated in an epitope-specific manner.

**Figure 1.**
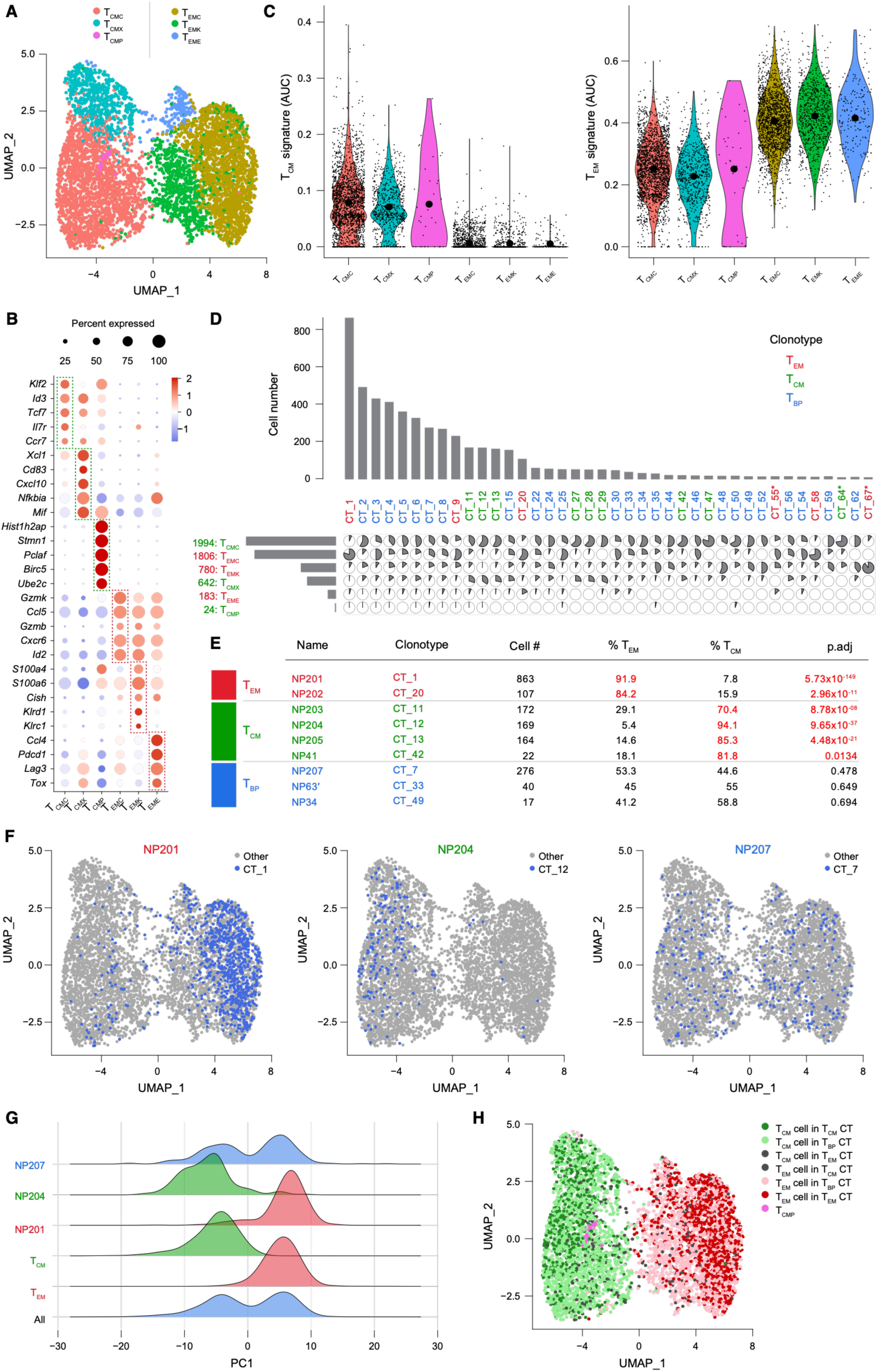
TCRαβ clonotypes intrinsically influence memory CD8^+^ T cell fate. (A) UMAP clustering of NP_366-374_-specific memory CD8⁺ T cells. (B) Dotplot of select immune-related genes. Circles are proportional to the fraction of cells with expression of the indicated gene. Red indicates high expression relative to other clusters while blue indicates low expression. (C) Violin plots of T_CM_ and T_EM_ gene signatures generated from an independent experiment and applied to the six clusters. The black dots show the medians. (D) Correlation matrix of the 40 most abundant clonotypes. Pie slice sizes are proportional to the fraction of the clonotype in each cluster. The barplot above the matrix indicates the number of cells with the given clonotype while the barplot to the left indicates the number of cells in each cluster. (E) Table of select clonotypes and their distribution across T_CM_ and T_EM_ clusters. T_CM_ and T_EM_ clonotypes are those for which more than 70% of cells are in T_CM_ or T_EM_ cluster, respectively. T_BP_ clonotypes are those for which fewer than 70% of cells are in a T_CM_ or T_EM_ cluster. P-values were generated using a binomial test (with adjustment by the Benjamini-Hochberg procedure) relative to the overall distribution T_CM_ or T_EM_ cells. The NP63’ designation refers to the clonotypes usage of TRAV16 for CT_33, which differs at two residues from TRAV16N used by NP63 (14V→M and 61V→E), sites distal to the pMHC interface. (F) Blue dots on UMAPs showing which cells bear the indicated clonotypes. (G) Ridge plot indicating the position of cells across the first principal component. Sets of cells include all cells, cells from T_CM_ clusters, T_EM_ clusters, and select clonotypes. (H) UMAP with cells colored by the indicated CT category and memory cell types comprising each.

We analyzed the distribution of clonotypes across the UMAP clusters to determine whether TCRs significantly influence memory fate (Figure 1D), identifying those with single TCRα and TCRβ chains after filtration (Data S2 and Methods). Among the top 40 clonotypes, 25 were distributed comparably amongst subsets of T_CM_ vs T_EM_ clusters. In contrast, the remaining 15 clonotypes were highly skewed toward one or the other subsets, suggesting that TCRs *per se* impact memory T cell fate as indicated by the low adjusted p-values (Figure 1E and Table S1). Accordingly, clonotypes were categorized as belonging to the T_CM_ vs T_EM_ type when >70% of the cells in each clone’s progeny were polarized toward one or more of the three respective subclusters therein, as defined in Figure 1A. Other clonotypes with a more equivalent distribution not achieving the 70% threshold heuristic were classified as bipolar (T_BP_). Representative clonotypes listed in Figure 1E underwent further analysis. The NP41, as well as NP34 and NP63, TCRs identified in a previous study of the IAV memory recall response ^30^ were also observed here in the primary memory pool, as one T_CM_ and two T_BP_ clonotypes, respectively. Moreover, since both analogue NP34 and digital NP63 are categorized as T_BP_, their different performance under pN forces (i.e. digital TCRs need only a few pMHC for activation while analogue TCRs require dozens to hundreds ^30^), is not linked to the T memory subset determination. The cell projection on the UMAP (Figure 1F) and the Euclidean distance of principal component-1 (PC1) for NP201, NP204, and NP207 (Figure 1G) confirmed their locations in T_CM,_ T_EM,_ or both clusters, highlighting gene expression similarity between individual clonotypes and the overall T_CM_ and T_EM_ populations as well as total T cells.

We further examined the localization of all cell types belonging to each memory clonotype category (CT) and projected their distribution on the UMAP of the entire CD8 repertoire (Figure 1H). “CT-matched” cells (i.e., T_CM_ cells in T_CM_ CTs and T_EM_ cells in T_EM_ CTs) are localized primarily toward the periphery of the respective T_CM_ and T_EM_ regions, whereas T_CM_ and T_EM_ cells in T_BP_ CTs are diffusely spread but with more concentrated toward the center of the UMAP. This differential distribution suggests that CT-matched cells express their respective signature genes more strongly than T_BP_ clonotypes or those rare “CT-mismatched” cells (i.e., T_CM_ cells in T_EM_ CTs and T_EM_ cells in T_CM_ CTs), thereby enhancing their fate determination. Consistent with this, RNA velocity analysis supports the notion that T_CM_ and T_EM_ clonotypes represent more terminally differentiated states, as directional flow arrows from CT-matched cells are either absent or pointed toward the outer UMAP edges (Figure S2A). In contrast, T_BP_ clonotypes appear more interchangeable, as the arrows from T_CM_ and T_EM_ cells in T_BP_ CTs indicate some bidirectional flow.

To determine whether the expression of all signature memory genes or only a specific subset underlie the memory polarization, we analyzed differentially expressed genes in T_EM_ and T_CM_ clusters (Data S3) across CT-matched cells, CT-mismatched cells, and T_CM_ and T_EM_ cells in T_BP_ CTs. Both aggregate T_EM_ and T_CM_ signature gene expressions increased progressively with clonotype polarization (Figure S2B and S2C). Of the 126 T_EM_ signature genes elevated in the T_EM_ cluster, only 45 genes, including *Ccl5*, *Gzmb*, *Gzmk*, and *Fasl* showed a gradual increase in expression corresponding to the degree of clonotype polarization, suggesting that these genes may contribute directly to clonotype fate commitment (Figure S2B). Similarly, of 140 T_CM_ genes, only 65 genes, such as *Lef1*, *Tcf7*, and *Tnfsf8* exhibited graded expression aligned with clonotype polarity, implying they are key regulators of T_CM_ fate (Figure S2C). How gene expression polarity is mediated by TCRs and their signal transduction pathways in mechanistic terms is a subject we shall return to.

### The T_CM_ repertoire is the most variegated

Having defined clonotype “polarity” from paired TCR-transcriptome data, we next asked whether TCR sequence architecture itself predicts these fate-linked programs across the repertoire. We therefore interrogated TCR gene usage, clonal expansion (“burst size”), and sequence relatedness to identify repertoire-level features associated with the T_CM_, T_EM_, and/or T_BP_ transcriptional phenotype. V and J gene usage analysis revealed the preferences for TRAV16N/TRAV16, TRBJ2-2/TRBJ2-7, and TRBV13-1 among all 242 NP-specific clonotypes (Figure 2A), consistent with a previous study ^32^. Notably, the 140 T_CM_ clonotypes exhibited greater diversity in V and J gene usage, while 44 T_BP_ clonotypes showed a more restricted repertoire, with a higher frequency of the dominant VJ combinations found across the NP-specific TCRs. The 58 T_EM_ clonotypes displayed an intermediate pattern. Entropy analysis confirmed the differences in VJ diversity among the memory subsets (Figure S3A).

**Figure 2.**
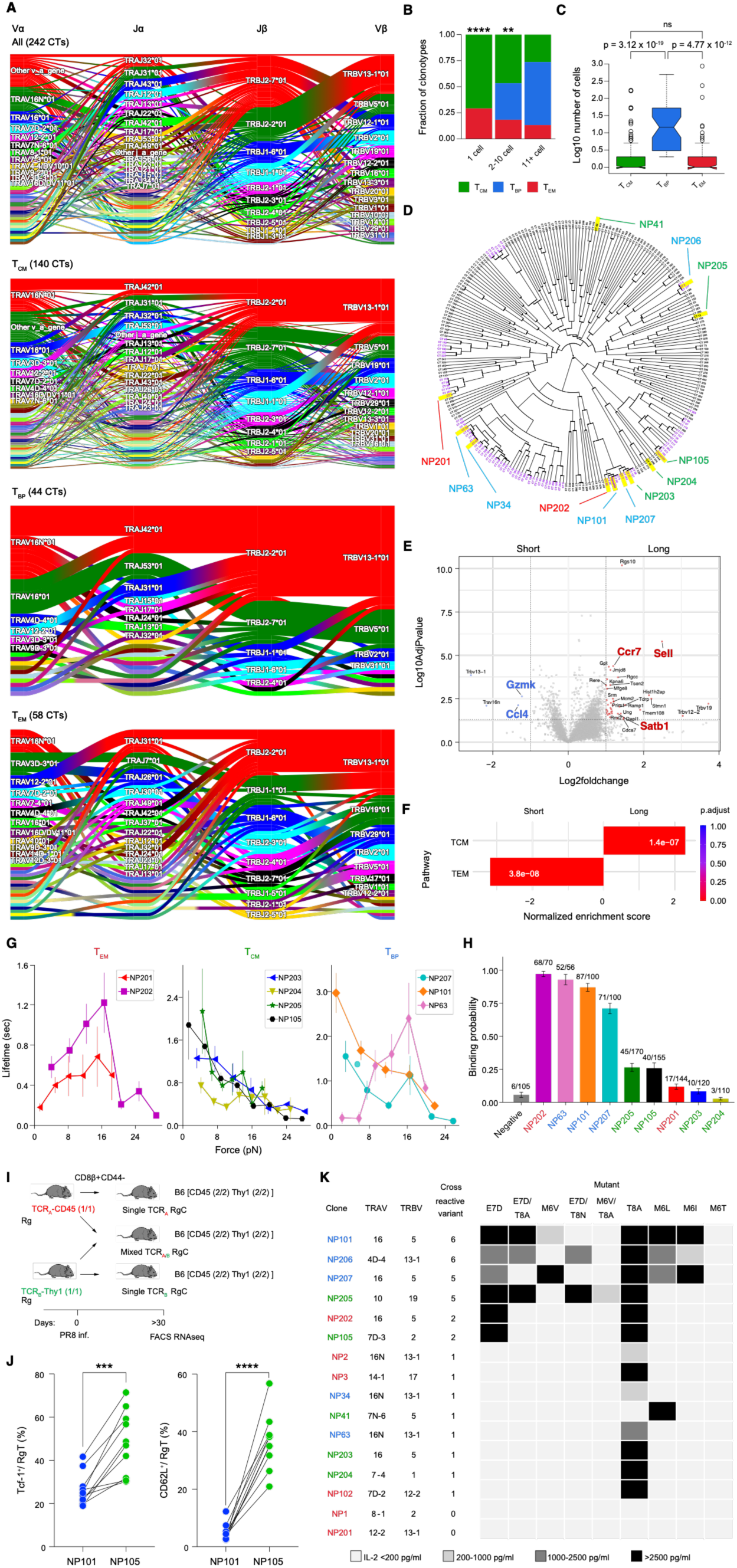
Distinct features of memory T cell subsets according to their polarities. (A) Sankey plot of V and J-gene usage by clonotype category. (B) Stacked barplot of clonotype type distribution binned by cell number. The p-values by chi-squared test for comparison between T_CM_ and T_EM_ are: 1 cell vs overall distribution of cells: ****; p = 7.58 x 10^-7^, 2-10 cells: **; p = 0.0093, and 11+ cells: p = 0.29. (C) Boxplot of cell number by clonotype type. P-values for comparisons were calculated by Mann-Whitney-U test. (D) Phylogram of clonotypes generated by hierarchical clustering of TCR distances with complete linkage. Clonotypes that are sequence similar to multiple other clonotypes (“short branch”) are shaded in purple while the remaining clonotypes (“long branch”) are in black. Representative T_EM_ clonotypes are highlighted in red, T_CM_ in green, and T_BP_ in blue. (E) Volcano plot of differential expression of clonotypes in the long branch category versus short branch category. (F) Gene Set Enrichment Analysis of the differential expression in E using the T_CM_ and T_EM_ gene signatures. (G) Force vs. lifetime plot for T_EM_ clones NP201 (red curve, N=56), NP202 (magenta curve, N=136); T_CM_ clones NP203 (blue curve, N=58), NP204 (yellow curve, N=110), NP205 (green curve, N=79), NP105 (black curve, N=101); and T_BP_ clones NP101 (orange curve, N=103), NP207 (cyan curve, N=100) And NP63 (pink curve, N=40) with its cognate ligand NP_366-374_/Db. Error bars indicate SEM. (H) Tether forming probability for the SM optical tweezers assay under identical conditions of tethers per bead and surface ligand density. Error bars indicate SEM. (I) Experimental scheme for generating single and mixed retrogenic (RgC) mice using congenic markers (i.e., CD45.1/1, CD45.1/2 or Thy1.1) is detailed in Methods. (J) Tcf-1 and CD62L expression on Rg T cells in mLN of mixed NP101/105 RgC mice that were recipients of 5,000 Rg T cells bearing each clonotype (i.e. 10,000 cells total). (K) Crossreactivity profiles of 16 NP-specific TCRs against a panel of naturally occurring IAV variants carrying mutated NP peptides. Reactivity was assessed by measuring IL-2 production from TCR-transduced BW cell lines following stimulation with peptide-loaded APC. Memory category color coding matches that in (D) Data is the representative of two or three independent experiments.

Next, we compared clonal burst sizes amongst memory populations. T_CM_ clonotypes were the largest fraction in lower cell number categories, particularly in the single-cell group, compared to T_EM_ clonotypes (Figure 2B). In contrast, T_BP_ clonotypes were significantly more expanded than either T_CM_ or T_EM_ clonotypes (Figure 2C). Since clonotypes with only one cell cannot be classified as T_BP_ by definition, we repeated the analysis excluding clonotypes with single-cell clonotypes and observed a consistent result (Figure S3B). These findings suggest that T_CM_ tend to persist as a small number of cells, whereas T_BP_ are more clonally expanded.

To distinguish clonotypes with similar versus divergent TCR sequences, we categorized branches of a TCR sequence tree into “short” and “long”, respectively, based on paired TCR sequence distance metrics (Figure 2D). For each clonotype, we calculated the distance to all other clonotypes and computed the mean of the top 10 shortest pairwise distances. For example, CT_25, categorized as a short-branch clonotype, had a mean top 10 distance of 7.8, a weighted amino acid sequence similarity metric, whereas CT_50, categorized as a long-branch clonotype, had a corresponding value of 196.4 (Figure S3C). A histogram of these mean top 10 distances across all clonotypes revealed a bimodal distribution, and we defined the valley between the two peaks (mean distance = 109) as the threshold distinguishing short- and long-branch clonotypes (Figure S3D). A TCR distance heatmap further supported the validity of this classification (Figure S3E). Unsurprisingly, analysis of short- and long-branch clonotypes showed that short-branch clonotypes exhibited more restricted V and J segments commonly used in NP-specific TCRs, whereas long-branch clonotypes displayed greater diversity (Figure S3F).

We then examined DEGs between short vs. long branch clonotypes. Short-branch clonotypes showed higher expression of T_EM_ genes such as *Gzmk* and *Ccl4*, whereas long-branch clonotypes expressed higher levels of T_CM_ genes, including *Sell*, *Ccr7*, and *Satb1* (Figure 2E and Data S4). Gene enrichment analysis confirmed a significant enrichment of T_EM_ gene signatures in short branches and T_CM_ gene signatures in long branches (Figure 2F). We further examined the distribution of representative memory clonotypes on the TCR tree, including NP101 and NP105, identified from an independent scRNA-Seq experiment (Methods) with NP101 showing identical amino acid sequence to CT_5 and CT_56 (Table S1) and NP105 bearing a significant relationship to NP204 described below. Representative T_EM_ clonotypes such as NP201 and NP202, as well as T_BP_ clonotypes including NP34, NP63, NP101, NP206, and NP207, fell within the short branch (Figure 2D). In contrast, T_CM_ clonotypes NP41, NP204, and NP205 were found in long branch, supporting the notion that TCR sequence similarity is associated with T_EM_ or T_BP_ phenotypes, while greater sequence divergence is linked to the T_CM_ phenotype.

To investigate the scope of this observation, we independently generated TCR distance trees for PA_224–233_/D^b^ [PA]-specific (Figure S3G) and PB1_703–711_/K^b^ [PB1]-specific TCRs (Figure S3H) from 10 and 35 B6 mice infected with PR8, analyzed at 34 dpi (PA) and 36 dpi (PB1), respectively, and identified short- and long-branch structures using the same criteria as in Figure S3C and S3D. Consistent with our initial findings, short-branch clonotypes were enriched for T_EM_ signature genes, whereas long-branch clonotypes preferentially expressed T_CM_ signature genes for both antigen-specific T cells (Figure S3I and S3J). Hence, TCR diversity is a hallmark of T_CM_ repertoires, challenging model 2 of memory fate determination. Nonetheless, departures can exist. Examples of deviations include T_CM_ clonotypes NP105 and NP203 belonging to the short branch (Figure 2D). In the case of NP203, this is probably explained by the TCR sequence differing by only 1-3 amino acids from short-branch T_BP_ clonotypes NP101 and NP207 and T_EM_ clonotype NP202. Hence, minor sequence differences can override the general TCR branching pattern to influence fate decisions despite overall amino acid similarity, as described in a later section.

### Weak TCR-NP_366-374_/D^b^ bonding profiles

Because sequence diversity alone does not specify mechanism, we next tested whether fate-biased clonotypes differ in the mechanics of their pMHC engagement under force. Single molecule (SM) TCRαβ-pMHC analysis was used to evaluate the performance and bonding characteristics of clonotypes belonging to each memory category (Figures S4A and S4B). T_CM_ clonotypes NP203, NP204, NP205, and NP105 were weak binders with all force lifetime profiles, except NP204, consistent with classic slip bonds, readily breaking with force application, unlike catch bonds that strengthen as pulled (Figure 2G). T_EM_ clonotypes NP202 and NP201 were also weak binders but exhibited more typical catch bond profiles with peaks around 15 piconewtons (pN). T_BP_ clonotypes NP101 and NP207 exhibited mixed profiles, with higher lifetime amplitudes at both the typical slip and catch force regions (5 and 15 pN, respectively), while NP63 was a catch bond as in previous work^30^. Additionally, overall lifetime amplitudes were weak relative to verified digital clonotypes such as NP63^30^ (Figure 2G and Figure S4C). All clonotypes showed the ability to undergo a conformational transition, described in detail below. The capacity to form a stable single-molecule TCRɑβ-pMHC interaction irrespective of load was determined by comparing the total number of binding opportunities allowed to the total number of events measured. Stable tethering formation varied substantially among the clonotypes, with T_CM_ clonotypes showing a notably lower tendency to form stable bonds, which also tended to bind in the slip-type manner (Figure 2H).

Excluding NP63, ranking relative performance based on lifetime amplitude around 15 pN in addition to the tendency to form stable tethers, suggests that T_EM_ clonotype NP202 and T_BP_ clonotypes NP207 and NP101 are a tier above T_CM_ clonotypes NP203, NP205, and NP105. NP204 and NP201 are the lowest performing clonotypes, exhibiting low lifetime amplitudes across all force ranges measured, coupled with a poor ability to form stable bonds measured by binding probability (Figure 2H). Binding probability was strongest for NP202 (and NP63) where multiple tethers were observed, compared to the other clonotypes profiled. Collectively from these data, TCR biophysical performance *per se* is not the singular correlate of the transcriptome of the cell displaying the receptor.

The slip bond character of most T_CM_ might suggest a greater role for CD8 co-receptor in the bidentate interaction with pMHC ^33^. This is the case for NP204 and NP205 but not for NP203 as revealed by FACS-based ligand binding to T cell transfectants using WT CD8 binding-competent NP_366-374_/D^b^ tetramers versus NP_366-374_/D^b^ α3 domain point mutant CD8 binding-defective NP_366-374_/D^b^ tetramers (Figure S4D). However, the NP206 T_BP_ (Table S1) is also impacted, excluding strict clonotype co-receptor dependency as limited to T_CM_. Notably, comparable micromolar affinity interactions at equilibrium between NP_366-374_/D^b^ and T_EM_ NP202, T_BP_ NP207 and T_CM_ NP105 TCRs were observed by Surface Plasmon Resonance (SPR) analysis (see kinetic and equilibrium data in Data S5), showing a trend consistent with the lowest force lifetime bin of each in Figure 2G.

### Retrogenic profiling reprises polarization

These results suggested that force-dependent binding metrics, while informative, are insufficient on their own to explain fate bias. We therefore asked whether the TCR is sufficient to impose polarization *in vivo*. To determine whether αβTCRs imbue the CD8 differentiative fate and memory phenotypes identified by scRNA-Seq following antigen encounter *in vivo*, we generated retrogenic chimera (RgC) mice. This was achieved by adoptively transferring naïve congenically marked CD8⁺ retrogenic (Rg) T cells from Rg donor mice into B6 [CD45 (2/2) Thy1.2] mice, followed by infection with the PR8 one day later (Figure 2I). At 30 dpi in mixed NP101/NP105 RgC mice, NP105 Rg T cells exhibited significantly higher expression of T_CM_ markers, CD62L and Tcf-1, compared to NP101 Rg T cells (Figure 2J and Figure S5A), indicating that the RgC system faithfully recapitulates clonotype polarization patterns identified by scRNA-Seq. In a second example, we assessed the rapid *in vivo* recall response of Rg T cells NP202, NP204, and NP207 at day 4 after secondary IAV challenge, given that T_CM_ characteristically expand rapidly ^8,34^. As shown in Figure S5B, the T_CM_-type NP204 manifest the best expansion, greater than T_EM_-type NP202 in LN and lung (p<0.01) and trended better than T_BP_-NP207, particularly in mLN. In sum, the findings using Rg mice support the view that the clonotype confers differentiative fate. in further support of model 1 where fate is predetermined.

### Memory IAV crossreactivities

We next assessed whether these fate-linked clonotype classes also segregate by functional breadth, specifically heterosubtypic recognition of naturally occurring NP variants. The role of T cells in providing heterosubtypic IAV protection to largely conserved epitopes is well established ^35^ despite limited structural analysis ^36^. To ascertain if such crossreactivity could be extended more broadly to various memory clonotypes recognizing NP_366-374_/D^b^, we selected 16 TCRs that were transduced into mCD8αβ^+^BW5147.3 (BW) cells and sorted for comparable TCR and CD8αβ expression. BW transductants were then stimulated with WT-PR8 NP_366-374_ peptide or 9 naturally occurring variant peptides derived from 11 other IAV strains ^37^ on APCs. IL-2 production was assayed as a readout of TCR activation (Figure 2K). The importance of the structurally featured residues of the peptide (p4E, p6M and p7E) were also quantified (Figures S6A and S6B) and referenced to the IAV variants (Figure S6C). Crossreactivity was observed in 14 of 16 TCRs, ranging from 1-6 of the 9 variants, with all TCRs responding to the isologous PR8 strain (Figure 2K and Figures S6B and S6D). As 0/6 T_EM_, 1/5 T_CM,_ and 3/5 T_BP_ recognized 5 or more viral epitopes, T_BP_ recognition trended as more crossreactive compared to the polar clonotypes (p = 0.0632 using a two-tailed Fisher’s exact test). The greatest crossreactivity was exclusive to the T_BP_ (NP101, NP206 and NP207). As an example, detailed IL-2 production from NP207 is further quantified by peptide titration (Figure S6E). Binding probabilities measured by the SM assay independently reflect the IL-2-based crossreactivity patterns for NP207, NP203, and NP202 (Figure S6F) along with bond lifetimes and/or structural transition frequencies in the memory pool (Figure S6G).

### Protein expression, crystallization, and analyses

To define structural features that could encode polarity and crossreactivity, we determined crystal structures of representative TCR-NP_366-374_/Dᵇ complexes spanning T_EM_, T_CM_, and T_BP_ phenotypes. To that end, we engineered mouse TCRs and TCR chimeras with murine V domains fused to human C domains (sequences in Data S6). All TCRαβ heterodimers were generated in eukaryotic Expi293F N-acetylglucoseaminotransferase I-deficient (GnTI^-^) cells to facilitate deglycosylation under native conditions. H-2D^b^ and β2m proteins were produced in *E. coli* and refolded *in vitro* with chemically synthesized NP_366-374_ peptide to generate pMHC complexes. Procedures for purifications, crystallization screening, and data collection are detailed in Methods. A series of crystals of unligated TCRs and their complexes were obtained and structurally determined. Crystallographic statistics, complex interface buried surface areas (BSA) and surface complementarity (Sc values) along with H-bonding and van der Waals interactions are given in Table S2-S4.

Figure 3A tabulates information on the six TCRs from B6 mice that recognize the D^b^-bound NP_366-374_ PR8 IAV epitope used for structural analysis. Four sequence-similar TCRs are representative of T_EM_ (NP202), T_BP_ (NP101, NP207), and T_CM_ (NP203). It is striking that the TCRs utilize identical TRAV and TRBV gene segments differing by only 1-3 CDR3 amino acid residues and yet belong to functionally distinct memory CD8 subsets (Figure 3A). We also selected the NP204 and NP105 T_CM_ TCRs comprising distinct V gene segments but with similar CDR3β loops given a shared TRBJ1-6 differing by just 4 of 17 residues. Note that CT_29 (Table S1) is a T_CM_ like NP105 and NP204. CT_29 is most similar to NP105 with utilization of the same TRAV, TRBV, TRBJ, but a different TRAJ.

**Figure 3.**
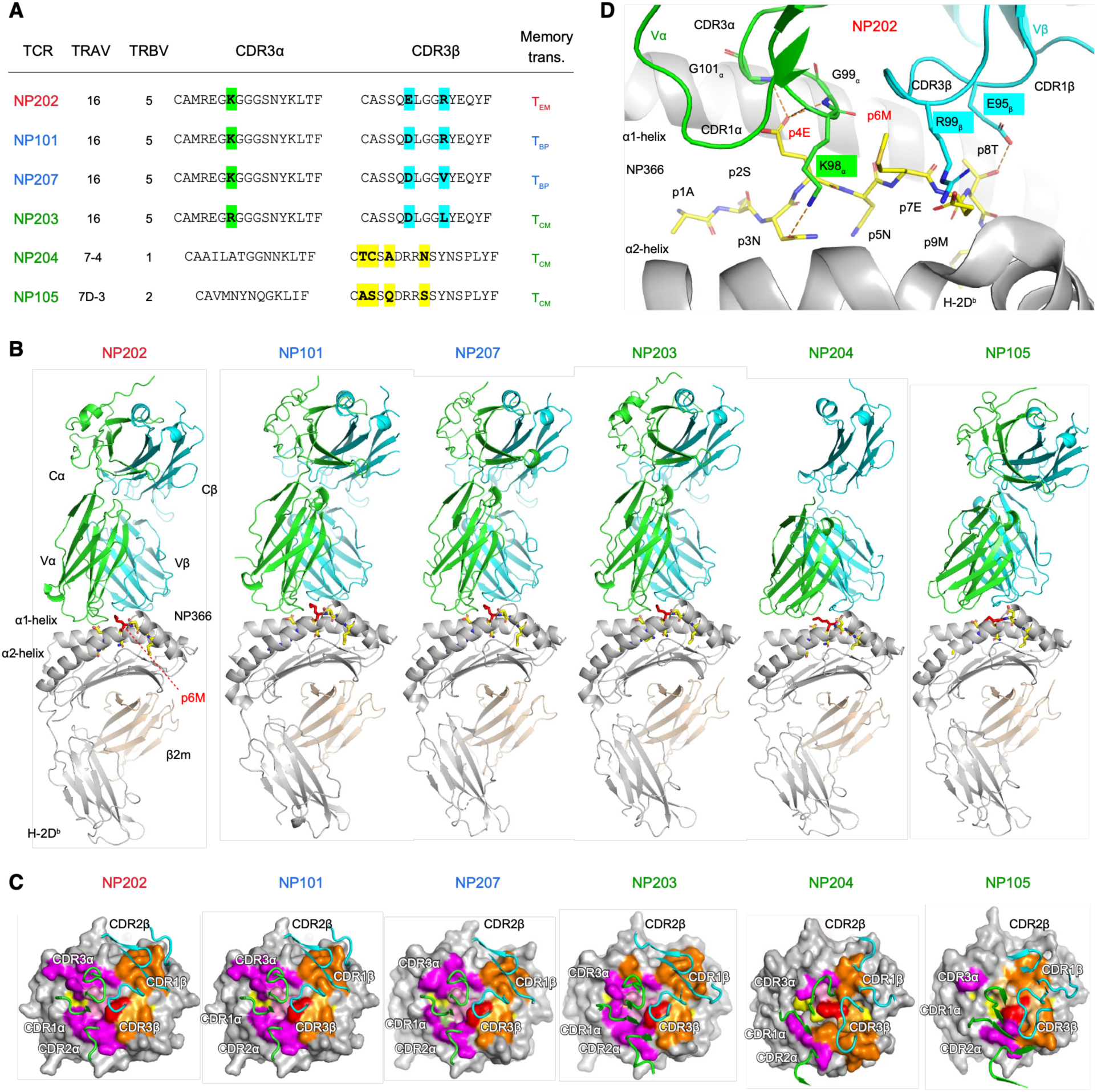
TCR sequences and structures of six NP TCR-NP_366-374_/D^b^ complexes derived from T_EM_, T_CM,_ and T_BP_ memory CD8 T cells. (A) The genes and CDR3 sequences of six NP TCRs. In the first four NP TCRs, the three residues that vary among their CDR3s are highlighted in green or cyan. Between NP105 and NP204 key CDR3β residues are in common as a result of a shared TRBJ1-6. Only varied residues between two CDR3β are highlighted in yellow. (B) Ribbon diagrams of six NP TCR-NP_366-374_ **/**D^b^ complexes arranged in the order from T_EM_ (α-dominant) to T_CM_ (β-dominant) from left to right. Their α1α2 domains of MHC are positioned in the same orientation. The NP_366-374_ peptide is displayed in sticks with the central residue p6M highlighted in red. TCR Vα, TCR Vβ, NP_366-374_ peptide, MHC α chain and MHC β2m are colored in green, blue, yellow, grey and light orange, respectively. The Cα domain of NP 204 TCR is absent due to structural disorder. (C) TCR CDR loops and their footprints on pMHC. Magenta and pink surface areas represent the MHC and peptide residues that interact with TCR Vα, respectively. Orange and bright orange surface represent the MHC and peptide residues that interact with TCR Vβ, respectively. (D) A zoomed-in view of the interaction between TCR and peptide in the NP202-NP_366-374_ complex. The three varied CDR3 residues noted in (A) are highlighted in the same color schemes to show their positions and roles. All peptide residues are labeled, with p4E and p6M highlighted in red for their prominent roles in TCR’s recognition.

Each of the six TCRs adopts a canonical binding mode in their respective complex such that Vβ overlies the C-terminal half of the peptide and the MHC α1 helix while Vα overlies the N-terminal half of the peptide and MHCα2 (Figure 3B). Amongst the various TCR-NP_366-374_/D^b^ complexes, the docking modes of the four closely related clonotypes, NP202, NP101, NP207 and NP203 appear quite similar with comparable footprints on NP_366-374_/D^b^ (Figure 3C), complex interfaces (BSA and Sc) (Table S3). Even docking parameters (incidence angle, twist and shift) are rather similar (Figures S7A-S7C). Therefore, the above widely employed structural parameters cannot account for the observed TCR-linked transcriptome differences. Amongst these four related NP TCRs, distinct interactions with ligand and downstream signaling consequences become apparent following detailed atomistic structural examination, molecular dynamics simulation under load, and biochemical studies described below.

Those four highly similar interfaces are obviously different from the two footprints of the “Vβ-centric” NP105 and NP204 TCRs analyzed using the same set of parameters. We refer to NP105 and NP204 as Vβ-centric since the TCR-pMHC complex in each is dominated by structural interactions involving their respective Vβ domain. For these two Vβ-centric TCRs, the ratio of TCRα/TCRβ BSA in the TCR-pMHC complex is 0.26-0.59. In contrast, for the four NP 202, NP101, NP207, and NP203, the ratio is 1.23-1.33, due to considerably greater Vα interactions (Table S3).

### Atomistic basis of CD8 memory fates

The primary overall crystallographic features of the NP202 T_EM_ TCR clonotype’s interaction with NP_366-374_/D^b^ are shared with NP101, NP207, and NP203 interactions. The most exposed NP_366-374_ peptide residues extending upward from the MHC groove are p4E and p6M, and to a lesser extent, p8T (Figure S6A), in agreement with a prior pMHC structure ^38^. When the NP202 TCR docks on NP_366-374_/D^b^, the negatively charged carboxylic group of p4E is positioned below the center of a glycine-rich loop from CDR3α, forming 2-3 H-bonds to the amide groups (Figure 3D). The central p6M sidechain is sandwiched by CDR3α and CDR3β and constrained in a hydrophobic pocket involving aliphatic sidechains of K98α and R99β. An additional specific interaction is a H-bond between side chains of p3N and CDR3 K98α.

The NP101 T_BP_ TCR differs from the NP202 T_EM_ TCR by only a single CDR3β residue (D95β vs. E95β), as highlighted in Figure 4A. Consequently, the two TCR docking angles and binding sites on NP_366-374_/D^b^ are very similar (Figures S7A-S7C). Structural alignment of their H-2D^b^ α1α2 domains demonstrates excellent overlay of the peptide binding grooves and corresponding NP_366-374_ peptides except at the central p6M residue differing by a slight shift and side chain rotation. Although the single-residue substitution resides in CDR3β, the Vβ domain including CDR3β exhibits limited deviation (shortened arrow). However, a surprising positional shift is evident in the Vα domain comparison as a combined result of slightly different TCR docking angles onto the pMHC and deviations at the Vα-Vβ interface between the two TCR V modules (two long red arrows). This shift brings important pocket-forming residues, including F33α and Y54α (in green), as well as the aliphatic part of R99β (in cyan), closer to p6M, which significantly reduces the size of the hydrophobic pocket. For example, the Cα-Cα distance between D52α and R99β, a representative distance between CDR2α and CDR3β, is reduced from 7.78Å in NP202 to 6.80Å in NP101. Furthermore, apposition of the Vα C’ strand to the Vβ G strand is reflected in the formation of two salt bridges, R50α to E101β and D52α to R99β in NP101. Shrinking of the p6M hydrophobic pocket strengthens hydrophobic and van der Waals interactions with NP101. Meanwhile, the interaction between p4E and G100α is lost in NP101 (Table S4A). Together, the overall structural alterations foster repositioning of the TCR-pMHC interaction center from the p4E to the p6M residue, thereby doubling the number of p6M van der Waals contacts made by NP101 relative to NP202 (3 vs 6, Table S4B). This subtle difference is not revealed by TCR docking angles or footprints onto pMHC (Figures S7A-S7C or Figure 3C). A second T_BP_ clonotype, namely NP207, manifests a similar shift, akin to NP101, associated with TCR crossreactivity against variant NP_366-374_ epitopes derived from heterotypic IAV strains, as noted in Figure 2K and Figures S6B-S6E. Thus, subtle structural changes meaningfully impact function.

**Figure 4.**
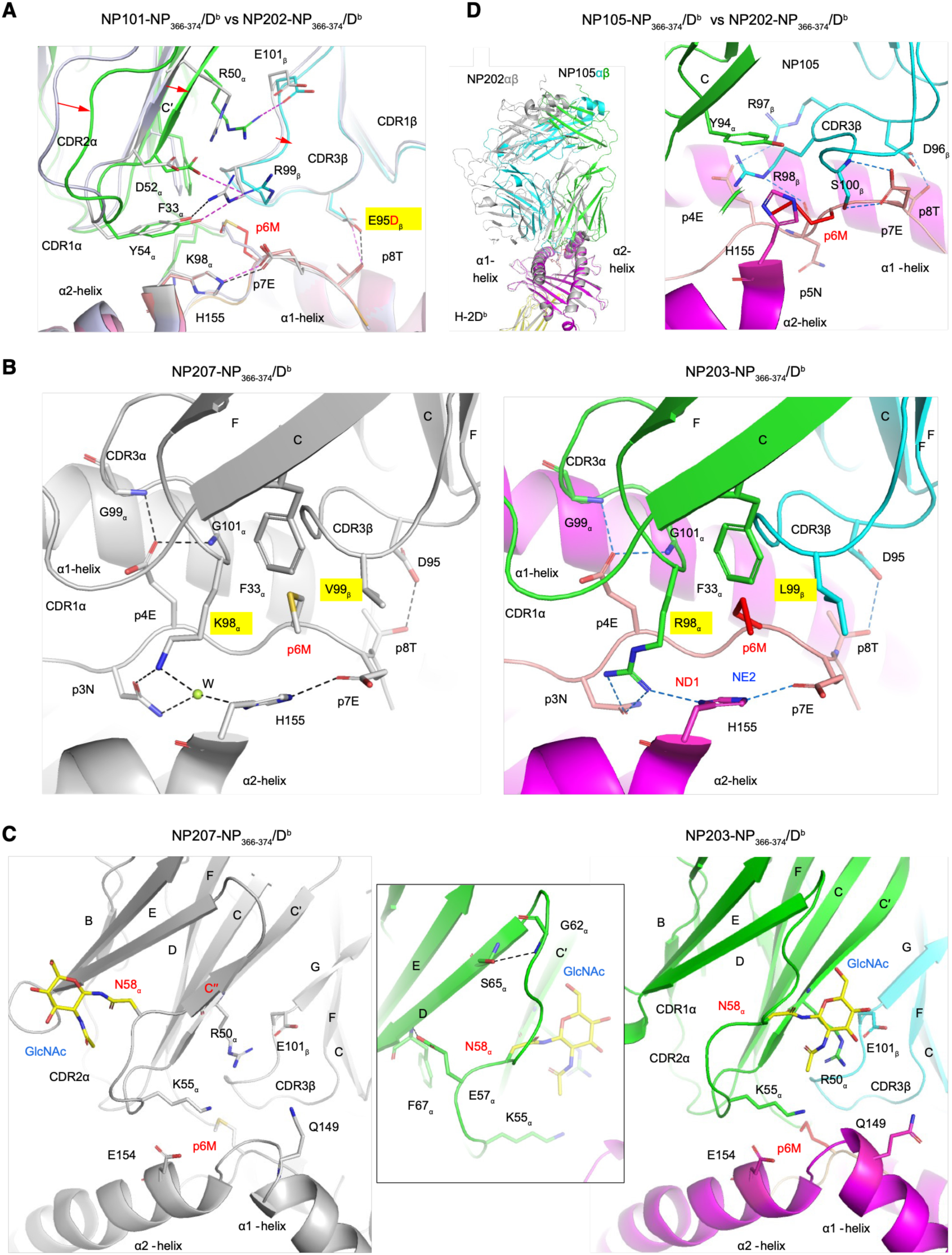
Fine structure details of TCR-ligand interactions in a single CD8 T cell repertoire linked to memory differentiation. (A) TCR-pMHC interface area of NP101 and NP202 complexes superimposed based on their D^b^ α1α2 MHC domains. The NP202 complex is colored in grey as a reference. The TCR α-chain, TCR β-chain, peptide and MHC of NP101 complex are colored in green, cyan, orange and pink, respectively. A few relevant residues are displayed in sticks for comparison. (B) Side-by-side comparison between NP207 and NP203 complexes. NP207 complex is colored in grey on the left with a structured water (W) represented as a green sphere. The TCR α-chain, TCR β-chain, peptide and MHC of NP203 complex are colored in green, cyan, pink, and magenta, respectively. Some key residues described in text are shown in sticks to display hydrogen bonds and salt bridges. The ND1 and NE2 atoms of H2-D^b^ H155 are labeled to show their potential protonation/deprotonation status. (C) Side-by-side comparison between NP207 and NP203 complexes in a view focusing on the C’(C”)-D edge of Vα domain. The NAG moiety attached to N58α is shown in sticks. In NP203 complex, the extended loop between C’ strand and D strand is stabilized by two hydrogen bonds between sidechain and mainchain as shown in the insert. (D) A docking comparison of β-dominated NP105 (in grey) with α-dominated NP202 (colored) on the left and a zoomed-in display of NP105 interaction with NP_366-374_ /D^b^ on the right, in which a part of α2-helix in front of peptide is omitted for clarity.

Consistent with this notion, NP207 (T_BP_) and NP203 (T_CM_) differ at two residues, K98α vs R98α in CDR3α and V99β vs L99β in CDR3β, respectively, as highlighted in Figure 4B. In the NP207-NP_366-374_/D^b^ complex, K98α forms a H-bond with p3N. A well-defined water molecule (labeled as W) also bridges the interactions between K98α and p3N to the ND1 atom of H155 in the D^b^ α2-helix. Meanwhile, p7E and NE2 of H155 form a salt bridge. However, in the NP203 complex, the longer R98α sidechain both pushes p3N slightly downward and displaces the water molecule found in the NP207 complex to form a salt bridge directly with the ND1 atom of H155 creating a thermodynamically altered interface. The ND1 atom of H155 is likely deprotonated to neutralize the extra positive charge introduced by the K→R substitution, while the NE2 of H155 is protonated and neutralized by p7E, thereby becoming indispensable in the NP203 TCR recognition of pMHC, in contrast to NP207, as elaborated below. The CDR3β L99 in NP203 (T_CM_) relative to CDR3β V99 in NP207 (T_BP_) augments van der Waals contacts with p6M, whereas the H-bonds of p4E to CDR3α are weakened with the bond length (p4E to G101α) increase from 2.97Å to 3.34Å (Table S4). These changes potentially contribute to an apparent shift of the NP203 TCR mechanical load from the α chain to the β chain.

Furthermore, Figure 4C reveals two additional differences between the NP207 (T_BP_) and NP203 (T_CM_) complexes related to their respective Vα domains and the single N-glycosylation site adduct. In NP207, as well as in NP101 and NP202, the Vα C” edge strand shifts from the GFCC’ to the ABED face as commonly observed in the TCR Vα domains ^39^. In NP207, the conserved N-glycan binding residue N58α is a part of CDR2α. The first N-acetylglucosamine (GlcNAc) attached to N58α is visible in the structure, pointing away from the ABEDC” sheet of the Vα domain. In this orientation, even a mature N-linked glycan attached to N58α is unlikely to extend to the TCR-pMHC interface. In contrast, in the NP203 complex, the C” of Vα domain is absent with no secondary structural element between the C’-strand and D-strand. Instead, the extended loop from C’- to D-strands is stabilized by two side chain-main chain H-bonds between the loop and the D-strand (boxed insert in Figure 4C). Because E57α points to the D-strand, N58α pivots from the D-strand to the interface edge between the Vα C’-stand and the Vβ G-strand, repositioning the attached GlcNAc to above the α2-helix apex. The N-glycan with its additional sugar adducts attached to N58α (absent here due to EndoH cleavage) may contact the pMHC, impacting TCR ligation, mechanotransduction involving the TCR-pMHC interface, and NP203 T_CM_ polarity.

These structural distinctions should impact gene activation considering the divergence in the transcriptomes of the NP202, NP101, NP207 and NP203 clonotypes. On this point, molecular dynamics simulations under force reveal key factors that determine the mechanical response and fine peptide TCR discrimination ^40,41^. As force moves through the TCRαβ ectodomains, we posit that the four related clonotypes would manifest differential responses to the physical load on the TCRαβ-pMHC bond despite their similarity in primary amino acid sequences. To test this idea, we measure differential physical stress responses of individual clonotypes by counting the number of TCRα and TCRβ residues that form 6 or more intradomain contacts with occupancy greater than 80% during a 700ns simulation. The larger the number of these ‘hub’ residues, the more tightly packed and rigid the corresponding domain is, an indication that the domain bears greater physical load. Under identical loading conditions, the number of hub residues varied considerably among the 4 subdomains (Figure S7D). In sum, precise alterations in interface mechanochemistry (i.e., bonding impacted by and/or related to mechanical forces) appear to influence pMHC ligation as well as the resultant physical load distribution through these TCRs in functionally consequential ways.

In contrast, the two NP204 and NP105 T_CM_ clonotypes distinguish themselves from NP202, NP101, NP207 and NP203 above by their Vβ-dominant docking footprints on NP_366-374_/D^b^ (Figures 3B and 3C), and as further visualized by the comparison of NP202 versus NP105 side views (Figure 4D, left panel). The NP105 TCR-pMHC interaction is dominated by Vβ, notably involving its CDR3β motif D96RRxS100β (Figure 4D, right panel). The NP105 CDR3β D96 forms a hydrogen bond to p8T, akin to D95β or E95β found in other NP TCRs. Meanwhile, R97β forms a salt bridge to p4E, which lacks interaction with the Vα domain in contrast to the four TRBV5-containing NP TCRs (Figures 4A and 4D). R98β forms a H-bond to the carbonyl group of p5N. The side chain and amide group of S100β form a pair of H-bonds with the p7E sidechain. Furthermore, p6M is no longer sandwiched between Vα and Vβ. Instead, it is surrounded by a part of CDR3β (from R98β to S100β) and MHC residues H155 and Y156 (not shown) and capped by Y94α of CDR3α. Strikingly, the Vα contribution to peptide recognition is minimal. The primary features of the Vβ-dominant docking in NP105 are largely conserved in the other T_CM_ NP204 that utilizes distinct Vα and Vβ genes but shares the CDR3β motif with NP105 (Figure S6H) and distinct from the others characterized (Figure 4 and Figure S6H). From the collective structural analyses of these six TCRβ-pMHC complexes we conclude that regardless of whether TCR footprints map to the same or different physical regions on the pMHC interface, it is the atomistic bonding features therein and each receptor’s mechanical properties that determine how load is transferred to and then through the TCR. Because these structural differences imply distinct coupling to the CD3 signaling apparatus, we tested whether the Vα- versus Vβ-centric engagement produces distinguishable proximal phosphorylation signatures.

### Differential tyrosine phosphorylation and V-centric TCR binding

Explicitly, the distinct transcriptional programs elicited by pMHC ligation of those TCRs correlating with “Vα- vs Vβ-centric” interactions, suggest that different downstream signaling pathways should be apparent, especially considering the known compartmentalization of CD3 dimers in the holoreceptor (Figures 5A and 5B). As CD3ζ tyrosine phosphorylation is maximal at 1 minute following pMHC ligation of TCRs in triggered T cells^42^, we used this as a proxy to investigate BW5147 transductants of the NP202 versus NP105 and NP204 derived from T_EM_ and T_CM_ subpopulations, respectively. As shown in Figure 5C and Data S7, the NP202-associated CD3ζ subunit is significantly more phosphorylated following NP_366-374_/D^b^ tetramer stimulation compared to that of NP105 and NP204, as measured by anti-phospho-Tyr 142-specific antibody under non-reducing conditions required for its detection. This difference in three independent experiments is not due to diminished NP105 or NP204 signaling ability, as higher molecular weight proteins in the 100-120 kDa range are as or more heavily phosphorylated in NP105 and NP204 compared to NP202 (Figure 5C, right). Yet, within the ∼20-25 kDa range, where the CD3 subunits are found to migrate upon reduction, the NP202 results in a greater level of phosphorylation. Thus, while the T_EM_ NP202 α-dominant TCR results in higher levels of ∼20 kDa CD3 phosphorylation, the NP105 and NP204 β-dominant T_CM_ TCRs show phosphorylation of alternative signaling proteins. A similar relationship was observed between the T_EM_ NP201 and T_CM_ NP204 transductants (Figure 5D). NP201 shows higher levels of CD3ζ phosphorylation compared to the β-dominant NP204 TCR, whereas NP204 is more heavily phosphorylated in the 100-120 kDa range. Thus, the downstream signaling outcomes are woven into the structural alignment of pMHC binding to the TCR.

**Figure 5.**
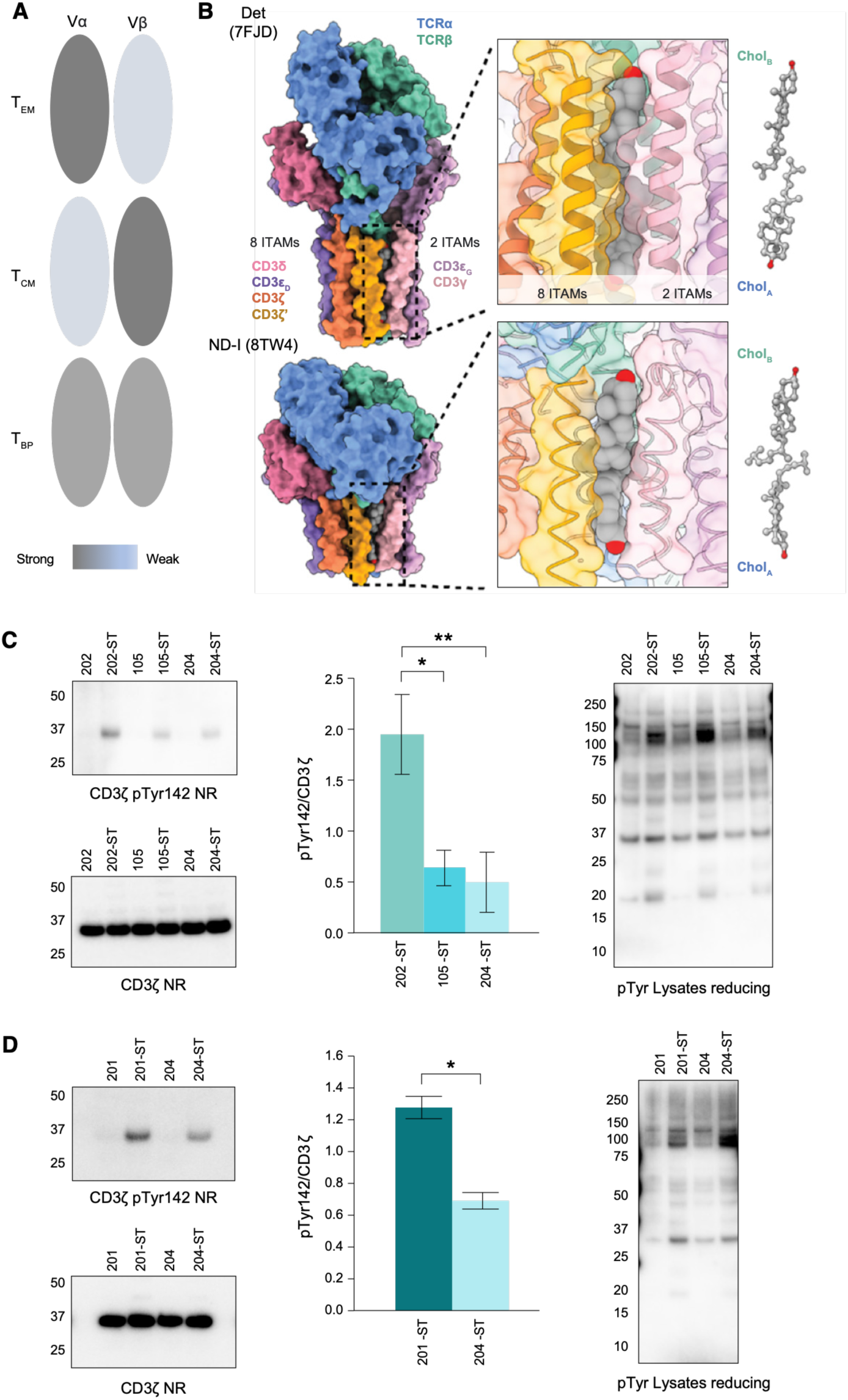
Differential tyrosine phosphorylation of CD3ζ and cellular proteins in TCR-transduced BW5147 cells derived from T_EM_ vs T_CM_ T cells. (A) Cartoon depicting polar and bipolar V domain interactions differing in their relative strength and distribution linked to indicated T memory types. (B) View of compartmentalization and sidedness of representative αβTCR holoreceptors (comprising eight subunits consisting of four dimers TCRαβ, CD3εγ, CD3εδ, and CD3ζζ) based on cryoEM structures in detergent micelle [Det (PDB ID:7FJD)] and nanodiscs [ND-Ⅰ (PDB ID:8TW4)] with TM segments. The position of two cholesterol molecules intercalated between dimeric CD3ζζ and CD3εγ signaling subunits are shown, as adapted from Notti et al. ^52^ (C) Western blot analysis of total CD3ζ (left lower blot) and CD3ζ pTyr 142 (left, upper blot) from NP202, NP105, and 204 cell lysates pre-and post-NP_366-374_ /D^b^ tetramer stimulation under non-reducing conditions (NR) required for pTyr142 antibody detection. The ST designation represents tetramer stimulated cells. The bar graph (center) displays the normalized, relative ratio of CD3ζ pTyr 142 to CD3ζ calculated to be 1.95 +/- 0.39, 0.64 +/- 0.18, and 0.50 +/- 0.29 for the NP202-ST, NP105-ST, and NP204-ST cell lines, respectively. Data are means +/-SD of three independent experiments and a calculated P value of 0.011 for NP202 vs. NP 105, and 0.008 for NP202 vs. NP204 determined by paired, two-tailed T-test using linear fitting to adjust for inter-experimental variations. Western blot analysis (right) of total pTyr for NP202, NP105, and NP204 cell lysates under reducing conditions, where the approximately 100 kDa region displays several more prominent bands in NP105-ST. (D) Western blot analysis of total CD3ζ (left, lower blot) and CD3ζ pTyr 142 (left, upper blot) from NP201, and 204 cell lysates pre-and post-NP_366-374_ /D^b^ tetramer stimulation under non-reducing conditions required for pTyr142 antibody detection. The bar graph (center) displays the normalized, relative ratio of CD3ζ pTyr 142 to CD3ζ calculated to be 1.29 +/- 0.07, and 0.69 +/- 0.05 for the NP201-ST, and NP204-ST cell lines, respectively. Data are means +/- SD of three independent experiments and a calculated P value of 0.014. Western blot analysis (right) of total pTyr for NP201 and NP204 cell lysates under reducing conditions, where the approximately 100 kDa region displays several more prominent bands in NP204-ST.

### Structural basis of heterosubtypic immunity

In parallel, the same structural framework provides an opportunity to rationalize why specific clonotypes tolerate (or fail to tolerate) naturally occurring NP substitutions. Structural analysis delineates the basis for differential TCR heterosubtypic immunity in some instances. For example, NP105 tolerates a peptide E7D viral mutation, whereas NP204 does not (Figure 2K). The weaker H-bonds between pE7 and CDR3β residues in NP204 (approximated as 3.10 Å and 3.49 Å) would have a greater impact than those between pE7 and CDR3β residues in NP105, which are shorter (2.77 Å and 2.82 Å) (Figure S6H and Table S4). Pointedly, the E7D mutation results in a 1.54Å side-chain shortening given a loss of one carbon-carbon bond. Distances between hydrogen bonding atoms would likely become too weak and break for NP204. Likewise, the tilt observed in NP101 (Figure 4A) and NP207 (Figure S6I) relative to the T_EM_ NP202, makes both T_BP_ clonotypes more crossreactive to p6M variants given that shorter sidechains such as I, L and V (Figure 2K) readily accommodate the reduction in volume of the hydrophobic pocket hosting the central p6 residue.

### Clonotypes shape transcriptomes: Rg T analysis

To further connect clonotype-based signaling to durable transcriptional programs under competitive *in vivo* conditions, we profiled bulk RNA-seq responses from adoptively transferred retrogenic T cells. Differential TCR clonotype-mediated biochemical signaling implies that resultant differential gene expression follows. This hypothesis was directly examined using RgT cells. Because high numbers of transferred T cells modulate memory phenotypes during antigen competition ^14–16^, we asked whether TCRs would continue to dictate intrinsic differentiation bias even under such challenging conditions. 100,000 Rg T cells bearing NP101 or NP105 TCRs were adoptively transferred to B6 hosts and infected with PR8 24 hrs later. At 35 dpi mLNs were isolated and Rg T cells were sorted based on congenic markers. PCA of gene expression following RNA-Seq for the 14 individual lymph node samples (7 for NP101 [C101 series], 7 for NP105 [C105 series]) delineated 3 clusters, 2 consisting only of NP101 (Cluster 1) or NP105 (Cluster 3), and an intermediate cluster (Cluster 2) of mixed clonotypes (Figure 6A). Differential expression of the top 500 genes identified 2 clear sets of genes partitioning with high probability between the NP101 and NP105 clusters (Figure 6B). Manual curation of the gene list for those reported to have T cell functionality reduced the set to 59 expressed genes segregating with high probability between the NP101 cluster and the NP105 cluster (Figure 6C). Application of this gene list to all 3 clusters (Figure 6D) identified the NP101 cluster 1 as displaying an exhaustion signature, as generally found in T_EM,_ especially in chronic infection and tumor environments ^43,44^. In contrast, the NP105 cluster expressing higher *Sell, Il7r* and *Bcl2* transcripts together with cytotoxic transcripts (*Gzma/Gzmb)* and active metabolic programming (*Myc*) as well as stem cell proliferative genes (*Placd8, Pim1*) suggested an activated T_CM_ state with effector functionality. Meanwhile, Cluster 2 with mixed NP101/NP105 samples revealed a representation of both activated T_CM_ and exhaustion signatures.

**Figure 6.**
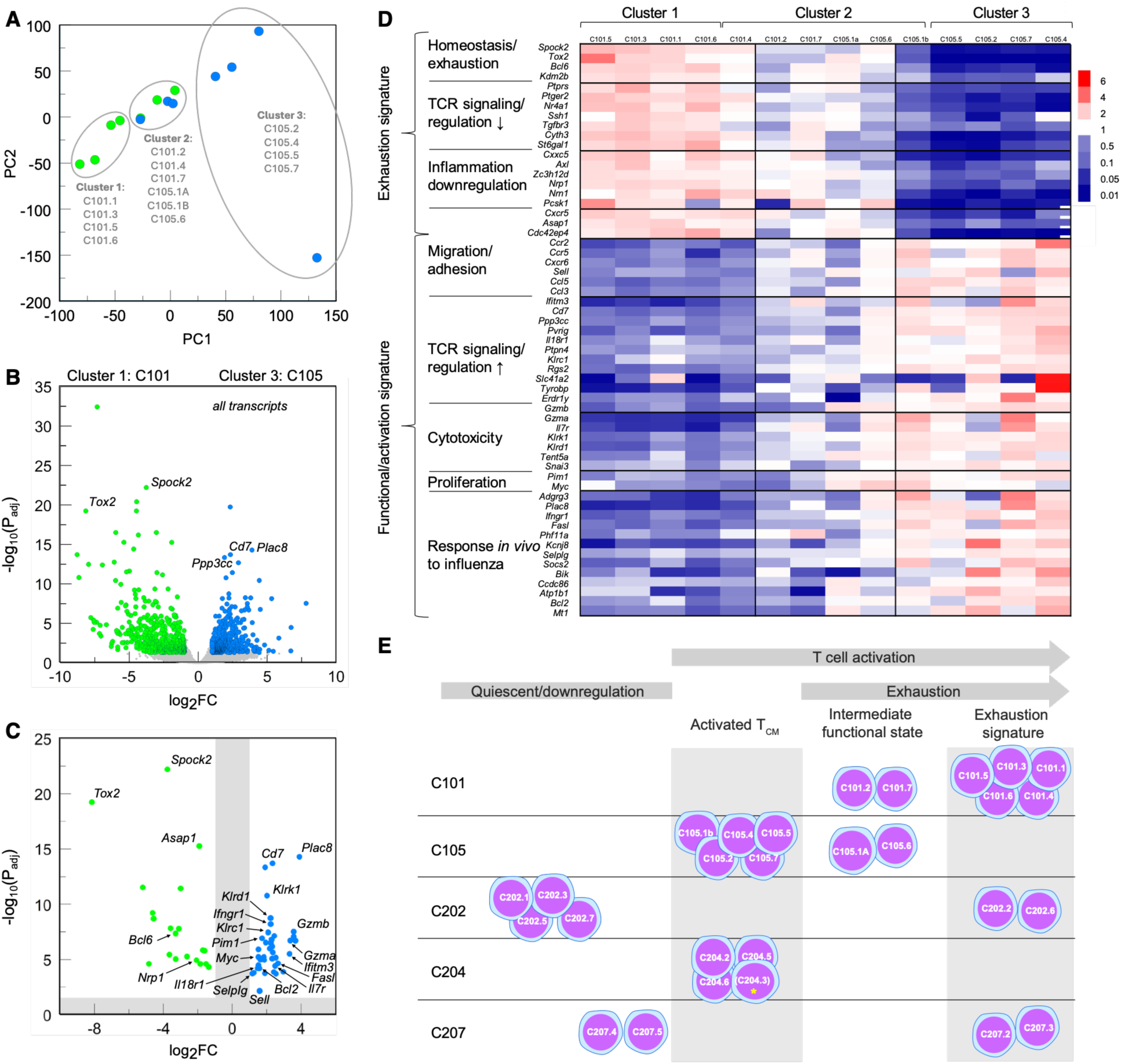
Functional characterization by transcriptome of Rg T cells isolated from mediastinal lymph nodes of IAV-infected host animals. (A) PCA for gene expression of Rg mLN-derived T cells from B6 mice adoptively transferred with either 10^5^ C101 (green) or C105 (blue) retrogenic CD8^+^ T cells expressing NP101 or NP105 TCRs, respectively, 30 d after IAV infection. Each data point represents results from an individual mouse (n = 14 mice). (B) Volcano plot of significant differential gene expression (962 genes) between Cluster 1 Rg cells (all C101, green) and Cluster 3 (all C105, blue). (C) Volcano plot of a 59 differentially expressed gene subset flagged for known T cell functionality (T cell panel) where Cluster 1 (C101.n) exhibits an “exhausted” signature and Cluster 3 (C105.n) exhibits a functionally activated T_CM_ transitioning to T_EM_ signature. (D) Hierarchical clustering/heat map of 59 gene T cell panel reclusters samples to 5 exhibiting an “exhausted” signature (all C101), 5 with activated T_CM_ signature (all C105), and a mixed C101/C105 cluster of 4 with an intermediate signature. Each row scaled where 1 (white) = base gene mean, red = fold-change above, and blue = fold-change below, the base mean. **e,** Schematic of merged C100.n [expressing NP101 and NP105 TCRs] and C200.n [expressing NP202, NP204, and NP207 TCRs] samples (see Figures S8A-S8E), shows skewed distribution of clonotypic Rg T cells by exhaustion signature, activated T_CM_ functionality, and activation state (n = 28 RNA-Seq analyses). The asterisk for C204.3 indicates quiescent signature by activation state (Figure S8C) but T_CM_ signature by differential T cell panel (Figure S8D).

To extend these findings, we performed a similar independent RNA-Seq for NP202, NP204, and NP207 Rg T cells (C202/C204/C207 series). Clustering of this dataset revealed a graded activation signature in addition to the T_CM_ and exhaustion signature (Figures S8A-S8E). Aggregation of the two data sets identified differential partitioning by cluster of the clonotypic retrogenic cells, implying this was a property of each unique NP-specific TCR (Figure 6E). By χ^2^ analysis, C101 and C105 samples partition differentially between clusters 1-3 (χ^2^ = 19.14, dof = 6, P < 0.004), and the C202, C204, and C207 dataset also partition differentially between clusters 1-4 (χ^2^ = 23.33, dof = 9, P <0.006). Notably, only NP105 and NP204 manifest as activated T_CM_ (Figure 6E). Collectively, the deep bulk RNA-Seq analyses identified TCR-linked CD8 IAV memory responses of adoptively transferred Rg T cells to manifest greater exhaustion of T_BP_ and T_EM_ counterparts relative to that of T_CM_ under adverse conditions.

Given TCRαβ clonotype-linked transcriptome differences at the T-cell memory phase, we next assessed whether distinctions would be evident following naïve Rg T cell activation within hours of stimulation, a time window that precedes overt memory differentiation and therefore reports early TCR-imposed transcriptional divergence. Examination of naïve NP202, NP204, and NP207 CD8 Rg T cells at time 0 or after 12, 36, or 66 hours of exposure *in vitro* to 1μg/ml NP_366-374_ peptide on splenocytes as APCs (Figure S8F) revealed clear differential gene expression by bulk RNA-Seq (Figures S8G-S8I), even after just 12 hours of antigen stimulation, the first timepoint assessed (Figure S8G). Notably, NP204 selectively inhibits glycolysis (*Nkg7*) (Data S8) whilst NP202 and NP207 augment glycolysis (*Hif1a* and *Snmp200*), reflecting known features of T_CM_ and T_EM_ functions, respectively ^45–48^. Since high copy numbers of NP_366-374_/D^b^ were arrayed on APCs akin to levels observed by IAV infection ^49^ for all three TCRs, the differential transcriptome outcome here supports specific TCR interaction with pMHC sculpting the T-cell developmental program, in apparent exclusion of the signal strength model 3 postulated in the introduction. Differences at 0 hours (Figure S8H) likely reflect differential clonotype-mediated homeostatic proliferation (Data S8) ^50^.

## Discussion

Our data support the conclusion that TCR-pMHC interactions directly shape CD8 memory fate, as shown by single-cell transcriptomic/paired TCR analyses in B6 mice and in TCR retrogenic chimeras. We identify structural mechanisms that appear to underlie CD8 memory fate through analysis of six distinct TCR-NP_366-374_/D^b^ complexes. The structural and molecular dynamics analyses indicate that asymmetric load transfer across the TCR may contribute to these differences, providing a plausible mechanistic basis for clonotype-linked signaling and memory bias. We interpret our data as suggesting that TCR-pMHC interaction geometry and mechanochemical organization are important determinants of memory fate. More broadly, these findings suggest that clonal diversity does not simply variegate antigen recognition; it also diversifies differentiation potential. The linkage between receptor architecture, signaling bias, and memory output may help explain how an epitope-specific response distributes cells across persistence- and effector-oriented niches. A recent non-mechanistic study ^51^ focused primarily on the acute responses against additional bacterial and viral infections distinct from IAV in the current T-memory investigation collectively support the broad conclusion that TCR-pMHC interaction is a key determinant of naïve CD8 T cell differentiation.

Two tiers of differential TCR-pMHC interactions emerged from our analyses. In the <u>first tier</u>, marked differences in docking position, specifically in the relative footprints of the Vα versus Vβ domains and in the buried surface area (BSA) contributed by each, distinguished T_EM_- and T_CM_-associated clonotypes. NP202, a T_EM_-associated TCR, was dominated by Vα contacts centered over the α1 MHC helix, whereas NP105 and NP204, both associated with T_CM_, were dominated by Vβ contacts centered over the α2 MHC helix. We term these distinct ligand-contacting topologies Vα-centric for NP202 and Vβ-centric for NP105 and NP204. This distinction appears functionally important because the αβTCR holoreceptor is itself structurally asymmetric: eight of the ten cytoplasmic tail ITAMs (CD3εδ and CD3ζζ) reside on the TCRα side, whereas only two (CD3εγ) lie on the TCRβ side. These signaling modules are further segregated by transmembrane segments separated by intercalated cholesterol molecules [^52^ and references therein] (Figures 5B and 7A). Such physical partitioning, together with asymmetric ligand engagement, likely promotes divergent signaling outputs following ITAM release (reviewed in ^29^), thereby affecting both the magnitude and quality of downstream responses. This interpretation is consistent with the bulk RNA-Seq data from naïve retrogenic T cells 12 hours after antigen exposure, with the differential gene signatures observed at one week ^51^ and one month in the current study, and with the distinct CD3ζζ tyrosine phosphorylation patterns reported here. These signaling differences may also contribute to the greater memory expansion observed for T_BP_ clonotypes and to the stronger tonic stimulation gene set found in naïve resting T_BP_ retrogenic T cells (Figure S8 and Data S8).

**Figure 7.**
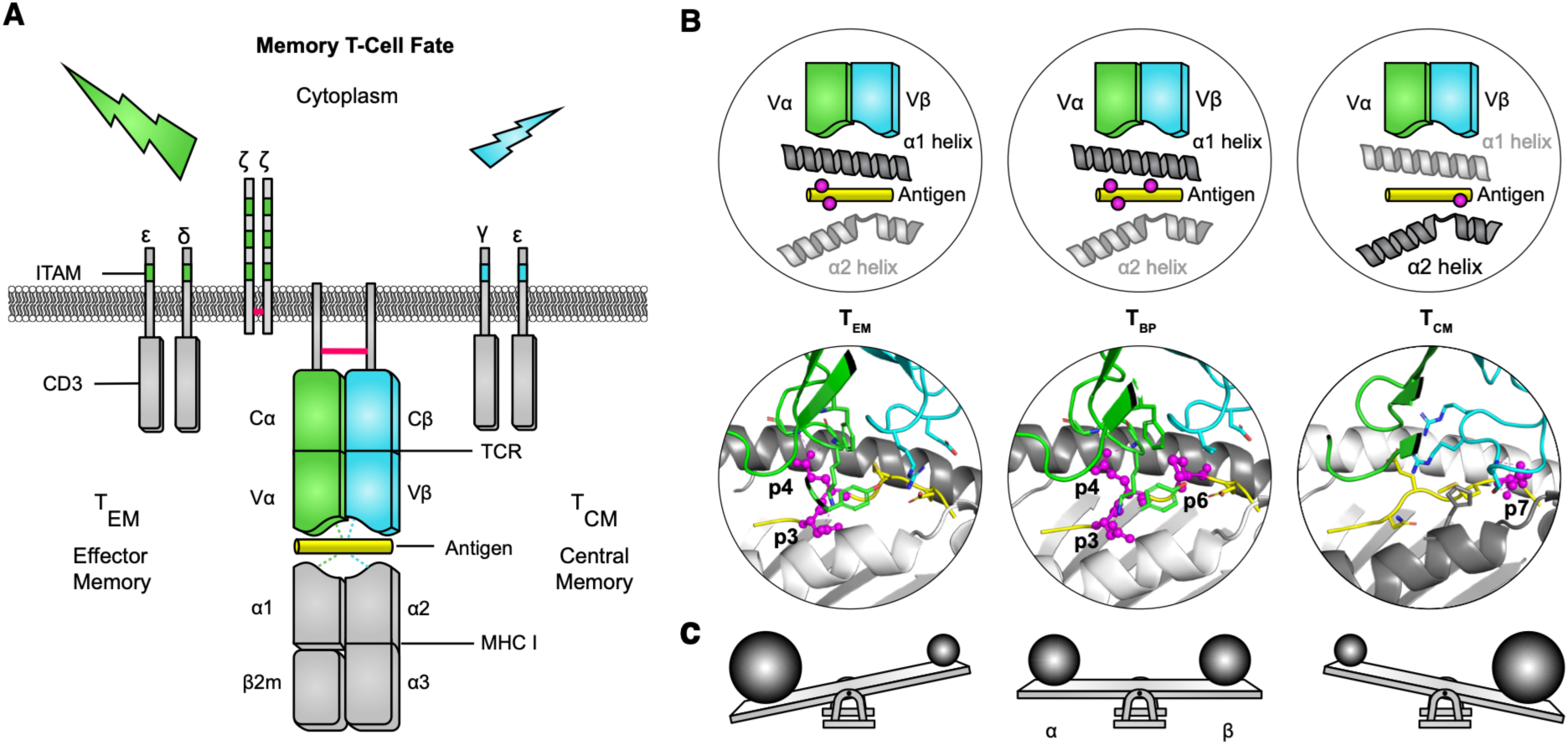
Schematic representation of memory T-cell fate and relationships to signaling balance or asymmetry. (A) Differential signaling related to TCR sidedness as a cartoon representation with CD3 elements and their ITAMs exposed to yield activation as shown. For ease of viewing ITAMs, the CD3 cytoplasmic tails are depicted on their respective sides but away from the inner membrane leaflet, rather than in their native membrane-associated positions. (B) Graphic depicting key peptide residues involved in TCR interaction illustrated as magenta balls on the peptide backbone (simple cylinder) that sits in the antigen presenting platform between the labeled MHC helices in the top row. The bottom row shows graphics based upon X-ray crystallographic structures of those residues in magenta ball and stick display with CDR3α and CDR3β (green and cyan) at the top. Both rows are based on the TCRs NP202, NP207, and NP105 from left to right. Other neighboring residues are not labeled for clarity. In both rows, the MHC helix closest to the TCR center of mass is indicated by the dark color in each complex. (C) Force on the TCRα and TCRβ subunits are balanced for T_BP_, in contrast to the indicated imbalances associated with T_EM_ and T_CM_ represented as balls of equivalent or differing sizes on a seesaw in gravitation pull to the ground. With respect to TCR mechanobiology, however, the seesaws would be flipped (rotated 180 degrees around its long axis) as the balls represent sizes of pMHC-related forces impacting the indicated TCR subunits.

The preTCR provides a potentially informative analogy. Because the preTCR lacks a Vα domain and instead uses the surrogate pTα chain paired with a canonical β chain before αβTCR development in the thymus ^53^, we propose that the preTCR pTα-β heterodimer represents an extreme form of Vβ-centric interaction linked to asymmetric mechanotransduction ^54–56^. The preTCR engages self-pMHC to support survival ^57^ and promotes β chain incorporation and persistence during thymocyte development. By analogy, Vβ-centricity in mature peripheral T cells may favor long-term T_CM_ development and renewal, that is, a more stem-like state. Consistent with this idea, five key genes involved in transcriptional programming and/or survival are upregulated both at the preTCR-dependent β-selection checkpoint and during CD8 T_CM_ generation: Tcf7 ^5,58^, Lef1 ^6,58^, Il7r ^59,60^, Id3 ^7,61^, and Bcl2 ^62,63^. By contrast, T_EM_ cells support rapid effector function but generally display more limited survival and plasticity ^1–3^.

The <u>second tier</u> of docking differences is more subtle. The overall footprints of NP202, NP101, NP207, and NP203 are largely similar, yet their atomistic-level differences manifest in functional distinctions. We suggest that NP203 is less able to transduce TCRα-associated signals for two reasons. First, Vα rigidity is reduced because NP203 lacks the C” strand. Second, the associated fully hydrated glycan adduct may intercalate between the TCR and pMHC surfaces, damping the effective physical load transmitted across the interface to the Vα domain. We therefore postulate that this reduced TCRα mechanotransduction favors T_CM_ development. In contrast, the T_BP_ clonotypes NP101 and NP207 lack these structural impediments, but both are shifted toward the Vβ side relative to the Vα-dominant NP202 TEM clonotype. We propose that the distinct pMHC binding topologies of NP101 and NP207 (Figures S7A-S7C), including their tilt and crossing angles, alter the direction of force transmission such that the resulting load is more evenly distributed between Vα and Vβ. This more balanced force vector, centered on the smaller TCR pocket surrounding p6M, may favor T_BP_ development, whereas the more Vα-dominant NP202 geometry may bias differentiation toward T_EM_.

Figure 7 summarizes these concepts. Panel A highlights the asymmetric positioning of CD3 signaling modules relative to the TCRα and TCRβ subunits. Panel B illustrates how peptide residues and MHC helices are contacted in distinct patterns across the TCR-pMHC interface, using NP202, NP207, and NP105 as examples spanning N-terminal to C-terminal peptide engagement and α1 to α2 MHC helical interactions. Panel C presents a seesaw analogy for how mechanical load may be distributed across the two TCR chains, with ball size representing relative mechanical force applied by the pMHC. As with a seesaw, unequal loading is expected to bias which signaling routes are preferentially engaged. When the load is balanced, as in T_BP_ clonotypes, stochastic outcomes may arise during repeated cycles of attachment to and detachment from antigen-presenting cells (APCs) throughout clonal expansion ^64–66^. Cell division requires such detachment, and thus variable shear forces generated at the T cell-APC interface, particularly at microvillar contact points, may influence whether progeny adopt T_EM_ or T_CM_ fates. This context-dependent flexibility is a defining feature of T_BP_ differentiation. By contrast, when the load is strongly biased, as in polar T_EM_ and T_CM_ clonotypes, the influence of stochastic variation is reduced and the cell is more likely to follow its dominant signaling trajectory. This interpretation is consistent with the known anisotropic response of the TCR to force, in which the direction of applied load influences triggering efficiency ^25^.

TCR mechanotransduction can be exquisitely sensitive: a single CDR3α amino acid difference between two T_BP_-derived TCRs (NP63 versus NP34) binding the same NP_366-374_/D^b^ ligand can determine whether signaling behaves in a more digital or analog manner ^30^. Conversely, a single amino acid substitution in a peptide ligand (i.e., an APL) presented by one MHC molecule can alter activation through the same TCR ^67^. In the latter setting, molecular dynamics simulations under load revealed that interdomain motion, molecular compliance, fluctuating forces, and interfacial contacts together define the mechanical response and thereby sharpen peptide discrimination ^40,41^. In the present study, internal stress distribution measured by molecular dynamics simulations likewise discloses divergent responses to force even among closely related TCRs (Figure S7D). More detailed simulations shall probably expose even greater differences among tier 2 TCRs than are apparent from static X-ray crystallographic structures.

Because a single epitope-specific repertoire contains many different clonotypic TCR sequences, those TCRs are likely to experience distinct patterns of stress and strain upon mechanical loading. Accordingly, load propagation pathways across the ectodomains of the αβTCR holoreceptor and into the membrane and cytoplasmic tails may also differ. These variations likely influence how the ten ITAMs within the αβTCR complex are released and engaged, thereby driving distinct downstream signaling programs, as reflected in our tyrosine phosphorylation data and transcriptomic analyses. Notably, CD3ζ Tyr-142 lies on the C-terminal side of ITAM-3, the most membrane-buried of the three CD3ζ ITAMs and the last to become phosphorylated ^68^. The greater ITAM-3 phosphorylation seen in T_EM_ relative to T_CM_ (Figure 5) suggests that Vα-centric force may preferentially perturb nearby lipids and thereby promote release of the adjacent CD3ζ tail from the membrane.

The distinction between balanced (bipolar) and polar (unbalanced) signaling has precedent in G-protein-coupled receptor (GPCR) biology where signaling through both G-protein and β-arrestin pathways, versus β-arrestin alone, can alter functional outcomes in μ-opioid and chemokine receptors (reviewed in ^69^). In GPCR systems, however, different ligands acting on the same receptor typically generate the distinct outputs. In contrast, in the T cell memory system studied here, the receptors themselves are different TCR clonotypes engaging the same pMHC ligand. Moreover, unlike most of the ∼800 GPCRs, TCRs are mechanosensors. Thus, although the systems are not identical, the conceptual parallel is biologically meaningful and not unprecedent ^70,71^.

The NP_366-374_/D^b^-specific repertoire is generated in response to a ligand expressed at ∼2000 copies per antigen-presenting cell ^49^ and is composed largely of analog TCRs ^30^. The fact that ∼20% of these TCRs are crossreactive (4 out of 16 TCRs are reactive to 5 or more NP peptide variants) offers a teleological explanation for why highly diverse clonotypes are retained within the memory niche: to preserve recognition of naturally evolving viral variants. In our dataset, T_BP_ clonotypes displayed the greatest crossreactivity, the largest clonal sizes, and rapid cytotoxicity through their T_EM_ progeny, all of which suggest substantial functional advantages. At the same time, T_CM_ clonotypes showed the least exhaustion and the greatest aggregate repertoire diversity potential for recognition of viral peptide variants, while typically existing as single-cell clonotypes occupying compact memory niche “real estate.” A similar organization is apparent in other CD8 repertoires, including PA_224-233_/D^b^, where exclusively digital performers arise because of sparse antigen display ^30^, as well as PB1_703-711_/K^b^, and likely also in CD4 repertoires directed against pMHC class II ligands.

Each TCR clonotype can be viewed as a distinct chemical entity responsive to force, that is, as a mechanophore. A repertoire directed against a single pMHC therefore consists of many mechanophores whose signaling polarity and mechanotransduction properties shape outcomes through intrinsic programs of stemness, exhaustion susceptibility, or balanced fate potential. This multidimensional information creates a compelling framework for immunotherapy design, including both engineered TCRs and synthetic receptors that must combine survival with durable effector function. Incorporating receptor-pMHC binding centricity into design principles may improve upon current AI-based receptor engineering strategies ^72–74^ and advance the clinical performance of next-generation cellular therapies.

### Limitations of the study

This study defines mechanistic links between TCR-pMHC interaction and CD8 memory fate within a single, deeply analyzed repertoire of 242 clonotypes targeting NP_366-374_/Dᵇ. However, the extent to which these principles generalize across additional antigen-specific repertoires remains to be determined. Although distance-tree analyses of independent PA_224–233_/Dᵇ- and PB1_703–711_/Kᵇ-specific repertoires support an association between long-branched clonotypes and T_CM_-like transcriptional programs, broader validation will be required. In addition, the structural analysis was limited to six TCR-NP_366–374_/Dᵇ complexes. Expanded structural sampling will be necessary to determine how broadly the interaction modes described here apply. Current AI-based methods are insufficiently accurate for reliable prediction of TCRαβ-pMHC interfaces, reinforcing the continued need for experimentally determined structures.

Our biophysical, structural, transcriptomic, and *in vivo* retrogenic datasets strongly support the mechanistic model proposed here, but they do not yet establish direct causality. Defining how force is transmitted through individual TCR mechanophores will require future atomistic molecular dynamics studies, initially spanning the TCRαβ ectodomains and ultimately extending to the full TCRαβ-CD3 holoreceptor complex, including transmembrane segments, surrounding lipids, and cytoplasmic tails. These analyses should be coupled with targeted mutagenesis of candidate pathway residues and structural elements, functional *in vitro* and *in vivo* testing of naïve CD8 T cells and memory differentiation programs expressing such variants, and detailed phosphoproteomic or ITAM-resolved signaling studies. Such future work may ultimately enable experimental manipulation of binding centricity and its associated memory bias.

Finally, tonic TCR-self-pMHC signaling may contribute to the basal state of each clonotypic T cell before foreign antigen encounter. Mechanotransduction-dependent differences in this homeostatic fitness could therefore influence chromatin state, survival, and subsequent differentiation potential. The cross-reactivity observed in T_BP_ clonotypes may extend to tonic signaling and, if so, could also contribute to this preconditioning.

## Resource availability

## Lead contact

Further information and requests for resources and reagents should be directed to and will be fulfilled by the lead contact, Ellis L. Reinherz, or Matthew J. Lang.

## Data and materials availability

Sequence files for scRNA-Seq of NP_366-374_/D^b^-specific memory CD8 T cells, non-tetramer sorted CD8 T cells at memory phase, and at recall response have been deposited in NCBI Gene Expression Omnibus (GEO) under accession GSE302680. Bulk RNA-Seq data for *in vivo* RgC mice are accession GSE302856. Bulk RNA-Seq data for *in vitro* Rg T cell culture are accession GSE313272. PDB accession code will be available at the time of publication for all X-ray crystallographic structures. PDB X-ray Structure Validation Reports are in Data S10. Materials and protocols are available on request to scientists through the corresponding authors.

## Acknowledgments

We gratefully acknowledge Dr. Jia-huai Wang for carefully reviewing the manuscript and Dr. Mikyung Kim (Dana-Farber Cancer Institute) for advice on SPR. We thank Carolyn Barbie for assistance with Figure 7 design.The use of SBC 19-ID at Argonne National Laboratory and the use of NYX 19-ID and FMX 17ID-2 at Brookhaven National Laboratory were supported by DOE contract no. DE-AC02-06CH11357 and DE-SC0012704, respectively. Monomers and tetramers listed in Data S9 were obtained through the NIH Tetramer Core Facility. This work was supported by National Institutes of Health grant 1P01AI143565 (M.J.L, R.J.M, W.H, and E.L.R) and T32 DK101003 (A.G.S). Simulations were performed by using computers at the Texas A&M High Performance Research Computing facility.

## Author contributions

A.A and K.T contributed to conceptualization, methodology, investigation, formal analysis, visualization, data curation, writing the original draft, and reviewing and editing the manuscript. R.J.M contributed to methodology, investigation, reviewing and editing the manuscript. M.A.B contributed to formal analysis, visualization, data curation, reviewing and editing the manuscript. J.S.D-C, contributed to methodology, investigation, formal analysis, visualization, data curation, reviewing and editing the manuscript. K.N.B contributed to methodology, investigation, formal analysis, visualization, reviewing and editing the manuscript. E.H.K and A.G.S contributed to methodology, investigation, formal analysis, and visualization. A.N.P contributed to methodology, formal analysis, visualization, reviewing and editing the manuscript. S.A contributed to investigation. V.C contributed to investigation and formal analysis. J.J.L, K.I.U, J.K.K, M.B, and C.M.M contributed to investigation. P.H.L contributed to data curation. M.Y.T contributed to data curation, supervision, reviewing and editing the manuscript. W.H contributed to formal analysis, Visualization, Funding acquisition, reviewing, and editing the manuscript. M.J.L contributed to conceptualization, visualization, funding acquisition, supervision, reviewing and editing the manuscript. E.L.R contributed to conceptualization, methodology, visualization, funding acquisition, supervision, writing the original draft, and reviewing and editing the manuscript.

## Competing interests

Authors declare that they have no competing interests.

## Methods

### Mice and IAV infection

C57BL/6N (B6), B6.129S6-Rag2^tm1Fwa^N12 (*Rag2*^-/-^), and B6.SJL-*Ptprc^a^*/BoyAiTac (CD45.1/1) mice were purchased from Taconic Biosciences, Inc. B6.PL-*Thy1^a^*/CyJ (Thy1.1) were obtained from The Jackson Laboratory. CD45.1/1-*Rag2*^-/-^ and CD45.1/2-*Rag2*^-/-^ were generated by intercrossing CD45.1/1 and *Rag2*^-/-^ strains. Thy1.1-*Rag2*^-/-^ mice were generated by intercrossing Thy1.1 and *Rag2*^-/-^ mice. All mice were housed and bred under specific pathogen-free conditions at the Dana-Farber Cancer Institute (DFCI) Animal Facility, accredited by the Association for Assessment and Accreditation of Laboratory Animal Care (AAALAC). Euthanasia was performed by CO_2_ inhalation followed by cervical dislocation. Mice aged 6-10 weeks were used for experiments. Sex-matched mice were used for each experiment. No gender preference was applied, and the sex in each experiment was not deliberately selected.

Mice were infected intranasally with 20 EID_50_ of Influenza A/PR/8/34 virus (PR8, H1N1, Charles River Laboratories) as a primary infection under anesthesia with intraperitoneal injection of ketamine/xylazine (120 mg/kg ketamine, 10 mg/kg xylazine). For selected experiments, PR8-infected mice were rechallenged intranasally with 1×10^7^ EID_50_ of serologically distinct X:31, A/Aichi/68 (X31, H3N2, Charles River Laboratories) >30 days after the initial infection. These viral doses were used consistently across all experiments. All mouse maintenance, breeding, and experimental procedures were approved under Dana-Farber Cancer Institute Institutional Animal Care and Use Committee (IACUC) protocol 04-113.

### Cell lines and cell culture

BW5147.3 cells and CD3δγεζ pMIY vector were a gift from the Vignali lab (St. Jude Children’s Research Hospital, Memphis, Tennessee), and mCD8αβ^+^ BW5147.3 cells were generated as previously described ^20^. The cells were maintained in D10 [DMEM medium (Gibco), 10% fetal bovine serum (FBS) (Sigma-Aldrich), 100 IU Penicillin and 100μg/mL Streptomycin (Corning), 2 mM L-glutamine (Corning) and 55 μM 2-mercaptoethanol (Gibco)] with 400 μg/mL Hygromycin B (Gibco) and 400 μg/mL Geneticin (Gibco). R8 cells were maintained in R10 (RPMI-1640 (Gibco), 10% FBS, 100 IU Penicillin and 100 μg/mL Streptomycin, and 55 μM 2-mercaptoethanol). Mouse Rg T cells were cultured in X-vivo 15 Hematopoietic Serum-Free Culture Media (LONZA) with 2% FBS (Sigma-Aldrich), 100 IU Penicillin and 100μg/mL Streptomycin (Corning), 1% Glutamax (Gibco), and 55 μM 2-mercaptoethanol (Gibco). The sorted Rg T cells from CD45.1/2 Rg mice were cultured with CD45.2/2 splenocytes as APC with 1μg/mL NP_366-374_ peptide (ASNENMETM) (United Biosystems) on 96-well plates. All cells were grown in a 5% CO_2_ incubator at 37°C.

### Cell isolation, flow cytometry, and cell sorting

Mediastinal lymph nodes (mLNs) and spleens were harvested, mashed with a 3 mL syringe on a 24-well plate with RPMI-1640, filtered with 80 μm mesh, washed with RPMI-1640. Spleen samples underwent red blood cell lysis using hemolysis buffer (140 mM NH₄Cl, 17 mM Tris-HCl, pH 7.2), followed by washing with RPMI-1640. Resident CD8^+^ T cells from lung were isolated as previously described ^30^. Briefly, mice were intravenously injected with 0.8 μg PE-Cy7-conjugated anti-CD8α mAb in 200 μL PBS 5 minutes before euthanized to distinguish CD8^+^ T cells residing in lung tissue from those in lung vasculature ^75^. Subsequently, lung blood vessels were gently perfused with 60 mL PBS through the right ventricle to wash out the residual injected antibody and then lung tissues were harvested. After mincing lungs with scissors, the chopped tissues were digested with 2 mg/mL collagenase D and 80 U/mL DNase I in HBSS at 37 °C for 1 hour with manual rotation every 10 minutes. Digested tissues were dissociated by gentleMACS Dissociator (Miltenyi Biotec), and the cells were filtered through a 70 μm cell strainer. Red blood cells were eliminated from the cell suspension by treating with hemolysis buffer. Cells were then resuspended with FACS buffer (2% FBS/0.05% NaN_3_/PBS) or appropriate buffer as indicated for the following experiments. Cell suspensions were first treated with anti-mouse CD16/CD32 mAbs in FACS buffer to block FcR binding for 10 minutes at 4°C and then stained with antibodies indicated in each figure legend in FACS buffer for 20 minutes at 4°C. For intracellular staining, cells were fixed with 4% PFA/PBS and permeabilized using permeabilization buffer following the manufacturer’s protocol (Invitrogen). After washing, cells were incubated with anti-mouse TCF-1 mAb for 30 minutes at 4 °C. For scRNA-Seq of NP_366-374_ /H-2D^b^-specific memory T cells, cell suspensions from pooled mLNs from 10 mice was stained with APC-conjugated NP_366-374_ /H-2D^b^ tetramer (MBL International Corporation) for 30 min at RT followed by staining with Zombie Aqua, and CD8β and CD44 mAb. NP_366-374_-tetramer^+^ cells were sorted after gating on Zombie Aqua^-^ CD8β^+^ CD44^+^ cells. For scRNA-Seq of non-tetramer sorted T cells at memory and recall phase, Zombie Aqua^-^ CD8β^+^ and CD44^+^ cells were sorted from pooled mLNs from four and two mice, respectively. Rg T cells for bulk RNA-Seq were isolated from individual RgC mouse mLN as Zombie⁻ CD8β⁺ cells in combination with congenic markers CD45.1, CD45.2, and Thy1.1. Naive Rg T cells for *in vitro* culture were isolated from the spleens of CD45.1/2 Rg mice (pooled from 2-5 mice, as indicated in Figure S8F). The cells were sorted as Zombie^-^ CD8β⁺ CD44^-^populations. Following *in vitro* culture, Rg T cells were harvested at the indicated time points, pooled from 8-10 wells of 96-well plates/sample, and purified for bulk RNA-Seq by sorting Zombie⁻ CD8β⁺ CD45.1⁺ cells. All antibodies and the concentration used are listed in Data S9. All cells were analyzed on a BD LSR Fortessa™ Cell Analyzer (BD Biosciences) or sorted on a FACS Aria II (BD Biosciences). Data were analyzed with FlowJo software (FlowJo, LLC).

### Single-cell RNA-Seq library construction and sequencing

NP_366-374_-specific memory CD8^+^ cells were sorted into 500 mL R10 in 1.5 mL tube. Cell density was adjusted to analyze 10,000 cells per sample. Single-cell gene expression (scRNA-Seq) was performed using Chromium Single Cell 5’ Library and Gel Bead Kit with V(D)J enrichment kit (10x Genomics). Gel Bead-in-Emulsion reverse transcription (GEM-RT) reaction, clean-up, and PCR amplification were performed to obtain cDNA for library generation. Single cell sequencing libraries were generated using 5’GEM-X v1 according to manufacturer’s protocol. Libraries were uniquely indexed for multiplexed sequencing. Libraries were single i7-indexed using the Chromium i7 Multiplex kit (indices listed in table). Following isolation and clean-up of library DNA, concentration and fragment size were assessed using an Agilent Bioanalyzer and quantification by Qubit analysis (ThermoFisher). 150bp paired-end sequencing was performed on a Hiseq 3000 utilizing the following allocation parameters per lane: two 5’GEM-X libraries (2 x 40% of reads) and two V(D)J libraries (2 x 10% of reads).

### Single-cell RNA-Seq data analysis

Gene expression and TCR FASTQ files were processed using CellRanger version 3.1.0 ^76^ using the GRCm38 mouse genome reference. The R package Seurat (version 5.1.0) ^77^ was used to filter out cells, cluster the data using 7 principal components, and perform downstream analyses. Cells were required to have less than 10% of mitochondrial content, between 5 and 7,000 features, and a minimum of 1,000 UMIs. Potential doublets were removed using the Python package “scrublet” with default parameters ^78^. Additionally, only those cell with exactly one alpha TCR chain and exactly one beta TCR chain were retained. Gene signature scores were calculated using the package AUCell (version 1.26) ^79^. The first principal component was used to generate a cell density ridge plot. UMAP approach was used for data visualization ^80^.

### Verification of memory clusters

scRNA-Seq was performed on non-tetramer sorted (NS) memory CD8 T cells from mLNs, and these were processed as described above with cluster resolution set to “0.2”. Previously defined gene sets associated with T_CM_ and T_EM_ cells ^81^ and AUCell were used to determine which clusters in the NS data constituted T_CM_ and T_EM_ clusters. Differential gene expression between NS T_EM_ clusters and T_CM_ clusters was performed using “FindMarkers” from the “Seurat” package with “min.pct” set to 0.3. Final T_EM_ and T_CM_ signatures were those with an adjusted p-value <= 0.05, and, for T_EM_, (pct.1 – pct.2) >= 0.3 and pct.2 < 0.2 and, for T_CM_, (pct.1 – pct.2) <= -0.3 and pct.1 < 0.2. The final T_EM_ gene signature includes *“Gzmk”, “Itga1”, “S100a4”, “Id2”, “S100a6”, “Mdfic”, “Rgs1”, “Fasl”, “Pdcd1”, “Ly6a”,* and *“AU020206*”. The final T_CM_ gene signature includes *“Sell”, “Satb1”, “Tmem108”, “Txk”, “Nsg2”, “S1pr1”, “Cmah”, “Pde2a”, “Ccr7”, “Treml2”, “Rras2”, “Dapl1”, “Il6st”, “Pdlim1”, “Id3”,* and *“Dennd2d”*. These gene signatures were used to verify T_CM_ and T_EM_ clusters on scRNA-Seq of NP_366-374_/D^b^-specific memory CD8 T cells using “AUCell”.

### Memory clonotype definition

T_CM_ and T_EM_ clonotypes were broadly defined as those in which more than 70% of cells localized to the polarized T_CM_ or T_EM_ cluster, respectively. T_BP_ clonotypes were those in which fewer than 70% of cells localized to either cluster. Polarized clonotypes manifest statistically distinct and non-random distributions in contrast to bipolar clonotypes.

### Clonotype origin

Clonotypes NP201-207 were isolated from pooled mLNs of 10 B6 mice at day 34 following primary infection with the PR8 strain of IAV. Clonotypes NP101 and NP105 were derived from pooled mLNs of 2 B6 mice undergoing a memory recall response, 4 days after heterologous challenge with X31. Clonotypes NP34, NP41, and NP63 were previously identified from lungs 5 days after secondary X31 infection ^30^. Clonotypes NP1-3 were isolated from the T_EM_ cluster of CD44⁺ memory CD8β⁺ T cells sorted from pooled mLNs of four B6 mice at day 33 post-infection.

### Gene enrichment analysis

Gene Set Enrichment analysis was performed on log_2_ fold changes using the “GSEA” function of the R package “clusterProfiler” (version 4.12.6) ^82^ with parameters minGSSize = 6 and pAdjustMethod = “fdr”.

### RNA velocity analysis

Intronic read counts were calculated using “velocyto” (version 0.17.17) ^83^. The Python package “scVelo” (version 0.2.5) ^84^ was used to perform velocity analysis using parameter mode set to ‘stochastic’.

### Generation of TCR sequence tree and analysis

Vα and Vβ sequence similarities (TCR distances) were calculated using the package “TCRDist3” (version 0.2.2) ^85^ with the distance matrices for the two summed. Clonotypes with the TRBV gene “TRBV12-2+TRBV13-2” were excluded from the analysis due to the absence of this V-gene from the TCRDist3 database. A distance cladogram was generated using the R package “hclust” using complete linkage. The R package “pheatmap” (version 1.0.12) was used to create a heatmap of TCR distances.

### Definition of short and long branch

Short and long branches of the TCR tree are defined based on pairwise TCR sequence distance metrics. First, the mean of the top 10 shortest pairwise distances was calculated for each clonotype. Then the threshold value was determined based on the bimodal distribution of the mean distances for all clonotypes (mean distance = 109.0, Figure S3D). This threshold value was used to define ‘short’ and ‘long’ branches. For example, CT_25, categorized as a short-branch clonotype, had a mean top 10 distance of 7.8 < 109.0, whereas CT_50, categorized as a long-branch clonotype, had a corresponding value of 196.4 > 109.0.

### Entropy analysis

Shannon entropy of V and J gene usage was calculated using the “Entropy” function of R package DescTools (version 0.99.58). Entropy values were normalized by calculating the maximum possible entropy for given numbers of V and J genes and independent clonotypes using a custom script. The numbers of possible V and J genes were retrieved from CellRanger 3.1.0 VDJ database.

### Retrovirus production and transduction

To generate non-fluorescent TCRαβ-expressing vectors, the IRES-EGFP site was cut out from retroviral vector pMSCV-IRES-GFP Ⅱ (pMIG Ⅱ, Addgene, #52107) with restriction enzymes and replaced with synthesized DNA encoding TCRβ-P2A-TCRα. The vector was transfected into Plat-E packaging cells using the calcium phosphate precipitation method. A transfection mixture was prepared by combining 35 μg of plasmid DNA, 125 μl of 2.5 M CaCl₂, and 1090 μl of 0.1× TE buffer. After incubation at room temperature for 15 minutes, 1250 μl of 2× HBS buffer (100 mM HEPES, 281 mM NaCl, 1.5 mM Na₂HPO₄) was added dropwise to the DNA/CaCl₂ solution while vortexing. The resulting DNA/calcium phosphate precipitate was immediately added to Plat-E cells. Retrovirus in the cultured supernatant was collected 48 hours post-transfection and frozen at -80°C until use. Thawed retrovirus was transduced with Retronectin (Takara Bio) using Retronectin-bound virus infection methods according to the manufacturer’s protocol. Retrovirus was transduced into mCD8αβ^+^ BW5147.3 cells to generate TCRαβ-expressing cell lines or into mouse bone marrow (BM) cells to generate Rg mice. To establish TCRαβ-expressing BW cell lines, the transduced cells were sorted by FACS Aria Ⅱ (BD Biosciences) to match the TCR expression based on the cell surface expression of CD3ε. Before all assays were performed, CD3ε or TCRβ surface level was confirmed to be matched in a group.

### Generation of Rg mice

Rg mice were generated as previously described ^30,86^. Briefly, BM cells were harvested from CD45.1/1-*Rag2*^-/-^, CD45.1/2-*Rag2*^-/-^, or Thy1.1-*Rag2*^-/-^ mice as indicated in figure legends, and hematopoietic stem cells (HSC) were enriched by EasySep Mouse Hematopoietic Progenitor Cell Isolation Kit (STEMCELL Technologies) followed by expansion in Stem cell medium (StemPro™-34 SFM (Gibco), 5% FBS, 100 IU Penicillin and 100 μg/mL Streptomycin (Corning), 2 mM L-glutamine, with 50 ng/mL mIL-3 (STEMCELL Technologies), 50 ng/mL hIL-6 (STEMCELL Technologies) and 50 ng/mL mSCF (STEMCELL Technologies)) for 3 days. Subsequently, HSC were transduced with retrovirus encoding TCRαβ, cultured in Stem cell medium for 3 days, and then transferred into sublethally irradiated (600 rad) *Rag2*^-/-^ mice using a Gamma Cell 40 Cs^137^ Irradiator (Thratonics) one day before. Rg mouse blood was analyzed for CD8 development and TCRβ expression by flow cytometry 6 weeks after transplantation, and mice expressing more than 10% of CD8 T cells with comparable surface expression levels of the introduced TCRs in CD8^+^ cells were used for the generation of RgC mice.

### Generation of RgC mice

RgC mice were generated by adoptive transfer of naïve CD8β⁺CD44⁻ Rg T cells, isolated from pooled peripheral lymph nodes and spleens of Rg donor mice, into B6 (CD45.2/2, Thy1.2) recipient mice. The number of transferred cells ranged from 0.5 to 10 × 10⁴ per mouse, as indicated in figure legends. To avoid potential immunogenicity associated with fluorescent proteins such as EGFP ^87^, we used congenic surface markers, such as CD45.1/1, CD45.1/2, and Thy1.1, as identifiers for donor cells, allowing for reliable tracking of individual clones. Mixed RgC mice were generated by co-transferring equal numbers of Rg T cells expressing different TCRs to minimize variability across individual mice. The equal input ratios of each clone were verified by flow cytometry using congenic marker expression. Single RgC mice received Rg T cells expressing a single TCR to eliminate interclonal competition amongst different Rg T cell clonotypes. All RgC mice were intranasally infected with PR8 one day after adoptive transfer. At >30 days post-infection, transferred Rg T cells were analyzed as memory T cells by flow cytometry and RNA-Seq to assess phenotype and gene expression profiles.

### TCR Ectodomain Constructs for X-ray Crystallography

The design and expression of TCRαβ heterodimers has been described previously ^21,30,88,89^. Briefly, DNA vectors containing TCRα or TCRβ (i.e. V(D)J-C) ectodomains, a HRV3C protease cleavage site ^90^, a base or acid leucine zipper (LZ) motif to facilitate heterodimer formation upon expression ^91^, and a stop codon, were designed utilizing human cytomegalovirus promoter. TCRα and β vectors were constructed as separate eukaryotic expression vectors using the pGDom vector from prior work with TCRβ expression in mammalian cells ^92,93^, retaining the vector signal sequence and encoding the murine Cα-HRV3C-basic-LZ or Cβ-HRV3C-acid-LZ, named as pGDomCα-LZ or pGDomCβ-LZ, respectively. A novel cloning region was inserted between the signal sequence and the start of the C domain utilizing two BsaI type IIS restriction endonuclease sites. This allowed in-frame insertion, using Golden Gate Cloning strategies ^94^, of fragments encoding only VJ-α or VDJ-β. These were isolated by PCR amplification from plasmids of full-length constructs used in retroviral transductions or purchased as double stranded DNA fragments (Azenta) and subsequently ligated into the respective pGDomCα-LZ or pGDomCβ-LZ vector. TCR clonotypes with insufficient protein yields were alternately cloned into similar vectors encoding human C-domains along with the addition of a non-native disulfide between the Cα and Cβ, resulting in a chimeric construct ^95–97^. Annotated DNA and translated amino acid sequences of all constructs are included in a Data S6.

### TCR Protein Expression and Purification

The plasmids containing TCRα and TCRβ constructs were transfected together into a culture containing 2×10^6^ cells/mL Expi293F glucoseaminotransferase deficient (GnTi^-^) cells (ThermoFisher) in fresh Expi293 Expression Medium (ThermoFisher) at a final concentration of 0.5 µg vector/mL culture each, using ExpiFectamine transfection reagent (ThermoFisher) and following the manufacturer’s protocol. The Expi293F GnTi^-^ variant produces only high mannose-modified N-linked glycans, allowing the removal of glycans under native conditions using endoglycosidase H (EndoH) ^93,98^. Transfected cells were cultured at 37°C with rotary shaking at 125 rpm, per manufacturer recommendation, for 7 days for expression of protein. The supernatant was then collected and clarified by centrifugation and filtration before being passed at least twice over protein G resin (GammaBind Pl; Cytiva) coupled with the anti-LZ mAb 2H11, utilizing a gravity flow apparatus at a flow rate of approximately 1 mL/min or slower ^89^. Following washing with 10 column volumes of TBS pH 8.0, bound protein was eluted with 50 mM Citrate, 500 mM NaCl buffer, pH 2.9 (adjusted with Tris base), in six 9 mL fractions, and neutralized immediately upon collection with 1 mL (i.e., 1/10 vol) 1M Tris-HCl, pH 8.0. The fractions containing eluted protein were concentrated and buffer exchanged into TBS and purified via size exclusion chromatography (SEC) utilizing a HiLoad 16/600 Superdex 200 pg column (S200P; Cytiva). Fractions containing the TCRαβ heterodimer were concentrated to 1 mg/mL, calculated via absorbance at 280 nm (A_280_), prior to digestion with HRV3C protease (1 µg enzyme/50 µg TCR) at 25°C for 2 hours for complete removal of the LZ. Glycans were then digested with EndoH at 1U enzyme/1 μg TCR with incubation at 25°C for up to 24 hours. Completeness of the digestions was confirmed with SDS-PAGE. The fully digested protein was then purified with anion-exchange chromatography utilizing a MonoQ 5/50 column (MonoQ; Cytiva) and eluted as a single peak. Final protein purity was assessed by non-reducing (NR) and reducing SDS-PAGE prior to crystallization trials. For crystallization screening, the purified protein was buffer exchanged into 10mM Tris-HCl, pH 8.0, 100 mM NaCl and concentrated to 10 mg/mL, per A_280_ measurements.

For SM use, all constructs were the murine C domain variants with the single C-terminal interchain disulfide only. Expression and purification mirrored that for structural work with the following exceptions. Expression utilized the wildtype Expi293F cells (ThermoFisher), retaining complex glycan modifications. Following elution from 2H11 affinity columns, the proteins were buffer exchanged into PBS pH 7.4 using SEC chromatography with a 21 mL analytical-scale S200 column (Cytiva). The sample was then concentrated to approximately 1 mg/mL concentration and flash frozen with LZ regions intact for SM analysis.

TCR production for SPR was similar to the above procedures. C domain murine constructs were transfected into Expi293 Pro cells (ThermoFisher) adjusted to 5×10^6^ cells/mL using the accompanying Expi293 Pro expression medium. Cells were transfected with plasmids encoding for the TCRα and TCRβ construct using the Expi293 Pro transfection kit, following the manufacturer’s protocol with a few exceptions. 24 hours after transfection, the cells were placed in an incubator at 30°C and the supernatant was collected and clarified by centrifugation and filtered after 5 days. Subsequently, the protein was purified, and the LZ was removed as previously described, conserving the complex glycans present. All TCRs were buffer exchanged into PBS (pH 7.4) and concentrated to 500μM via centrifugal concentration.

### pMHC expression, refolding and complex formation, and purification

NP_366-374_ (NP366) peptide (ASNENMETM) was chemically synthesized (United Biosystems, Inc) and NP366/H-2D^b^ production was executed similarly to that previously described ^55,99,100^. Plasmids containing H-2D^b^ or murine β2m were transformed into BL21-DE3 *E.Coli* for expression. Transformed cells were grown to an OD_600_ of 0.6-0.8 in Luria broth (LB) at 37°C under agitation at which point expression was induced with the addition of 0.5mM IPTG. After 3 hours, cells were harvested by centrifugation, supernatant was discarded and the cell pellet was resuspended in 30 mL of TBS, pH 8.0, 1 mg/mL lysozyme was added, and the resuspension was frozen at -80°C for at least 2 hr. The lysate was then thawed and 0.3 mg/mL of DNAse and EDTA free protease inhibitor (Roche) were added before being frozen again at -80°C > 2 hr or overnight. The lysate was thawed, and inclusion bodies were purified via centrifugation and resuspension cycles, washing twice with TBS, twice with TBS + 0.1% TritonX-100, and once with TBS. The final product was a pure inclusion body pellet, which was dissolved in 10 mL of 5.4M Guanidine HCl, 0.1M Tris, pH 8.0. The solution was centrifuged and the supernatant was collected before the addition of dithiothreitol to 2 mM and storage at -80°C. H-2D^b^, β2M, and NP366 peptide were added to 300 mL 20 mM Tris, 8M Urea at a 60:20:10 w/w ratio before serial dialysis (6-8 kDa nominal molecular weight cutoff; Spectrum Laboratories, Inc.) against 4 L of 20 mM Tris + 2 M, 1M, 0.5M and 0M Urea, with each step greater than 2 h or overnight for a total of at least 48 hours dialysis/refold time. The dialyzed refolded protein was then concentrated for purification via SEC (S200P; Cytiva) followed by anion-exchange chromatography (MonoQ; Cytiva). The final purification yielded a mono-dispersed peak and subsequent NR SDS-PAGE revealed two bands, corresponding to H-2D^b^ and β2M. For crystallization screening, the purified protein was buffer exchanged into 10 mM Tris-HCl, pH 8.0, 100 mM NaCl and concentrated to 10 mg/mL, per A_280_ measurements.

### SM assay

Tether geometry and optical tweezers measurements were carried out essentially as detailed in ^21^, and as shown schematically in Figure (Figure S4A). Streptavidin-coated polystyrene beads (1.29 μm diameter, Spherotech Inc) were connected to biotin terminated 1010-bp double-stranded DNA containing covalently bound 2H11 antibody that binds to the TCRαβ construct paired LZ region. Incubation time and DNA concentration were varied to control tether density around the bead, which was optimized for the binding probability of each individual TCRαβ-pMHC pair. After incubation at RT with DNA tethers, DNA-labeled beads were washed then resuspended in 15 μL of TCRαβ (∼0.1 mg/mL) and incubated at RT for an additional 1 hr. The tethered beads were then washed a final time before resuspension in PBS buffer.

A 10-μL flow cell was prepared using two pieces of double-sided sticky tape. Cover-glass surfaces were coated with functionalized PEG silane 99% mixed with 1% biotin-PEG silane like previously described work ^21^. Casein blocker (0.1 mg/mL) was introduced to the flow cell for 10 minutes then washed out with PBS buffer. Streptavidin (0.05 mg/mL) was introduced into the flow cell, incubated for 30 min, and washed with PBS buffer. Biotinylated pMHC (∼0.05 mg/mL, CD8-binding site-mutated monomer, NIH tetramer core facility) was introduced and incubated for 30 min and washed again with PBS buffer. Finally, tethered bead solution was introduced to the flow cell for bond lifetime measurements.

Tethers were formed by scanning a bead near the cover-glass surface using 300 nm steps of the piezo stage in the x-y plane, pausing for several seconds between stage movements to allow for TCRαβ-pMHC bond formation (Figure S4B). Successful tether formation was indicated by bead displacement relative to the trap center. After rupture, beads were mapped for position sensing using automated procedures followed by stiffness calibration. Control experiments where TCRαβ was omitted from the assay resulted in minimal, atypical tether formation. Binding probability (Figure 2H and Figure S6F) can be assessed by comparing the number of stage movements which resulted in a binding event to the total number of stage movements. This is performed as a group for a given panel under identical conditions of tethers per bead and surface ligand density.

### Surface Plasmon Resonance (SPR)

SPR experiments were performed at 25°C using a Biacore 1K (Cytiva) with PBS, pH 7.4 buffer. ∼3000 RU of streptavidin (New England Biolabs) was fixed to a CM5 chip in both the reference and experiment flow cell using an amine coupling kit (Cytiva), following the manufacturer’s procedure. 300 RU of biotinylated WT-NP_366-374_/D^b^ monomer (NIH tetramer facility) was bound to the streptavidin coupled experiment flow cell. Kinetic and steady state parameters were determined by flowing threefold serial dilutions ranging from 33.33 to 1.37μM for NP207 and NP105 or 100 to 1.37μM for NP202 at a rate of 30 μL/minute with contact and disassociation times of 180 seconds. Running buffer was used for regeneration. Each experiment was repeated in duplicate and was analyzed with Biacore Insight Evaluation Software (version 5.0.18.22102) using 1:1 binding model.

### Peptide stimulation and IL-2 measurement

1×10^5^ NP-specific TCR-transduced BW cells were cultured in D10 with 1μg/mL WT-NP_366-374_, 1μg/mL NP_366-374_ peptide variants, or titrated theses peptides (from 1×10^-5^ to 1×10^4^ ng/mL) at 37°C for 16-18 hours overnight. 1×10^5^ R8 cells used as APC were treated with Mitomycin C (Sigma-Aldrich) and washed with PBS three times prior to use. IL-2 concentration in culture supernatant was measured by ELISA assay according to the manufacturer’s instruction. EC_50_ of peptide response to each BW cells was calculated by Prism 10 (GraphPad Software).

### Tetramer binding assay

5×10^5^ NP-TCR transduced-BW cells were plated on 96-well plates, and 10μg/ml WT- or CD8 binding site-point mutant (D227K) NP_366-374_/H-2D^b^ tetramer (NIH tetramer core facility) in FACS buffer was added. The cells were incubated at RT for 30 minutes, washed with FACS buffer twice, and the fluorescence intensity was analyzed by flow cytometry. The tetramers used in the experiment are listed in Data S9.

### Tetramer activation assay and pTyr analysis

Because pMHC tetramers crosslink TCRs that are associated with the cytoskeleton, bioforces in the pN range are applied and thereby facilitate physical load-driven T cell activation. Detection of CD3 subunits, pTyr, and CD3ζ pTyr142 pre- and post-tetramer activation was carried out using 5 x 10^6^ cells/ml in DMEM. WT NP_366-374_/D^b^ tetramer (0.75 μg/ml final concentration) was added to the NP202 or NP105 cell samples matched for cell-surface TCR level expression prior to use. Control samples were simultaneously prepared for each cell line without the addition of tetramer at time zero. All cell samples were incubated on ice for 20 min and then washed with DMEM to remove unbound tetramer. The cells were resuspended at a final concentration of 2 x 10^6^ cells per 0.1 mL and incubated at 37◦C for 1 min. Then, 1 mL of ice-cold PBS was added to the cells and the cells were pelleted at 1200 rpm for 5 min. The PBS wash was removed and lysis buffer (1% Triton X-100, 0.1% SDS, 50 mM Tris (pH 7.4), 150 mM NaCl, 2 mM NaVO3, 1 mM N-ethylmaleimide, and Roche cØmplete protease cocktail) was added to the cell pellet. The resuspended samples were immediately placed on dry ice to await further processing.

For analysis of CD3 and phosphorylated proteins, cell samples were then thawed for 10 min on ice and centrifuged at 13,000 rpm, 4◦C for 15 min. The clarified lysate was transferred to a clean tube. Aliquots were separated non-reducing with 4-12% bis-tris NuPAGE gels, transferred to polyvinylidene difluoride (PVDF) membrane, and detected with anti-CD3ζ pTyr142 (clone EM-54), and anti-CD3ζ (clone H146-968). The western blots were imaged using a Bio-Rad ChemiDoc imaging system. Band density was measured using the Image Lab software, with band densities normalized to the median within each experimental set to mitigate antibody staining artifacts The level of phosphorylated CD3ζ Tyr142 was determined in proportion to CD3ζ for each sample. Three biological replicates were prepared for each activation assay with tetramer activation being carried out on three separate days with three separate cell culture preparations. To examine overall phosphotyrosine levels, the cell lysates were separated on a reducing gel, transferred to PVDF membrane, and assayed for pTyr (clone 4G10).

### Bulk RNA-Seq analyses

For transcriptome analysis of RgC CD8^+^ T cells responding to influenza virus *in vivo*, lymph node cells of retrogenic origin (see table below), cells were sorted into 1.5 mL tubes containing TCL buffer x2 (350 µL; Qiagen) supplemented with 2-mercaptoethanol (1%).

Gating parameters and cell yields for retrogenic cells:

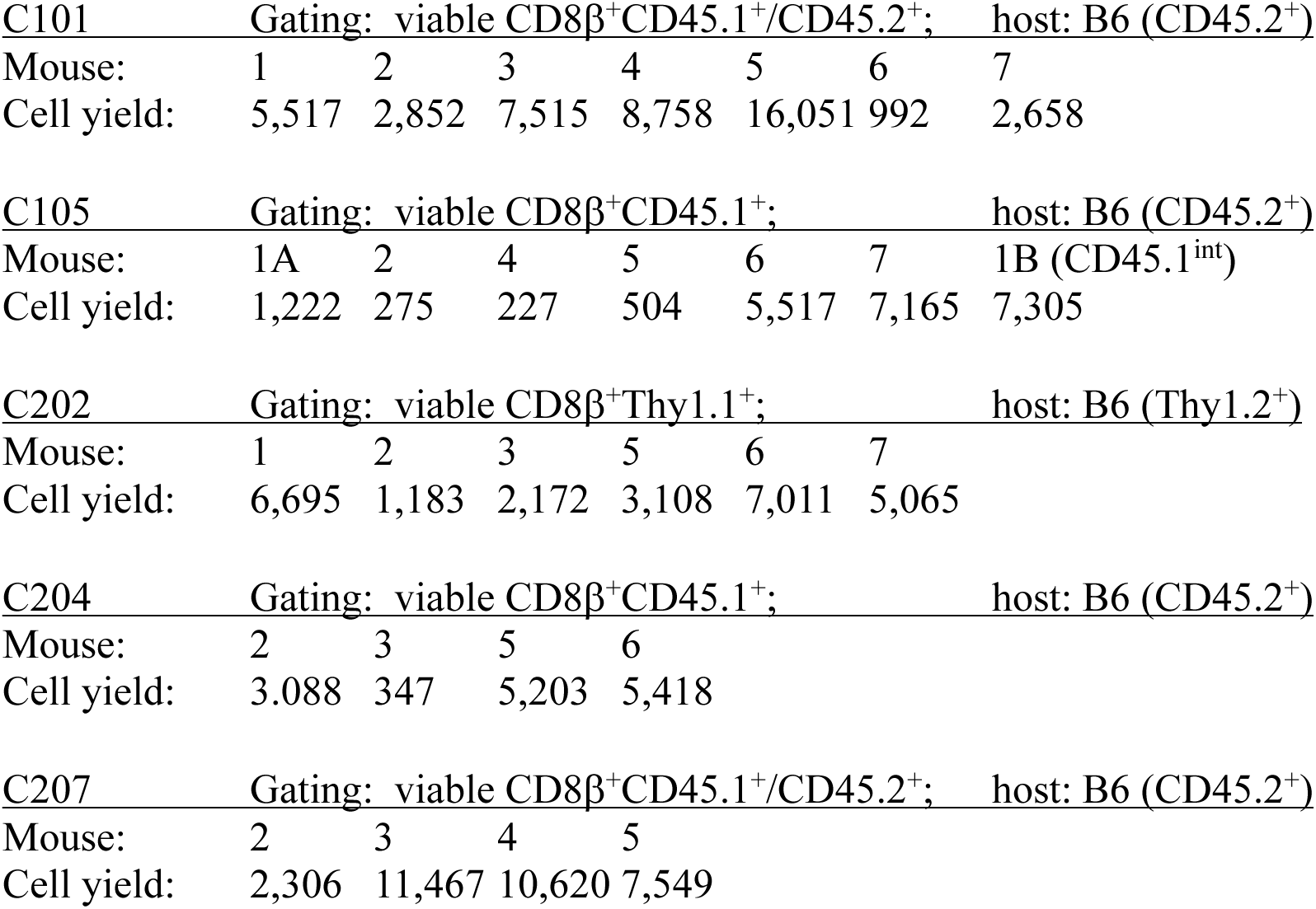

| C101 | Gating: viable CD8β<sup>+</sup>CD45.1<sup>+</sup>/CD45.2<sup>+</sup>; host: B6 (CD45.2<sup>+</sup>) | | | | | | |
| --- | --- | --- | --- | --- | --- | --- | --- |
| Mouse: | 1 | 2 | 3 | 4 | 5 | 6 | 7 |
| Cell yield: | 5,517 | 2,852 | 7,515 | 8,758 | 16,051 | 992 | 2,658 |

| C105 | Gating: viable CD8β<sup>+</sup>CD45.1<sup>+</sup>; host: B6 (CD45.2<sup>+</sup>) | | | | | | |
| --- | --- | --- | --- | --- | --- | --- | --- |
| Mouse: | 1A | 2 | 4 | 5 | 6 | 7 | 1B (CD45.1<sup>int</sup>) |
| Cell yield: | 1,222 | 275 | 227 | 504 | 5,517 | 7,165 | 7,305 |

| C202 | Gating: viable CD8β<sup>+</sup>Thy1.1<sup>+</sup>; host: B6 (Thy1.2<sup>+</sup>) | | | | | | |
| --- | --- | --- | --- | --- | --- | --- | --- |
| Mouse: | 1 | 2 | 3 | 5 | 6 | 7 |  |
| Cell yield: | 6,695 | 1,183 | 2,172 | 3,108 | 7,011 | 5,065 |  |

| C204 | Gating: viable CD8β<sup>+</sup>CD45.1<sup>+</sup>; host: B6 (CD45.2<sup>+</sup>) | | | | | | |
| --- | --- | --- | --- | --- | --- | --- | --- |
| Mouse: | 2 | 3 | 5 | 6 |  |  |  |
| Cell yield: | 3,088 | 347 | 5,203 | 5,418 |  |  |  |

| C207 | Gating: viable CD8β<sup>+</sup>CD45.1<sup>+</sup>/CD45.2<sup>+</sup>; host: B6 (CD45.2<sup>+</sup>) | | | | | | |
| --- | --- | --- | --- | --- | --- | --- | --- |
| Mouse: | 2 | 3 | 4 | 5 |  |  |  |
| Cell yield: | 2,306 | 11,467 | 10,620 | 7,549 |  |  |  |

For transcriptome analysis of naïve Rg CD8^+^ cells responding to NP_366-374_ pMHC *in vitro*, cell cultures were initiated as described above and cells were harvested at 0 h, 12 h, 36 h, and 66 h. Cells were sorted (Data S8) into 1.5 mL tubes containing TCL buffer x2 (350 µL; Qiagen) supplemented with 2-mercaptoethanol (1%).

For all sorted cell samples, total RNA was extracted from each sample using the Arcturus Pico Pure RNA kit (ThermoFisher) following the manufacturer’s protocol including on-column RNase-free DNase treatment (27 Kunitz units/sample; Qiagen), and eluting into 11 µL elution buffer, snap freezing at -80°C and storage at the same temperature until library preparation and sequencing (Admera Health, LLC). Following initial RNA preparation quality control, libraries were prepared using the SMART-Seq v4 Ultra Low Input kit with Tagmentation (Takara Bio USA, Inc.), Illumina adapters added, and sequenced on the NovaSeq X Plus platform (2 x 250 bp paired end sequencing, 40 x 10^6^ reads/sample). The paired output fasta files for each sample were checked for quality using FastQC, and adapters trimmed using FastqMcf and Cutadapt ^101^. Contaminating non-polyA-tailed RNA sequence, mitochondrial genome sequences, ribosomal RNAs and transfer RNAs were then removed using Bowtie2 ^102^ and the resulting filtered reads aligned to the mouse reference genome assembly GRCm39 using STAR (v2.7.3a) ^103^. HTSeq-count was then used to count the reads aligned to each gene. Gene expression in transcripts per million (tpm) normalizing for transcript length and library size, was obtained using the DESeq2 package under Bioconductor version 3.21 running on the R version 4.5 platform. All further processing for hierarchical clustering and heat maps, and differential gene expression (DESeq2), was performed on the same platform. Note that for Data S8E, the heat maps represent transcripts per million (tpm) for a reduced panel of transcripts representing genes validated as upregulated on T cell activation. In this instance, tpm can be used as a broad measure for comparison of the two data sets (C101/C105 and C202/C204/C207) since all samples were of similar origin (mLN CD8^+^ retrogenic T cells), and identical RNA preparation, sequencing, and quality control statistics (e.g. mitochondrial transcript representation) ^104^. For the NP202, NP204, NP207 Rg cells developing *in vitro*, this consideration does not apply as all samples were sorted and total RNAs were purified simultaneously. All data was organized/plotted using MS-Excel or plotted using PSI-Plot version 9 (Poly Software International).

### Crystallization

Each NP TCR was mixed with NP366-H-2D^b^ (NP_366-374_/D^b^) in a 1:1 molar ratio. The mixed TCR-pMHC solution was concentrated to about 10-20mg/ml. Sitting drop vapor diffusion technique in 96-well CrystalQuick plates (Greiner) was used for crystallization trials with the help of a Mosquito nanoliter liquid handler (TTP LabTech) at 16^◦^C. For each condition, 0.4 mL of crystallization screening formulation was added to 0.4 ml of concentrated TCR-pMHC solution; the mixture was equilibrated against 140 mL of the same screening buffer in each reservoir well. The crystallization screens used were MCSG-1-4 (Anatrace), Peg/Ion (Hampton Research) and Top96 (Molecular Dimensions). Crystals appeared under the conditions 1-8 as listed for each construct:

1. NP101-NP366/H-2D^b^ (mouse/human hybrid TCR) = NP101-NP_366-374_/D^b^ 0.2 M Ammonium iodide pH 6.2, 20% (w/v) PEG 3,350.
2. NP105-NP366/H-2D^b^ (mouse/human hybrid TCR) = NP105-NP_366-374_/D^b^ 0.1 M Sodium citrate, 5% (v/v) 2-Propanol, 20% (w/v) PEG 4,000,
3. NP202 apo form (mouse TCR) 0.2 M Ammonium nitrate pH 6.2, 20% (w/v) PEG 3,350.
4. NP202-NP366/H-2D^b^ (mouse TCR) = NP202-NP_366-374_/D^b^ 0.1 M Tris hydrochloride pH 8.5, 8% (w/v) PEG 8,000.
5. NP203-NP366/H-2D^b^ (mouse/human hybrid TCR) = NP203-NP_366-374_/D^b^ 0.2 M Ammonium fluoride pH 6.2, 20% (w/v) PEG 3,350.
6. NP204-NP366/H-2D^b^ (mouse TCR) = NP204-NP_366-374_/D^b^ 1% (w/v) Tryptone, 0.05 M HEPES sodium pH 7.0, 12% (w/v) PEG 3,350.
7. NP207 apo form (mouse TCR) 0.2 M Potassium Nitrate: HCl pH 6.9, 20% (w/v) PEG 3,350.
8. NP207-NP366/H-2D^b^ (mouse/human hybrid TCR) = NP207-NP_366-374_/D^b^ 0.6 M Sodium Chloride, 0.1 M MES:NaOH pH 6.5, 20% (w/v) PEG 4,000.

### **X-** ray diffraction data collection and processing

All crystals were harvested and treated with a cryoprotectant solution (25% glycerol or 25% ethylene glycol in mother liquor) and then flash-frozen in liquid nitrogen before X-ray diffraction data collection. Those data were collected at 100 K from the cryocooled crystals at the 19-ID at the 19-ID (SBC) beamline of the Advanced Photon Source at Argonne National Laboratory or 19-ID (NYX) beamline or 17-ID-2 (FMX) beamline of National Synchrotron Light Source II at the Brookhaven National Laboratory. The intensities of each data set were integrated, scaled, and merged with the HKL-3000 program suite ^105^. The structure of NP202-NP366-H-2D^b^ (NP202-NP_366-374_/D^b^) was first determined using the molecular replacement (MR) method ^106^, in which the murine P14 TCR and the H-2D^b^ from their complex (PDB code: 5M01) were used as search templates (Table S2). The following structural determination of NP TCR apo forms and other complexes were performed with the same MR method using NP202 TCR and H-2D^b^ obtained in NP202-NP366-H-2D^b^ complex (NP202-NP_366-374_/D^b^) that was determined and refined in the first place. All the models from structural determination were rebuilt using the program Coot ^107^. The final models was refined using the program phenix.refine ^108^. Structural validation of each model was performed using the program MolProbity ^109^.

### Molecular dynamics simulation

Molecular dynamics simulation under force was performed based on the method described in ^41^. Structure preparation, preparatory simulation, and trajectory analysis were performed by using CHARMM ^110^ and production runs were performed by using OpenMM ^111^. Crystal structures of NP202 (3.60Å), NP101 (2.20Å), NP207 (1.95Å), and NP203 (2.10Å) complexed with NP_366-374_/H-2D^b^ were prepared by converting the Cα, and Cβ sequences from human (used for x-ray crystallography) to mouse (used for other experiments). The structure of NP202 had the full mouse sequence except for S184 of Cβ, which was mutated to C184 for mouse. The resulting structure of the C-module was aligned and copied to NP101, NP207, and NP203 to generate the mouse versions. 5-residue long linkers were added to the C-terminal ends of MHC and the TCR based on their amino acid sequences so that the last residues were Y283 for MHC, T214 for Cα, and A245 for Cβ, respectively.

Without structures known, the linkers built by CHARMM are initially in fully straight conformations. They were relaxed by performing a short 120-ps simulation while the C-terminal Y283 C_α_ atom of MHC and the center of mass of the two C-terminal C_α_ atoms of TCRαβ (T214 and A245, respectively) are pulled in opposite directions by a 10-pN force in the FACTS implicit solvent environment of CHARMM ^112^. The force prevents collapse of the initially straight linkers due to entropic elasticity. Relaxation of linkers on MHC and TCRαβ was done in two separate simulations while coordinates of the protein except the linker were fixed in space. After the short 120-ps period, the linkers take reasonably relaxed conformations. The TCR-pMHC complex was then oriented such that the terminal C_α_ atoms used for relaxation are aligned along the x-axis. After this, solvation, neutralization, stepwise energy minimization, heating and equilibration of each system followed the protocol described in ^41^. The solvated system contained 204,000-268,000 atoms. The span of the terminal C_α_ atoms after the equilibration run was used to determine the 168-Å extension used to impose 1 kcal/[mol*Å^2^] harmonic restraints on them to apply fluctuating loads, as done previously ^24,41^. After additional 2-ns simulation at 300K with weak 0.001 kcal/[mol*Å^2^] harmonic restraints applied to to other C_α_ atoms for further equlibration in CHARMM, 600ns-700ns long production runs were performed in OpenMM for its faster performance in GPUs. Except for the relaxation run of added linkers in implicit solvent, the CHARMM36 force field was used in all simulations ^113^. For accuracy in simulation, we used an Ewald error tolerance of 10^-5^ which is 1/50 of the default value in OpenMM, and we used a 1.2-nm cutoff for short-range nonbonded interactions instead of OpenMM’s default value, 1.0 nm. We modified the contact analysis method developed in ^24^ to examine intradomain contacts as shown in Figure S7D. Additional simulations and more extensive analyses will be published elsewhere.

### Statistics

Statistical tests used throughout this study are indicated in figure legends or the main text. Paired t-tests were used for RgC mouse and p-tyrosine experiments. Chi-squared tests were employed for analysis of T_CM_ vs. T_EM_ clonotype distributions, Shannon entropy, and RNA-Seq clustering. Mann-Whitney U-tests evaluated differences in cell counts. Trend tests assessed graded gene expression across cell-clonotype categories. Two-sided Fisher’s exact tests were used for cross-reactive clonotype comparisons. Gene Set Enrichment Analysis (GSEA) was evaluated by a permutation test.

## Notes

### Competing Interest Statement

The authors have declared no competing interest.

### Summary of Updates

Figure 5 has been moved from the Extended Data section to the Main Data section and updated to clarify the TCR downstream mechanisms and their generality by increasing the number of TCRs examined. In addition, an author is added, and the supplemental files have been updated. The latter now includes further generalization of TCR distance analysis and linkage to TCM vs TEM branching, long and short, respectively, through incorporation of TCR[alpha][beta] clonotypes derived from two additional T cell repertoires directed at other epitopes of the IAV proteome that are distinct from that of NP_366-374_/D^b^. Also, note the title has a minor modification.

