## Supplemental information for "Atomistic TCR-ligand interactions shape memory T-cell differentiation"

[1]Laboratory of Immunobiology, Dana-Farber Cancer Institute; Boston, MA 02115, USA.
[2]Department of Medical Oncology, Dana-Farber Cancer Institute; Boston, MA 02115, USA.
[3]Department of Medicine, Harvard Medical School; Boston; MA 02115, USA.
[4]Structural Biology Center, X-Ray Science Division, Advanced Photon Source, Argonne National Laboratory; Lemont, IL 60439, USA.
[5]Department of Dermatology, Harvard Medical School; Boston, MA 02115, USA.
[6]Department of Informatics and Analytics, Dana-Farber Cancer Institute; Boston, MA 02115, USA.
[7]Department of Chemical and Biomolecular Engineering, Vanderbilt University; Nashville, TN 37212, USA
[8]Belfer Center for Applied Cancer Science, Dana-Farber Cancer Institute; Boston, MA, USA.
[9]Departments of Biomedical Engineering, Materials Science & Engineering, Physics & Astronomy, Texas A&M University; College Station, TX 77843, USA
[10]Center for AI and Natural Sciences, Korea Institute for Advanced Study; Seoul 02455, Republic of Korea
[11]Department of Molecular Physiology and Biophysics; Vanderbilt University School of Medicine; Nashville, TN 37232, USA

Table of contents:

Figures S1-S8

Tables S1-S4

Supplemental References

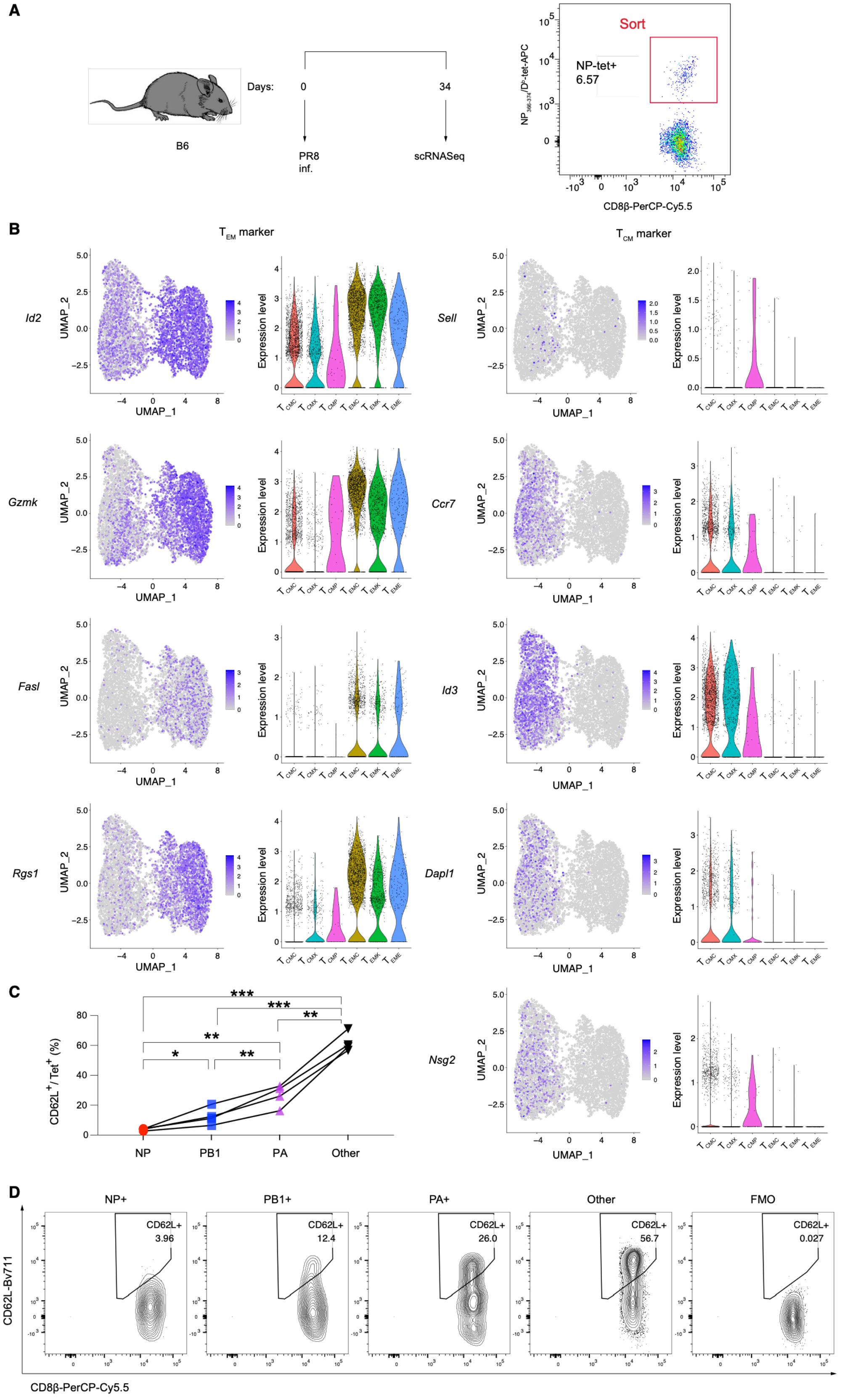
A
B6
Days:
0
34
PR8
inf.
scRNASeq
Sort
NP-tet+
6.57
CD8β-PerCP-Cy5.5
B
T EM marker
T CM marker
Id2
Gzmk
Fasl
Rgs1
Sell
Ccr7
Id3
Dapl1
Nsg2
UMAP_1
UMAP_2
Expression level
C
CD62L+/Tet+ (%)
NP
PB1
PA
Other
D
NP+
PB1+
PA+
Other
FMO
CD62L+
3.96
12.4
26.0
56.7
0.027
CD62L-Bv711
CD8β-PerCP-Cy5.5

**Figure S1. Schematic of scRNA-seq workflow and analysis of $T_{CM}$ and $T_{EM}$ marker expression on $NP_{366\text{-}374}/D^b$-specific T cells, related to Figure 1.** (A) Experimental workflow for analyzing $NP_{366\text{-}374}/D^b$-specific memory TCRs. mLNs were isolated from B6 mice 34 days after PR8 infection using the designated sort window in red. $CD8\beta^+$ $CD44^+$ $NP_{366\text{-}374}/D^b$-tetramer$^+$ cells were sorted, and scRNA-seq was performed. (B) Expression of representative $T_{EM}$ and $T_{CM}$ signature genes derived from $T_{CM}$ and $T_{EM}$ clusters, identified by separate scRNA-seq analysis on $CD8\beta^+CD44^+$ memory cells isolated by FACS from pooled mLNs of four B6 mice at dpi 33. The projection on the UMAP and violin plots showing the expression of representative genes. (C and D) Quantification (C) and representative FACS plots (D) showing CD62L expression on specific tetramer reactive $NP_{366\text{-}374}/D^b$-, $PB1_{703\text{-}711}/K^b$-, $PA_{224\text{-}233}/D^b$- T cells (NP, PB1 and PA), and NP/PB1/PA aggregate-negative T cells (Other) from mLN of four B6 mice at 33 days post-infection. FMO: fluorescence-minus one control. Data are representative of three independent experiments. ***P <0.001, **P <0.01, *P<0.05. P values were calculated by paired t-test.

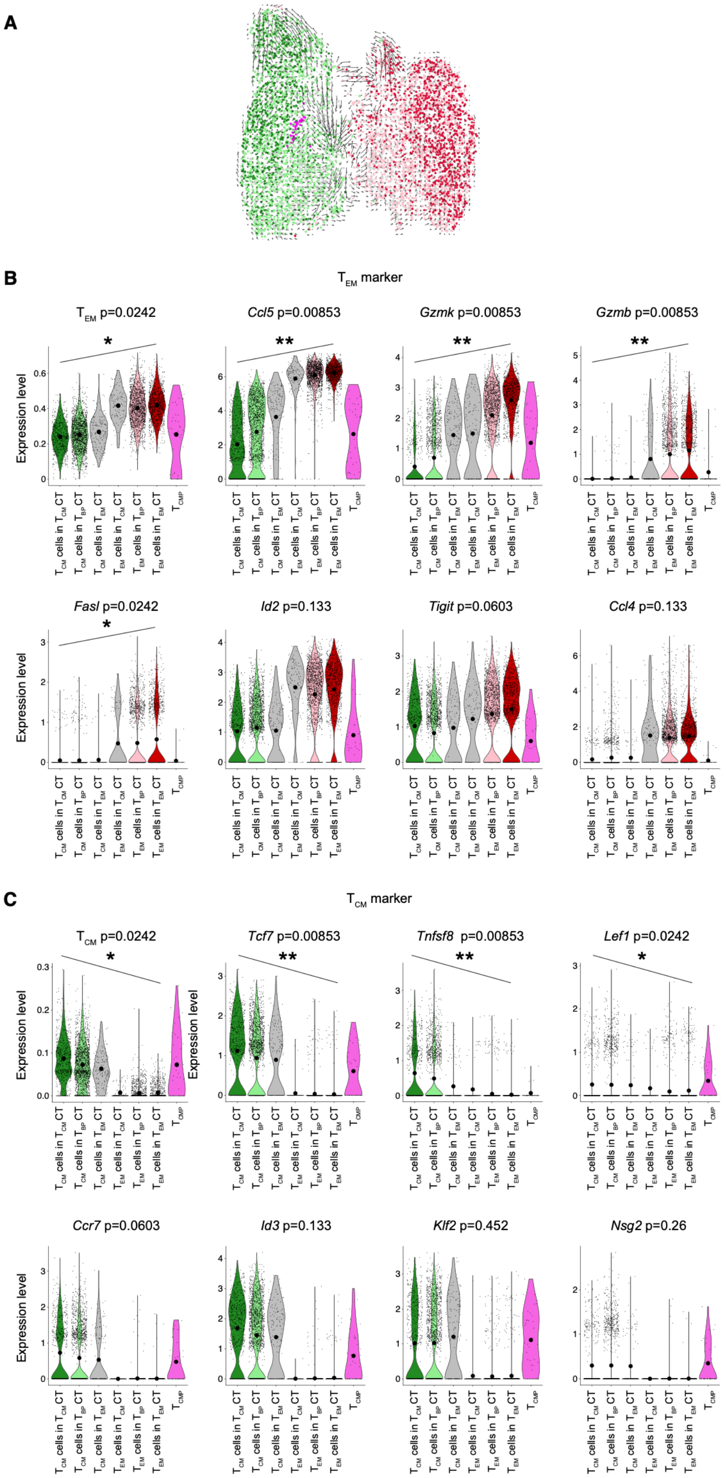
A
B
$T_{EM}$ marker
$T_{EM}$ p=0.0242
Ccl5 p=0.00853
Gzmk p=0.00853
Gzmb p=0.00853
Fasl p=0.0242
Id2 p=0.133
Tigit p=0.0603
Ccl4 p=0.133
C
$T_{CM}$ marker
$T_{CM}$ p=0.0242
Tcf7 p=0.00853
Tnfsf8 p=0.00853
Lef1 p=0.0242
Ccr7 p=0.0603
Id3 p=0.133
Klf2 p=0.452
Nsg2 p=0.26
Expression level
$T_{CM}$ cells in $T_{CM}$ CT
$T_{CM}$ cells in $T_{BP}$ CT
$T_{CM}$ cells in $T_{EM}$ CT
$T_{EM}$ cells in $T_{CM}$ CT
$T_{EM}$ cells in $T_{BP}$ CT
$T_{EM}$ cells in $T_{EM}$ CT
$T_{CMP}$

**Figure S2. RNA velocity analysis and $T_{EM}$ and $T_{CM}$ gene expression in clonally categorized memory T cells, related to Figure 1.** (A) UMAP showing RNA Velocity. (B) Violin plots showing the expression of aggregate and representative $T_{EM}$ signature genes among clonotype category (CT). Significance of trend across categories was generated using the Mann-Kendall trend test. (C) Violin plots showing the expression of aggregate and representative $T_{CM}$ signature genes among clonotype category, with Mann-Kendall test p-values indicated. **P <0.01, *P<0.05.

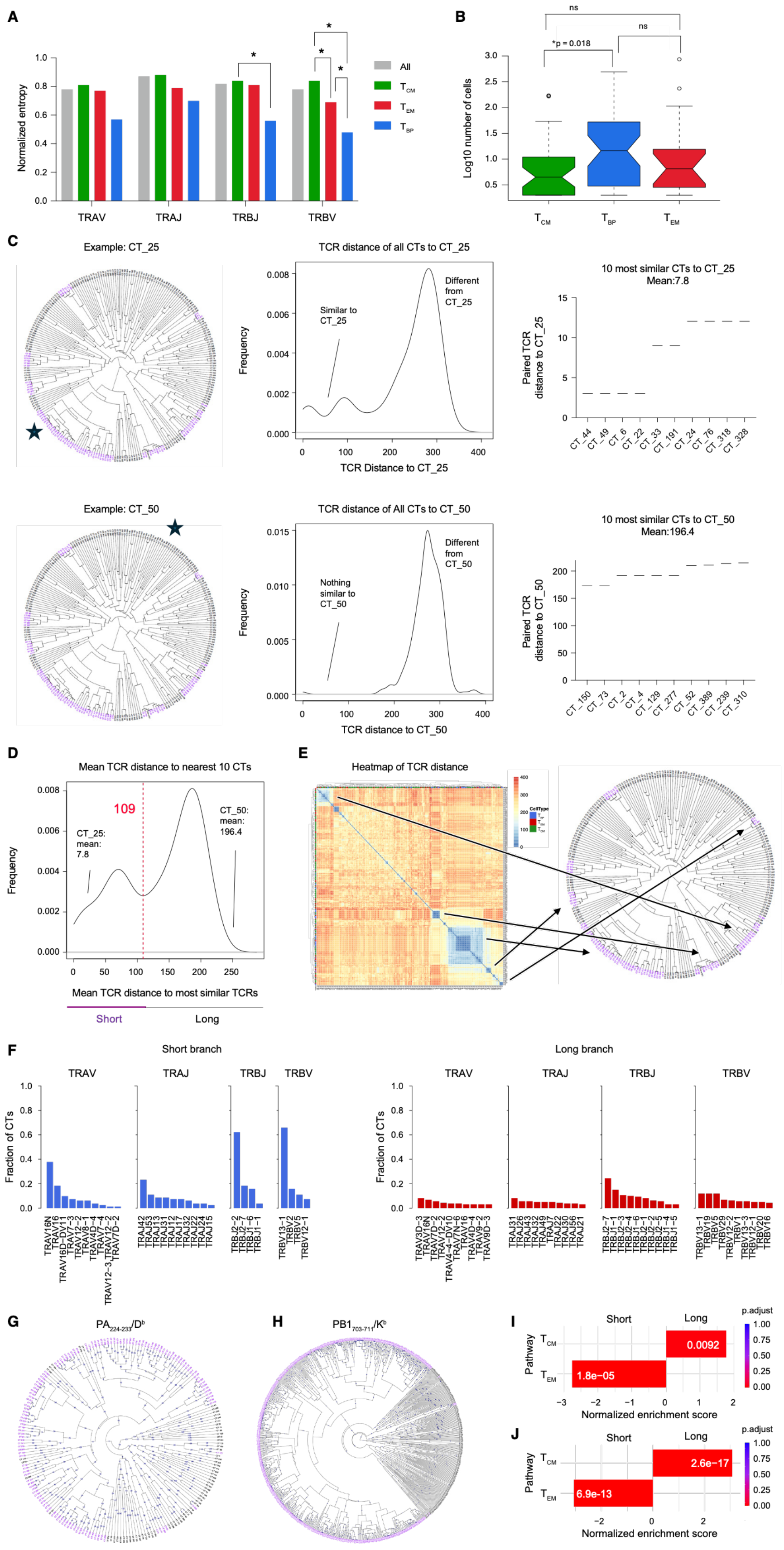

A
B
C
Example: CT_25
TCR distance of all CTs to CT_25
10 most similar CTs to CT_25
Mean:7.8
Example: CT_50
TCR distance of All CTs to CT_50
10 most similar CTs to CT_50
Mean:196.4
D
Mean TCR distance to nearest 10 CTs
Mean TCR distance to most similar TCRs
Short
Long
E
Heatmap of TCR distance
F
Short branch
Long branch
G
H
I
J
Normalized enrichment score

**Figure S3. $T_{CM}$ TCR sequences are more diverse whereas $T_{BP}$ clonotypes show greater expansion, related to Figure 2.** (A) Barplot of normalized Shannon entropy of V- and J-gene usage. Chi-square tests were used for statistics. *P<0.05. (B) Boxplot of cell number by clonotype polarity. Clonotypes used in the analysis were restricted to those with two or more cells. P-values for comparisons were calculated by Mann-Whitney-U. ns= not significant. (C) Sequence similarity analyses examples of two clonotypes: CT_25 ("short branch") and CT_50 ("long branch"). Density plots show the TCR Distance of the indicated clonotype to all other clonotypes. The TCR distances of the 10 most sequence similar clonotypes CT_25 and CT_50 are also shown**.** (D) Density plot of the mean TCR distance of the 10 most similar clonotypes for all clonotypes. The value of 109 (between the two peaks of the plot) was used as the threshold between short branch and long branch clonotypes**.** (E) Heatmap of pairwise TCR distances for all clonotypes. For a select subset of groups of clonotypes, arrows indicate the location of those clonotypes on the phylogram relative to the heatmap. (F) Bar plots indicating TRAV, TRAJ, TRBV, and TRBJ gene usage in clonotypes categorized as short- or long-branch based on TCR sequence similarity. The ten most frequently used gene segments are shown. (G and H) TCR sequence distant tree are shown for $PA_{224\text{-}233}/D^b$- (G), and $PB1_{703\text{-}711}/K^b$-specific TCRs (H). Long-branch clonotypes are shown in black and short-branch clonotypes are shown in purple. (I and J) $T_{EM}$ and $T_{CM}$ gene enrichment analysis for short and long branch are shown for PA- (I), and PB1-specific T cells (J).

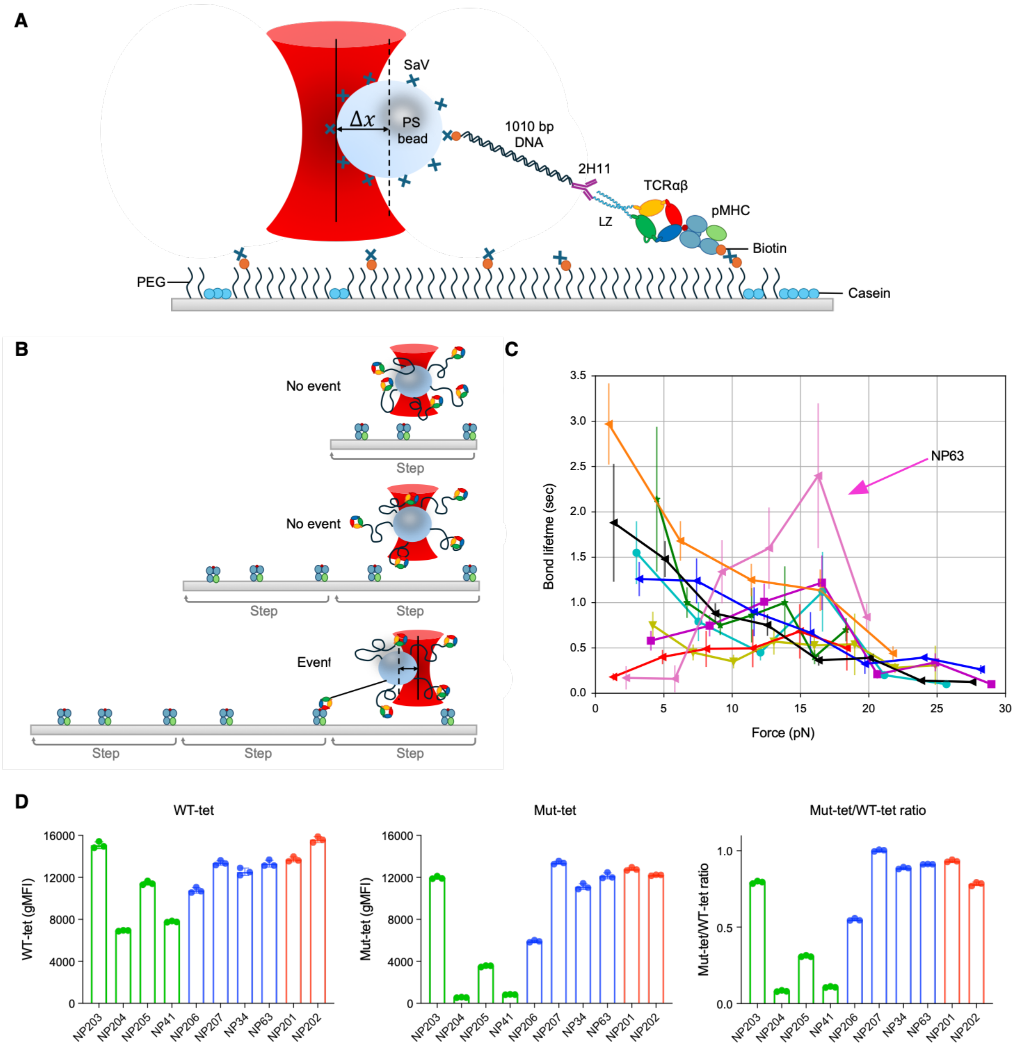


**Figure S4. SM assay details, clonotype force-bond-lifetimes, and CD8 dependency of NP$_{366-374}$/D$^b$ tetramer binding by NP-specific TCRs, related to Figure 2.** (A) Cartoon schematic depicting tether assay geometry and connectivity for probing the lifetime of TCRαβ-pMHC bonds. 'ΔX' denotes bead displacement from the trap center. (B) Cartoon schematic depicting binding probability assay monitoring bead position and tether formation during repetitive translation steps of the stage relative to the trap. Total steps versus those indicating binding events can be tallied to assess binding probability. (C) Force versus bond-lifetime curves for all TCRαβ-pMHC pairs shown in the same plot. Here, digital NP63 (indicated in pink with arrow) shows stronger bond lifetime at critical force of ~15 pN relative to other clonotypes with colors as indicated in Figure 2G. (D) BW5148CD8αβ cells expressing individual NP-specific TCRs were incubated with 10 μg/mL wild-type NP$_{366-374}$/D$^b$ tetramer (left, WT-tet) or CD8-binding site deficient (D227K-mutated) $NP_{366\text{-}374}/D^b$ tetramer (middle, mut-tet) for 30 min at RT. Fluorescence intensity was quantified by flow cytometry. The ratio of binding (mut-tet/ WT-tet) was calculated for each TCR (right). $T_{EM}$ TCRs are shown in red, $T_{CM}$ in green, and $T_{BP}$ in blue. The data show three technical replicates and are representative of two independent experiments.

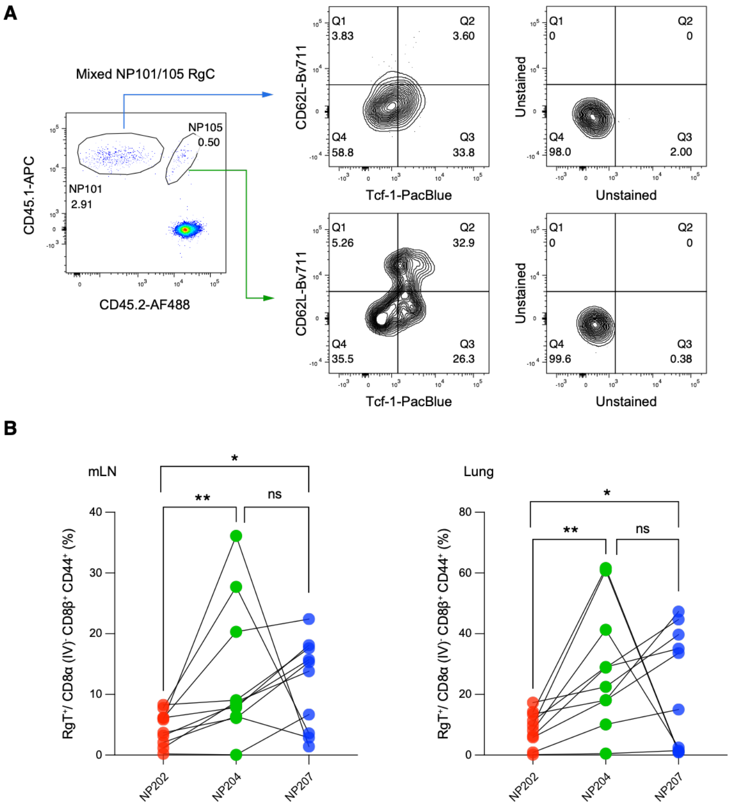


**Figure S5. Rg mouse analysis recapitulates clonotype polarization patterns identified by scRNA-seq, related to Figure 2.** (A) Assessment of memory marker expression in mixed NP101/105 RgC mice shown in Figure 2J. Representative flow cytometry plots showing the gating strategy used to evaluate $T_{CM}$ cells ($CD62L^+$ and/or Tcf-$1^+$) and $T_{EM}$ cells ($CD62L^-$ Tcf-$1^-$) in transferred NP101 ($CD45.1^+$ $CD45.2^-$) and NP105 ($CD45.1^+$ $CD45.2^+$) Rg T cells from mixed NP101/ NP105 RgC mice in mLN at 30 days post-PR8 infection. Data are gated $Live^+$ $CD8\beta^+$ $CD44^+$ cells. (B) Frequency of Rg T cells in mixed NP202/204/207 RgC mice after secondary infection. Mixed NP202-CD45 (1/1)/NP204-CD45 (1/2)/NP207-Thy1 (1/1) RgC mice were first infected with PR8 and, 33 days later, received a secondary X31 infection. Four days after the secondary challenge, the frequency of Rg T cells in the mLN (left) and lung (right) was quantified based on congenic marker expression by flow cytometry. **P <0.01, *P<0.05. ns=not significant. P values were calculated by paired t-test.

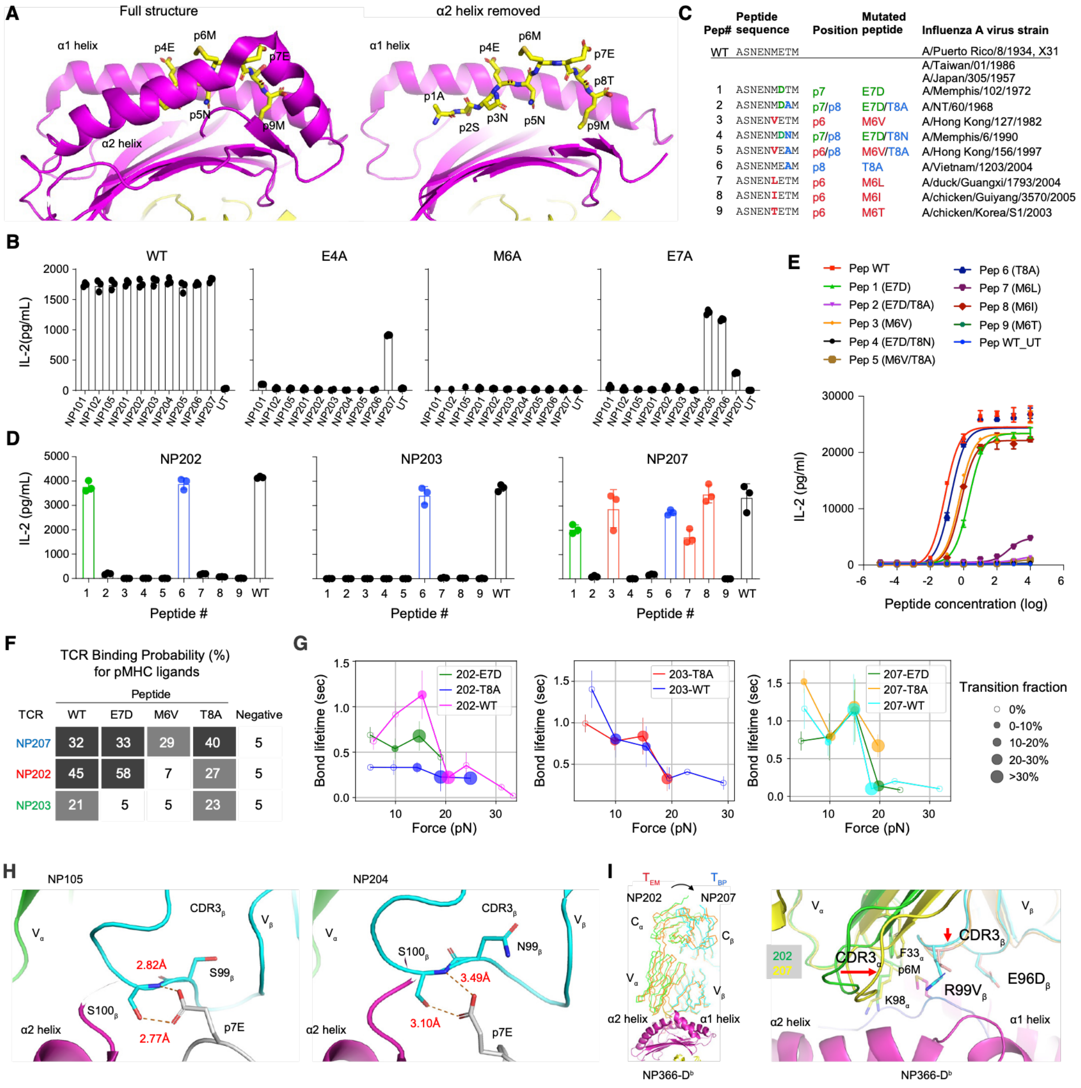


**Figure S6. Assessment and structural explanation of crossreactivity in memory clonotypes, related to Figure 2.** (A) NP366-374/D$^b$ structure with peptide (yellow) and antigen presenting platform (magenta) shown. The α2 helix has been removed on the right subpanel to display more clearly those featured peptide sidechains. (B) IL-2 production from culture supernatants of BW cells transduced with 10 different NP-specific TCRs, or untransduced (UT), following stimulation with 1 μg/mL of wild-type (WT) or point-mutated NP peptides (E4A, M6A, and E7A) loaded onto APCs overnight. The range of detection of IL-2 is 0-1861 pg/mL. (C) List of naturally occurring variant NP366-374/D$^b$ peptides used in Figure 2K and Figure S6D-S6G. Sequences were derived from IAV strains reported in Zhong et al. [1]. (D) Representative IL-2 production from the supernatants of NP-specific TCR-transduced BW cells (NP202, NP203, and NP207) stimulated with 1 μg/mL of wild-type (WT) or naturally occurring mutated NP peptides listed in Figure S8C. The range of detection of IL-2 is 0-4548 pg/mL. (E) IL-2 production from NP207-transduced BW cells stimulated with titrated concentrations of WT and mutated NP peptides. On the X-axis, 0=1ng/mL. (F) SM optical tweezer tether forming probability of TCR-pMHC binding for indicated peptide epitope variants under identical experimental conditions. Probability represents the percentage of piezo stage steps resulting in a binding event (N=200 per condition). Values not different from the negative control are indicate as a white background, while 17-29% are in grey, and 30% or greater in black. (G) Force-bond lifetime curves for indicated clonotypes binding with WT and variant ligands E7D and T8A, with noted fraction undergoing transitions. NP202 binding is shown for WT peptide (magenta curve, N=136), E7D (green curve, N=97), and T8A (blue curve, N=69). NP203 binding is shown for WT peptide (blue curve, N=58) and T8A (red curve, N=187). NP207 binding is shown for WT (cyan curve, N=100), E7D (green curve, N=65), and T8A (orange curve, N=136). In the case of NP203 and NP207 crossreactivity profiles were strikingly similar to WT. For NP202, crossreactivity lifetimes were somewhat muted relative to WT yet performed as well or better at higher force ranges with respect to transition fraction. (H) NP105 (left) and NP204 (right) depicting the interactions with $NP_{366-374}/D^b$, both of which manifest β chain dominance utilizing a related but distinct CDR3β and entirely different Vα and Vβ germline-encoded segments. The p7E residue of peptide plays a critical role in the interaction with the CDR3β of both TCRs.  Apparently, their divergent hydrogen bond lengths, determine their p7E to D mutation responsiveness. In the NP105-$NP_{366-374}/D^b$ complex, p7E clearly forms two short H-bonds. This conformation may explain why NP105 is still reactive to the p7D mutant even though bond strength is likely reduced. In NP204-$NP_{366-374}/D^b$ complex, p7E appears to form two long H-bonds to CDR3β with bond lengths equal to or larger than 3.10Å. The lower resolution of the NP204-$NP_{366-374}/D^b$ structure makes the latter estimate an approximation. Nonetheless, a p7D mutation likely diminishes these H-bonds further, abrogating crossreactivity with the shorter peptide D side chain. (I) Structural differences in NP202 versus NP207 regarding tilt onto $NP_{366-374}/D^b$ as shown both overall (left) and at the interface (right) emphasizing the sizeable shift of CDR3α and with indicated residues shown.

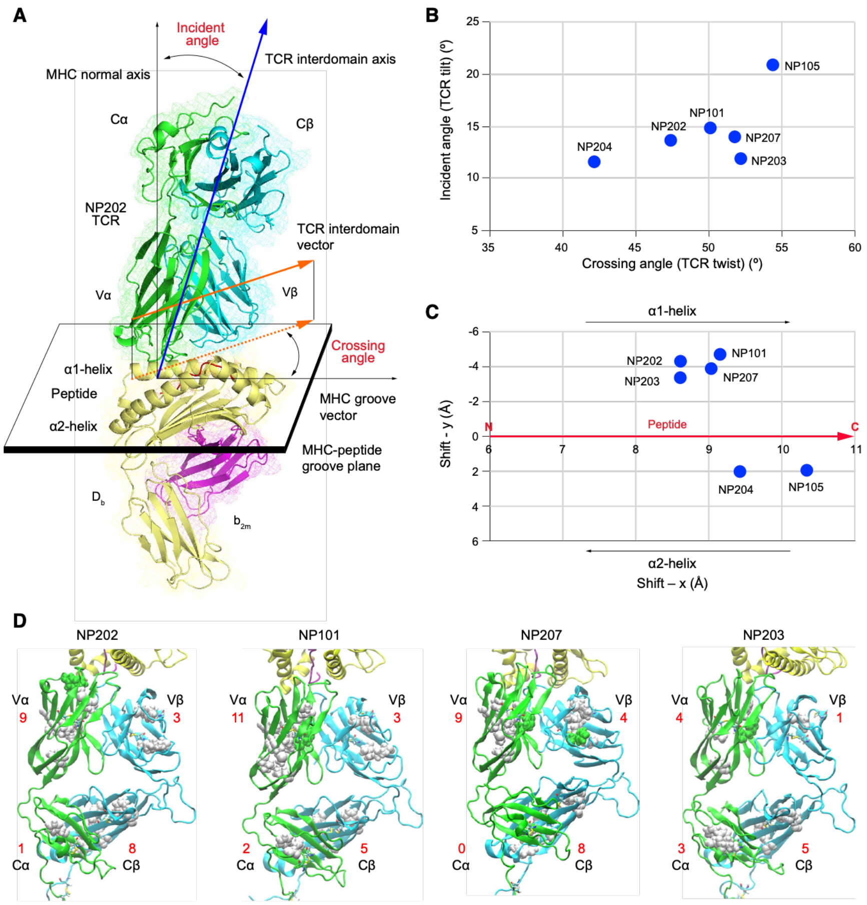


**Figure S7. NP TCR docking angles and shifts on pMHC and divergent mechanical responses of four NP TCRs binding $NP_{366\text{-}374}/D^b$, related to Figure 3**. (A) The definitions of TCR docking angles are illustrated using the NP202-$NP_{366\text{-}374}/D^b$ complex structure. The angles shown in the model are detailed in the TCR3d database (https://tcr3d.ibbr.umd.edu/). In short, TCR angles include crossing and incident angles, which represent the orientation of the TCR on the pMHC platform. A small crossing angle indicates a small twist of the TCR relative to the long axis of the peptide. Meanwhile, a small incident angle indicates a slight tilt from a vertical direction normal to the pMHC platform. (B) The crossing and incident angles of the six NP TCRs on the pMHC plane. The docking angles seemingly have no direct correlation with the polarization of NP TCRs. (C) The shifts of NP TCRs on the pMHC plane. The definition of these shifts is detailed in the TCR3d database. For clarity, three vectors, red for peptide and two black for α1- and α2-helices are drawn relative orientation and directions of increasing sequential residue numbers. The closeness of NP105 and NP204 in this diagram contrasts sharply with their well-separated positions in the docking angle diagram as shown in Figure S7B. The shifts of these two NP TCRs more toward the C-terminal peptide position and/or their a2 helix proximity apparently relates to their common β-dominant features and $T_{CM}$ polarity. In contrast, the clustering of four related TCRs NP101, NP202, NP203 and NP207 that share almost identical amino acid sequences suggests that other factors play roles in their distinct TCR polarities. (D) Simulations of the NP TCR-pMHC complexes were performed under load following the protocol described in Chang-Gonzalez et al.[2]. $T_{EM}$ NP202 is followed by $T_{BP}$ NP101 and $T_{BP}$ NP207 that differ from NP202 by 1 and 2 residues, respectively, and finally by $T_{CM}$ NP203 with 3 residue differences. For each of the 4 subdomains, the number of residues that form 6 or more intra-domain contacts with occupancy greater than 80% during the simulation (600ns-700ns in duration) are shown (red). Space filling representations of hydrophobic and polar residues are shown in white and green, respectively. Despite very similar TCR sequences, the number as a measure of the packing or rigidity of the domain varies significantly, suggesting differential responses to load.

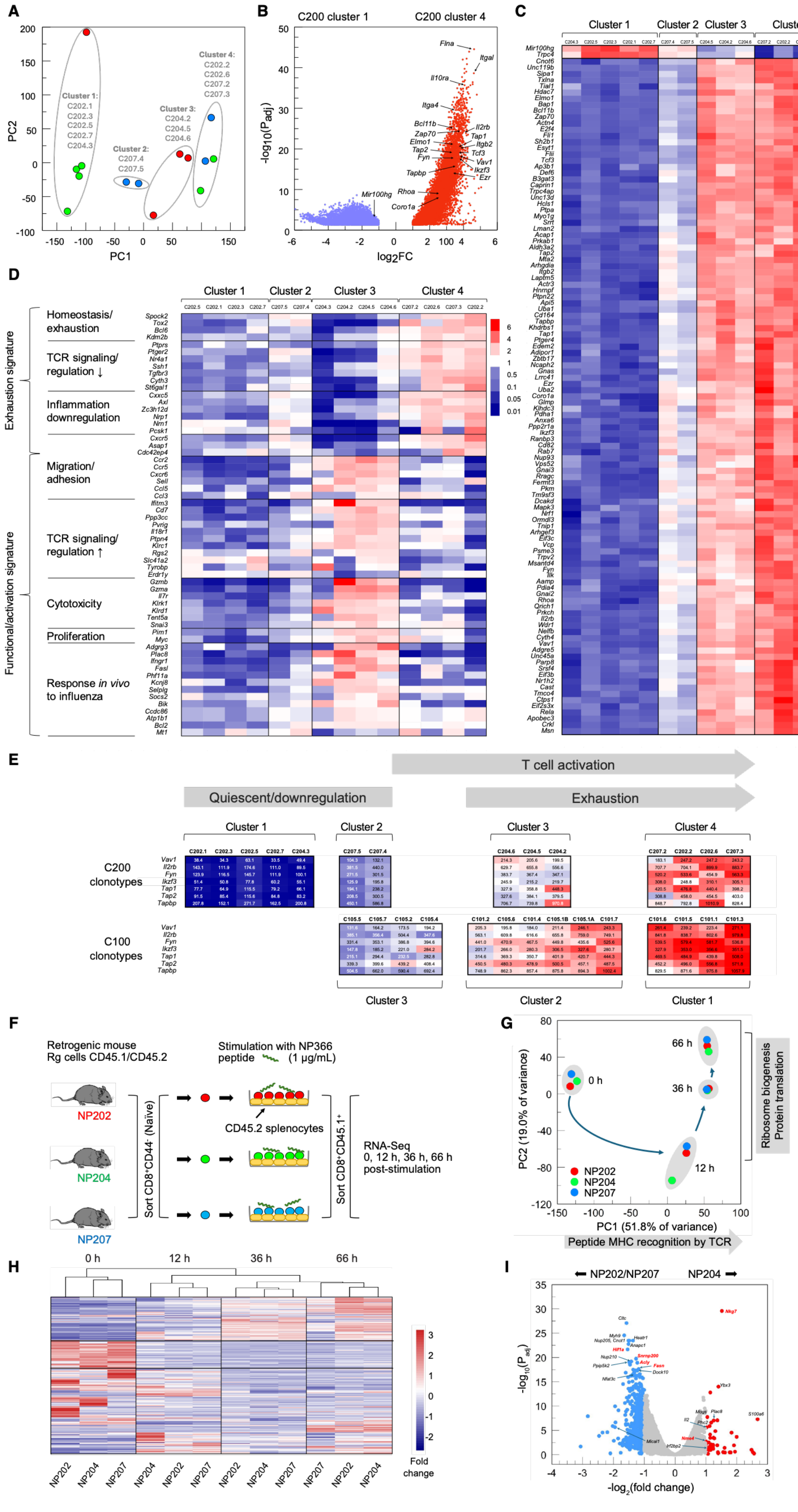
A
Cluster 1:
Cluster 2:
Cluster 3:
Cluster 4:
PC1
PC2
B
C200 cluster 1
C200 cluster 4
log2FC
-log10(Padj)
C
Cluster 1
Cluster 2
Cluster 3
Cluster 4
D
Homeostasis/ exhaustion
TCR signaling/ regulation ↓
Inflammation downregulation
Migration/ adhesion
TCR signaling/ regulation ↑
Cytotoxicity
Proliferation
Response in vivo to influenza
Exhaustion signature
Functional/activation signature
E
T cell activation
Quiescent/downregulation
Exhaustion
C200 clonotypes
C100 clonotypes
F
Retrogenic mouse Rg cells CD45.1/CD45.2
Stimulation with NP366 peptide (1 μg/mL)
NP202
NP204
NP207
CD45.2 splenocytes
RNA-Seq 0, 12 h, 36 h, 66 h post-stimulation
G
PC1 (51.8% of variance)
PC2 (19.0% of variance)
Ribosome biogenesis Protein translation
Peptide MHC recognition by TCR
H
0 h
12 h
36 h
66 h
Fold change
I
NP202/NP207
NP204
-log2(fold change)

**Figure S8. Retrogenic T cell functionality among memory clonotypes 30 days following IAV infection (related to Figure 6) or following high copy number pMHC stimulation *in vitro* of their naïve counterparts.** ***A-E in vivo stimulated***. (A) PCA of mLN-localized retrogenic T cells from mice adoptively transferred with 2.5 x $10^4$ C202 (green), C204 (red), or C207 (blue), cells expressing individual NP202, NP204 or NP207 TCR clonotypes (n = 14 mice), and analyzed by RNA-seq 30 days after IAV infection as in Figure 6. Each data point represents cells isolated from an individual mouse. Four clusters are identified. (B) Volcano plot of DEG between Cluster 1 and Cluster 4 identifies significant gene upregulation in cluster 4, where the majority relate to T cell functionality including migration, TCR signaling, and responses to viral infection. (C) Manual curation of the top 250 differentially regulated genes for those with known T cell functionality as described above contracted the list to 111 transcripts. Hierarchical clustering of the curated list delineated the same 4 clusters observed in panel A where a gradient of minimal to high T cell functionality was observed from cluster 1 to cluster 4. The lncRNA *Mir100hg* and *Trpc*, both associated with downregulation of T cell activation, demonstrate an inverse expression pattern to that of the activation-related transcripts. The z-score is depicted for each gene. (D) Application of the curated C100 T cell 59 gene transcript list (Figure 6D) to the C200 series identifies an exhaustion signature for Cluster 4 and an activated $T_{CM}$ signature for Cluster 3. Each row scaled where 1 (white) = base gene mean, red = fold-change above, and blue = fold-change below, the base mean. (E) Expression (transcripts per million; tpm) for a T cell activation-related panel to identify a broad cluster correspondence between the C100 and C200 series by activation (further details in Methods). White indicates the base gene expression mean for each gene across the aggregated data set, while blue and red define the lower and upper tpm bounds, respectively, for that gene (n = 28). Each column for the 7 activated T cell transcripts represents clonotypic cell expression from a single mouse (n = 28). ***F-I in vitro stimulated***.(F) Schematic depicting the procedure for isolation of 3 retrogenic naïve $CD8^+$ CD45 (1/2) T cell populations, each bearing a unique TCR recognizing $NP_{366\text{-}374}$ peptide presented by $H2\text{-}D^b$ on CD45 (2/2) splenocytes as APC, and FACS sorting at varying times up to 66 h for transcriptome analysis by RNA-Seq. Gating for FACS sorting of the retrogenic naïve $CD8^+$ T cells prior to seeding on $NP_{366\text{-}374}$ peptide-presenting cells is depicted in Data S8. (G) Principal component analysis mapping at each time point for the transcriptomes of the three unique TCR-bearing retrogenic $CD8^+$ T cells. Gene ontology (GO) analysis of the top 500 genes contributing to PC1 and PC2, respectively, identifies the main identified biological processes (see Data S8 for full listing of GO output). Note that the ribosome biogenesis signature, indicating increased protein translation, was not well-represented in NP204 at 12 h (see panel I for differential gene expression). (H) Hierarchical clustering delineates the samples by time point. The upper row, bounded by black lines identifies a cluster of genes upregulating with time. The second row, bounded by black lines indicates a cluster of genes downregulated after peptide presentation. (I) Differential gene expression between NP204 and NP202/NP207 after peptide-MHC presentation for 12 h. Genes well expressed in NP202 and NP207 $CD8^+$ T cells are indicated as blue dots with representative genes annotated and selected genes of interest highlighted in red font. Genes well expressed in NP204 are depicted as red dots with representative genes annotated and selected genes of interest highlighted in red font. The grey points indicate genes with a $\log_2$ fold-change less than 2 and/or a $P_{adj} < 0.05$ (see Data S8 for full listing of gene expression data).

| | Name | Clonotype | Cell # | % TEM | % TCM | p.adj | TRAV | TRBV | CDR3α | CDR3β | TRAJ | TRBJ |
|---|---|---|---|---|---|---|---|---|---|---|---|---|
| | NP201 | CT_1 | 863 | 91.9 | 7.8 | 5.73E-149 | TRAV12-2 | TRBV13-1 | CAPNSNNRIFF | CASKGGGVTGQLYF | TRAJ31 | TRBJ2-2 |
| | | CT_9 | 234 | 74.7 | 24.4 | 1.20E-12 | TRAV16N | TRBV13-1 | CAMRGNSGTYQRF | CASSGGSNTGQLYF | TRAJ13 | TRBJ2-2 |
| $T_{EM}$ | NP202 | CT_20 | 107 | 84.2 | 15.9 | 2.96E-11 | TRAV16 | TRBV5 | CAMREGKGGGSNYKLTF | CASSQELGGRYEQYF | TRAJ53 | TRBJ2-7 |
| | | CT_55 | 16 | 81.3 | 18.8 | 0.098690892 | TRAV16D-DV11 | TRBV13-1 | CAMREGTGTGSNRLTF | CASSGGSNTGQLYF | TRAJ28 | TRBJ2-2 |
| | | CT_58 | 15 | 86.6 | 13.4 | 0.042434702 | TRAV16N | TRBV13-1 | CAMREGGTGKLTF | CASRGGSNTGQLYF | TRAJ27 | TRBJ2-2 |
| | | CT_67 | 10 | 90 | 10 | 0.098690892 | TRAV7-4 | TRBV2 | CAASPSSGSWQLIF | CASSQDRRNSYNSPLYF | TRAJ22 | TRBJ1-6 |
| | NP203 | CT_11 | 172 | 29.1 | 70.4 | 8.78E-08 | TRAV16 | TRBV5 | CAMREGRGGGSNYKLTF | CASSQDLGGLYEQYF | TRAJ53 | TRBJ2-7 |
| | NP204 | CT_12 | 169 | 5.4 | 94.1 | 9.65E-37 | TRAV7-4 | TRBV1 | CAAILATGGNNKLTF | CTCSADRRNSYNSPLYF | TRAJ56 | TRBJ1-6 |
| | NP205 | CT_13 | 164 | 14.6 | 85.3 | 4.48E-21 | TRAV10 | TRBV19 | CAASDSNNRIFF | CASRDWRQNTLYF | TRAJ31 | TRBJ2-4 |
| $T_{CM}$ | | CT_27 | 54 | 11.1 | 88.9 | 7.81E-09 | TRAV9N-4 | TRBV16 | CALSMSSNNRIFF | CASSLARQGKYF | TRAJ31 | TRBJ2-7 |
| | | CT_28 | 52 | 11.5 | 88.4 | 4.08E-08 | TRAV16 | TRBV5 | CAMREGRGGGSNYKLTF | CASSQDLGGLYEQYF | TRAJ53 | TRBJ2-7 |
| | | CT_29 | 51 | 15.7 | 84.3 | 1.50E-06 | TRAV7D-3 | TRBV2 | CAVMNYGSSGNKLIF | CASSQDRRGSYNSPLYF | TRAJ32 | TRBJ1-6 |
| | NP41 | CT_42 | 22 | 18.1 | 81.8 | 0.013417636 | TRAV7N-6 | TRBV5 | CAVSMNQGGRALIF | CASSQDRWGNYAEQFF | TRAJ15 | TRBJ2-1 |
| | | CT_47 | 20 | 10 | 90 | 0.001180773 | TRAV8D-1 | TRBV12-1 | CASRTGGYKVVF | CASSLGPSSYEQYF | TRAJ12 | TRBJ2-7 |
| | | CT_64 | 11 | 27.3 | 72.7 | 0.319323145 | TRAV4-4-DV10 | TRBV13-1 | CAARRPGYQNFYF | CASSDARQTEVFF | TRAJ49 | TRBJ1-1 |
| | NP206 | CT_2 | 492 | 30.5 | 69.1 | 7.90E-19 | TRAV4D-4 | TRBV13-1 | CAASNSGTYQRF | CASSARTANTEVFF | TRAJ13 | TRBJ1-1 |
| | | CT_3 | 431 | 61.5 | 38.3 | 0.00013793 | TRAV16N | TRBV13-1 | CAMREAVGDNSKLIW | CASKGGGNTGQLYF | TRAJ38 | TRBJ2-2 |
| | | CT_4 | 415 | 46.8 | 52.5 | 0.284843022 | TRAV4D-4 | TRBV13-1 | CAASNSGTYQRF | CASSARTANTEVFF | TRAJ13 | TRBJ1-1 |
| | NP101" | CT_5 | 362 | 40.9 | 59.1 | 0.000730254 | TRAV16 | TRBV5 | CAMREGKGGGSNYKLTF | CASSQDLGGRYEQYF | TRAJ53 | TRBJ2-7 |
| | NP34* | CT_6 | 329 | 33.7 | 65.6 | 5.62E-09 | TRAV16N | TRBV13-1 | CAMRVSGGSNAKLTF | CASSGGSNTGQLYF | TRAJ42 | TRBJ2-2 |
| | NP207 | CT_7 | 276 | 53.3 | 44.6 | 0.477927988 | TRAV16 | TRBV5 | CAMREGKGGGSNYKLTF | CASSQDLGGVYEQYF | TRAJ53 | TRBJ2-7 |
| | | CT_8 | 271 | 54.3 | 45.3 | 0.477927988 | TRAV16N | TRBV13-1 | CAMRTASLGKLQF | CASSGGSNTGQLYF | TRAJ24 | TRBJ2-2 |
| | | CT_15 | 158 | 45 | 55.1 | 0.310195462 | TRAV16N | TRBV19 | CAMRVKGNTGKLIF | CASSPLGGGAETLYF | TRAJ37 | TRBJ2-3 |
| | NP34* | CT_22 | 63 | 47.6 | 52.4 | 0.768008573 | TRAV16N | TRBV13-1 | CAMRVSGGSNAKLTF | CASSGGSNTGQLYF | TRAJ42 | TRBJ2-2 |
| | | CT_24 | 58 | 34.5 | 65.5 | 0.064091157 | TRAV16 | TRBV13-1 | CAMRTSGGSNAKLTF | CASSGGSNTGQLYF | TRAJ42 | TRBJ2-2 |
| | NP34* | CT_25 | 55 | 63.6 | 34.5 | 0.2020416 | TRAV16 | TRBV13-1 | CAMRVSGGSNAKLTF | CASSGGSNTGQLYF | TRAJ42 | TRBJ2-2 |
| | | CT_30 | 50 | 68 | 32 | 0.098690892 | TRAV16 | TRBV5 | CAMREGRGGGSNYKLTF | CASSQDLGGRYEQYF | TRAJ53 | TRBJ2-7 |
| $T_{BP}$ | NP63' | CT_33 | 40 | 45 | 55 | 0.649391811 | TRAV16 | TRBV13-1 | CAMRVAGGSNAKLTF | CASSGGSNTGQLYF | TRAJ42 | TRBJ2-2 |
| | | CT_34 | 34 | 35.3 | 64.7 | 0.273761197 | TRAV16N | TRBV13-1 | CAMRRSGGSNAKLTF | CASSGGGNTGQLYF | TRAJ42 | TRBJ2-2 |
| | | CT_35 | 33 | 60.7 | 36.3 | 0.396413252 | TRAV16N | TRBV13-1 | CAMREANTGKLTF | CASRGGSNTGQLYF | TRAJ27 | TRBJ2-2 |
| | NP34* | CT_44 | 23 | 43.4 | 56.5 | 0.719835214 | TRAV16N | TRBV13-1 | CAMRVSGGSNAKLTF | CASSGGSNTGQLYF | TRAJ42 | TRBJ2-2 |
| | | CT_46 | 21 | 52.4 | 47.7 | 1 | TRAV12D-1 | TRBV5 | CALSERTEGADRLTF | CASSQDLGGRYEQYF | TRAJ45 | TRBJ2-7 |
| | | CT_48 | 18 | 55.6 | 44.4 | 0.970878204 | TRAV16N | TRBV13-1 | CAMRDPNSAGNKLTF | CASRGGSNTGQLYF | TRAJ17 | TRBJ2-2 |
| | | CT_50 | 18 | 27.8 | 66.7 | 0.280379199 | TRAV4D-4 | TRBV2 | CAAALTSGGNYKPTF | CASSPRTRNTEVFF | TRAJ6 | TRBJ1-1 |
| | NP34 | CT_49 | 17 | 41.2 | 58.8 | 0.694008343 | TRAV16N | TRBV13-1 | CAMRVSGGSNAKLTF | CASSGGSNTGQLYF | TRAJ42 | TRBJ2-2 |
| | | CT_52 | 16 | 50 | 50 | 1 | TRAV4D-4 | TRBV13-3 | CAARGPGSGGKLTL | CASRDSANTEVFF | TRAJ44 | TRBJ1-1 |
| | NP101" | CT_56 | 16 | 43.7 | 56.3 | 0.768008573 | TRAV16 | TRBV5 | CAMREGKGGGSNYKLTF | CASSQDLGGRYEQYF | TRAJ53 | TRBJ2-7 |
| | | CT_54 | 15 | 60.1 | 40 | 0.768008573 | TRAV9D-3 | TRBV13-1 | CAVSMTNSAGNKLTF | CASRGGSNTGQLYF | TRAJ17 | TRBJ2-2 |
| | | CT_59 | 14 | 50 | 50 | 1 | TRAV16N | TRBV13-1 | CAMRDYQGGRALIF | CASSGGGRSAEQFF | TRAJ15 | TRBJ2-1 |
| | | CT_62 | 10 | 40 | 60 | 0.719835214 | TRAV10D | TRBV14 | CAASWGGLSGKLTF | CASSRDRGRGEVFF | TRAJ2 | TRBJ1-1 |

**Table S1. Representative $T_{EM}$, $T_{CM}$, and $T_{BP}$ clonotypes, related to Figure 1.** Clonotypes represented by more than 10 cells showing V-gene, J-gene, and CDR3 usage. Fraction of cells in $T_{EM}$ and $T_{CM}$ clusters are given with p-values generated by a binomial test versus overall cell distribution. Clonotypes are grouped by $T_{EM}$, $T_{CM}$, and TBP categories. Clonotypes were categorized as $T_{CM}$- or $T_{EM}$-type if more than 70% of their progeny were polarized toward one or more of the three respective subcluster, as defined in Figure 1A, with a statistically significant difference (p.adj < 0.05). Clonotypes with a more balanced distribution that did not meet this threshold were classified as $T_{BP}$. The asterisk (*) indicates that the clonotypes have the same amino acid sequence as NP34 but different nucleotide sequences. The prime (') dictates the usage of TRAV16 for CT_33, which differs at two residues from TRAV16N used by NP63 (V→ M and V→ E) at sites distal to the pMHC interface. The double prime (") indicates that the clonotypes have the same amino acid sequence as NP101 but different nucleotide sequence.

| *Data collection* | NP202 TCR | NP202-$NP_{366}$-H-$2D^b$ | NP207 TCR | NP207-$NP_{366}$-H-$2D^b$ | NP101-$NP_{366}$-H-$2D^b$ | NP105-$NP_{366}$-H-$2D^b$ | NP203-$NP_{366}$-H-$2D^b$ | NP204-$NP_{366}$-H-$2D^b$ |
|---|---|---|---|---|---|---|---|---|
| Space group | $P2_1$ | $P2_1$ | $P2_1$ | $P1$ | $P1$ | $P2_12_12_1$ | $P1$ | $P2_12_12_1$ |
| Unit Cell dimensions | | | | | | | | |
| *a, b,* c (Å) | 62.07, 127.4, 66.56 | 94.29, 89.59, 104.3 | 62.12, 126.0, 66.63 | 57.53, 68.02, 74.68 | 51.10, 53.59, 101.5 | 58.17, 131.7, 141.9 | 56.78, 66.62, 76.78 | 91.09, 146.4, 154.0 |
| *α, β, γ* (°) | 90, 110.1, 90 | 90, 96.17, 90 | 90, 110.2, 90 | 72.83, 89.36, 67.21 | 79.28, 88.37, 65.64 | 90, 90, 90 | 68.78, 89.27, 66.42 | 90, 90, 90 |
| Protein MW Da (No. of a.a.) | 52,838.4 (469) | 98,190.4 (859) | 52,767.2 (469) | 99,026.1 (867) | 99,083.1 (867) | 98,667.6 (863) | 99,068.1 (867) | 97,881.0 (857) |
| Complex (AU) | 2 | 1.5 [1] | 2 | 1 | 1 | 1 | 1 | 2 |
| Wavelength (Å) | 0.9795 | 0.9792 | 0.9795 | 0.9786 | 0.9786 | 0.9786 | 0.9786 | 0.9792 |
| Resolution (Å) | 2.56 – 32.6 | 3.60 – 52.0 | 2.50 – 30.9 | 1.95 – 45.0 | 2.20 – 48.0 | 2.80 – 50.0 | 2.10 – 50.0 | 4.45 – 44.0 |
| Number of Unique | 30,273 (1,226) [2] | 18,351 (1,168) [3] | 32,374 (1,329) [4] | 68,675 (2,903) [5] | 47,985 (2,413) [6] | 26,780 (1,758) [7] | 53,182 (2,438) [8] | 13,111 (636) [9] |
| Completeness (%) | 97.5 (81.0) [2] | 96.7 (92.0) [3] | 97.7 (80.2) [4] | 95.0 (80.3) [5] | 97.4 (96.4) [6] | 95.8 (97.9) [7] | 96.1 (87.8) [8] | 96.7 (57.1) [9] |
| Redundancy | 5.9 (4.2) [2] | 4.2 (3.6) [3] | 4.3 (3.6) [4] | 2.3 (2.1) [5] | 2.3 (2.2) [6] | 5.6 (5.7) [7] | 1.9 (1.7) [8] | 5.7 (4.2) [9] |
| $R_{merge}$ | 0.144 (0.850) [2] | 0.282 (0.997) [3] | 0.079 (0.477) [4] | 0.052 (0.669) [5] | 0.055 (0.651) [6] | 0.160 (0.980) [7] | 0.044 (0.462) [8] | 0.169 (1.117) [9] |
| $R_{pim}$ | 0.063 (0.430) [2] | 0.145 (0.580) [3] | 0.042 (0.273) [4] | 0.041 (0.553) [5] | 0.045 (0.430) [6] | 0.070 (0.418) [7] | 0.039 (0.443) [8] | 0.076 (0.583) [9] |
| $CC_{1/2}$ | 0.992 (0.590) [2] | 0.970 (0.369) [3] | 0.993 (0.820) [4] | 0.994 (0.622) [5] | 0.988 (0.541) [6] | 0.985 (0.628) [7] | 0.999 (0.723) [8] | 0.967 (0.571) [9] |
| $I/\sigma(I)$ | 13.1 (1.1) [2] | 6.2 (1.1) [3] | 13.6 (1.32) [4] | 17.5 (0.93) [5] | 23.8 (1.6) [6] | 8.0 (1.0) [7] | 15.7 (1.19) [8] | 9.2 (0.89) [9] |
| Wilson B-factors (Å$^2$) | 45.0 | 111.2 | 52.9 | 37.8 | 48.6 | 54.9 | 39.7 | 171 |
| ***Phasing*** [10] | | | | | | | | |
| Resolution (Å) | 2.56 – 32.6 | 3.60 – 52.0 | 2.50 – 30.9 | 1.95 – 45.0 | 2.20 – 48.0 | 2.80 – 50.0 | 2.10 – 50.0 | 4.45 – 44.0 |
| Correlation coefficient | 0.474 | 0.704 | 0.698 | 0.703 | 0.728 | 0.681 | 0.678 | 0.603 |
| ***Refinement*** | | | | | | | | |
| Resolution (Å) | 2.56 – 32.6 | 3.60 – 52.0 | 2.50 – 30.9 | 1.95 – 45.0 | 2.20 – 48.0 | 2.80 – 50.0 | 2.10 – 50.0 | 4.45 – 44.0 |
| No. reflections (work/test) | 28,404/1,883 | 18,260/924 | 32,298/1,627 | 68,646/3,337 | 47,957/2,189 | 26,736/1,293 | 53,156/2,635 | 12,908/559 |
| $R_{work}/R_{free}$ (%) | 22.3/27.4 | 24.2/30.0 | 21.57/26.36 | 19.4/22.8 | 19.9/25.1 | 21.5/26.3 | 21.13/24.6 | 29.4/32.9 |
| No.of atoms | | | | | | | | |
| Protein | 6,798 | 9,846 | 6,839 | 6,638 | 6,636 | 6,751 | 6,745 | 10,901 |
| Water/Others | 55/90 | 0/56 | 58/192 | 251/138 | 88/58 | 44/39 | 143/75 | 0/0 |
| B-factors (Å$^2$) | | | | | | | | |
| Protein | 67.3 | 129.7 | 74.2 | 58.3 | 75.1 | 70.3 | 65.9 | 240.6 |
| Water/Others | 47.9/66.7 | NA/133.5 | 48.8/87.5 | 47.3/69.1 | 53.7/84.0 | 47.9/77.0 | 45.2/63.1 | NA/NA |
| R.m.s deviation [11] | | | | | | | | |
| Bond length (Å) | 0.002 | 0.003 | 0.002 | 0.008 | 0.004 | 0.002 | 0.002 | 0.002 |
| Bond angle (°) | 0.564 | 0.522 | 0.533 | 0.907 | 0.596 | 0.474 | 0.504 | 0.426 |
| Ramachandran Plot (%) | | | | | | | | |
| Preferred regions | 94.63 | 92.77 | 93.20 | 96.18 | 96.59 | 94.54 | 95.66 | 91.35 |
| Allowed regions | 5.37 | 7.15 | 6.80 | 3.70 | 3.41 | 5.34 | 4.10 | 8.37 |
| Outliers | 0.00 | 0.00 | 0.00 | 0.12 | 0.00 | 0.12 | 0.24 | 0.28 |
| **PDB code** | 9NTL | 9NU7 | 9NUA | 9NV2 | 9NV6 | 9NVA | 9NVE | 9NW2 |

**Table S2. Data collection and refinement statistics for X-ray crystallography of NP TCRs and their pMHC complexes, related to Figure 3.** [1] One NP202-$NP_{366\text{-}374}/D^b$ complex and one NP202 TCR**;** [2] (Last resolution bin, 2.56 – 2.62 Å); [3] (Last resolution bin, 3.60 – 3.68 Å); [4] (Last resolution bin, 2.50 – 2.54 Å); [5] (Last resolution bin, 1.95 – 1.98 Å); [6] (Last resolution bin, 2.20 – 2.24 Å); [7] (Last resolution bin, 2.80 – 2.87 Å); [8] (Last resolution bin, 2.10 – 2.14 Å); [9] (Last resolution bin, 4.45 – 4.53 Å); [10] Molecular replacement method; [11] Root mean square Deviation.

NP202-$NP_{366-374}/D^b$ ($T_{EM}$)

| BSA (Å$^2$) | $D^b$ | $NP_{366}$ | $NP_{366}/D^b$ | $TCR_\alpha/TCR_\beta$ | Sc | $NP_{366}/D^b$ |
|---|---|---|---|---|---|---|
| $TCR_\alpha$ | 849.0 | 352.8 | 1201.8 | | $TCR_\alpha$ | 0.63 |
| $TCR_\beta$ | 643.8 | 298.0 | 941.8 | | $TCR_\beta$ | 0.54 |
| $TCR_{\alpha\beta}$ | 1492.8 | 650.8 | 2143.6 | 1.28 | $TCR_{\alpha\beta}$ | 0.61 |
| $NP_{366}$ | 1751.4 | | | | | |

NP101-$NP_{366-374}/D^b$ ($T_{BP}$)

| BSA (Å$^2$) | $D^b$ | $NP_{366}$ | $NP_{366}/D^b$ | $TCR_\alpha/TCR_\beta$ | Sc | $NP_{366}/D^b$ |
|---|---|---|---|---|---|---|
| $TCR_\alpha$ | 930.6 | 359.8 | 1290.4 | | $TCR_\alpha$ | 0.73 |
| $TCR_\beta$ | 687.0 | 282.4 | 969.4 | | $TCR_\beta$ | 0.76 |
| $TCR_{\alpha\beta}$ | 1617.6 | 642.2 | 2,259.8 | 1.33 | $TCR_{\alpha\beta}$ | 0.75 |
| $NP_{366}$ | 1710.4 | | | | | |

NP207-$NP_{366-374}/D^b$ ($T_{BP}$)

| BSA (Å$^2$) | $D^b$ | $NP_{366}$ | $NP_{366}/D^b$ | $TCR_\alpha/TCR_\beta$ | Sc | $NP_{366}/D^b$ |
|---|---|---|---|---|---|---|
| $TCR_\alpha$ | 711.8 | 376.6 | 1088.4 | | $TCR_\alpha$ | 0.63 |
| $TCR_\beta$ | 620.6 | 263.2 | 883.8 | | $TCR_\beta$ | 0.80 |
| $TCR_{\alpha\beta}$ | 1332.4 | 639.8 | 1972.2 | 1.23 | $TCR_{\alpha\beta}$ | 0.71 |
| $NP_{366}$ | 1700.6 | | | | | |

NP203-$NP_{366-374}/D^b$ ($T_{CM}$)

| BSA (Å$^2$) | $D^b$ | $NP_{366}$ | $NP_{366}/D^b$ | $TCR_\alpha/TCR_\beta$ | Sc | $NP_{366}/D^b$ |
|---|---|---|---|---|---|---|
| $TCR_\alpha$ | 858.0 | 367.8 | 1225.8 | | $TCR_\alpha$ | 0.70 |
| $TCR_\beta$ | 696.0 | 275.0 | 971.0 | | $TCR_\beta$ | 0.76 |
| $TCR_{\alpha\beta}$ | 1554.0 | 642.8 | 2196.8 | 1.26 | $TCR_{\alpha\beta}$ | 0.73 |
| $NP_{366}$ | 1643.2 | | | | | |

NP105-$NP_{366-374}/D^b$ ($T_{CM}$)

| BSA (Å$^2$) | $D^b$ | $NP_{366}$ | $NP_{366}/D^b$ | $TCR_\alpha/TCR_\beta$ | Sc | $NP_{366}/D^b$ |
|---|---|---|---|---|---|---|
| $TCR_\alpha$ | 279.8 | 44.2 | 324.0 | | $TCR_\alpha$ | 0.57 |
| $TCR_\beta$ | 769.8 | 490.8 | 1260.6 | | $TCR_\beta$ | 0.68 |
| $TCR_{\alpha\beta}$ | 1049.6 | 535.0 | 1584.6 | 0.26 | $TCR_{\alpha\beta}$ | 0.68 |
| $NP_{366}$ | 1692.0 | | | | | |

NP204-$NP_{366-374}/D^b$ ($T_{CM}$)

| BSA (Å$^2$) | $D^b$ | $NP_{366}$ | $NP_{366}/D^b$ | $TCR_\alpha/TCR_\beta$ | Sc | $NP_{366}/D^b$ |
|---|---|---|---|---|---|---|
| $TCR_\alpha$ | 344.6 | 234.0 | 578.6 | | $TCR_\alpha$ | 0.57 |
| $TCR_\beta$ | 545.0 | 437.2 | 982.2 | | $TCR_\beta$ | 0.66 |
| $TCR_{\alpha\beta}$ | 889.6 | 671.2 | 1560.8 | 0.59 | $TCR_{\alpha\beta}$ | 0.58 |
| $NP_{366}$ | 1646.4 | | | | | |

**Table S3. Buried surface area (BSA) and Surface complementarity (Sc) score across the interface between NP TCR and $NP_{366-374}/D^b$, related to Figure 3.** Individual contributions of TCR α- or β-chain to $NP_{366-374}$ peptide or H-2$D^b$ are also provided. BSA values were calculated with online PDBePISA service (https://www.ebi.ac.uk/pdbe/pisa/). Sc values were obtained by using Sc program[3] within CCP4 suite.

**A H-bond/Salt Bridge (Å)**

| TCR \ Peptide residue | p1A | p2S | p3N | p4E | p5N | p6M | p7E | p8T | p9M |
|---|---|---|---|---|---|---|---|---|---|
| **NP202** | | | K98α<br>(3.30) | G99α<br>(2.99)<br>G100α<br>(3.48)[1]<br>G101α<br>(3.05) | | | | E95β<br>(2.68) | |
| **NP101** | | | K98α<br>(2.77) | G99α<br>(2.97)<br>G101α<br>(3.13) | | | | D95β<br>(3.21) | |
| **NP207** | | | K98α<br>(2.75) | G99α<br>(2.83)<br>G101α<br>(2.97) | | | | D95β<br>(3.05) | |
| **NP203** | | | R98α<br>(3.16)<br>R98α<br>(3.25) | G99α<br>(2.83)<br>G101α<br>(3.34) | | | | D95β<br>(3.34) | |
| **NP204[2]** | | | | R97β<br>(2.81) | R97β<br>(2.83) | | S100β<br>(3.10)<br>S100β<br>(3.49) | D96β<br>(3.20) | |
| **NP105** | | | | R97β<br>(3.32) | R98β<br>(3.38) | | S100β<br>(2.77)<br>S100β<br>(2.82) | D96β<br>(2.87) | |

**B Van der Waals Contact**

| TCR \ Peptide residue | p1A | p2S | p3N | p4E | p5N | p6M | p7E | p8T | p9M |
|---|---|---|---|---|---|---|---|---|---|
| **NP202** | | | | K98α | | F33α<br>K98α<br>R99β | Y100β | | |
| **NP101** | | | | K98α | | F33α<br>Y54α<br>K98α<br>G97β<br>G98β<br>R99β | R99β<br>Y100β | | |
| **NP207** | | | | K98α | | F33α<br>Y54α<br>K98α<br>G97β<br>G98β<br>V99β | Y100β | | |
| **NP203** | | | | R98α | | F33α<br>Y54α<br>R98α<br>G98β<br>L99β | L99β<br>Y100β | | |
| **NP204[2]** | | | | R97β | | A94α<br>N98α<br>D96β | R98β | | |
| **NP105** | | | | R98β | | R98β<br>S100β | D96β<br>S99β | | |

**Table S4. TCRαβ-$NP_{366\text{-}374}/D^b$ H-bonding and Van der Waals interactions between each specified TCRαβ and the $NP_{366\text{-}374}$ peptide, related to Figure 3.** [1]Considering possible hydrogen bond geometry and potential electrostatic interaction, the interaction between p4E and G100α is weak. [2]The complex including ABCDE chains in NP204-$NP_{366\text{-}374}/D^b$ structure (PDB code: 9NU7) was used in the analysis.
