## Supplementary material for "Atomistic TCR-ligand interactions shape memory T-cell differentiation": Data S7

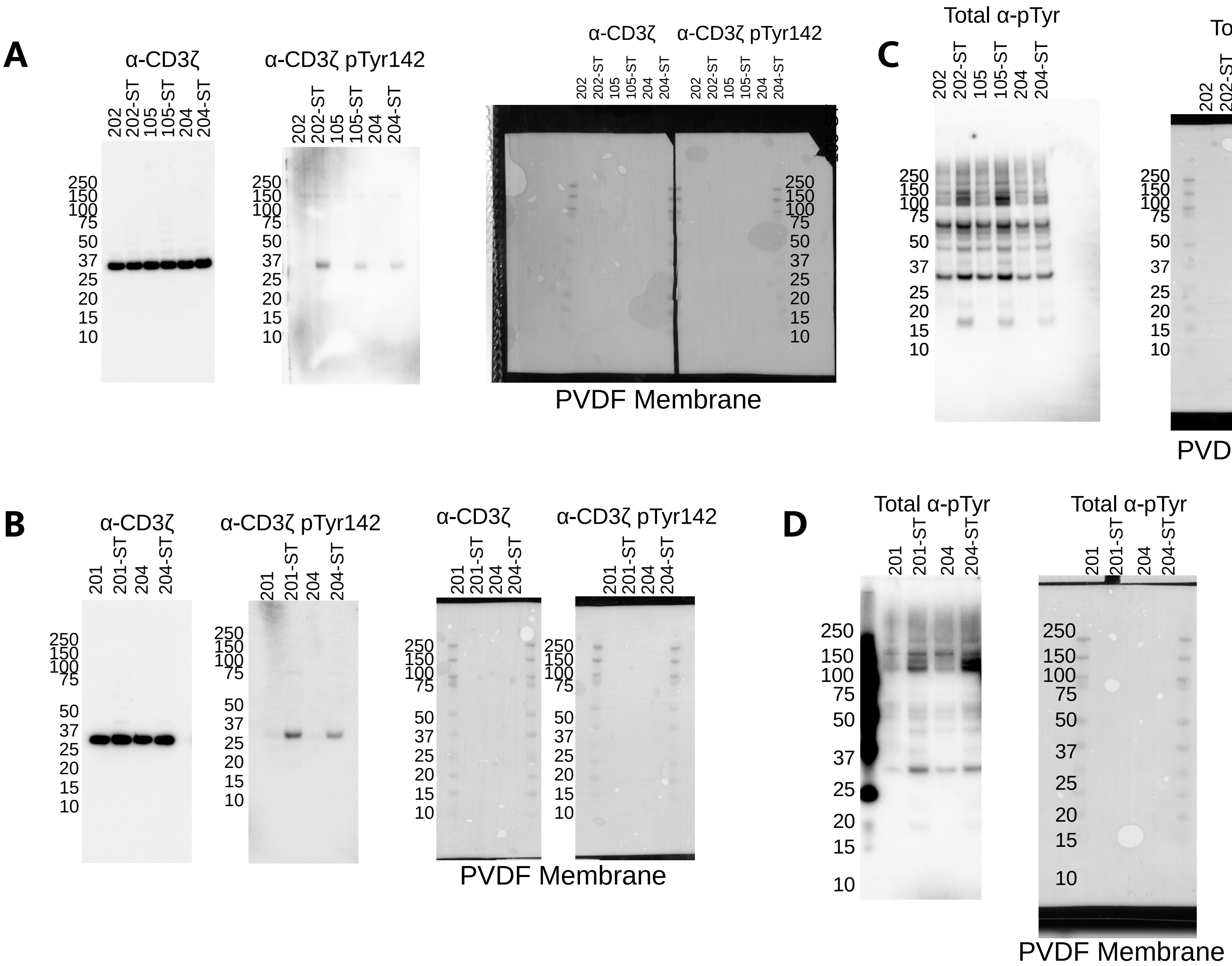


**Data S6. Original PVDF membranes and full blots, related to Figure 5. (A and B) Detection of CD3ζ and CD3ζpTyr 142 for the 202, 105 and 204 cell lysates samples (A) and the 201 and 204 cell lysate samples (B) under non-reducing conditions. The PVDF membrane used in the western blot analysis is shown, along with the molecular weight marker. The cell lysates samples were run on the same gel, and CD3ζ serves as the loading control. Western blots were cropped to the range of 50 kDa to 25 kDa. (C and D) Total pTyr detection in 202, 105 and 204 cell lysates (C) and and the 201 and 204 cell lysates (D) in reducing conditions, with the PVDF membrane and molecular weight marker also shown.**
