## Supplementary material for "Atomistic TCR-ligand interactions shape memory T-cell differentiation": Data S10

### Full wwPDB X-ray Structure Validation Report ⓘ

Mar 20, 2025 – 03:02 PM EDT

| | | |
|---|---|---|
| PDB ID | : | 9NTL |
| Title | : | Crystal structure of NP202 TCR |
| Deposited on | : | 2025-03-18 |
| Resolution | : | 2.56 Å(reported) |

**This wwPDB validation report is for manuscript review**

This is a Full wwPDB X-ray Structure Validation Report.

This report is produced by the wwPDB biocuration pipeline after annotation of the structure.

We welcome your comments at **
A user guide is available at
https://www.wwpdb.org/validation/2017/XrayValidationReportHelp
with specific help available everywhere you see the ⓘ symbol.

The types of validation reports are described at
http://www.wwpdb.org/validation/2017/FAQs#types.

The following versions of software and data (see references ⓘ) were used in the production of this report:

| | | |
|---|---|---|
| MolProbity | : | 4.02b-467 |
| Mogul | : | 2022.3.0, CSD as543be (2022) |
| Xtriage (Phenix) | : | 1.21 |
| EDS | : | 3.0 |
| buster-report | : | 1.1.7 (2018) |
| Percentile statistics | : | 20231227.v01 (using entries in the PDB archive December 27th 2023) |
| CCP4 | : | 9.0.004 (Gargrove) |
| Density-Fitness | : | 1.0.11 |
| Ideal geometry (proteins) | : | Engh & Huber (2001) |
| Ideal geometry (DNA, RNA) | : | Parkinson et al. (1996) |

For Manuscript Review

### 1 Overall quality at a glance (i)

The following experimental techniques were used to determine the structure:
*X-RAY DIFFRACTION*

The reported resolution of this entry is 2.56 Å.

Percentile scores (ranging between 0-100) for global validation metrics of the entry are shown in the following graphic. The table shows the number of entries on which the scores are based.

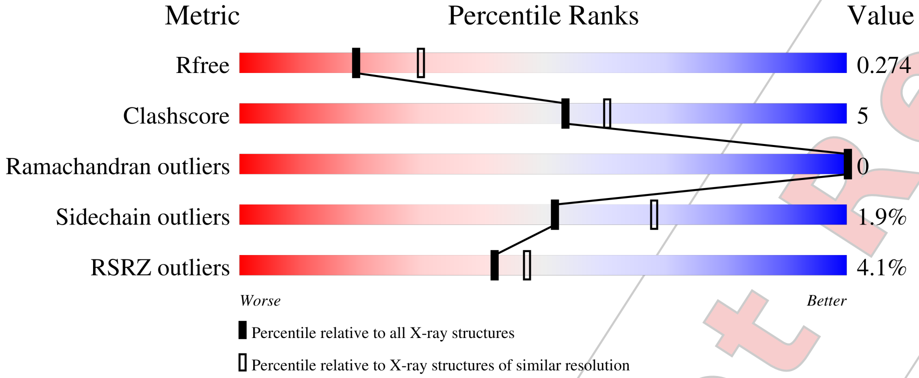


| Metric | Whole archive (#Entries) | Similar resolution (#Entries, resolution range(Å)) |
|---|---|---|
| $R_{free}$ | 164625 | 1685 (2.58-2.54) |
| Clashscore | 180529 | 1779 (2.58-2.54) |
| Ramachandran outliers | 177936 | 1766 (2.58-2.54) |
| Sidechain outliers | 177891 | 1766 (2.58-2.54) |
| RSRZ outliers | 164620 | 1685 (2.58-2.54) |

The table below summarises the geometric issues observed across the polymeric chains and their fit to the electron density. The red, orange, yellow and green segments of the lower bar indicate the fraction of residues that contain outliers for >=3, 2, 1 and 0 types of geometric quality criteria respectively. A grey segment represents the fraction of residues that are not modelled. The numeric value for each fraction is indicated below the corresponding segment, with a dot representing fractions <=5% The upper red bar (where present) indicates the fraction of residues that have poor fit to the electron density. The numeric value is given above the bar.

| Mol | Chain | Length | Quality of chain |
|---|---|---|---|
| 1 | A | 219 | 3% / 82% 11% 7% |
| 1 | C | 219 | 7% / 73% 16% 11% |
| 2 | B | 250 | 83% 10% • 5% |
| 2 | D | 250 | 5% / 84% 10% 6% |

Validation Pipeline (wwPDB-VP) : 2.41.5

For Manuscript Review

### 2 Entry composition (i)

There are 5 unique types of molecules in this entry. The entry contains 6943 atoms, of which 0 are hydrogens and 0 are deuteriums.

In the tables below, the ZeroOcc column contains the number of atoms modelled with zero occupancy, the AltConf column contains the number of residues with at least one atom in alternate conformation and the Trace column contains the number of residues modelled with at most 2 atoms.

- Molecule 1 is a protein called NP202 TCR alpha chain.

| Mol | Chain | Residues | Atoms | | | | | ZeroOcc | AltConf | Trace |
|---|---|---|---|---|---|---|---|---|---|---|
| 1 | A | 204 | Total<br>1583 | C<br>1002 | N<br>254 | O<br>316 | S<br>11 | 0 | 0 | 0 |
| 1 | C | 195 | Total<br>1476 | C<br>934 | N<br>236 | O<br>297 | S<br>9 | 0 | 0 | 0 |

- Molecule 2 is a protein called NP202 TCR beta chain.

| Mol | Chain | Residues | Atoms | | | | | ZeroOcc | AltConf | Trace |
|---|---|---|---|---|---|---|---|---|---|---|
| 2 | B | 237 | Total<br>1900 | C<br>1208 | N<br>331 | O<br>354 | S<br>7 | 0 | 0 | 0 |
| 2 | D | 236 | Total<br>1839 | C<br>1172 | N<br>320 | O<br>339 | S<br>8 | 0 | 0 | 0 |

- Molecule 3 is 2-acetamido-2-deoxy-beta-D-glucopyranose (three-letter code: NAG) (formula: $C_8H_{15}NO_6$) (labeled as "Ligand of Interest" by depositor).

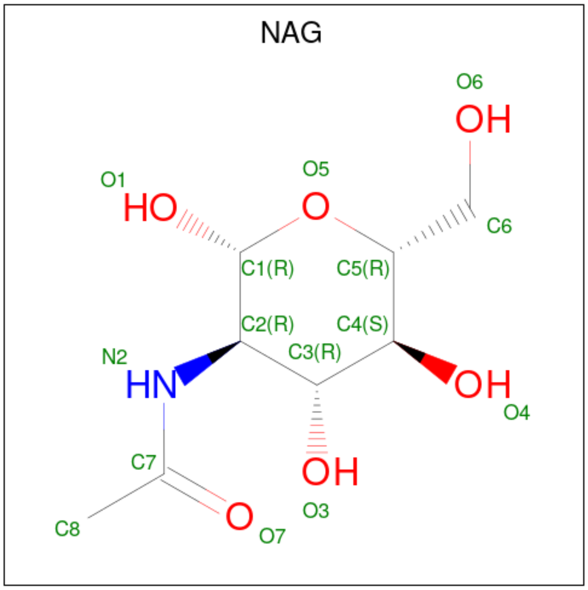


For Manuscript Review

| Mol | Chain | Residues | Atoms | ZeroOcc | AltConf |
|---|---|---|---|---|---|
| 3 | A | 1 | Total C N O<br>14 8 1 5 | 0 | 0 |
| 3 | B | 1 | Total C N O<br>14 8 1 5 | 0 | 0 |
| 3 | B | 1 | Total C N O<br>14 8 1 5 | 0 | 0 |
| 3 | B | 1 | Total C N O<br>14 8 1 5 | 0 | 0 |
| 3 | D | 1 | Total C N O<br>14 8 1 5 | 0 | 0 |
| 3 | D | 1 | Total C N O<br>14 8 1 5 | 0 | 0 |

- Molecule 4 is GLYCEROL (three-letter code: GOL) (formula: $C_3H_8O_3$).

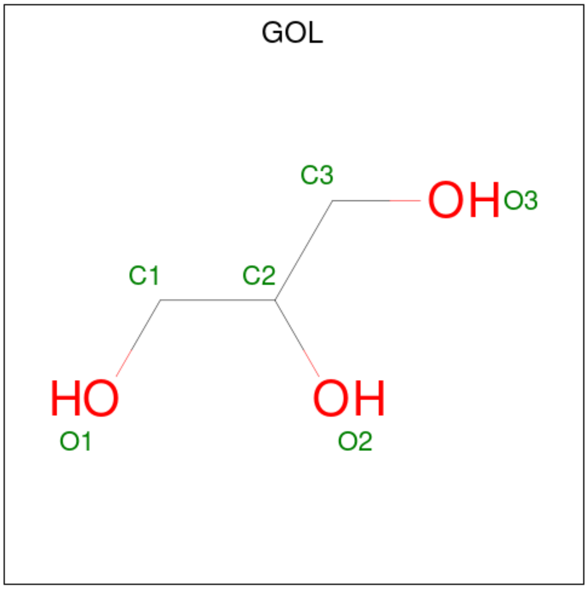


| Mol | Chain | Residues | Atoms | ZeroOcc | AltConf |
|---|---|---|---|---|---|
| 4 | D | 1 | Total C O<br>6 3 3 | 0 | 0 |

- Molecule 5 is water.

| Mol | Chain | Residues | Atoms | ZeroOcc | AltConf |
|---|---|---|---|---|---|
| 5 | A | 28 | Total O<br>28 28 | 0 | 0 |
| 5 | B | 17 | Total O<br>17 17 | 0 | 0 |
| 5 | C | 4 | Total O<br>4 4 | 0 | 0 |

*Continued on next page...*

For Manuscript Review

*Continued from previous page...*

| Mol | Chain | Residues | Atoms | | ZeroOcc | AltConf |
|---|---|---|---|---|---|---|
| 5 | D | 6 | Total<br>6 | O<br>6 | 0 | 0 |

For Manuscript Review

### 3 Residue-property plots (i)

These plots are drawn for all protein, RNA, DNA and oligosaccharide chains in the entry. The first graphic for a chain summarises the proportions of the various outlier classes displayed in the second graphic. The second graphic shows the sequence view annotated by issues in geometry and electron density. Residues are color-coded according to the number of geometric quality criteria for which they contain at least one outlier: green = 0, yellow = 1, orange = 2 and red = 3 or more. A red dot above a residue indicates a poor fit to the electron density (RSRZ > 2). Stretches of 2 or more consecutive residues without any outlier are shown as a green connector. Residues present in the sample, but not in the model, are shown in grey.

- Molecule 1: NP202 TCR alpha chain

Chain A:

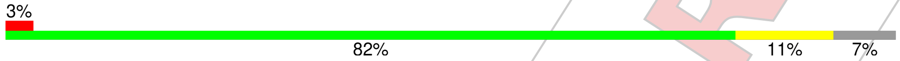


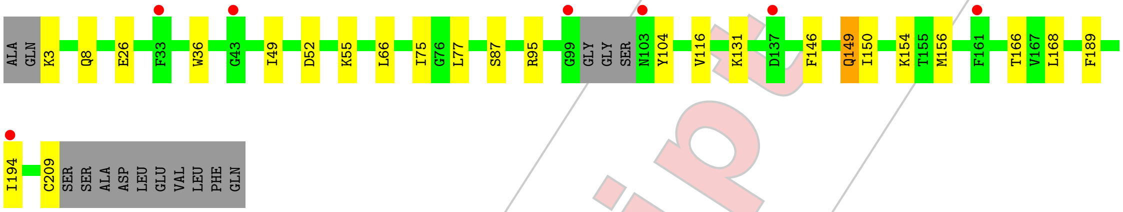


- Molecule 1: NP202 TCR alpha chain

Chain C:

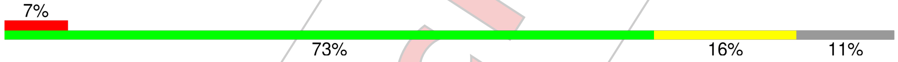


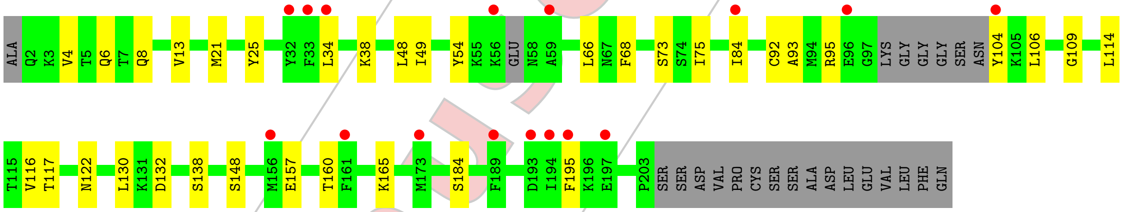


- Molecule 2: NP202 TCR beta chain

Chain B:

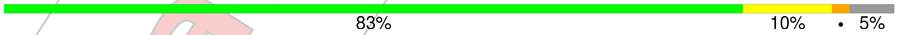


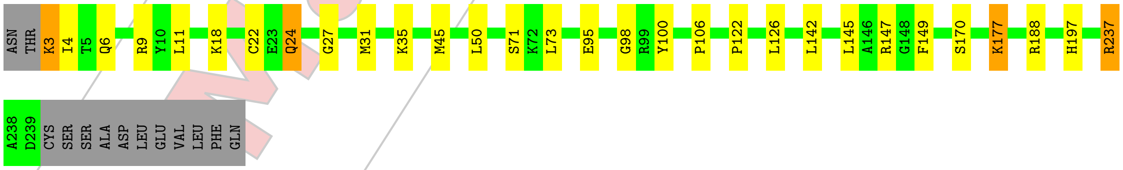


- Molecule 2: NP202 TCR beta chain

Chain D:

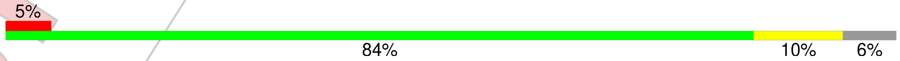


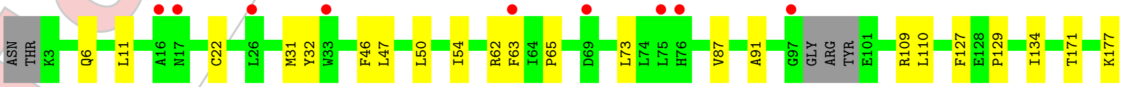


For Manuscript Review

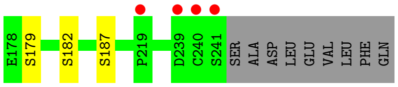


For Manuscript Review

### 4 Data and refinement statistics (i)

| Property | Value | Source |
|---|---|---|
| Space group | P 1 21 1 | Depositor |
| Cell constants<br>a, b, c, $\alpha$, $\beta$, $\gamma$ | 62.07Å 127.39Å 66.56Å<br>90.00° 110.11° 90.00° | Depositor |
| Resolution (Å) | 32.60 – 2.56<br>32.60 – 2.56 | Depositor<br>EDS |
| % Data completeness<br>(in resolution range) | 90.8 (32.60-2.56)<br>90.8 (32.60-2.56) | Depositor<br>EDS |
| $R_{merge}$ | 0.14 | Depositor |
| $R_{sym}$ | (Not available) | Depositor |
| $< I/\sigma(I) >$ [1] | 1.85 (at 2.54Å) | Xtriage |
| Refinement program | PHENIX (1.21.2_5419: ???) | Depositor |
| R, $R_{free}$ | 0.223 , 0.274<br>0.223 , 0.274 | Depositor<br>DCC |
| $R_{free}$ test set | 29297 reflections (6.63%) | wwPDB-VP |
| Wilson B-factor (Å$^2$) | 44.9 | Xtriage |
| Anisotropy | 0.821 | Xtriage |
| Bulk solvent $k_{sol}$(e/Å$^3$), $B_{sol}$(Å$^2$) | 0.33 , 57.6 | EDS |
| L-test for twinning[2] | $< \|L\| >= 0.48$, $< L^2 >= 0.31$ | Xtriage |
| Estimated twinning fraction | No twinning to report. | Xtriage |
| $F_o$,$F_c$ correlation | 0.92 | EDS |
| Total number of atoms | 6943 | wwPDB-VP |
| Average B, all atoms (Å$^2$) | 67.0 | wwPDB-VP |

Xtriage's analysis on translational NCS is as follows: *The largest off-origin peak in the Patterson function is 7.04% of the height of the origin peak. No significant pseudotranslation is detected.*

[1] Intensities estimated from amplitudes.

[2] Theoretical values of $< |L| >$, $< L^2 >$ for acentric reflections are 0.5, 0.333 respectively for untwinned datasets, and 0.375, 0.2 for perfectly twinned datasets.

For Manuscript Review

### 5 Model quality (i)

#### 5.1 Standard geometry (i)

Bond lengths and bond angles in the following residue types are not validated in this section: GOL, NAG

The Z score for a bond length (or angle) is the number of standard deviations the observed value is removed from the expected value. A bond length (or angle) with $|Z| > 5$ is considered an outlier worth inspection. RMSZ is the root-mean-square of all Z scores of the bond lengths (or angles).

| Mol | Chain | Bond lengths | | Bond angles | |
|---|---|---|---|---|---|
| | | RMSZ | $\#\|Z\| > 5$ | RMSZ | $\#\|Z\| > 5$ |
| 1 | A | 0.26 | 0/1614 | 0.50 | 0/2188 |
| 1 | C | 0.26 | 0/1503 | 0.51 | 0/2043 |
| 2 | B | 0.26 | 0/1956 | 0.50 | 0/2658 |
| 2 | D | 0.25 | 0/1893 | 0.49 | 0/2580 |
| All | All | 0.26 | 0/6966 | 0.50 | 0/9469 |

There are no bond length outliers.

There are no bond angle outliers.

There are no chirality outliers.

There are no planarity outliers.

#### 5.2 Too-close contacts (i)

In the following table, the Non-H and H(model) columns list the number of non-hydrogen atoms and hydrogen atoms in the chain respectively. The H(added) column lists the number of hydrogen atoms added and optimized by MolProbity. The Clashes column lists the number of clashes within the asymmetric unit, whereas Symm-Clashes lists symmetry-related clashes.

| Mol | Chain | Non-H | H(model) | H(added) | Clashes | Symm-Clashes |
|---|---|---|---|---|---|---|
| 1 | A | 1583 | 0 | 1529 | 18 | 0 |
| 1 | C | 1476 | 0 | 1385 | 20 | 0 |
| 2 | B | 1900 | 0 | 1825 | 21 | 0 |
| 2 | D | 1839 | 0 | 1728 | 14 | 0 |
| 3 | A | 14 | 0 | 13 | 0 | 0 |
| 3 | B | 42 | 0 | 39 | 0 | 0 |
| 3 | D | 28 | 0 | 26 | 0 | 0 |
| 4 | D | 6 | 0 | 8 | 0 | 0 |
| 5 | A | 28 | 0 | 0 | 0 | 0 |

*Continued on next page...*

For Manuscript Review

*Continued from previous page...*

| Mol | Chain | Non-H | H(model) | H(added) | Clashes | Symm-Clashes |
|---|---|---|---|---|---|---|
| 5 | B | 17 | 0 | 0 | 0 | 0 |
| 5 | C | 4 | 0 | 0 | 0 | 0 |
| 5 | D | 6 | 0 | 0 | 0 | 0 |
| All | All | 6943 | 0 | 6553 | 65 | 0 |

The all-atom clashscore is defined as the number of clashes found per 1000 atoms (including hydrogen atoms). The all-atom clashscore for this structure is 5.

All (65) close contacts within the same asymmetric unit are listed below, sorted by their clash magnitude.

| Atom-1 | Atom-2 | Interatomic distance (Å) | Clash overlap (Å) |
|---|---|---|---|
| 1:A:95:ARG:HH21 | 1:A:104:TYR:HB2 | 1.44 | 0.83 |
| 2:D:32:TYR:HB2 | 2:D:91:ALA:HB3 | 1.72 | 0.70 |
| 1:A:168:LEU:HB3 | 2:B:170:SER:HB2 | 1.73 | 0.69 |
| 1:C:8:GLN:HE22 | 1:C:21:MET:HA | 1.60 | 0.67 |
| 1:C:84:ILE:HD13 | 1:C:116:VAL:HG12 | 1.77 | 0.66 |
| 2:B:22:CYS:HB3 | 2:B:73:LEU:HB2 | 1.77 | 0.66 |
| 1:A:166:THR:HG23 | 2:B:188:ARG:HH11 | 1.61 | 0.65 |
| 2:B:11:LEU:HD11 | 2:B:18:LYS:HD3 | 1.81 | 0.63 |
| 2:D:129:PRO:HB2 | 2:D:134:ILE:HD11 | 1.83 | 0.59 |
| 1:C:21:MET:HE1 | 1:C:114:LEU:HD22 | 1.84 | 0.59 |
| 2:B:122:PRO:HB3 | 2:B:149:PHE:HB3 | 1.86 | 0.56 |
| 1:C:157:GLU:HB3 | 1:C:160:THR:HB | 1.88 | 0.56 |
| 1:C:6:GLN:NE2 | 1:C:92:CYS:H | 2.03 | 0.56 |
| 1:A:66:LEU:HD13 | 1:A:75:ILE:HD11 | 1.89 | 0.54 |
| 1:C:49:ILE:HD13 | 1:C:66:LEU:HG | 1.89 | 0.54 |
| 1:A:3:LYS:HB3 | 1:A:26:GLU:HB3 | 1.90 | 0.54 |
| 1:C:117:THR:HB | 1:C:148:SER:HB3 | 1.90 | 0.53 |
| 1:C:132:ASP:HB3 | 1:C:138:SER:HB3 | 1.91 | 0.53 |
| 1:C:34:LEU:HD22 | 1:C:75:ILE:HG21 | 1.93 | 0.50 |
| 1:A:8:GLN:HG2 | 1:C:13:VAL:HG12 | 1.93 | 0.50 |
| 1:C:95:ARG:HH12 | 1:C:104:TYR:HA | 1.77 | 0.50 |
| 1:A:95:ARG:HD3 | 2:B:98:GLY:HA2 | 1.95 | 0.49 |
| 1:A:49:ILE:HD13 | 1:A:66:LEU:HG | 1.94 | 0.49 |
| 2:D:62:ARG:HG3 | 2:D:63:PHE:CD1 | 2.48 | 0.49 |
| 1:C:38:LYS:HB3 | 1:C:48:LEU:HD21 | 1.96 | 0.48 |
| 2:D:171:THR:HG23 | 2:D:187:SER:HB2 | 1.95 | 0.48 |
| 2:B:35:LYS:HB2 | 2:B:45:MET:SD | 2.53 | 0.48 |
| 1:A:146:PHE:CD2 | 1:A:150:ILE:HD11 | 2.49 | 0.47 |
| 2:D:62:ARG:HG3 | 2:D:63:PHE:HD1 | 1.80 | 0.46 |
| 1:C:130:LEU:HB3 | 2:D:127:PHE:HB3 | 1.96 | 0.46 |

*Continued on next page...*

For Manuscript Review

*Continued from previous page...*

| Atom-1 | Atom-2 | Interatomic distance (Å) | Clash overlap (Å) |
|---|---|---|---|
| 2:B:126:LEU:HD11 | 2:B:142:LEU:HD23 | 1.98 | 0.46 |
| 2:B:177:LYS:H | 2:B:177:LYS:HG3 | 1.51 | 0.46 |
| 2:B:3:LYS:HG3 | 2:B:4:ILE:H | 1.82 | 0.45 |
| 2:D:87:VAL:HG22 | 2:D:109:ARG:HA | 1.97 | 0.45 |
| 1:C:6:GLN:HE21 | 1:C:109:GLY:HA3 | 1.81 | 0.45 |
| 2:B:24:GLN:O | 2:B:71:SER:HB2 | 2.17 | 0.45 |
| 1:C:117:THR:HG22 | 1:C:165:LYS:HE2 | 1.98 | 0.44 |
| 1:A:168:LEU:CB | 2:B:170:SER:HB2 | 2.43 | 0.44 |
| 2:B:6:GLN:HB3 | 2:B:106:PRO:HG2 | 2.00 | 0.44 |
| 2:D:47:LEU:HD22 | 2:D:54:ILE:HD11 | 2.00 | 0.44 |
| 1:A:131:LYS:HD3 | 1:A:131:LYS:HA | 1.69 | 0.43 |
| 1:A:168:LEU:HD23 | 2:B:188:ARG:CZ | 2.48 | 0.43 |
| 1:C:160:THR:HA | 1:C:184:SER:HA | 2.00 | 0.43 |
| 1:A:52:ASP:HB2 | 1:A:55:LYS:HG3 | 1.99 | 0.43 |
| 2:B:197:HIS:HA | 2:B:237:ARG:O | 2.19 | 0.43 |
| 1:A:189:PHE:HB3 | 1:A:194:ILE:HD11 | 2.00 | 0.43 |
| 1:C:25:TYR:CZ | 1:C:73:SER:HA | 2.55 | 0.42 |
| 1:A:156:MET:SD | 2:D:177:LYS:HG2 | 2.59 | 0.42 |
| 1:C:6:GLN:HE22 | 1:C:92:CYS:H | 1.66 | 0.42 |
| 2:D:179:SER:HB2 | 2:D:182:SER:HB3 | 2.01 | 0.42 |
| 1:A:87:SER:HB3 | 1:A:116:VAL:H | 1.85 | 0.42 |
| 2:B:95:GLU:HB2 | 2:B:100:TYR:CE1 | 2.54 | 0.42 |
| 2:D:46:PHE:CD1 | 2:D:65:PRO:HG3 | 2.55 | 0.42 |
| 1:A:149:GLN:HE21 | 1:A:149:GLN:HB3 | 1.66 | 0.42 |
| 2:B:31:MET:HB3 | 2:B:73:LEU:HD11 | 2.02 | 0.42 |
| 1:C:93:ALA:HB1 | 1:C:106:LEU:HD11 | 2.02 | 0.41 |
| 2:B:145:LEU:HG | 2:B:147:ARG:HG3 | 2.02 | 0.41 |
| 1:C:4:VAL:HG11 | 1:C:92:CYS:O | 2.20 | 0.41 |
| 2:D:11:LEU:HD22 | 2:D:110:LEU:HG | 2.01 | 0.41 |
| 2:B:24:GLN:NE2 | 2:B:27:GLY:H | 2.19 | 0.41 |
| 1:A:36:TRP:CE2 | 1:A:77:LEU:HB2 | 2.55 | 0.41 |
| 2:D:6:GLN:HG3 | 2:D:22:CYS:HB2 | 2.03 | 0.41 |
| 2:D:31:MET:HB3 | 2:D:73:LEU:HD11 | 2.02 | 0.41 |
| 2:B:3:LYS:HD2 | 2:B:3:LYS:HA | 1.85 | 0.41 |
| 2:B:122:PRO:CB | 2:B:149:PHE:HB3 | 2.52 | 0.40 |

There are no symmetry-related clashes.

For Manuscript Review

#### 5.3 Torsion angles (i)

##### 5.3.1 Protein backbone (i)

In the following table, the Percentiles column shows the percent Ramachandran outliers of the chain as a percentile score with respect to all X-ray entries followed by that with respect to entries of similar resolution.

The Analysed column shows the number of residues for which the backbone conformation was analysed, and the total number of residues.

| Mol | Chain | Analysed | Favoured | Allowed | Outliers | Percentiles | |
|---|---|---|---|---|---|---|---|
| 1 | A | 200/219 (91%) | 188 (94%) | 12 (6%) | 0 | 100 | 100 |
| 1 | C | 189/219 (86%) | 181 (96%) | 8 (4%) | 0 | 100 | 100 |
| 2 | B | 235/250 (94%) | 227 (97%) | 8 (3%) | 0 | 100 | 100 |
| 2 | D | 232/250 (93%) | 214 (92%) | 18 (8%) | 0 | 100 | 100 |
| All | All | 856/938 (91%) | 810 (95%) | 46 (5%) | 0 | 100 | 100 |

There are no Ramachandran outliers to report.

##### 5.3.2 Protein sidechains (i)

In the following table, the Percentiles column shows the percent sidechain outliers of the chain as a percentile score with respect to all X-ray entries followed by that with respect to entries of similar resolution.

The Analysed column shows the number of residues for which the sidechain conformation was analysed, and the total number of residues.

| Mol | Chain | Analysed | Rotameric | Outliers | Percentiles | |
|---|---|---|---|---|---|---|
| 1 | A | 178/196 (91%) | 175 (98%) | 3 (2%) | 56 | 71 |
| 1 | C | 159/196 (81%) | 155 (98%) | 4 (2%) | 42 | 58 |
| 2 | B | 208/222 (94%) | 202 (97%) | 6 (3%) | 37 | 52 |
| 2 | D | 194/222 (87%) | 193 (100%) | 1 (0%) | 86 | 92 |
| All | All | 739/836 (88%) | 725 (98%) | 14 (2%) | 52 | 68 |

All (14) residues with a non-rotameric sidechain are listed below:

| Mol | Chain | Res | Type |
|---|---|---|---|
| 1 | A | 149 | GLN |
| 1 | A | 154 | LYS |

*Continued on next page...*

For Manuscript Review

*Continued from previous page...*

| Mol | Chain | Res | Type |
|---|---|---|---|
| 1 | A | 209 | CYS |
| 2 | B | 3 | LYS |
| 2 | B | 9 | ARG |
| 2 | B | 24 | GLN |
| 2 | B | 50 | LEU |
| 2 | B | 177 | LYS |
| 2 | B | 237 | ARG |
| 1 | C | 54 | TYR |
| 1 | C | 68 | PHE |
| 1 | C | 122 | ASN |
| 1 | C | 195 | PHE |
| 2 | D | 50 | LEU |

Sometimes sidechains can be flipped to improve hydrogen bonding and reduce clashes. All (7) such sidechains are listed below:

| Mol | Chain | Res | Type |
|---|---|---|---|
| 1 | A | 63 | HIS |
| 1 | A | 149 | GLN |
| 2 | B | 24 | GLN |
| 2 | B | 174 | GLN |
| 1 | C | 6 | GLN |
| 1 | C | 8 | GLN |
| 2 | D | 49 | ASN |

##### 5.3.3 RNA (i)

There are no RNA molecules in this entry.

#### 5.4 Non-standard residues in protein, DNA, RNA chains (i)

There are no non-standard protein/DNA/RNA residues in this entry.

#### 5.5 Carbohydrates (i)

There are no oligosaccharides in this entry.

#### 5.6 Ligand geometry (i)

7 ligands are modelled in this entry.

For Manuscript Review

In the following table, the Counts columns list the number of bonds (or angles) for which Mogul statistics could be retrieved, the number of bonds (or angles) that are observed in the model and the number of bonds (or angles) that are defined in the Chemical Component Dictionary. The Link column lists molecule types, if any, to which the group is linked. The Z score for a bond length (or angle) is the number of standard deviations the observed value is removed from the expected value. A bond length (or angle) with $|Z| > 2$ is considered an outlier worth inspection. RMSZ is the root-mean-square of all Z scores of the bond lengths (or angles).

| Mol | Type | Chain | Res | Link | Bond lengths | | | Bond angles | | |
|---|---|---|---|---|---|---|---|---|---|---|
| | | | | | Counts | RMSZ | #$\|Z\| > 2$ | Counts | RMSZ | #$\|Z\| > 2$ |
| 3 | NAG | B | 302 | 2 | 14,14,15 | 0.73 | 0 | 17,19,21 | 0.80 | 0 |
| 4 | GOL | D | 303 | - | 5,5,5 | 0.35 | 0 | 5,5,5 | 0.49 | 0 |
| 3 | NAG | B | 303 | 2 | 14,14,15 | 0.71 | 0 | 17,19,21 | 0.78 | 0 |
| 3 | NAG | B | 301 | 2 | 14,14,15 | 0.72 | 0 | 17,19,21 | 0.99 | 1 (5%) |
| 3 | NAG | A | 301 | 1 | 14,14,15 | 0.74 | 0 | 17,19,21 | 0.95 | 0 |
| 3 | NAG | D | 301 | 2 | 14,14,15 | 0.84 | 0 | 17,19,21 | 1.57 | 2 (11%) |
| 3 | NAG | D | 302 | 2 | 14,14,15 | 0.73 | 0 | 17,19,21 | 0.88 | 0 |

In the following table, the Chirals column lists the number of chiral outliers, the number of chiral centers analysed, the number of these observed in the model and the number defined in the Chemical Component Dictionary. Similar counts are reported in the Torsion and Rings columns. '-' means no outliers of that kind were identified.

| Mol | Type | Chain | Res | Link | Chirals | Torsions | Rings |
|---|---|---|---|---|---|---|---|
| 3 | NAG | B | 302 | 2 | - | 0/6/23/26 | 0/1/1/1 |
| 4 | GOL | D | 303 | - | - | 0/4/4/4 | - |
| 3 | NAG | B | 303 | 2 | - | 0/6/23/26 | 0/1/1/1 |
| 3 | NAG | B | 301 | 2 | - | 2/6/23/26 | 0/1/1/1 |
| 3 | NAG | A | 301 | 1 | - | 2/6/23/26 | 0/1/1/1 |
| 3 | NAG | D | 301 | 2 | - | 0/6/23/26 | 0/1/1/1 |
| 3 | NAG | D | 302 | 2 | - | 2/6/23/26 | 0/1/1/1 |

There are no bond length outliers.

All (3) bond angle outliers are listed below:

| Mol | Chain | Res | Type | Atoms | Z | Observed($^o$) | Ideal($^o$) |
|---|---|---|---|---|---|---|---|
| 3 | D | 301 | NAG | C1-O5-C5 | 4.99 | 118.87 | 112.19 |
| 3 | D | 301 | NAG | C2-N2-C7 | 2.64 | 126.44 | 122.90 |
| 3 | B | 301 | NAG | C1-O5-C5 | 2.08 | 114.97 | 112.19 |

There are no chirality outliers.

All (6) torsion outliers are listed below:

For Manuscript Review

| Mol | Chain | Res | Type | Atoms |
|---|---|---|---|---|
| 3 | B | 301 | NAG | C4-C5-C6-O6 |
| 3 | B | 301 | NAG | O5-C5-C6-O6 |
| 3 | D | 302 | NAG | O5-C5-C6-O6 |
| 3 | A | 301 | NAG | O5-C5-C6-O6 |
| 3 | A | 301 | NAG | C4-C5-C6-O6 |
| 3 | D | 302 | NAG | C4-C5-C6-O6 |

There are no ring outliers.

No monomer is involved in short contacts.

The following is a two-dimensional graphical depiction of Mogul quality analysis of bond lengths, bond angles, torsion angles, and ring geometry for all instances of the Ligand of Interest. In addition, ligands with molecular weight > 250 and outliers as shown on the validation Tables will also be included. For torsion angles, if less then 5% of the Mogul distribution of torsion angles is within 10 degrees of the torsion angle in question, then that torsion angle is considered an outlier. Any bond that is central to one or more torsion angles identified as an outlier by Mogul will be highlighted in the graph. For rings, the root-mean-square deviation (RMSD) between the ring in question and similar rings identified by Mogul is calculated over all ring torsion angles. If the average RMSD is greater than 60 degrees and the minimal RMSD between the ring in question and any Mogul-identified rings is also greater than 60 degrees, then that ring is considered an outlier. The outliers are highlighted in purple. The color gray indicates Mogul did not find sufficient equivalents in the CSD to analyse the geometry.

**Ligand NAG B 302**

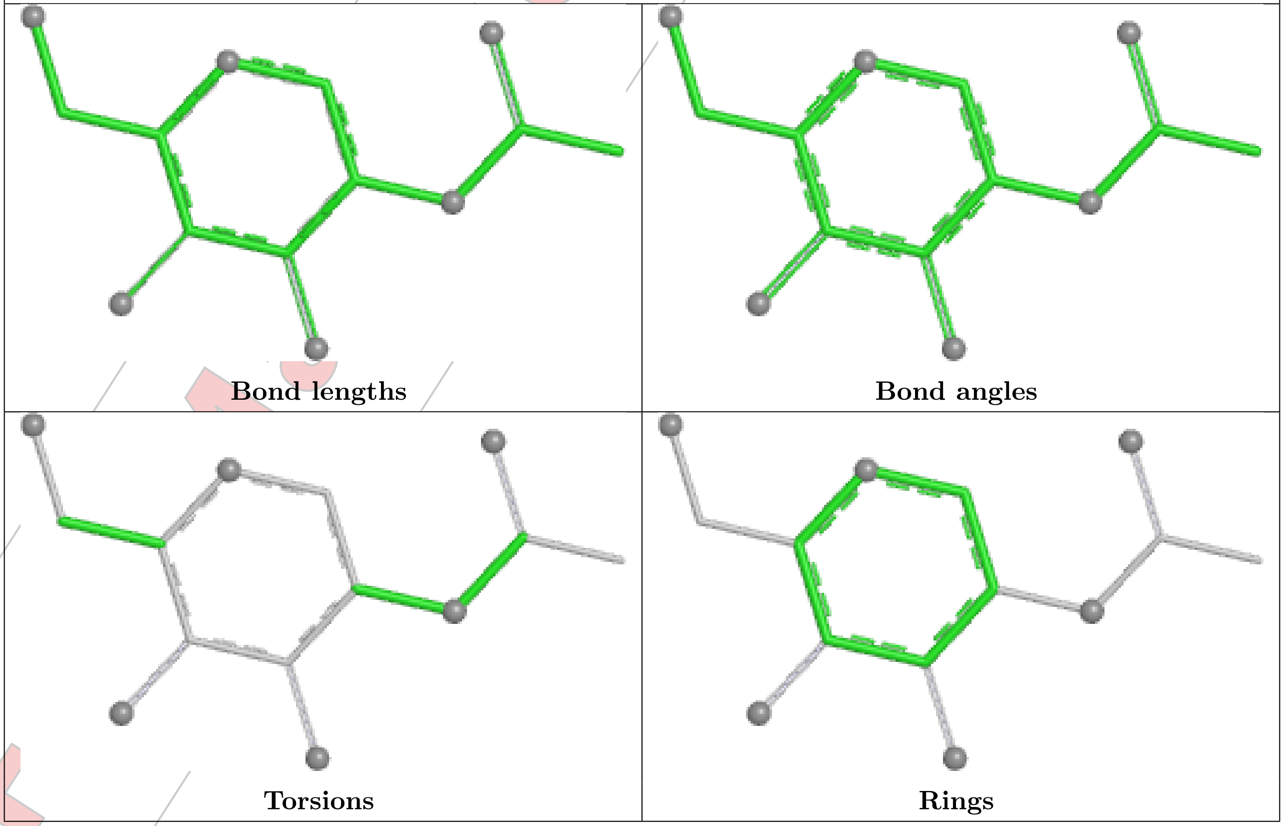


For Manuscript Review

#### Ligand NAG B 303

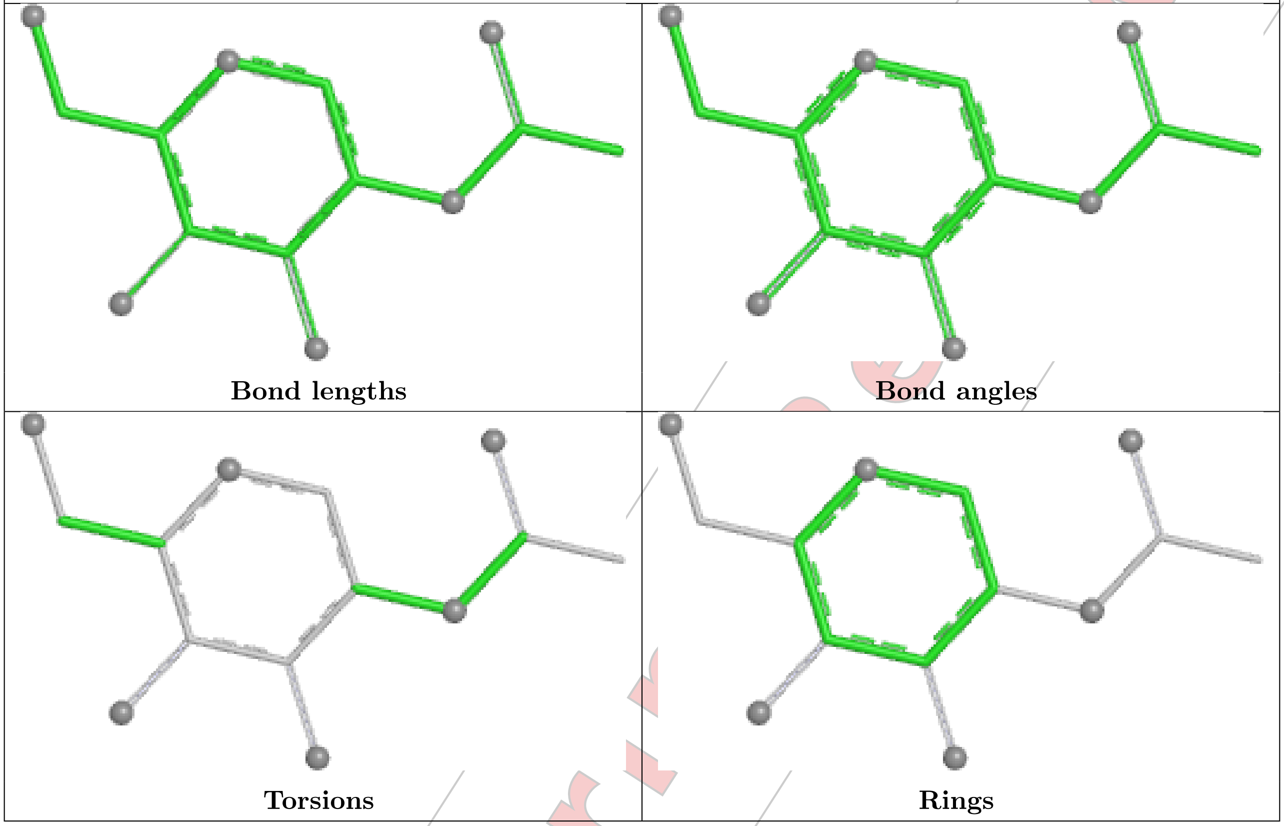


#### Ligand NAG B 301

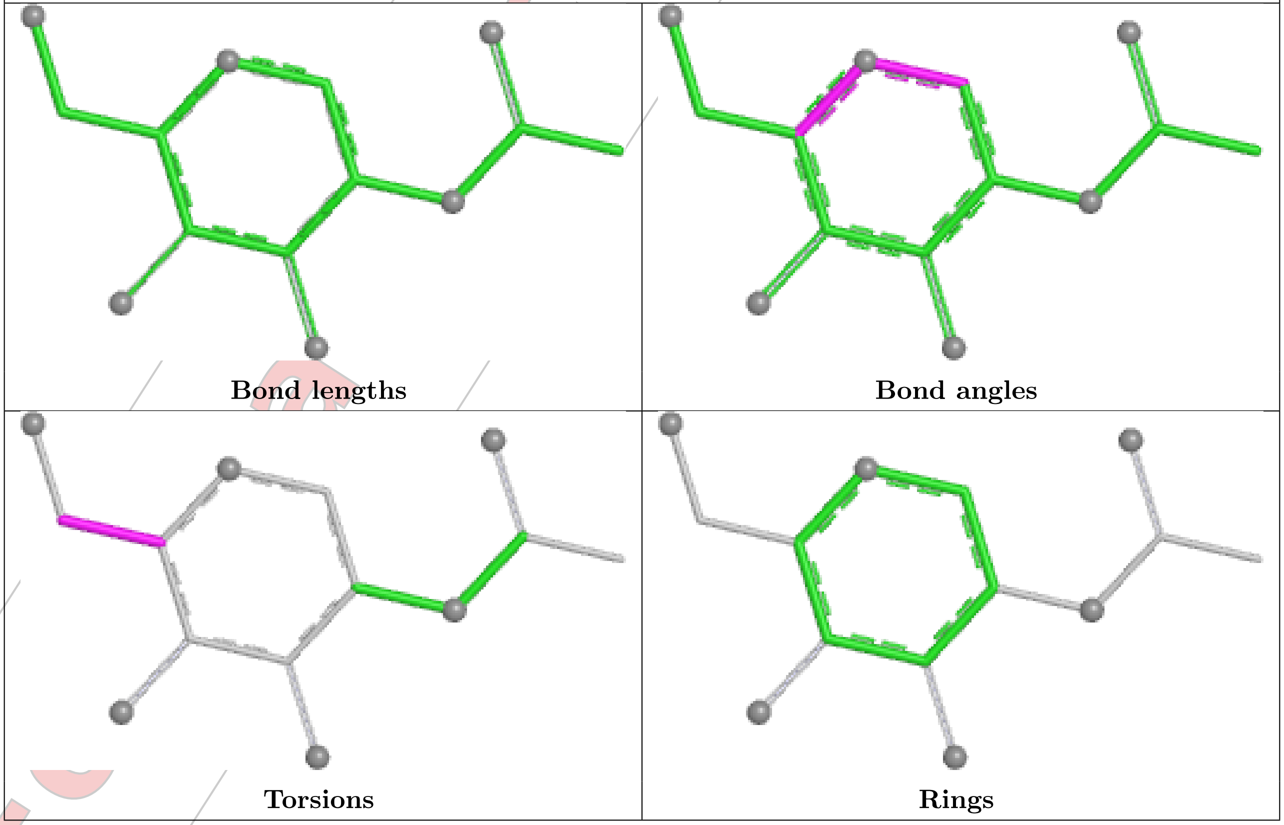

##### Ligand NAG A 301

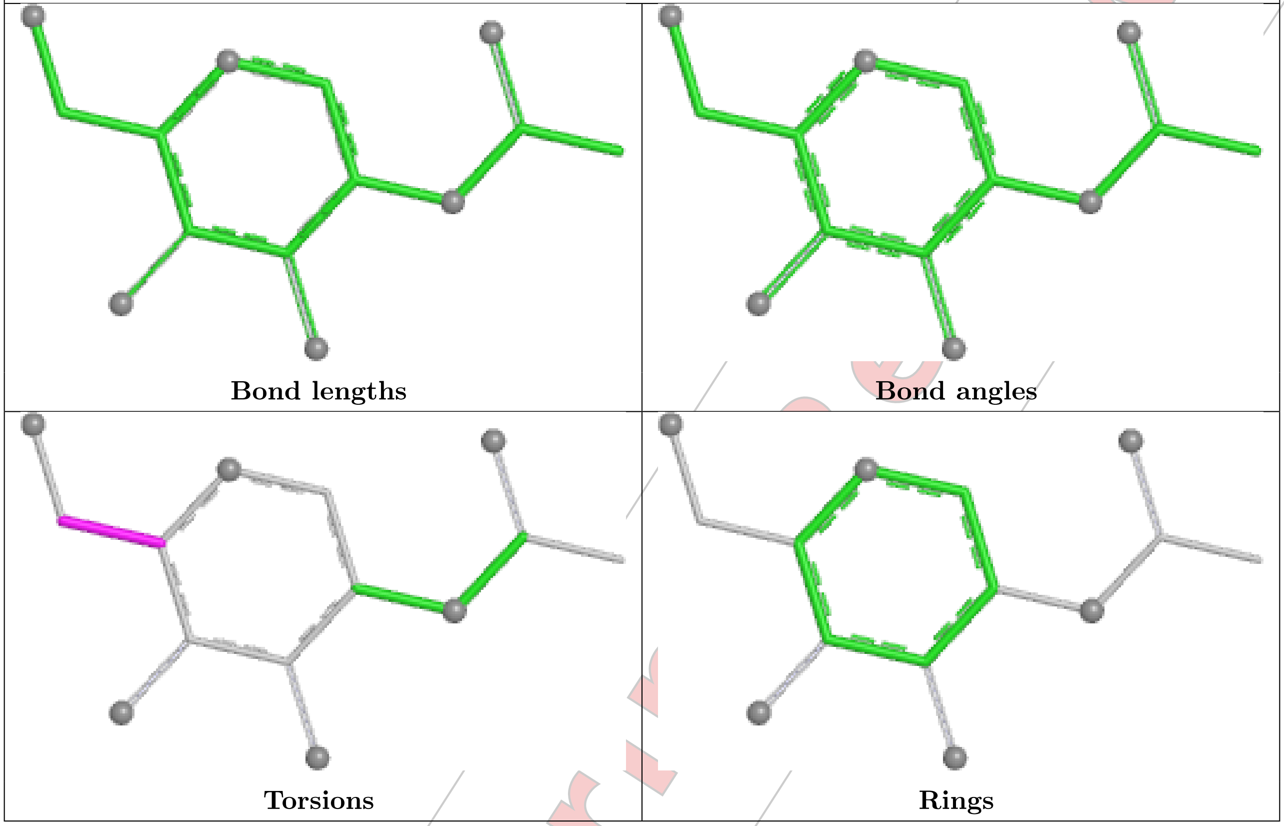


##### Ligand NAG D 301

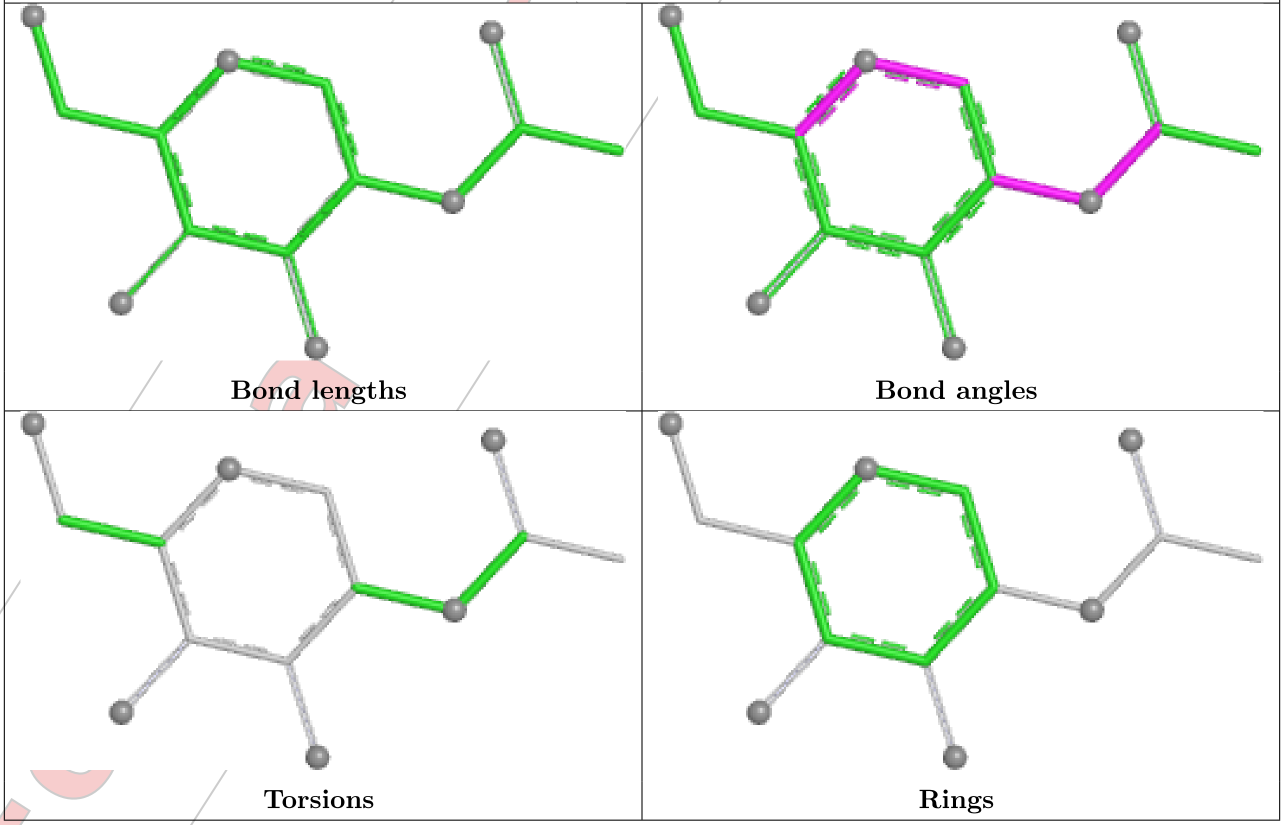

**Ligand NAG D 302**

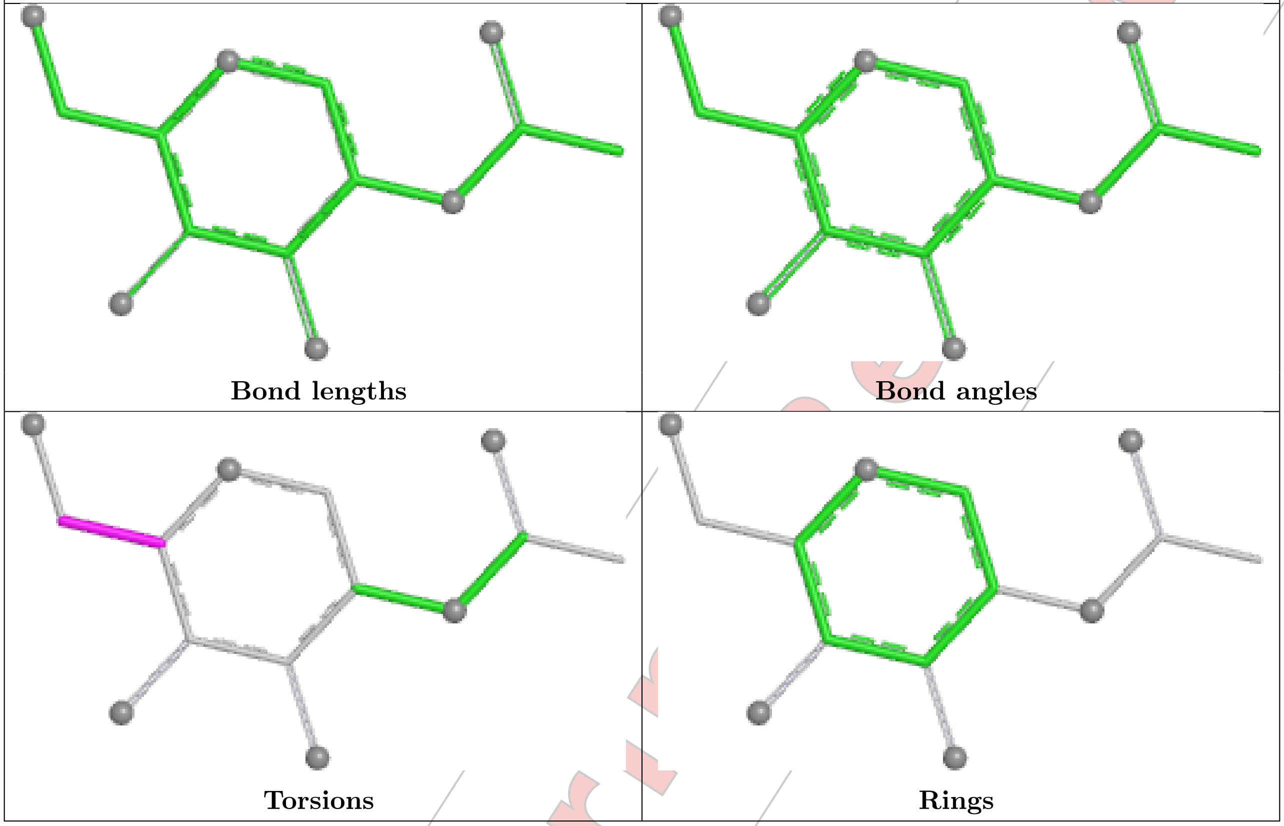


#### 5.7 Other polymers (i)

There are no such residues in this entry.

#### 5.8 Polymer linkage issues (i)

There are no chain breaks in this entry.

For Manuscript Review

### 6 Fit of model and data (i)

#### 6.1 Protein, DNA and RNA chains (i)

In the following table, the column labelled '#RSRZ> 2' contains the number (and percentage) of RSRZ outliers, followed by percent RSRZ outliers for the chain as percentile scores relative to all X-ray entries and entries of similar resolution. The OWAB column contains the minimum, median, $95^{th}$ percentile and maximum values of the occupancy-weighted average B-factor per residue. The column labelled 'Q< 0.9' lists the number of (and percentage) of residues with an average occupancy less than 0.9.

| Mol | Chain | Analysed | <RSRZ> | #RSRZ>2 | | | OWAB(Å$^2$) | Q<0.9 |
|---|---|---|---|---|---|---|---|---|
| 1 | A | 204/219 (93%) | 0.37 | 7 (3%) | 48 | 53 | 33, 53, 108, 145 | 0 |
| 1 | C | 195/219 (89%) | 0.83 | 16 (8%) | 19 | 22 | 43, 79, 128, 146 | 0 |
| 2 | B | 237/250 (94%) | 0.05 | 0 | 100 | 100 | 34, 48, 78, 122 | 0 |
| 2 | D | 236/250 (94%) | 0.70 | 13 (5%) | 32 | 36 | 46, 73, 118, 149 | 0 |
| All | All | 872/938 (92%) | 0.48 | 36 (4%) | 42 | 47 | 33, 63, 115, 149 | 0 |

All (36) RSRZ outliers are listed below:

| Mol | Chain | Res | Type | RSRZ |
|---|---|---|---|---|
| 1 | C | 104 | TYR | 5.1 |
| 2 | D | 97 | GLY | 4.3 |
| 1 | A | 99 | GLY | 4.0 |
| 1 | C | 161 | PHE | 4.0 |
| 1 | C | 34 | LEU | 3.4 |
| 2 | D | 63 | PHE | 3.4 |
| 1 | A | 137 | ASP | 3.2 |
| 1 | C | 194 | ILE | 3.1 |
| 1 | C | 96 | GLU | 3.0 |
| 2 | D | 26 | LEU | 3.0 |
| 1 | C | 59 | ALA | 3.0 |
| 1 | C | 32 | TYR | 2.9 |
| 2 | D | 75 | LEU | 2.7 |
| 2 | D | 17 | ASN | 2.6 |
| 1 | A | 194 | ILE | 2.6 |
| 2 | D | 239 | ASP | 2.5 |
| 2 | D | 241 | SER | 2.5 |
| 1 | C | 84 | ILE | 2.5 |
| 1 | C | 193 | ASP | 2.5 |
| 1 | A | 103 | ASN | 2.4 |
| 2 | D | 76 | HIS | 2.4 |

Continued on next page...

For Manuscript Review

*Continued from previous page...*

| Mol | Chain | Res | Type | RSRZ |
|---|---|---|---|---|
| 2 | D | 33 | TRP | 2.4 |
| 2 | D | 69 | ASP | 2.4 |
| 1 | C | 56 | LYS | 2.3 |
| 1 | C | 156 | MET | 2.3 |
| 2 | D | 16 | ALA | 2.3 |
| 1 | A | 43 | GLY | 2.2 |
| 2 | D | 240 | CYS | 2.1 |
| 1 | C | 189 | PHE | 2.1 |
| 1 | C | 195 | PHE | 2.1 |
| 1 | C | 197 | GLU | 2.1 |
| 1 | C | 173 | MET | 2.1 |
| 2 | D | 219 | PRO | 2.0 |
| 1 | A | 33 | PHE | 2.0 |
| 1 | A | 161 | PHE | 2.0 |
| 1 | C | 33 | PHE | 2.0 |

#### 6.2 Non-standard residues in protein, DNA, RNA chains (i)

There are no non-standard protein/DNA/RNA residues in this entry.

#### 6.3 Carbohydrates (i)

There are no monosaccharides in this entry.

#### 6.4 Ligands (i)

In the following table, the Atoms column lists the number of modelled atoms in the group and the number defined in the chemical component dictionary. The B-factors column lists the minimum, median, $95^{th}$ percentile and maximum values of B factors of atoms in the group. The column labelled 'Q< 0.9' lists the number of atoms with occupancy less than 0.9.

| Mol | Type | Chain | Res | Atoms | RSCC | RSR | B-factors($Å^2$) | Q<0.9 |
|---|---|---|---|---|---|---|---|---|
| 3 | NAG | D | 301 | 14/15 | 0.50 | 0.16 | 76,78,86,88 | 0 |
| 3 | NAG | B | 303 | 14/15 | 0.74 | 0.11 | 76,79,82,84 | 0 |
| 3 | NAG | D | 302 | 14/15 | 0.74 | 0.12 | 80,85,89,90 | 0 |
| 4 | GOL | D | 303 | 6/6 | 0.74 | 0.14 | 61,67,68,71 | 0 |
| 3 | NAG | B | 302 | 14/15 | 0.81 | 0.10 | 48,53,58,58 | 0 |
| 3 | NAG | A | 301 | 14/15 | 0.88 | 0.09 | 49,57,64,67 | 0 |
| 3 | NAG | B | 301 | 14/15 | 0.90 | 0.10 | 38,43,48,51 | 0 |

For Manuscript Review

The following is a graphical depiction of the model fit to experimental electron density of all instances of the Ligand of Interest. In addition, ligands with molecular weight > 250 and outliers as shown on the geometry validation Tables will also be included. Each fit is shown from different orientation to approximate a three-dimensional view.

**Electron density around NAG D 301:**

2mF$_o$-DF$_c$ (at 0.7 rmsd) in gray
mF$_o$-DF$_c$ (at 3 rmsd) in purple (negative)
and green (positive)

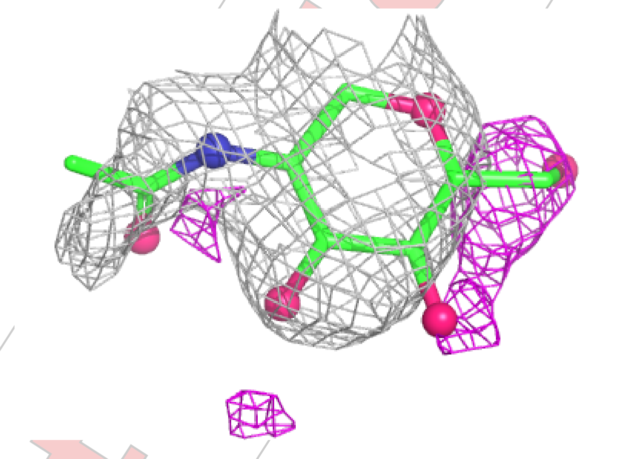

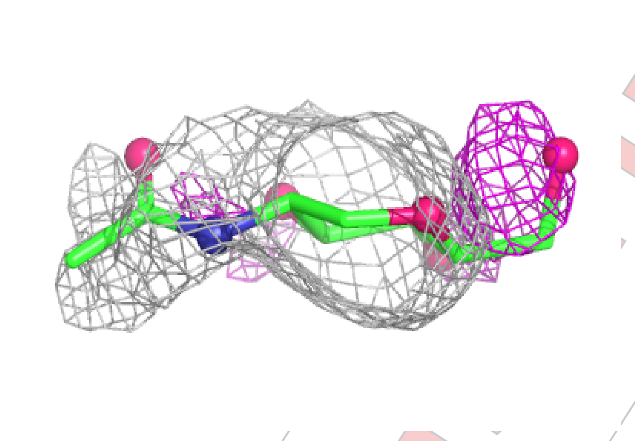

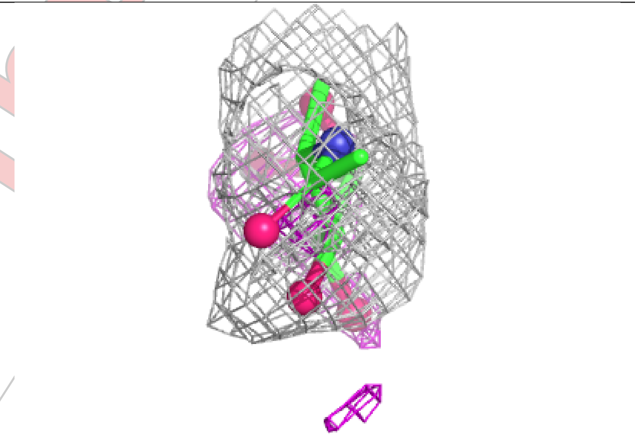

For Manuscript Review

**Electron density around NAG B 303:**

$2mF_o$-$DF_c$ (at 0.7 rmsd) in gray
$mF_o$-$DF_c$ (at 3 rmsd) in purple (negative)
and green (positive)

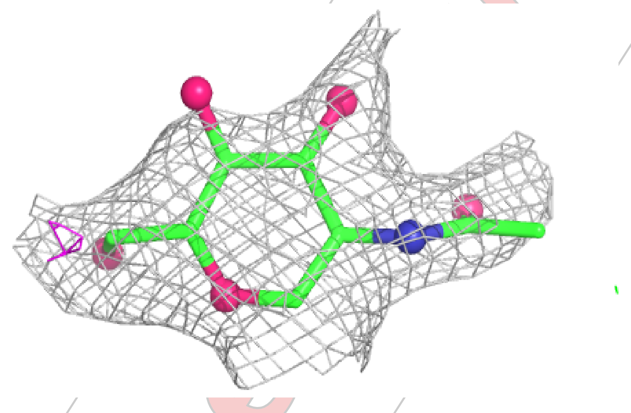

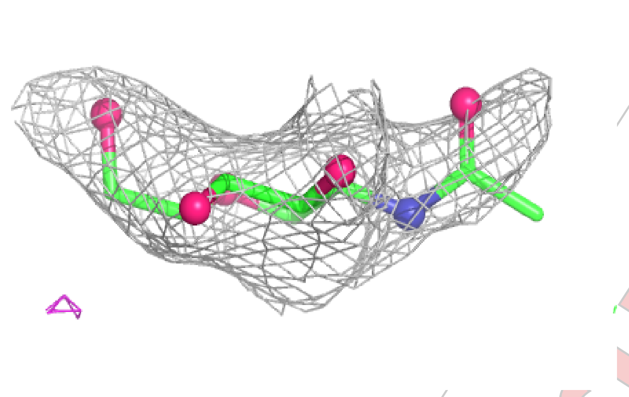

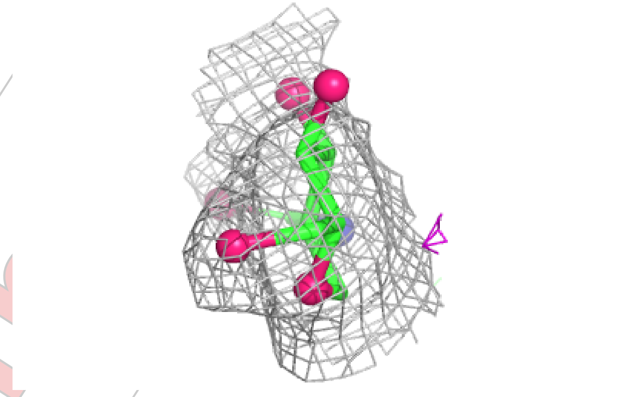

**Electron density around NAG D 302:**

$2mF_o$-$DF_c$ (at 0.7 rmsd) in gray
$mF_o$-$DF_c$ (at 3 rmsd) in purple (negative)
and green (positive)

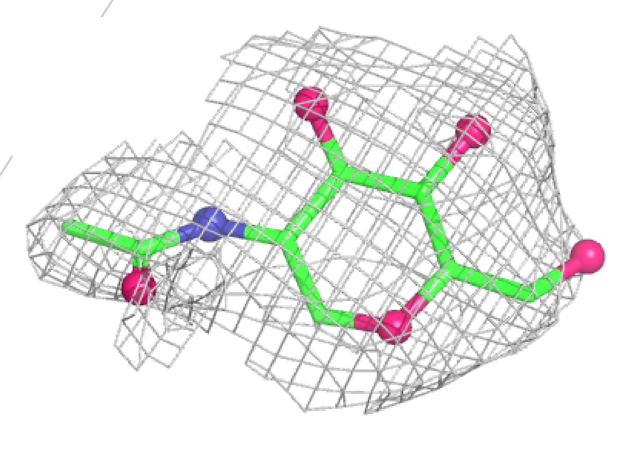

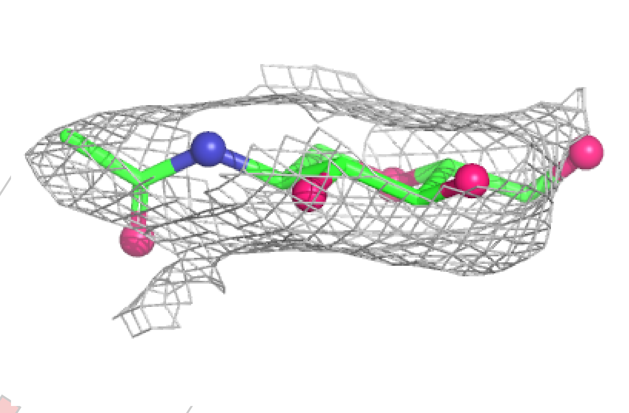

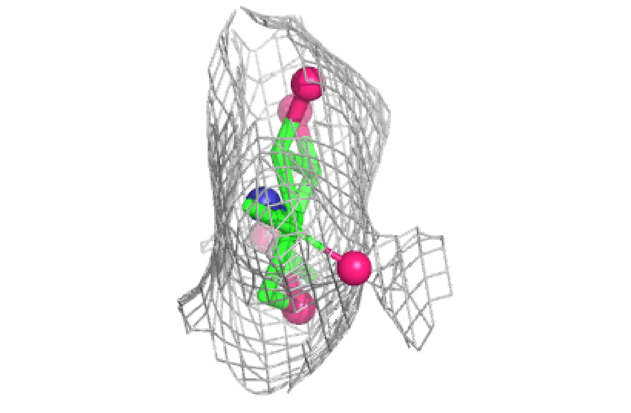

**Electron density around NAG B 302:**

$2mF_o$-$DF_c$ (at 0.7 rmsd) in gray
$mF_o$-$DF_c$ (at 3 rmsd) in purple (negative)
and green (positive)

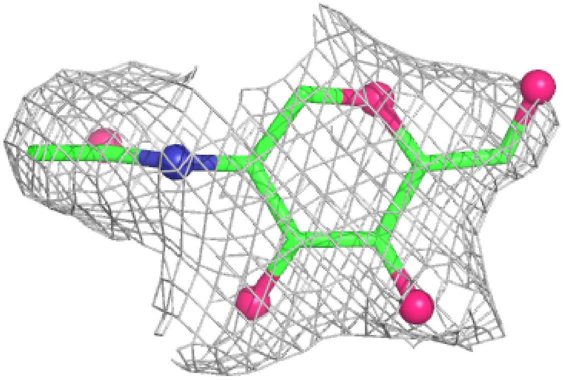

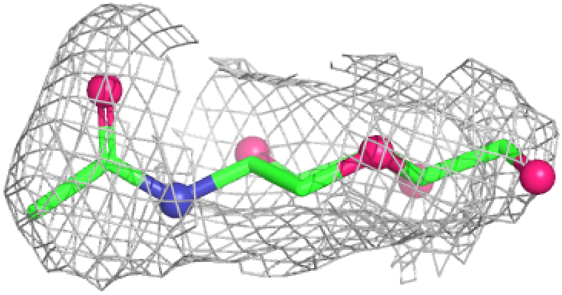

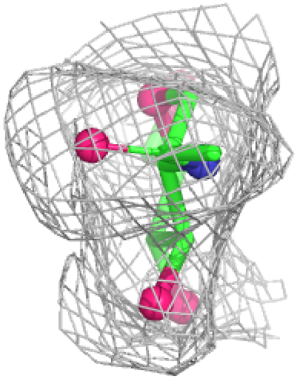

**Electron density around NAG A 301:**

$2mF_o$-$DF_c$ (at 0.7 rmsd) in gray
$mF_o$-$DF_c$ (at 3 rmsd) in purple (negative)
and green (positive)

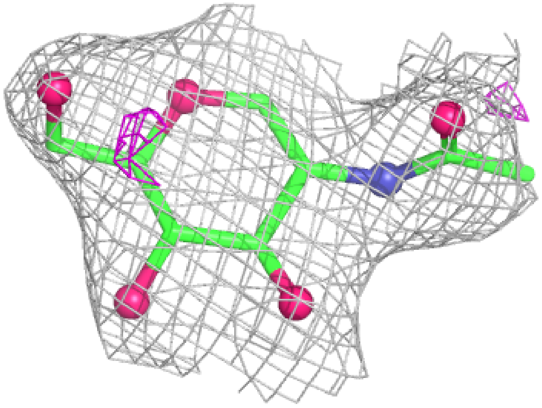

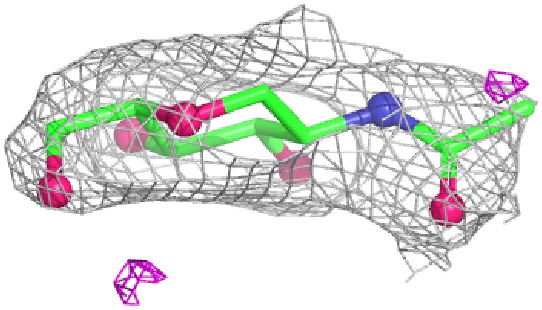

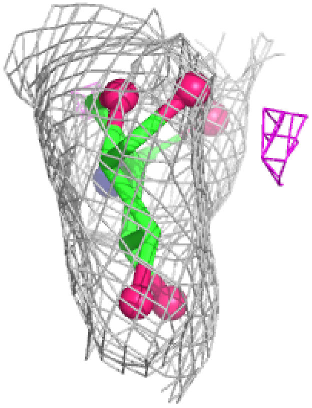

**Electron density around NAG B 301:**

$2mF_o$-$DF_c$ (at 0.7 rmsd) in gray
$mF_o$-$DF_c$ (at 3 rmsd) in purple (negative) and green (positive)

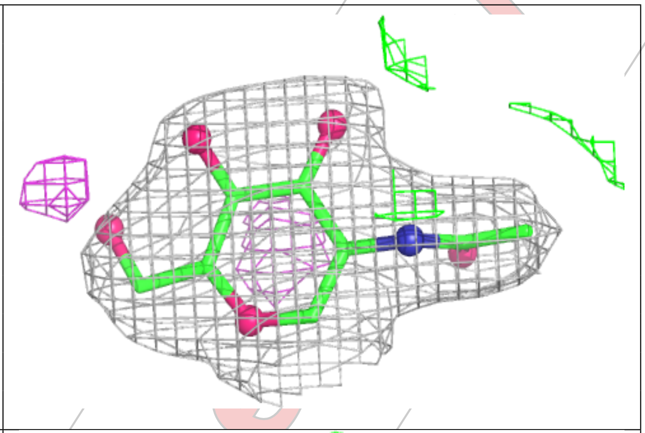

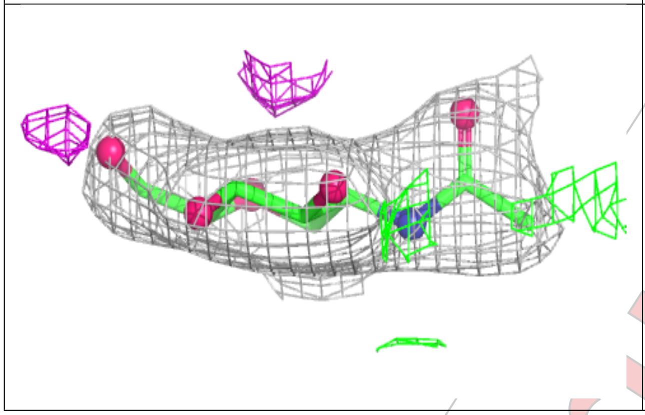

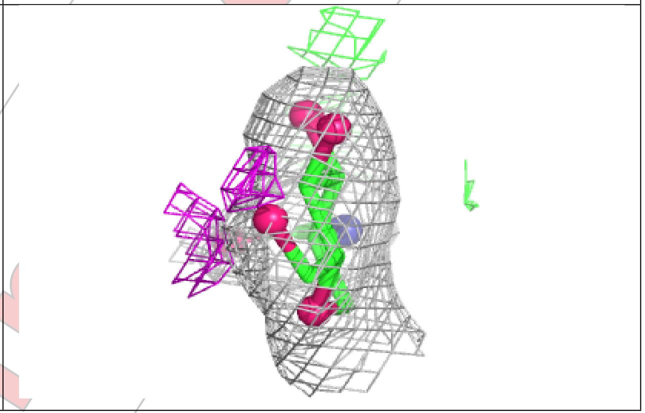

#### 6.5 Other polymers ⓘ

There are no such residues in this entry.

For Manuscript Review

### Full wwPDB X-ray Structure Validation Report (i)

Jun 30, 2025 – 10:50 AM EDT

| | | |
|---|---|---|
| PDB ID | : | 9NU7 / pdb_00009nu7 |
| Title | : | Crystal structure of NP202 TCR in complex with NP366-H-2Db |
| Deposited on | : | 2025-03-19 |
| Resolution | : | 3.60 Å(reported) |

**This wwPDB validation report is for manuscript review**

This is a Full wwPDB X-ray Structure Validation Report.

This report is produced by the wwPDB biocuration pipeline after annotation of the structure.

We welcome your comments at **
A user guide is available at
https://www.wwpdb.org/validation/2017/XrayValidationReportHelp
with specific help available everywhere you see the (i) symbol.

The types of validation reports are described at
http://www.wwpdb.org/validation/2017/FAQs#types.

The following versions of software and data (see references (i)) were used in the production of this report:

| | | |
|---|---|---|
| MolProbity | : | 4-5-2 with Phenix2.0rc1 |
| Mogul | : | 2022.3.0, CSD as543be (2022) |
| Xtriage (Phenix) | : | 2.0rc1 |
| EDS | : | 3.0 |
| Percentile statistics | : | 20231227.v01 (using entries in the PDB archive December 27th 2023) |
| CCP4 | : | 9.0.006 (Gargrove) |
| Density-Fitness | : | 1.0.12 |
| Ideal geometry (proteins) | : | Engh & Huber (2001) |
| Ideal geometry (DNA, RNA) | : | Parkinson et al. (1996) |
| Validation Pipeline (wwPDB-VP) | : | 2.44 |

For Manuscript Review

### 1 Overall quality at a glance (i)

The following experimental techniques were used to determine the structure:
*X-RAY DIFFRACTION*

The reported resolution of this entry is 3.60 Å.

Percentile scores (ranging between 0-100) for global validation metrics of the entry are shown in the following graphic. The table shows the number of entries on which the scores are based.

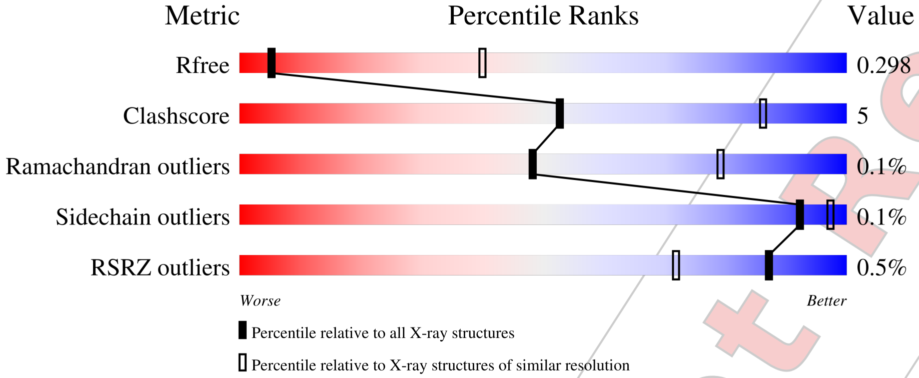


| Metric | Whole archive (#Entries) | Similar resolution (#Entries, resolution range(Å)) |
|---|---|---|
| $R_{free}$ | 164625 | 1563 (3.70-3.50) |
| Clashscore | 180529 | 1665 (3.70-3.50) |
| Ramachandran outliers | 177936 | 1641 (3.70-3.50) |
| Sidechain outliers | 177891 | 1640 (3.70-3.50) |
| RSRZ outliers | 164620 | 1562 (3.70-3.50) |

The table below summarises the geometric issues observed across the polymeric chains and their fit to the electron density. The red, orange, yellow and green segments of the lower bar indicate the fraction of residues that contain outliers for >=3, 2, 1 and 0 types of geometric quality criteria respectively. A grey segment represents the fraction of residues that are not modelled. The numeric value for each fraction is indicated below the corresponding segment, with a dot representing fractions <=5% The upper red bar (where present) indicates the fraction of residues that have poor fit to the electron density. The numeric value is given above the bar.

| Mol | Chain | Length | Quality of chain |
|---|---|---|---|
| 1 | A | 219 | 81% 13% 5% |
| 1 | S | 219 | 79% 14% 7% |
| 2 | B | 250 | % / 89% 6% 5% |
| 2 | T | 250 | % / 84% 11% 5% |

*Continued on next page...*

For Manuscript Review

*Continued from previous page...*

| Mol | Chain | Length | Quality of chain |
|---|---|---|---|
| 3 | C | 281 | 89% 10% • |
| 4 | D | 99 | 77% 23% |
| 5 | P | 9 | 100% |

For Manuscript Review

#### 2 Entry composition (i)

There are 6 unique types of molecules in this entry. The entry contains 9902 atoms, of which 0 are hydrogens and 0 are deuteriums.

In the tables below, the ZeroOcc column contains the number of atoms modelled with zero occupancy, the AltConf column contains the number of residues with at least one atom in alternate conformation and the Trace column contains the number of residues modelled with at most 2 atoms.

- Molecule 1 is a protein called NP202 TCR alpha-chain.

| Mol | Chain | Residues | Atoms | | | | | ZeroOcc | AltConf | Trace |
|---|---|---|---|---|---|---|---|---|---|---|
| | | | Total | C | N | O | S | | | |
| 1 | A | 207 | 1540 | 973 | 243 | 315 | 9 | 0 | 0 | 0 |
| 1 | S | 204 | 1542 | 973 | 246 | 313 | 10 | 0 | 0 | 0 |

- Molecule 2 is a protein called NP202 TCR beta-chain.

| Mol | Chain | Residues | Atoms | | | | | ZeroOcc | AltConf | Trace |
|---|---|---|---|---|---|---|---|---|---|---|
| | | | Total | C | N | O | S | | | |
| 2 | B | 238 | 1913 | 1215 | 335 | 356 | 7 | 0 | 0 | 0 |
| 2 | T | 238 | 1894 | 1206 | 329 | 352 | 7 | 0 | 0 | 0 |

- Molecule 3 is a protein called H-2 class I histocompatibility antigen, D-B alpha chain.

| Mol | Chain | Residues | Atoms | | | | | ZeroOcc | AltConf | Trace |
|---|---|---|---|---|---|---|---|---|---|---|
| | | | Total | C | N | O | S | | | |
| 3 | C | 278 | 2092 | 1316 | 370 | 396 | 10 | 0 | 0 | 0 |

There is a discrepancy between the modelled and reference sequences:

| Chain | Residue | Modelled | Actual | Comment | Reference |
|---|---|---|---|---|---|
| C | 0 | MET | - | initiating methionine | UNP P01899 |

- Molecule 4 is a protein called Beta-2-microglobulin.

| Mol | Chain | Residues | Atoms | | | | | ZeroOcc | AltConf | Trace |
|---|---|---|---|---|---|---|---|---|---|---|
| | | | Total | C | N | O | S | | | |
| 4 | D | 99 | 796 | 509 | 130 | 150 | 7 | 0 | 0 | 0 |

There is a discrepancy between the modelled and reference sequences:

For Manuscript Review

| Chain | Residue | Modelled | Actual | Comment | Reference |
|---|---|---|---|---|---|
| D | 85 | ASP | ALA | engineered mutation | UNP P01887 |

- Molecule 5 is a protein called Nucleoprotein peptide.

| Mol | Chain | Residues | Atoms | ZeroOcc | AltConf | Trace |
|---|---|---|---|---|---|---|
| 5 | P | 9 | Total C N O S<br>69 38 11 18 2 | 0 | 0 | 0 |

- Molecule 6 is 2-acetamido-2-deoxy-beta-D-glucopyranose (CCD ID: NAG) (formula: $C_8H_{15}NO_6$).

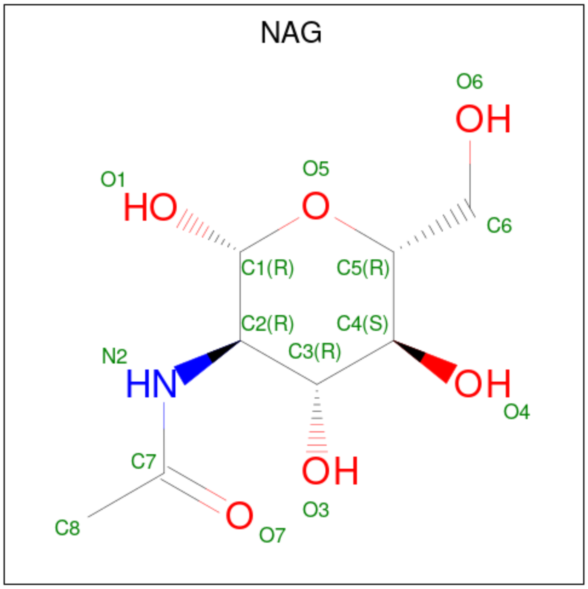


| Mol | Chain | Residues | Atoms | ZeroOcc | AltConf |
|---|---|---|---|---|---|
| 6 | A | 1 | Total C N O<br>14 8 1 5 | 0 | 0 |
| 6 | B | 1 | Total C N O<br>14 8 1 5 | 0 | 0 |
| 6 | B | 1 | Total C N O<br>14 8 1 5 | 0 | 0 |
| 6 | T | 1 | Total C N O<br>14 8 1 5 | 0 | 0 |

For Manuscript Review

### 3 Residue-property plots

These plots are drawn for all protein, RNA, DNA and oligosaccharide chains in the entry. The first graphic for a chain summarises the proportions of the various outlier classes displayed in the second graphic. The second graphic shows the sequence view annotated by issues in geometry and electron density. Residues are color-coded according to the number of geometric quality criteria for which they contain at least one outlier: green = 0, yellow = 1, orange = 2 and red = 3 or more. A red dot above a residue indicates a poor fit to the electron density (RSRZ > 2). Stretches of 2 or more consecutive residues without any outlier are shown as a green connector. Residues present in the sample, but not in the model, are shown in grey.

- Molecule 1: NP202 TCR alpha-chain

Chain A:

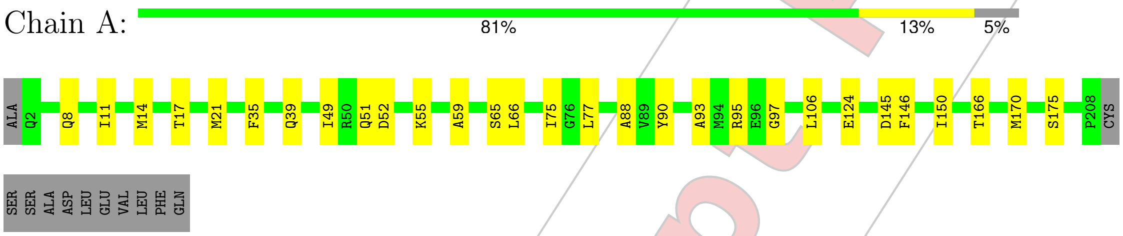


- Molecule 1: NP202 TCR alpha-chain

Chain S:

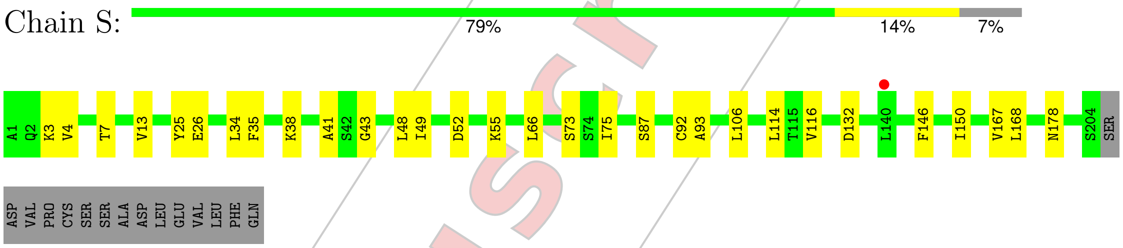


- Molecule 2: NP202 TCR beta-chain

Chain B:

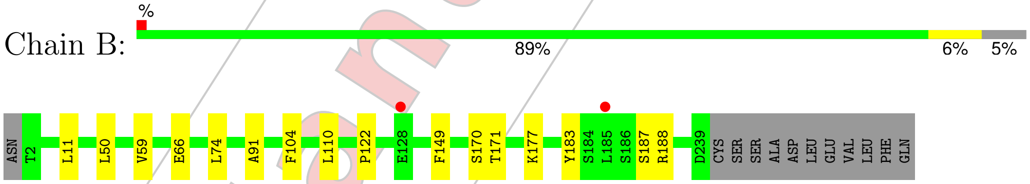


- Molecule 2: NP202 TCR beta-chain

Chain T:

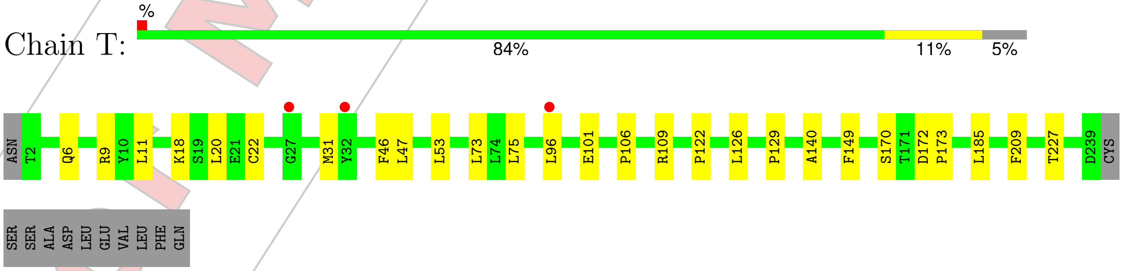


- Molecule 3: H-2 class I histocompatibility antigen, D-B alpha chain

For Manuscript Review

Chain C: 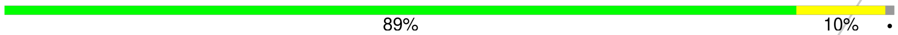


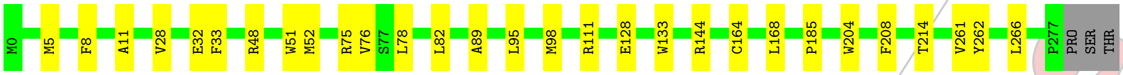


• Molecule 4: Beta-2-microglobulin

Chain D: 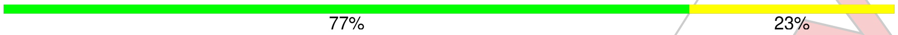


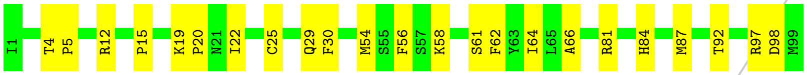


• Molecule 5: Nucleoprotein peptide

Chain P: 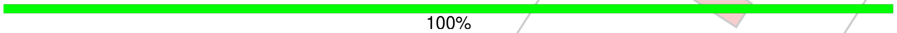


There are no outlier residues recorded for this chain.

For Manuscript Review

### 4 Data and refinement statistics (i)

| Property | Value | Source |
|---|---|---|
| Space group | P 1 21 1 | Depositor |
| Cell constants<br>a, b, c, $\alpha$, $\beta$, $\gamma$ | 94.29Å 89.59Å 104.27Å<br>90.00° 96.17° 90.00° | Depositor |
| Resolution (Å) | 51.83 – 3.60<br>51.83 – 3.63 | Depositor<br>EDS |
| % Data completeness<br>(in resolution range) | 92.6 (51.83-3.60)<br>92.6 (51.83-3.63) | Depositor<br>EDS |
| $R_{merge}$ | 0.28 | Depositor |
| $R_{sym}$ | (Not available) | Depositor |
| $< I/\sigma(I) >$ [1] | 1.65 (at 3.67Å) | Xtriage |
| Refinement program | PHENIX (1.21.2_5419: ???) | Depositor |
| R, $R_{free}$ | 0.242 , 0.300<br>0.244 , 0.298 | Depositor<br>DCC |
| $R_{free}$ test set | 1013 reflections (5.06%) | wwPDB-VP |
| Wilson B-factor (Å$^2$) | 111.2 | Xtriage |
| Anisotropy | 0.163 | Xtriage |
| Bulk solvent $k_{sol}$(e/Å$^3$), $B_{sol}$(Å$^2$) | 0.31 , 93.7 | EDS |
| L-test for twinning[2] | $< \|L\| > = 0.44$, $< L^2 > = 0.26$ | Xtriage |
| Estimated twinning fraction | No twinning to report. | Xtriage |
| $F_o$,$F_c$ correlation | 0.88 | EDS |
| Total number of atoms | 9902 | wwPDB-VP |
| Average B, all atoms (Å$^2$) | 129.0 | wwPDB-VP |

Xtriage's analysis on translational NCS is as follows: *The largest off-origin peak in the Patterson function is 4.67% of the height of the origin peak. No significant pseudotranslation is detected.*

[1]Intensities estimated from amplitudes.

[2]Theoretical values of $< |L| >$, $< L^2 >$ for acentric reflections are 0.5, 0.333 respectively for untwinned datasets, and 0.375, 0.2 for perfectly twinned datasets.

For Manuscript Review

### 5 Model quality (i)

#### 5.1 Standard geometry (i)

Bond lengths and bond angles in the following residue types are not validated in this section: NAG

The Z score for a bond length (or angle) is the number of standard deviations the observed value is removed from the expected value. A bond length (or angle) with $|Z| > 5$ is considered an outlier worth inspection. RMSZ is the root-mean-square of all Z scores of the bond lengths (or angles).

| Mol | Chain | Bond lengths | | Bond angles | |
|---|---|---|---|---|---|
| | | RMSZ | #\|Z\| >5 | RMSZ | #\|Z\| >5 |
| 1 | A | 0.17 | 0/1572 | 0.33 | 0/2147 |
| 1 | S | 0.15 | 0/1572 | 0.30 | 0/2137 |
| 2 | B | 0.14 | 0/1969 | 0.28 | 0/2675 |
| 2 | T | 0.14 | 0/1950 | 0.28 | 0/2652 |
| 3 | C | 0.17 | 0/2150 | 0.29 | 0/2934 |
| 4 | D | 0.15 | 0/822 | 0.32 | 0/1121 |
| 5 | P | 0.23 | 0/68 | 0.43 | 0/88 |
| All | All | 0.16 | 0/10103 | 0.30 | 0/13754 |

There are no bond length outliers.

There are no bond angle outliers.

There are no chirality outliers.

There are no planarity outliers.

#### 5.2 Too-close contacts (i)

In the following table, the Non-H and H(model) columns list the number of non-hydrogen atoms and hydrogen atoms in the chain respectively. The H(added) column lists the number of hydrogen atoms added and optimized by MolProbity. The Clashes column lists the number of clashes within the asymmetric unit, whereas Symm-Clashes lists symmetry-related clashes.

| Mol | Chain | Non-H | H(model) | H(added) | Clashes | Symm-Clashes |
|---|---|---|---|---|---|---|
| 1 | A | 1540 | 0 | 1406 | 20 | 0 |
| 1 | S | 1542 | 0 | 1454 | 20 | 0 |
| 2 | B | 1913 | 0 | 1844 | 11 | 0 |
| 2 | T | 1894 | 0 | 1817 | 18 | 0 |
| 3 | C | 2092 | 0 | 1820 | 20 | 0 |
| 4 | D | 796 | 0 | 733 | 19 | 0 |

*Continued on next page...*

For Manuscript Review

*Continued from previous page...*

| Mol | Chain | Non-H | H(model) | H(added) | Clashes | Symm-Clashes |
|---|---|---|---|---|---|---|
| 5 | P | 69 | 0 | 61 | 0 | 0 |
| 6 | A | 14 | 0 | 13 | 0 | 0 |
| 6 | B | 28 | 0 | 26 | 1 | 0 |
| 6 | T | 14 | 0 | 13 | 1 | 0 |
| All | All | 9902 | 0 | 9187 | 94 | 0 |

The all-atom clashscore is defined as the number of clashes found per 1000 atoms (including hydrogen atoms). The all-atom clashscore for this structure is 5.

All (94) close contacts within the same asymmetric unit are listed below, sorted by their clash magnitude.

| Atom-1 | Atom-2 | Interatomic distance (Å) | Clash overlap (Å) |
|---|---|---|---|
| 1:S:3:LYS:HB3 | 1:S:26:GLU:HB3 | 1.65 | 0.76 |
| 4:D:12:ARG:HH21 | 4:D:22:ILE:HG21 | 1.54 | 0.73 |
| 3:C:5:MET:HB2 | 3:C:168:LEU:HD13 | 1.72 | 0.71 |
| 3:C:133:TRP:HB2 | 3:C:144:ARG:HG3 | 1.74 | 0.70 |
| 2:T:6:GLN:HG3 | 2:T:106:PRO:HD2 | 1.73 | 0.69 |
| 4:D:5:PRO:HB3 | 4:D:30:PHE:HB3 | 1.75 | 0.68 |
| 2:B:50:LEU:HD21 | 3:C:76:VAL:HG22 | 1.75 | 0.68 |
| 3:C:98:MET:HE1 | 4:D:58:LYS:HG2 | 1.77 | 0.67 |
| 2:T:31:MET:HB3 | 2:T:73:LEU:HD11 | 1.76 | 0.66 |
| 2:T:53:LEU:HD23 | 6:T:301:NAG:H82 | 1.82 | 0.61 |
| 2:B:171:THR:HG23 | 2:B:187:SER:HB2 | 1.83 | 0.60 |
| 2:B:66:GLU:HB2 | 2:B:74:LEU:HB2 | 1.85 | 0.59 |
| 1:A:8:GLN:HG3 | 1:S:13:VAL:HG12 | 1.84 | 0.59 |
| 3:C:82:LEU:HD11 | 3:C:89:ALA:HA | 1.86 | 0.58 |
| 1:S:87:SER:HB3 | 1:S:116:VAL:H | 1.70 | 0.57 |
| 1:S:132:ASP:OD1 | 2:T:126:LEU:O | 2.23 | 0.57 |
| 4:D:30:PHE:HB2 | 4:D:84:HIS:CE1 | 2.40 | 0.57 |
| 4:D:19:LYS:HD2 | 4:D:20:PRO:HD2 | 1.87 | 0.56 |
| 4:D:54:MET:HE2 | 4:D:64:ILE:HD13 | 1.87 | 0.56 |
| 1:S:34:LEU:HD13 | 1:S:75:ILE:HG21 | 1.87 | 0.56 |
| 1:A:8:GLN:HG2 | 1:A:11:ILE:HD11 | 1.87 | 0.55 |
| 1:A:66:LEU:HD22 | 1:A:75:ILE:HD11 | 1.89 | 0.55 |
| 3:C:262:TYR:HA | 3:C:266:LEU:HD12 | 1.89 | 0.55 |
| 2:T:11:LEU:HD11 | 2:T:18:LYS:HD3 | 1.89 | 0.55 |
| 3:C:204:TRP:HH2 | 4:D:98:ASP:HB2 | 1.70 | 0.55 |
| 1:A:49:ILE:HD13 | 1:A:66:LEU:HG | 1.89 | 0.54 |
| 3:C:28:VAL:HG23 | 3:C:33:PHE:CE1 | 2.42 | 0.54 |
| 1:S:52:ASP:HB2 | 1:S:55:LYS:HG3 | 1.90 | 0.54 |
| 2:B:50:LEU:HB2 | 3:C:75:ARG:HH12 | 1.73 | 0.53 |

*Continued on next page...*

For Manuscript Review

Continued from previous page...

| Atom-1 | Atom-2 | Interatomic distance (Å) | Clash overlap (Å) |
|---|---|---|---|
| 4:D:29:GLN:HA | 4:D:61:SER:HB2 | 1.91 | 0.53 |
| 1:S:13:VAL:HG22 | 1:S:114:LEU:HD11 | 1.90 | 0.53 |
| 3:C:33:PHE:HB2 | 3:C:52:MET:HG3 | 1.90 | 0.52 |
| 1:A:14:MET:HE3 | 1:S:7:THR:HA | 1.90 | 0.52 |
| 1:S:38:LYS:HB2 | 1:S:48:LEU:HD11 | 1.92 | 0.52 |
| 3:C:51:TRP:CZ3 | 3:C:52:MET:HE3 | 2.46 | 0.51 |
| 1:A:170:MET:HB2 | 1:A:175:SER:HB3 | 1.94 | 0.50 |
| 1:S:93:ALA:HB1 | 1:S:106:LEU:HD11 | 1.94 | 0.50 |
| 4:D:54:MET:HE2 | 4:D:64:ILE:HG21 | 1.94 | 0.50 |
| 2:T:9:ARG:HH11 | 2:T:106:PRO:HB2 | 1.76 | 0.50 |
| 1:A:146:PHE:HB2 | 1:A:150:ILE:HD11 | 1.94 | 0.49 |
| 1:A:39:GLN:O | 1:A:88:ALA:HB1 | 2.13 | 0.49 |
| 3:C:52:MET:HA | 3:C:52:MET:HE2 | 1.94 | 0.49 |
| 1:A:95:ARG:HB2 | 1:A:106:LEU:HD13 | 1.94 | 0.49 |
| 1:A:124:GLU:HG3 | 1:A:145:ASP:HB3 | 1.95 | 0.49 |
| 4:D:4:THR:HG23 | 4:D:87:MET:SD | 2.52 | 0.49 |
| 1:A:52:ASP:HB2 | 1:A:55:LYS:HB2 | 1.96 | 0.48 |
| 2:T:47:LEU:HD21 | 2:T:96:LEU:HD11 | 1.94 | 0.48 |
| 1:A:35:PHE:HE2 | 1:A:95:ARG:HB3 | 1.79 | 0.47 |
| 3:C:185:PRO:HB3 | 3:C:208:PHE:HB3 | 1.95 | 0.47 |
| 1:A:166:THR:HG22 | 2:B:170:SER:HB2 | 1.97 | 0.47 |
| 4:D:15:PRO:HG3 | 4:D:98:ASP:OD1 | 2.15 | 0.46 |
| 3:C:11:ALA:HB3 | 3:C:95:LEU:HD11 | 1.98 | 0.46 |
| 3:C:78:LEU:HG | 3:C:95:LEU:HD21 | 1.97 | 0.46 |
| 1:A:21:MET:HE1 | 1:A:90:TYR:CG | 2.51 | 0.46 |
| 3:C:204:TRP:CH2 | 4:D:98:ASP:HB2 | 2.50 | 0.46 |
| 2:B:59:VAL:O | 6:B:301:NAG:H5 | 2.16 | 0.46 |
| 3:C:32:GLU:OE2 | 3:C:48:ARG:HD3 | 2.17 | 0.45 |
| 2:T:46:PHE:HB3 | 2:T:53:LEU:HD11 | 1.99 | 0.45 |
| 2:T:172:ASP:HB2 | 2:T:185:LEU:HD12 | 1.99 | 0.45 |
| 1:A:35:PHE:HB2 | 1:A:93:ALA:HB3 | 1.99 | 0.44 |
| 1:S:35:PHE:HB2 | 1:S:93:ALA:HB3 | 1.99 | 0.44 |
| 2:T:22:CYS:HB3 | 2:T:73:LEU:HB2 | 1.98 | 0.44 |
| 3:C:214:THR:O | 3:C:261:VAL:HA | 2.18 | 0.44 |
| 4:D:12:ARG:NH2 | 4:D:22:ILE:HD13 | 2.33 | 0.44 |
| 3:C:111:ARG:HD2 | 3:C:128:GLU:OE1 | 2.18 | 0.44 |
| 4:D:56:PHE:HB3 | 4:D:62:PHE:CD2 | 2.52 | 0.44 |
| 2:B:170:SER:OG | 2:B:188:ARG:HD3 | 2.18 | 0.43 |
| 2:B:177:LYS:HB3 | 2:B:183:TYR:CE2 | 2.52 | 0.43 |
| 4:D:12:ARG:HH21 | 4:D:22:ILE:HD13 | 1.83 | 0.43 |
| 4:D:25:CYS:HB3 | 4:D:66:ALA:HB3 | 2.00 | 0.43 |

Continued on next page...

For Manuscript Review

*Continued from previous page...*

| Atom-1 | Atom-2 | Interatomic distance (Å) | Clash overlap (Å) |
|---|---|---|---|
| 1:A:51:GLN:HB2 | 1:A:66:LEU:HD13 | 2.00 | 0.43 |
| 1:A:59:ALA:HB3 | 1:A:66:LEU:HD12 | 2.01 | 0.43 |
| 2:T:209:PHE:O | 2:T:227:THR:HA | 2.18 | 0.43 |
| 1:A:17:THR:HG21 | 1:S:7:THR:HB | 1.99 | 0.43 |
| 2:B:11:LEU:HD23 | 2:B:110:LEU:HD12 | 1.99 | 0.43 |
| 1:S:41:ALA:HB1 | 2:T:173:PRO:HB3 | 2.01 | 0.43 |
| 1:A:65:SER:O | 1:A:77:LEU:HD12 | 2.19 | 0.42 |
| 2:T:101:GLU:H | 2:T:101:GLU:CD | 2.27 | 0.42 |
| 1:S:146:PHE:HB2 | 1:S:150:ILE:HD11 | 2.02 | 0.42 |
| 2:B:122:PRO:HB3 | 2:B:149:PHE:HB3 | 2.02 | 0.42 |
| 1:S:49:ILE:HD13 | 1:S:66:LEU:HG | 2.02 | 0.42 |
| 2:B:91:ALA:HB2 | 2:B:104:PHE:CD1 | 2.55 | 0.42 |
| 2:T:20:LEU:HB2 | 2:T:75:LEU:HB3 | 2.01 | 0.42 |
| 4:D:81:ARG:HG2 | 4:D:92:THR:OG1 | 2.20 | 0.41 |
| 1:S:168:LEU:HB3 | 2:T:170:SER:HB3 | 2.02 | 0.41 |
| 3:C:8:PHE:HB3 | 4:D:56:PHE:CE1 | 2.56 | 0.41 |
| 1:S:4:VAL:HG11 | 1:S:92:CYS:O | 2.21 | 0.41 |
| 4:D:97:ARG:H | 4:D:97:ARG:HD2 | 1.86 | 0.41 |
| 1:S:25:TYR:CZ | 1:S:73:SER:HA | 2.56 | 0.41 |
| 2:T:122:PRO:HB3 | 2:T:149:PHE:HB3 | 2.02 | 0.41 |
| 1:S:43:GLY:HA3 | 2:T:109:ARG:HH21 | 1.86 | 0.40 |
| 1:S:167:VAL:HG22 | 1:S:178:ASN:OD1 | 2.22 | 0.40 |
| 2:T:129:PRO:HG3 | 2:T:140:ALA:HB1 | 2.03 | 0.40 |
| 1:A:55:LYS:HB3 | 1:A:55:LYS:HE2 | 1.86 | 0.40 |

There are no symmetry-related clashes.

#### 5.3 Torsion angles (i)

##### 5.3.1 Protein backbone (i)

In the following table, the Percentiles column shows the percent Ramachandran outliers of the chain as a percentile score with respect to all X-ray entries followed by that with respect to entries of similar resolution.

The Analysed column shows the number of residues for which the backbone conformation was analysed, and the total number of residues.

| Mol | Chain | Analysed | Favoured | Allowed | Outliers | Percentiles |
|---|---|---|---|---|---|---|
| 1 | A | 205/219 (94%) | 191 (93%) | 13 (6%) | 1 (0%) | 25 59 |

*Continued on next page...*

For Manuscript Review

*Continued from previous page...*

| Mol | Chain | Analysed | Favoured | Allowed | Outliers | Percentiles | |
|---|---|---|---|---|---|---|---|
| 1 | S | 202/219 (92%) | 187 (93%) | 15 (7%) | 0 | 100 | 100 |
| 2 | B | 236/250 (94%) | 222 (94%) | 14 (6%) | 0 | 100 | 100 |
| 2 | T | 236/250 (94%) | 222 (94%) | 14 (6%) | 0 | 100 | 100 |
| 3 | C | 276/281 (98%) | 255 (92%) | 21 (8%) | 0 | 100 | 100 |
| 4 | D | 97/99 (98%) | 85 (88%) | 12 (12%) | 0 | 100 | 100 |
| 5 | P | 7/9 (78%) | 6 (86%) | 1 (14%) | 0 | 100 | 100 |
| All | All | 1259/1327 (95%) | 1168 (93%) | 90 (7%) | 1 (0%) | 48 | 79 |

All (1) Ramachandran outliers are listed below:

| Mol | Chain | Res | Type |
|---|---|---|---|
| 1 | A | 97 | GLY |

##### 5.3.2 Protein sidechains ⓘ

In the following table, the Percentiles column shows the percent sidechain outliers of the chain as a percentile score with respect to all X-ray entries followed by that with respect to entries of similar resolution.

The Analysed column shows the number of residues for which the sidechain conformation was analysed, and the total number of residues.

| Mol | Chain | Analysed | Rotameric | Outliers | Percentiles | |
|---|---|---|---|---|---|---|
| 1 | A | 163/196 (83%) | 163 (100%) | 0 | 100 | 100 |
| 1 | S | 167/196 (85%) | 167 (100%) | 0 | 100 | 100 |
| 2 | B | 210/222 (95%) | 210 (100%) | 0 | 100 | 100 |
| 2 | T | 206/222 (93%) | 206 (100%) | 0 | 100 | 100 |
| 3 | C | 189/239 (79%) | 188 (100%) | 1 (0%) | 86 | 93 |
| 4 | D | 86/94 (92%) | 86 (100%) | 0 | 100 | 100 |
| 5 | P | 8/8 (100%) | 8 (100%) | 0 | 100 | 100 |
| All | All | 1029/1177 (87%) | 1028 (100%) | 1 (0%) | 92 | 97 |

All (1) residues with a non-rotameric sidechain are listed below:

| Mol | Chain | Res | Type |
|---|---|---|---|
| 3 | C | 164 | CYS |

For Manuscript Review

Sometimes sidechains can be flipped to improve hydrogen bonding and reduce clashes. All (11) such sidechains are listed below:

| Mol | Chain | Res | Type |
|---|---|---|---|
| 2 | B | 28 | HIS |
| 2 | B | 153 | HIS |
| 3 | C | 3 | HIS |
| 3 | C | 42 | ASN |
| 3 | C | 97 | GLN |
| 3 | C | 188 | HIS |
| 5 | P | 3 | ASN |
| 5 | P | 5 | ASN |
| 2 | T | 28 | HIS |
| 2 | T | 76 | HIS |
| 2 | T | 180 | ASN |

##### 5.3.3 RNA (i)

There are no RNA molecules in this entry.

#### 5.4 Non-standard residues in protein, DNA, RNA chains (i)

There are no non-standard protein/DNA/RNA residues in this entry.

#### 5.5 Carbohydrates (i)

There are no oligosaccharides in this entry.

#### 5.6 Ligand geometry (i)

4 ligands are modelled in this entry.

In the following table, the Counts columns list the number of bonds (or angles) for which Mogul statistics could be retrieved, the number of bonds (or angles) that are observed in the model and the number of bonds (or angles) that are defined in the Chemical Component Dictionary. The Link column lists molecule types, if any, to which the group is linked. The Z score for a bond length (or angle) is the number of standard deviations the observed value is removed from the expected value. A bond length (or angle) with $|Z| > 2$ is considered an outlier worth inspection. RMSZ is the root-mean-square of all Z scores of the bond lengths (or angles).

| Mol | Type | Chain | Res | Link | Bond lengths | | | Bond angles | | |
|---|---|---|---|---|---|---|---|---|---|---|
| | | | | | Counts | RMSZ | $\#\|Z\| > 2$ | Counts | RMSZ | $\#\|Z\| > 2$ |
| 6 | NAG | B | 301 | 2 | 14,14,15 | 0.74 | 0 | 17,19,21 | 0.90 | 0 |

For Manuscript Review

| Mol | Type | Chain | Res | Link | Bond lengths | | | Bond angles | | |
|---|---|---|---|---|---|---|---|---|---|---|
| | | | | | Counts | RMSZ | #\|Z\| > 2 | Counts | RMSZ | #\|Z\| > 2 |
| 6 | NAG | T | 301 | 2 | 14,14,15 | 0.69 | 0 | 17,19,21 | 0.99 | 1 (5%) |
| 6 | NAG | A | 301 | 1 | 14,14,15 | 0.75 | 0 | 17,19,21 | 0.84 | 0 |
| 6 | NAG | B | 302 | 2 | 14,14,15 | 0.75 | 0 | 17,19,21 | 1.29 | 2 (11%) |

In the following table, the Chirals column lists the number of chiral outliers, the number of chiral centers analysed, the number of these observed in the model and the number defined in the Chemical Component Dictionary. Similar counts are reported in the Torsion and Rings columns. '-' means no outliers of that kind were identified.

| Mol | Type | Chain | Res | Link | Chirals | Torsions | Rings |
|---|---|---|---|---|---|---|---|
| 6 | NAG | B | 301 | 2 | - | 0/6/23/26 | 0/1/1/1 |
| 6 | NAG | T | 301 | 2 | - | 0/6/23/26 | 0/1/1/1 |
| 6 | NAG | A | 301 | 1 | - | 2/6/23/26 | 0/1/1/1 |
| 6 | NAG | B | 302 | 2 | - | 0/6/23/26 | 0/1/1/1 |

There are no bond length outliers.

All (3) bond angle outliers are listed below:

| Mol | Chain | Res | Type | Atoms | Z | Observed($^o$) | Ideal($^o$) |
|---|---|---|---|---|---|---|---|
| 6 | B | 302 | NAG | C1-O5-C5 | 4.08 | 117.65 | 112.19 |
| 6 | T | 301 | NAG | O5-C1-C2 | -2.18 | 107.92 | 111.29 |
| 6 | B | 302 | NAG | C2-N2-C7 | 2.10 | 125.71 | 122.90 |

There are no chirality outliers.

All (2) torsion outliers are listed below:

| Mol | Chain | Res | Type | Atoms |
|---|---|---|---|---|
| 6 | A | 301 | NAG | O5-C5-C6-O6 |
| 6 | A | 301 | NAG | C4-C5-C6-O6 |

There are no ring outliers.

2 monomers are involved in 2 short contacts:

| Mol | Chain | Res | Type | Clashes | Symm-Clashes |
|---|---|---|---|---|---|
| 6 | B | 301 | NAG | 1 | 0 |
| 6 | T | 301 | NAG | 1 | 0 |

For Manuscript Review

#### 5.7 Other polymers (i)

There are no such residues in this entry.

#### 5.8 Polymer linkage issues (i)

There are no chain breaks in this entry.

For Manuscript Review

### 6 Fit of model and data (i)

#### 6.1 Protein, DNA and RNA chains (i)

In the following table, the column labelled '#RSRZ> 2' contains the number (and percentage) of RSRZ outliers, followed by percent RSRZ outliers for the chain as percentile scores relative to all X-ray entries and entries of similar resolution. The OWAB column contains the minimum, median, $95^{th}$ percentile and maximum values of the occupancy-weighted average B-factor per residue. The column labelled 'Q< 0.9' lists the number of (and percentage) of residues with an average occupancy less than 0.9.

| Mol | Chain | Analysed | <RSRZ> | #RSRZ>2 | | | OWAB(Å$^2$) | Q<0.9 |
|---|---|---|---|---|---|---|---|---|
| 1 | A | 207/219 (94%) | -0.42 | 0 | 100 | 100 | 63, 112, 170, 225 | 0 |
| 1 | S | 204/219 (93%) | -0.40 | 1 (0%) | 87 | 72 | 79, 128, 186, 238 | 0 |
| 2 | B | 238/250 (95%) | -0.39 | 2 (0%) | 82 | 64 | 81, 115, 164, 222 | 0 |
| 2 | T | 238/250 (95%) | -0.42 | 3 (1%) | 74 | 53 | 88, 139, 196, 305 | 0 |
| 3 | C | 278/281 (98%) | -0.35 | 0 | 100 | 100 | 65, 123, 211, 324 | 0 |
| 4 | D | 99/99 (100%) | -0.28 | 0 | 100 | 100 | 106, 153, 204, 262 | 0 |
| 5 | P | 9/9 (100%) | -0.21 | 0 | 100 | 100 | 86, 94, 109, 110 | 0 |
| All | All | 1273/1327 (95%) | -0.38 | 6 (0%) | 87 | 72 | 63, 126, 193, 324 | 0 |

All (6) RSRZ outliers are listed below:

| Mol | Chain | Res | Type | RSRZ |
|---|---|---|---|---|
| 2 | T | 96 | LEU | 2.8 |
| 2 | B | 185 | LEU | 2.6 |
| 1 | S | 140 | LEU | 2.3 |
| 2 | B | 128 | GLU | 2.3 |
| 2 | T | 27 | GLY | 2.3 |
| 2 | T | 32 | TYR | 2.3 |

#### 6.2 Non-standard residues in protein, DNA, RNA chains (i)

There are no non-standard protein/DNA/RNA residues in this entry.

#### 6.3 Carbohydrates (i)

There are no oligosaccharides in this entry.

For Manuscript Review

#### 6.4 Ligands (i)

In the following table, the Atoms column lists the number of modelled atoms in the group and the number defined in the chemical component dictionary. The B-factors column lists the minimum, median, $95^{th}$ percentile and maximum values of B factors of atoms in the group. The column labelled 'Q< 0.9' lists the number of atoms with occupancy less than 0.9.

| Mol | Type | Chain | Res | Atoms | RSCC | RSR | B-factors($Å^2$) | Q<0.9 |
|---|---|---|---|---|---|---|---|---|
| 6 | NAG | B | 302 | 14/15 | 0.57 | 0.09 | 118,126,135,137 | 0 |
| 6 | NAG | T | 301 | 14/15 | 0.63 | 0.07 | 165,168,176,178 | 0 |
| 6 | NAG | A | 301 | 14/15 | 0.76 | 0.09 | 115,128,137,150 | 0 |
| 6 | NAG | B | 301 | 14/15 | 0.93 | 0.06 | 99,105,111,114 | 0 |

#### 6.5 Other polymers (i)

There are no such residues in this entry.

For Manuscript Review

### Full wwPDB X-ray Structure Validation Report (i)

Mar 20, 2025 – 02:13 PM EDT

| | | |
|---|---|---|
| PDB ID | : | 9NUA |
| Title | : | Crystal structure of NP207 TCR |
| Deposited on | : | 2025-03-19 |
| Resolution | : | 2.50 Å(reported) |

**This wwPDB validation report is for manuscript review**

This is a Full wwPDB X-ray Structure Validation Report.

This report is produced by the wwPDB biocuration pipeline after annotation of the structure.

We welcome your comments at **
A user guide is available at
https://www.wwpdb.org/validation/2017/XrayValidationReportHelp
with specific help available everywhere you see the (i) symbol.

The types of validation reports are described at
http://www.wwpdb.org/validation/2017/FAQs#types.

The following versions of software and data (see references (i)) were used in the production of this report:

| | | |
|---|---|---|
| MolProbity | : | 4.02b-467 |
| Mogul | : | 2022.3.0, CSD as543be (2022) |
| Xtriage (Phenix) | : | 1.21 |
| EDS | : | 3.0 |
| buster-report | : | 1.1.7 (2018) |
| Percentile statistics | : | 20231227.v01 (using entries in the PDB archive December 27th 2023) |
| CCP4 | : | 9.0.004 (Gargrove) |
| Density-Fitness | : | 1.0.11 |
| Ideal geometry (proteins) | : | Engh & Huber (2001) |
| Ideal geometry (DNA, RNA) | : | Parkinson et al. (1996) |

For Manuscript Review

### 1 Overall quality at a glance (i)

The following experimental techniques were used to determine the structure:
*X-RAY DIFFRACTION*

The reported resolution of this entry is 2.50 Å.

Percentile scores (ranging between 0-100) for global validation metrics of the entry are shown in the following graphic. The table shows the number of entries on which the scores are based.

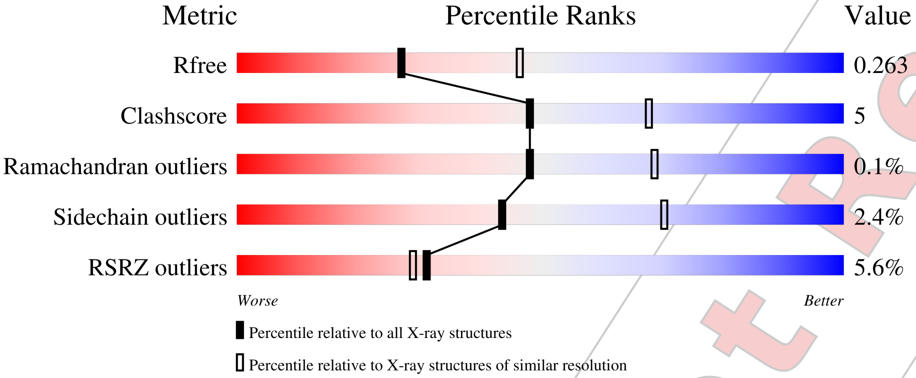


| Metric | Whole archive (#Entries) | Similar resolution (#Entries, resolution range(Å)) |
|---|---|---|
| $R_{free}$ | 164625 | 5504 (2.50-2.50) |
| Clashscore | 180529 | 6282 (2.50-2.50) |
| Ramachandran outliers | 177936 | 6191 (2.50-2.50) |
| Sidechain outliers | 177891 | 6193 (2.50-2.50) |
| RSRZ outliers | 164620 | 5504 (2.50-2.50) |

The table below summarises the geometric issues observed across the polymeric chains and their fit to the electron density. The red, orange, yellow and green segments of the lower bar indicate the fraction of residues that contain outliers for >=3, 2, 1 and 0 types of geometric quality criteria respectively. A grey segment represents the fraction of residues that are not modelled. The numeric value for each fraction is indicated below the corresponding segment, with a dot representing fractions <=5% The upper red bar (where present) indicates the fraction of residues that have poor fit to the electron density. The numeric value is given above the bar.

| Mol | Chain | Length | Quality of chain |
|---|---|---|---|
| 1 | A | 219 | 3%<br>82% 12% 5% |
| 1 | C | 219 | 9%<br>73% 16% 11% |
| 2 | B | 250 | 86% 9% • • |
| 2 | D | 250 | 9%<br>78% 18% • |

Validation Pipeline (wwPDB-VP) : 2.41.5

For Manuscript Review

#### 2 Entry composition (i)

There are 6 unique types of molecules in this entry. The entry contains 7094 atoms, of which 0 are hydrogens and 0 are deuteriums.

In the tables below, the ZeroOcc column contains the number of atoms modelled with zero occupancy, the AltConf column contains the number of residues with at least one atom in alternate conformation and the Trace column contains the number of residues modelled with at most 2 atoms.

- Molecule 1 is a protein called NP207 TCR alpha-chain.

| Mol | Chain | Residues | Atoms | ZeroOcc | AltConf | Trace |
|---|---|---|---|---|---|---|
| 1 | A | 208 | Total C N O S<br>1610 1013 258 328 11 | 0 | 0 | 0 |
| 1 | C | 194 | Total C N O S<br>1456 923 231 293 9 | 0 | 0 | 0 |

- Molecule 2 is a protein called NP207 TCR beta-chain.

| Mol | Chain | Residues | Atoms | ZeroOcc | AltConf | Trace |
|---|---|---|---|---|---|---|
| 2 | B | 239 | Total C N O S<br>1899 1209 330 352 8 | 0 | 0 | 0 |
| 2 | D | 239 | Total C N O S<br>1870 1186 325 351 8 | 0 | 0 | 0 |

- Molecule 3 is 2-acetamido-2-deoxy-beta-D-glucopyranose (three-letter code: NAG) (formula: $C_8H_{15}NO_6$) (labeled as "Ligand of Interest" by depositor).

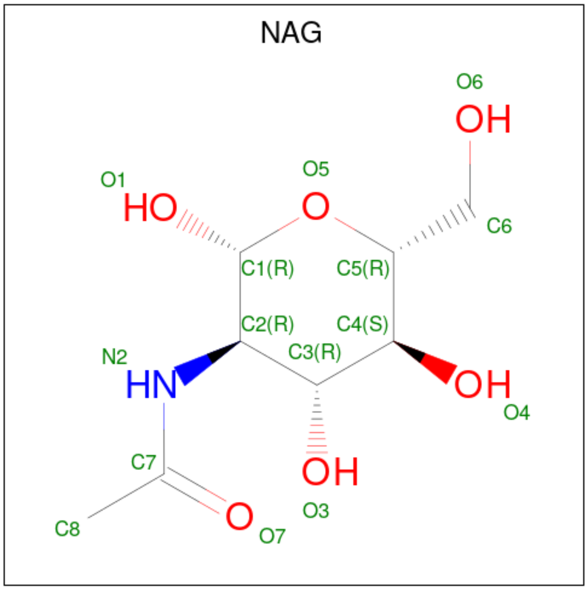


For Manuscript Review

| Mol | Chain | Residues | Atoms | ZeroOcc | AltConf |
|---|---|---|---|---|---|
| 3 | A | 1 | Total C N O<br>14 8 1 5 | 0 | 0 |
| 3 | A | 1 | Total C N O<br>14 8 1 5 | 0 | 0 |
| 3 | A | 1 | Total C N O<br>14 8 1 5 | 0 | 0 |
| 3 | B | 1 | Total C N O<br>14 8 1 5 | 0 | 0 |
| 3 | B | 1 | Total C N O<br>14 8 1 5 | 0 | 0 |
| 3 | B | 1 | Total C N O<br>14 8 1 5 | 0 | 0 |
| 3 | B | 1 | Total C N O<br>14 8 1 5 | 0 | 0 |
| 3 | C | 1 | Total C N O<br>14 8 1 5 | 0 | 0 |
| 3 | D | 1 | Total C N O<br>14 8 1 5 | 0 | 0 |
| 3 | D | 1 | Total C N O<br>14 8 1 5 | 0 | 0 |
| 3 | D | 1 | Total C N O<br>14 8 1 5 | 0 | 0 |

- Molecule 4 is GLYCEROL (three-letter code: GOL) (formula: $C_3H_8O_3$).

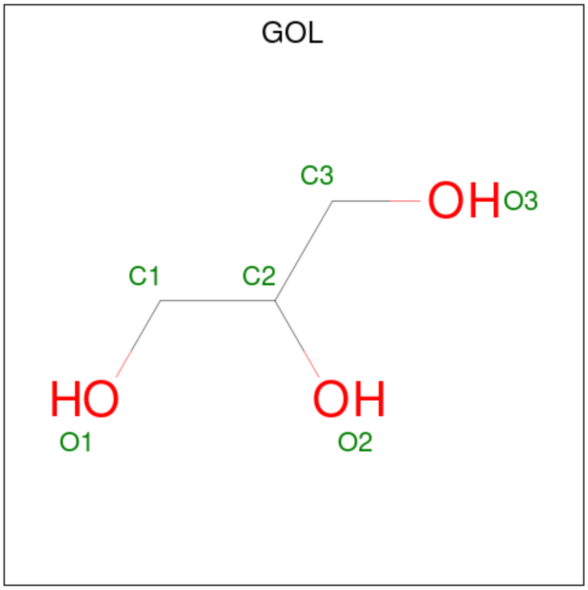


| Mol | Chain | Residues | Atoms | ZeroOcc | AltConf |
|---|---|---|---|---|---|
| 4 | A | 1 | Total C O<br>6 3 3 | 0 | 0 |

*Continued on next page...*

For Manuscript Review

*Continued from previous page...*

| Mol | Chain | Residues | Atoms | ZeroOcc | AltConf |
|---|---|---|---|---|---|
| 4 | B | 1 | Total C O<br>6 3 3 | 0 | 0 |
| 4 | B | 1 | Total C O<br>6 3 3 | 0 | 0 |
| 4 | B | 1 | Total C O<br>6 3 3 | 0 | 0 |
| 4 | C | 1 | Total C O<br>6 3 3 | 0 | 0 |

- Molecule 5 is CHLORIDE ION (three-letter code: CL) (formula: Cl).

| Mol | Chain | Residues | Atoms | ZeroOcc | AltConf |
|---|---|---|---|---|---|
| 5 | B | 5 | Total Cl<br>5 5 | 0 | 0 |
| 5 | D | 3 | Total Cl<br>3 3 | 0 | 0 |

- Molecule 6 is water.

| Mol | Chain | Residues | Atoms | ZeroOcc | AltConf |
|---|---|---|---|---|---|
| 6 | A | 22 | Total O<br>22 22 | 0 | 0 |
| 6 | B | 25 | Total O<br>25 25 | 0 | 0 |
| 6 | C | 7 | Total O<br>7 7 | 0 | 0 |
| 6 | D | 13 | Total O<br>13 13 | 0 | 0 |

For Manuscript Review

### 3 Residue-property plots (i)

These plots are drawn for all protein, RNA, DNA and oligosaccharide chains in the entry. The first graphic for a chain summarises the proportions of the various outlier classes displayed in the second graphic. The second graphic shows the sequence view annotated by issues in geometry and electron density. Residues are color-coded according to the number of geometric quality criteria for which they contain at least one outlier: green = 0, yellow = 1, orange = 2 and red = 3 or more. A red dot above a residue indicates a poor fit to the electron density (RSRZ > 2). Stretches of 2 or more consecutive residues without any outlier are shown as a green connector. Residues present in the sample, but not in the model, are shown in grey.

• Molecule 1: NP207 TCR alpha-chain

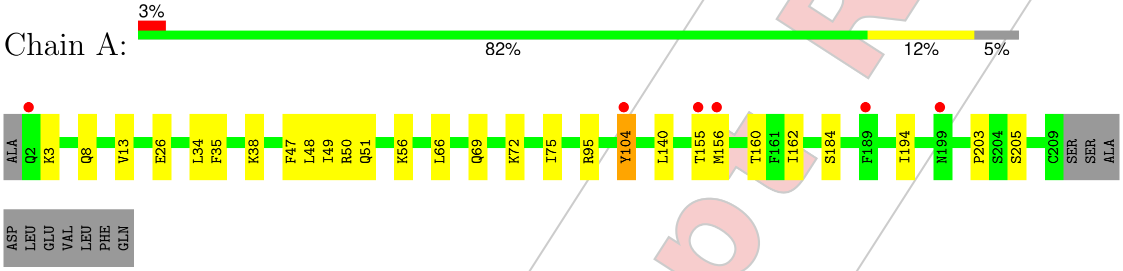


• Molecule 1: NP207 TCR alpha-chain

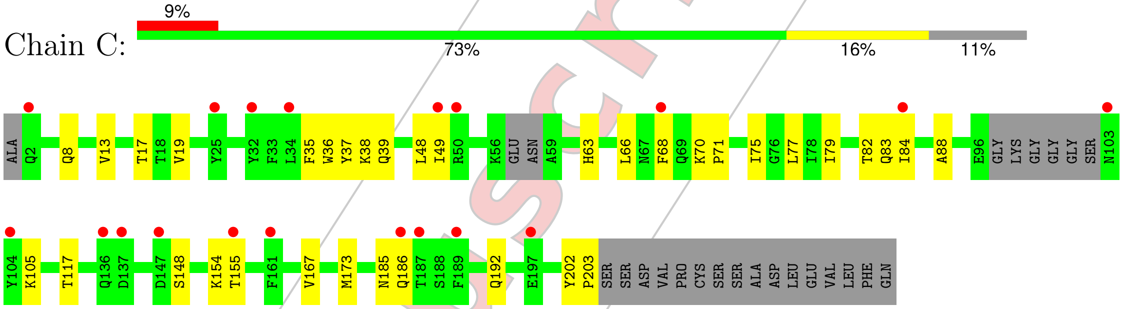


• Molecule 2: NP207 TCR beta-chain

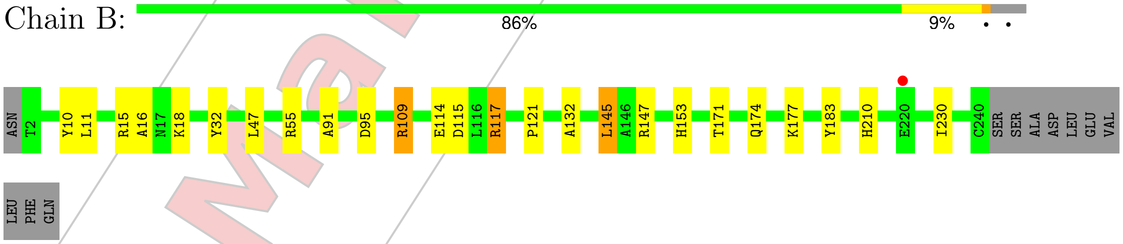


• Molecule 2: NP207 TCR beta-chain

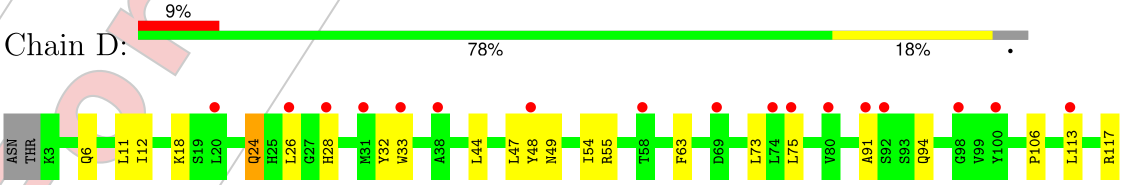


For Manuscript Review

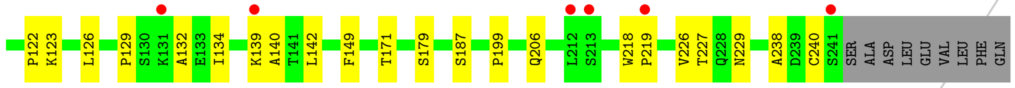


For Manuscript Review

### 4 Data and refinement statistics (i)

| Property | Value | Source |
|---|---|---|
| Space group | P 1 21 1 | Depositor |
| Cell constants<br>a, b, c, $\alpha$, $\beta$, $\gamma$ | 62.12Å 126.02Å 66.63Å<br>90.00° 110.23° 90.00° | Depositor |
| Resolution (Å) | 30.81 – 2.50<br>30.81 – 2.50 | Depositor<br>EDS |
| % Data completeness<br>(in resolution range) | 97.1 (30.81-2.50)<br>97.1 (30.81-2.50) | Depositor<br>EDS |
| $R_{merge}$ | 0.08 | Depositor |
| $R_{sym}$ | (Not available) | Depositor |
| $< I/\sigma(I) >$ [1] | 1.92 (at 2.51Å) | Xtriage |
| Refinement program | PHENIX 1.21.2_5419 | Depositor |
| R, $R_{free}$ | 0.214 , 0.263<br>0.213 , 0.263 | Depositor<br>DCC |
| $R_{free}$ test set | 1651 reflections (5.03%) | wwPDB-VP |
| Wilson B-factor (Å$^2$) | 53.0 | Xtriage |
| Anisotropy | 0.022 | Xtriage |
| Bulk solvent $k_{sol}$(e/Å$^3$), $B_{sol}$(Å$^2$) | 0.35 , 69.1 | EDS |
| L-test for twinning[2] | $< \|L\| > = 0.47$, $< L^2 > = 0.30$ | Xtriage |
| Estimated twinning fraction | No twinning to report. | Xtriage |
| $F_o$,$F_c$ correlation | 0.93 | EDS |
| Total number of atoms | 7094 | wwPDB-VP |
| Average B, all atoms (Å$^2$) | 75.0 | wwPDB-VP |

Xtriage's analysis on translational NCS is as follows: *The largest off-origin peak in the Patterson function is 4.10% of the height of the origin peak. No significant pseudotranslation is detected.*

[1] Intensities estimated from amplitudes.

[2] Theoretical values of $< |L| >$, $< L^2 >$ for acentric reflections are 0.5, 0.333 respectively for untwinned datasets, and 0.375, 0.2 for perfectly twinned datasets.

For Manuscript Review

### 5 Model quality (i)

#### 5.1 Standard geometry (i)

Bond lengths and bond angles in the following residue types are not validated in this section: NAG, CL, GOL

The Z score for a bond length (or angle) is the number of standard deviations the observed value is removed from the expected value. A bond length (or angle) with $|Z| > 5$ is considered an outlier worth inspection. RMSZ is the root-mean-square of all Z scores of the bond lengths (or angles).

| Mol | Chain | Bond lengths | | Bond angles | |
|---|---|---|---|---|---|
| | | RMSZ | #$\|Z\|$ >5 | RMSZ | #$\|Z\|$ >5 |
| 1 | A | 0.27 | 0/1642 | 0.49 | 0/2228 |
| 1 | C | 0.26 | 0/1484 | 0.48 | 0/2023 |
| 2 | B | 0.26 | 0/1955 | 0.48 | 0/2660 |
| 2 | D | 0.27 | 0/1925 | 0.51 | 0/2624 |
| All | All | 0.26 | 0/7006 | 0.49 | 0/9535 |

There are no bond length outliers.

There are no bond angle outliers.

There are no chirality outliers.

There are no planarity outliers.

#### 5.2 Too-close contacts (i)

In the following table, the Non-H and H(model) columns list the number of non-hydrogen atoms and hydrogen atoms in the chain respectively. The H(added) column lists the number of hydrogen atoms added and optimized by MolProbity. The Clashes column lists the number of clashes within the asymmetric unit, whereas Symm-Clashes lists symmetry-related clashes.

| Mol | Chain | Non-H | H(model) | H(added) | Clashes | Symm-Clashes |
|---|---|---|---|---|---|---|
| 1 | A | 1610 | 0 | 1539 | 17 | 0 |
| 1 | C | 1456 | 0 | 1341 | 22 | 0 |
| 2 | B | 1899 | 0 | 1818 | 15 | 0 |
| 2 | D | 1870 | 0 | 1748 | 27 | 0 |
| 3 | A | 42 | 0 | 39 | 0 | 0 |
| 3 | B | 56 | 0 | 52 | 1 | 0 |
| 3 | C | 14 | 0 | 13 | 0 | 0 |
| 3 | D | 42 | 0 | 39 | 2 | 0 |
| 4 | A | 6 | 0 | 8 | 0 | 0 |

*Continued on next page...*

For Manuscript Review

*Continued from previous page...*

| Mol | Chain | Non-H | H(model) | H(added) | Clashes | Symm-Clashes |
|---|---|---|---|---|---|---|
| 4 | B | 18 | 0 | 24 | 1 | 0 |
| 4 | C | 6 | 0 | 8 | 0 | 0 |
| 5 | B | 5 | 0 | 0 | 0 | 0 |
| 5 | D | 3 | 0 | 0 | 1 | 0 |
| 6 | A | 22 | 0 | 0 | 1 | 0 |
| 6 | B | 25 | 0 | 0 | 0 | 0 |
| 6 | C | 7 | 0 | 0 | 0 | 0 |
| 6 | D | 13 | 0 | 0 | 0 | 0 |
| All | All | 7094 | 0 | 6629 | 73 | 0 |

The all-atom clashscore is defined as the number of clashes found per 1000 atoms (including hydrogen atoms). The all-atom clashscore for this structure is 5.

All (73) close contacts within the same asymmetric unit are listed below, sorted by their clash magnitude.

| Atom-1 | Atom-2 | Interatomic distance (Å) | Clash overlap (Å) |
|---|---|---|---|
| 1:C:173:MET:HE1 | 2:D:139:LYS:HE2 | 1.59 | 0.85 |
| 2:B:47:LEU:HB3 | 2:B:55:ARG:HB2 | 1.67 | 0.75 |
| 2:B:32:TYR:HB2 | 2:B:91:ALA:HB3 | 1.75 | 0.69 |
| 1:C:203:PRO:HB2 | 2:D:132:ALA:HB1 | 1.77 | 0.66 |
| 2:D:179:SER:HB2 | 3:D:302:NAG:H62 | 1.80 | 0.63 |
| 2:D:126:LEU:HD21 | 2:D:142:LEU:HD23 | 1.80 | 0.62 |
| 2:D:12:ILE:HG23 | 2:D:113:LEU:HD23 | 1.82 | 0.61 |
| 2:D:24:GLN:HG2 | 2:D:26:LEU:H | 1.65 | 0.61 |
| 2:B:16:ALA:HB3 | 3:B:301:NAG:H82 | 1.80 | 0.61 |
| 2:D:33:TRP:CE2 | 2:D:73:LEU:HD23 | 2.35 | 0.60 |
| 2:D:32:TYR:HB2 | 2:D:91:ALA:HB3 | 1.83 | 0.59 |
| 1:A:38:LYS:HB2 | 1:A:48:LEU:HD11 | 1.86 | 0.58 |
| 2:B:145:LEU:HD22 | 2:B:147:ARG:HG3 | 1.86 | 0.58 |
| 1:A:69:GLN:HG2 | 1:A:72:LYS:HD2 | 1.86 | 0.56 |
| 2:D:24:GLN:HE21 | 2:D:26:LEU:N | 2.03 | 0.56 |
| 2:D:218:TRP:CD2 | 2:D:219:PRO:HD2 | 2.39 | 0.56 |
| 2:D:171:THR:HG23 | 2:D:187:SER:HB2 | 1.86 | 0.56 |
| 1:A:104:TYR:HB2 | 2:B:55:ARG:HE | 1.71 | 0.56 |
| 1:A:49:ILE:HD13 | 1:A:66:LEU:HG | 1.86 | 0.56 |
| 2:D:129:PRO:HB2 | 2:D:134:ILE:HD11 | 1.88 | 0.56 |
| 1:A:95:ARG:CZ | 1:A:104:TYR:HA | 2.40 | 0.53 |
| 1:C:66:LEU:HD12 | 1:C:75:ILE:HD11 | 1.90 | 0.52 |
| 1:C:192:GLN:HA | 1:C:202:TYR:CE2 | 2.45 | 0.52 |
| 2:D:28:HIS:H | 2:D:94:GLN:HB3 | 1.75 | 0.51 |
| 1:C:17:THR:O | 1:C:82:THR:HG23 | 2.11 | 0.50 |

*Continued on next page...*

For Manuscript Review

Continued from previous page...

| Atom-1 | Atom-2 | Interatomic distance (Å) | Clash overlap (Å) |
|---|---|---|---|
| 2:B:171:THR:H | 4:B:305:GOL:H2 | 1.75 | 0.50 |
| 1:C:49:ILE:HD13 | 1:C:66:LEU:HD23 | 1.94 | 0.50 |
| 2:D:11:LEU:HD21 | 2:D:18:LYS:HD2 | 1.93 | 0.50 |
| 1:C:68:PHE:HD2 | 1:C:75:ILE:HD13 | 1.77 | 0.49 |
| 2:D:122:PRO:HB3 | 2:D:149:PHE:HB3 | 1.94 | 0.49 |
| 1:C:68:PHE:CD2 | 1:C:75:ILE:HD13 | 2.48 | 0.49 |
| 1:A:51:GLN:HB2 | 1:A:66:LEU:HD13 | 1.95 | 0.48 |
| 1:C:36:TRP:HB3 | 1:C:77:LEU:HD12 | 1.95 | 0.48 |
| 1:A:13:VAL:HG12 | 1:C:8:GLN:HG2 | 1.94 | 0.48 |
| 1:C:19:VAL:HG12 | 1:C:79:ILE:HB | 1.94 | 0.48 |
| 2:B:177:LYS:HG3 | 2:B:183:TYR:CE2 | 2.49 | 0.47 |
| 1:A:8:GLN:HG2 | 1:C:13:VAL:HG12 | 1.96 | 0.47 |
| 1:A:34:LEU:HD13 | 1:A:75:ILE:HG21 | 1.97 | 0.47 |
| 1:A:160:THR:HG22 | 1:A:184:SER:HB2 | 1.96 | 0.47 |
| 2:B:10:TYR:CE2 | 2:B:109:ARG:HD3 | 2.50 | 0.47 |
| 1:C:38:LYS:HB2 | 1:C:48:LEU:HD11 | 1.97 | 0.46 |
| 2:D:129:PRO:HG2 | 2:D:140:ALA:HB1 | 1.97 | 0.46 |
| 2:D:199:PRO:HG3 | 2:D:238:ALA:HB2 | 1.98 | 0.46 |
| 2:D:226:VAL:HG12 | 2:D:227:THR:H | 1.81 | 0.46 |
| 1:C:84:ILE:HD12 | 1:C:84:ILE:H | 1.81 | 0.46 |
| 1:C:105:LYS:HA | 2:D:44:LEU:HD12 | 1.98 | 0.46 |
| 2:D:33:TRP:CD2 | 2:D:73:LEU:HD23 | 2.51 | 0.45 |
| 2:D:63:PHE:HD1 | 2:D:75:LEU:HD11 | 1.80 | 0.45 |
| 2:B:114:GLU:H | 2:B:114:GLU:CD | 2.19 | 0.45 |
| 1:A:3:LYS:HB3 | 1:A:26:GLU:HB2 | 1.99 | 0.45 |
| 2:B:153:HIS:HB3 | 2:B:210:HIS:HB2 | 1.99 | 0.45 |
| 1:A:35:PHE:CE2 | 1:A:50:ARG:HG3 | 2.52 | 0.44 |
| 1:C:117:THR:HB | 1:C:148:SER:HB3 | 1.99 | 0.44 |
| 1:A:162:ILE:HD11 | 1:A:194:ILE:HG22 | 1.99 | 0.43 |
| 1:A:203:PRO:HB2 | 2:B:132:ALA:HB1 | 2.00 | 0.43 |
| 1:A:140:LEU:HD23 | 2:B:145:LEU:HD12 | 2.01 | 0.43 |
| 2:D:47:LEU:HD12 | 2:D:54:ILE:HB | 2.00 | 0.43 |
| 2:B:115:ASP:OD1 | 2:B:117:ARG:HG2 | 2.18 | 0.43 |
| 1:C:70:LYS:N | 1:C:71:PRO:HD2 | 2.34 | 0.43 |
| 2:D:6:GLN:HB2 | 2:D:106:PRO:HG2 | 2.00 | 0.42 |
| 2:D:240:CYS:HA | 5:D:305:CL:CL | 2.56 | 0.42 |
| 1:A:47:PHE:HZ | 1:A:50:ARG:HB2 | 1.85 | 0.42 |
| 1:C:35:PHE:HB3 | 1:C:37:TYR:HE1 | 1.85 | 0.42 |
| 2:D:206:GLN:OE1 | 3:D:303:NAG:H82 | 2.19 | 0.42 |
| 1:C:84:ILE:HG21 | 1:C:167:VAL:HG11 | 2.01 | 0.41 |
| 1:C:154:LYS:H | 1:C:154:LYS:HG2 | 1.64 | 0.41 |

Continued on next page...

For Manuscript Review

*Continued from previous page...*

| Atom-1 | Atom-2 | Interatomic distance (Å) | Clash overlap (Å) |
|---|---|---|---|
| 1:A:205:SER:HB2 | 6:A:418:HOH:O | 2.21 | 0.41 |
| 2:D:47:LEU:HB3 | 2:D:55:ARG:HB2 | 2.03 | 0.41 |
| 2:D:134:ILE:HG13 | 2:D:139:LYS:O | 2.20 | 0.41 |
| 1:C:39:GLN:O | 1:C:88:ALA:HB1 | 2.21 | 0.41 |
| 2:B:11:LEU:HD11 | 2:B:18:LYS:HD3 | 2.02 | 0.41 |
| 1:C:185:ASN:HB3 | 1:C:186:GLN:H | 1.71 | 0.41 |
| 2:B:121:PRO:HB2 | 2:B:230:ILE:HD11 | 2.03 | 0.40 |

There are no symmetry-related clashes.

#### 5.3 Torsion angles (i)

##### 5.3.1 Protein backbone (i)

In the following table, the Percentiles column shows the percent Ramachandran outliers of the chain as a percentile score with respect to all X-ray entries followed by that with respect to entries of similar resolution.

The Analysed column shows the number of residues for which the backbone conformation was analysed, and the total number of residues.

| Mol | Chain | Analysed | Favoured | Allowed | Outliers | Percentiles | |
|---|---|---|---|---|---|---|---|
| 1 | A | 206/219 (94%) | 196 (95%) | 9 (4%) | 1 (0%) | 25 | 44 |
| 1 | C | 188/219 (86%) | 176 (94%) | 12 (6%) | 0 | 100 | 100 |
| 2 | B | 237/250 (95%) | 225 (95%) | 12 (5%) | 0 | 100 | 100 |
| 2 | D | 237/250 (95%) | 218 (92%) | 19 (8%) | 0 | 100 | 100 |
| All | All | 868/938 (92%) | 815 (94%) | 52 (6%) | 1 (0%) | 48 | 69 |

All (1) Ramachandran outliers are listed below:

| Mol | Chain | Res | Type |
|---|---|---|---|
| 1 | A | 104 | TYR |

##### 5.3.2 Protein sidechains (i)

In the following table, the Percentiles column shows the percent sidechain outliers of the chain as a percentile score with respect to all X-ray entries followed by that with respect to entries of similar resolution.

For Manuscript Review

The Analysed column shows the number of residues for which the sidechain conformation was analysed, and the total number of residues.

| Mol | Chain | Analysed | Rotameric | Outliers | Percentiles | |
|---|---|---|---|---|---|---|
| 1 | A | 182/196 (93%) | 179 (98%) | 3 (2%) | 58 | 80 |
| 1 | C | 155/196 (79%) | 152 (98%) | 3 (2%) | 52 | 77 |
| 2 | B | 207/222 (93%) | 201 (97%) | 6 (3%) | 37 | 64 |
| 2 | D | 199/222 (90%) | 193 (97%) | 6 (3%) | 36 | 63 |
| All | All | 743/836 (89%) | 725 (98%) | 18 (2%) | 44 | 70 |

All (18) residues with a non-rotameric sidechain are listed below:

| Mol | Chain | Res | Type |
|---|---|---|---|
| 1 | A | 56 | LYS |
| 1 | A | 155 | THR |
| 1 | A | 156 | MET |
| 2 | B | 15 | ARG |
| 2 | B | 95 | ASP |
| 2 | B | 109 | ARG |
| 2 | B | 117 | ARG |
| 2 | B | 145 | LEU |
| 2 | B | 174 | GLN |
| 1 | C | 63 | HIS |
| 1 | C | 83 | GLN |
| 1 | C | 155 | THR |
| 2 | D | 24 | GLN |
| 2 | D | 48 | TYR |
| 2 | D | 49 | ASN |
| 2 | D | 117 | ARG |
| 2 | D | 123 | LYS |
| 2 | D | 229 | ASN |

Sometimes sidechains can be flipped to improve hydrogen bonding and reduce clashes. All (6) such sidechains are listed below:

| Mol | Chain | Res | Type |
|---|---|---|---|
| 1 | A | 63 | HIS |
| 1 | C | 39 | GLN |
| 1 | C | 121 | GLN |
| 2 | D | 24 | GLN |
| 2 | D | 49 | ASN |
| 2 | D | 208 | GLN |

For Manuscript Review

##### 5.3.3 RNA (i)

There are no RNA molecules in this entry.

#### 5.4 Non-standard residues in protein, DNA, RNA chains (i)

There are no non-standard protein/DNA/RNA residues in this entry.

#### 5.5 Carbohydrates (i)

There are no oligosaccharides in this entry.

#### 5.6 Ligand geometry (i)

Of 24 ligands modelled in this entry, 8 are monoatomic - leaving 16 for Mogul analysis.

In the following table, the Counts columns list the number of bonds (or angles) for which Mogul statistics could be retrieved, the number of bonds (or angles) that are observed in the model and the number of bonds (or angles) that are defined in the Chemical Component Dictionary. The Link column lists molecule types, if any, to which the group is linked. The Z score for a bond length (or angle) is the number of standard deviations the observed value is removed from the expected value. A bond length (or angle) with $|Z| > 2$ is considered an outlier worth inspection. RMSZ is the root-mean-square of all Z scores of the bond lengths (or angles).

| Mol | Type | Chain | Res | Link | Bond lengths | | | Bond angles | | |
|---|---|---|---|---|---|---|---|---|---|---|
| | | | | | Counts | RMSZ | $\#\|Z\| > 2$ | Counts | RMSZ | $\#\|Z\| > 2$ |
| 4 | GOL | A | 304 | - | 5,5,5 | 0.31 | 0 | 5,5,5 | 0.37 | 0 |
| 3 | NAG | B | 303 | 2 | 14,14,15 | 0.68 | 0 | 17,19,21 | 0.97 | 1 (5%) |
| 3 | NAG | D | 303 | 2 | 14,14,15 | 0.77 | 0 | 17,19,21 | 0.85 | 0 |
| 3 | NAG | B | 302 | 2 | 14,14,15 | 0.73 | 0 | 17,19,21 | 0.93 | 1 (5%) |
| 4 | GOL | B | 306 | - | 5,5,5 | 0.33 | 0 | 5,5,5 | 0.44 | 0 |
| 4 | GOL | B | 305 | - | 5,5,5 | 0.34 | 0 | 5,5,5 | 0.38 | 0 |
| 4 | GOL | B | 307 | - | 5,5,5 | 0.32 | 0 | 5,5,5 | 0.43 | 0 |
| 4 | GOL | C | 302 | - | 5,5,5 | 0.35 | 0 | 5,5,5 | 0.44 | 0 |
| 3 | NAG | D | 301 | 2 | 14,14,15 | 0.78 | 0 | 17,19,21 | 1.16 | 2 (11%) |
| 3 | NAG | D | 302 | 2 | 14,14,15 | 0.70 | 0 | 17,19,21 | 0.82 | 0 |
| 3 | NAG | A | 301 | 1 | 14,14,15 | 0.75 | 0 | 17,19,21 | 0.88 | 0 |
| 3 | NAG | B | 301 | 2 | 14,14,15 | 0.70 | 0 | 17,19,21 | 0.87 | 0 |
| 3 | NAG | C | 301 | 1 | 14,14,15 | 0.71 | 0 | 17,19,21 | 0.80 | 0 |
| 3 | NAG | A | 302 | 1 | 14,14,15 | 0.70 | 0 | 17,19,21 | 0.82 | 0 |
| 3 | NAG | B | 304 | 2 | 14,14,15 | 0.71 | 0 | 17,19,21 | 0.82 | 0 |
| 3 | NAG | A | 303 | 1 | 14,14,15 | 0.77 | 0 | 17,19,21 | 1.27 | 1 (5%) |

For Manuscript Review

In the following table, the Chirals column lists the number of chiral outliers, the number of chiral centers analysed, the number of these observed in the model and the number defined in the Chemical Component Dictionary. Similar counts are reported in the Torsion and Rings columns. '-' means no outliers of that kind were identified.

| Mol | Type | Chain | Res | Link | Chirals | Torsions | Rings |
|---|---|---|---|---|---|---|---|
| 4 | GOL | A | 304 | - | - | 1/4/4/4 | - |
| 3 | NAG | B | 303 | 2 | - | 2/6/23/26 | 0/1/1/1 |
| 3 | NAG | D | 303 | 2 | - | 2/6/23/26 | 0/1/1/1 |
| 3 | NAG | B | 302 | 2 | - | 1/6/23/26 | 0/1/1/1 |
| 4 | GOL | B | 306 | - | - | 2/4/4/4 | - |
| 4 | GOL | B | 305 | - | - | 2/4/4/4 | - |
| 4 | GOL | B | 307 | - | - | 0/4/4/4 | - |
| 4 | GOL | C | 302 | - | - | 2/4/4/4 | - |
| 3 | NAG | D | 301 | 2 | - | 1/6/23/26 | 0/1/1/1 |
| 3 | NAG | D | 302 | 2 | - | 1/6/23/26 | 0/1/1/1 |
| 3 | NAG | A | 301 | 1 | - | 0/6/23/26 | 0/1/1/1 |
| 3 | NAG | B | 301 | 2 | - | 0/6/23/26 | 0/1/1/1 |
| 3 | NAG | C | 301 | 1 | - | 0/6/23/26 | 0/1/1/1 |
| 3 | NAG | A | 302 | 1 | - | 0/6/23/26 | 0/1/1/1 |
| 3 | NAG | B | 304 | 2 | - | 0/6/23/26 | 0/1/1/1 |
| 3 | NAG | A | 303 | 1 | - | 0/6/23/26 | 0/1/1/1 |

There are no bond length outliers.

All (5) bond angle outliers are listed below:

| Mol | Chain | Res | Type | Atoms | Z | Observed(°) | Ideal(°) |
|---|---|---|---|---|---|---|---|
| 3 | A | 303 | NAG | C1-O5-C5 | 3.35 | 116.67 | 112.19 |
| 3 | D | 301 | NAG | C1-O5-C5 | 3.01 | 116.22 | 112.19 |
| 3 | B | 303 | NAG | O5-C1-C2 | -2.44 | 107.51 | 111.29 |
| 3 | D | 301 | NAG | C2-N2-C7 | 2.19 | 125.83 | 122.90 |
| 3 | B | 302 | NAG | C1-O5-C5 | 2.04 | 114.92 | 112.19 |

There are no chirality outliers.

All (14) torsion outliers are listed below:

| Mol | Chain | Res | Type | Atoms |
|---|---|---|---|---|
| 4 | B | 305 | GOL | O1-C1-C2-C3 |
| 4 | B | 306 | GOL | O1-C1-C2-C3 |
| 4 | C | 302 | GOL | C1-C2-C3-O3 |
| 3 | B | 303 | NAG | C4-C5-C6-O6 |
| 3 | B | 303 | NAG | O5-C5-C6-O6 |

*Continued on next page...*

For Manuscript Review

*Continued from previous page...*

| Mol | Chain | Res | Type | Atoms |
|---|---|---|---|---|
| 4 | B | 305 | GOL | O1-C1-C2-O2 |
| 4 | B | 306 | GOL | O1-C1-C2-O2 |
| 4 | C | 302 | GOL | O2-C2-C3-O3 |
| 3 | D | 303 | NAG | C4-C5-C6-O6 |
| 3 | B | 302 | NAG | O5-C5-C6-O6 |
| 3 | D | 303 | NAG | O5-C5-C6-O6 |
| 3 | D | 302 | NAG | C1-C2-N2-C7 |
| 3 | D | 301 | NAG | O5-C5-C6-O6 |
| 4 | A | 304 | GOL | O1-C1-C2-C3 |

There are no ring outliers.

4 monomers are involved in 4 short contacts:

| Mol | Chain | Res | Type | Clashes | Symm-Clashes |
|---|---|---|---|---|---|
| 3 | D | 303 | NAG | 1 | 0 |
| 4 | B | 305 | GOL | 1 | 0 |
| 3 | D | 302 | NAG | 1 | 0 |
| 3 | B | 301 | NAG | 1 | 0 |

The following is a two-dimensional graphical depiction of Mogul quality analysis of bond lengths, bond angles, torsion angles, and ring geometry for all instances of the Ligand of Interest. In addition, ligands with molecular weight > 250 and outliers as shown on the validation Tables will also be included. For torsion angles, if less then 5% of the Mogul distribution of torsion angles is within 10 degrees of the torsion angle in question, then that torsion angle is considered an outlier. Any bond that is central to one or more torsion angles identified as an outlier by Mogul will be highlighted in the graph. For rings, the root-mean-square deviation (RMSD) between the ring in question and similar rings identified by Mogul is calculated over all ring torsion angles. If the average RMSD is greater than 60 degrees and the minimal RMSD between the ring in question and any Mogul-identified rings is also greater than 60 degrees, then that ring is considered an outlier. The outliers are highlighted in purple. The color gray indicates Mogul did not find sufficient equivalents in the CSD to analyse the geometry.

For Manuscript Review

#### Ligand NAG B 303

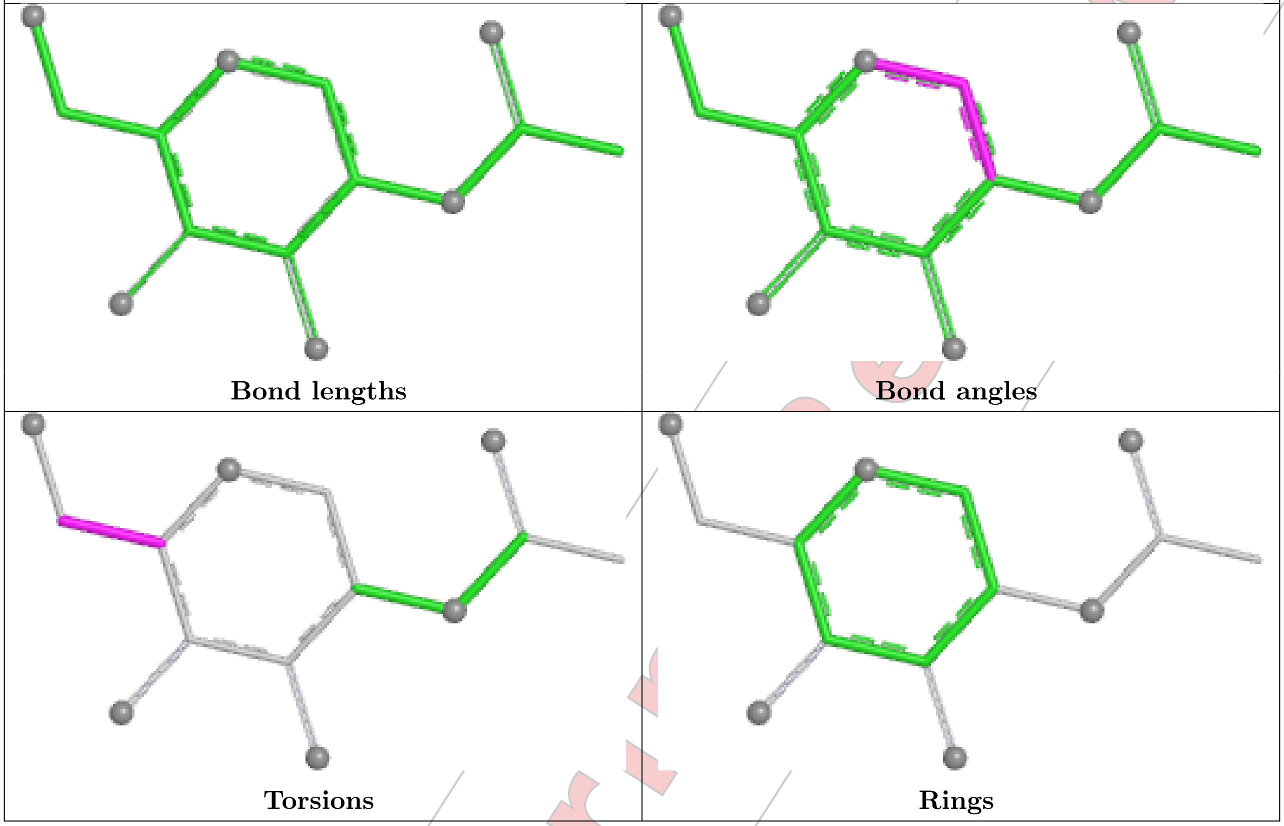


#### Ligand NAG D 303

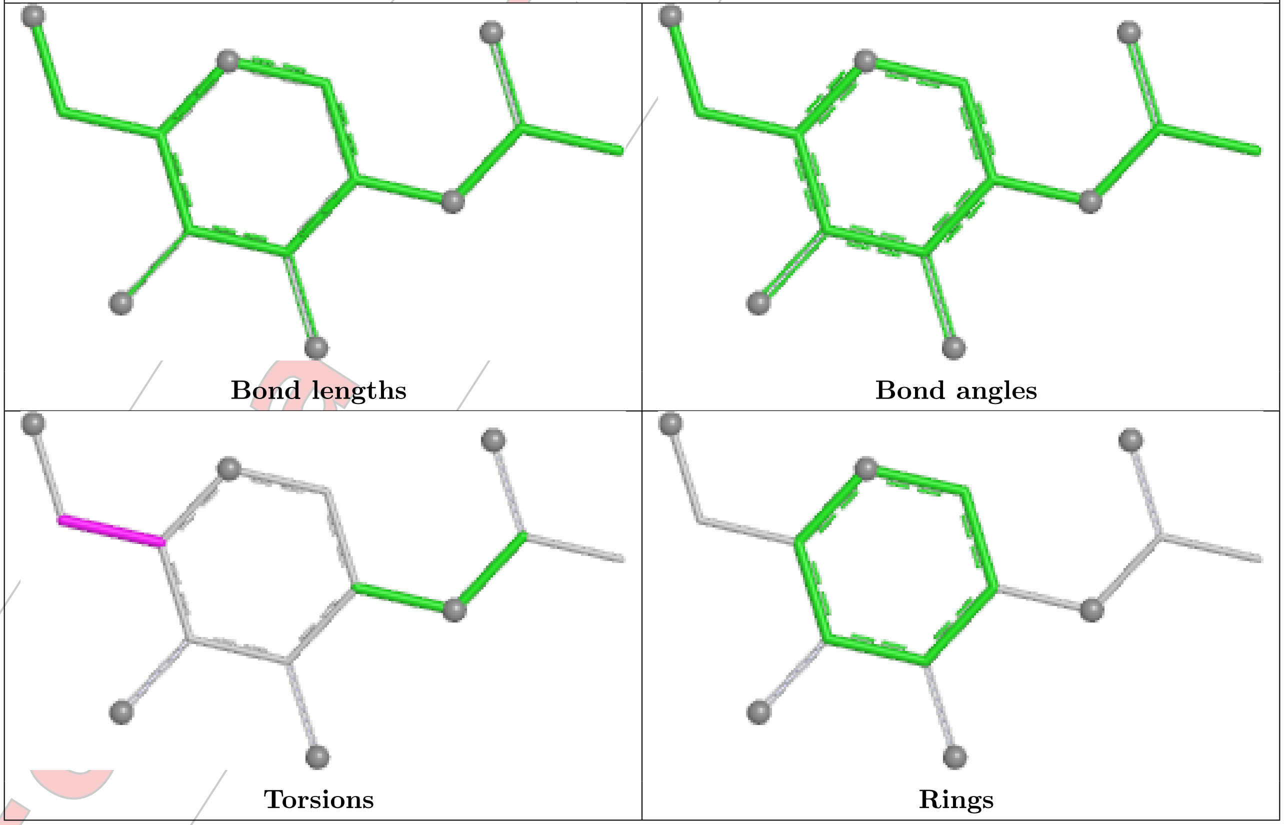

Ligand NAG B 302

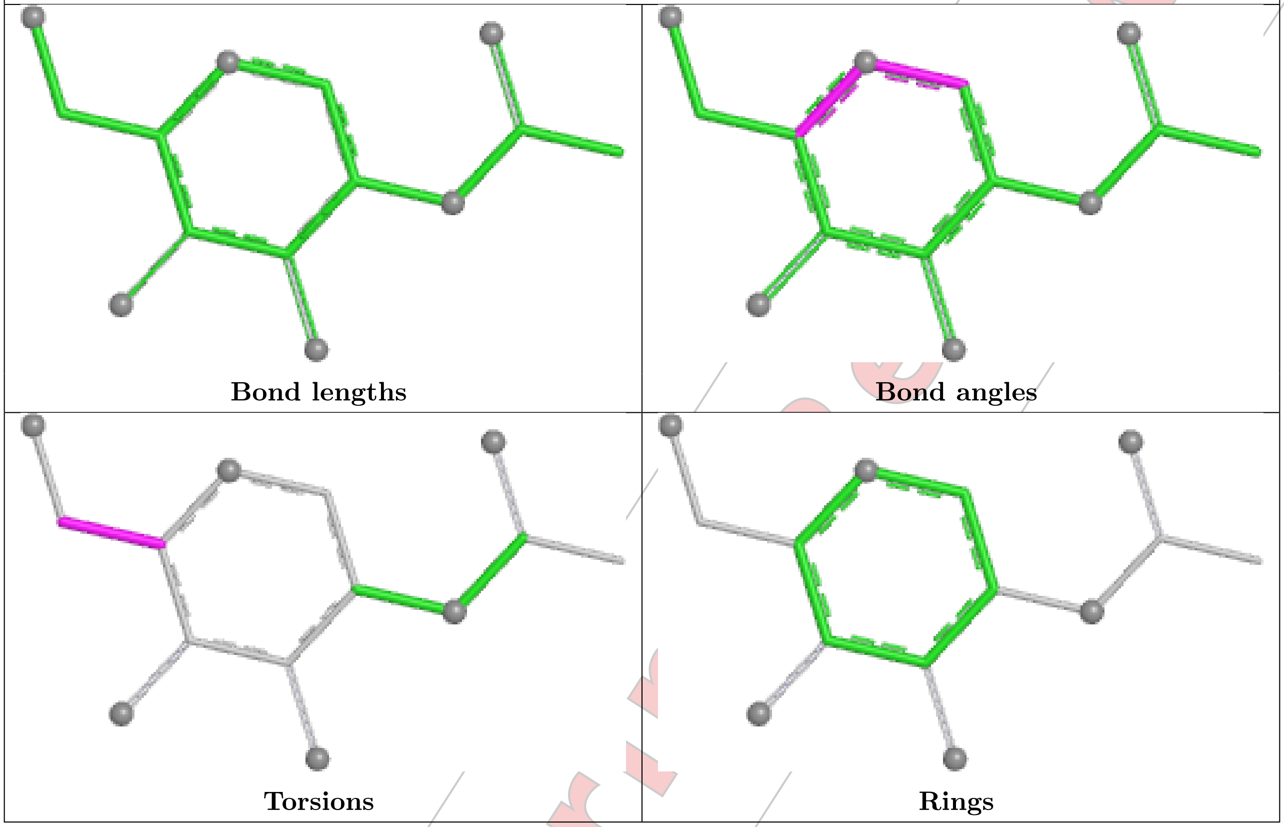


Ligand NAG D 301

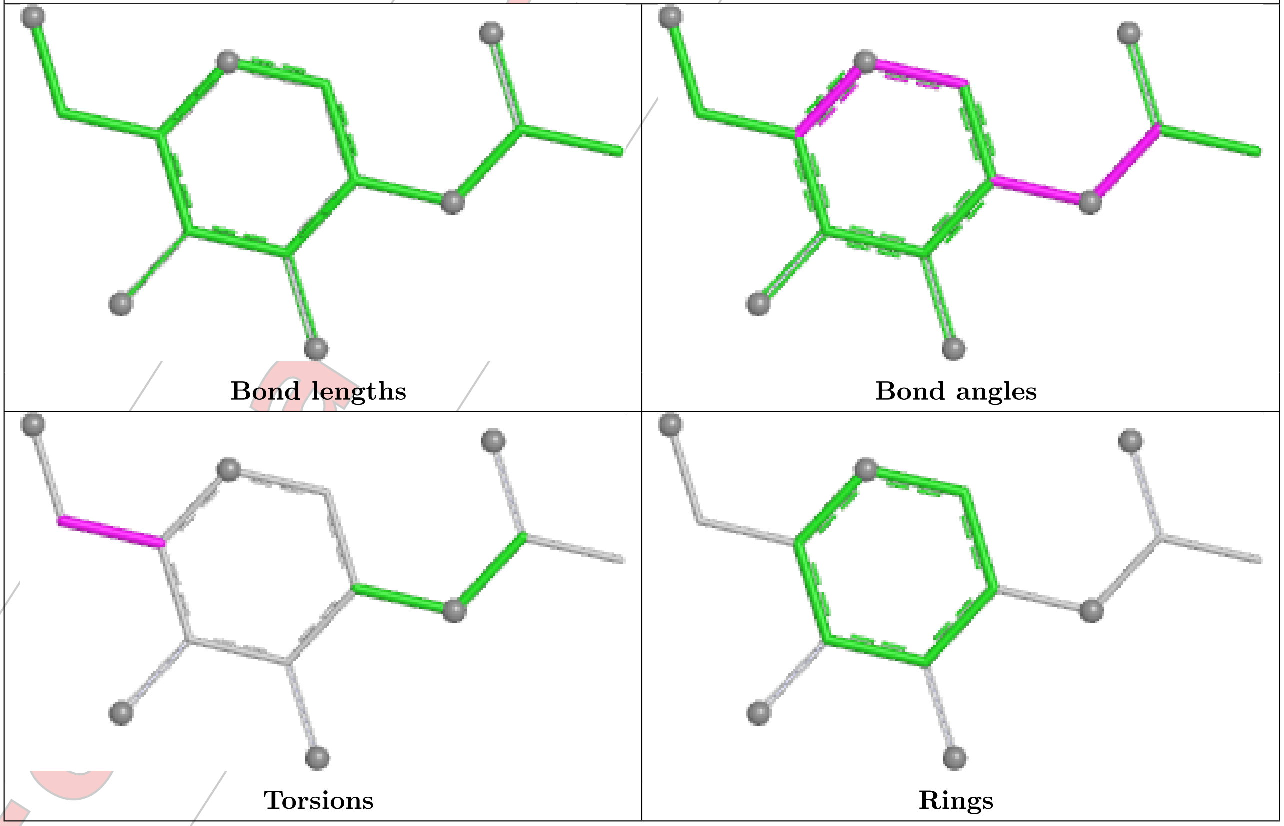

#### Ligand NAG D 302

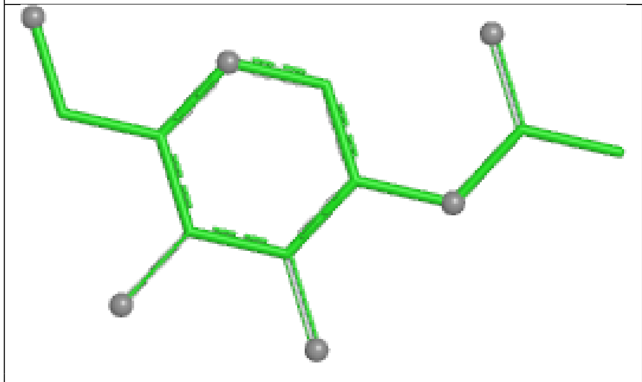

Bond lengths

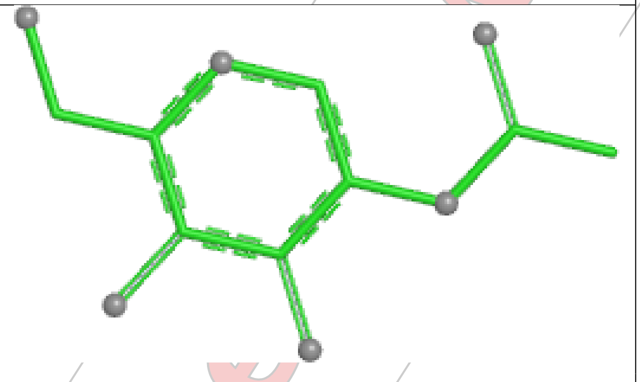

Bond angles

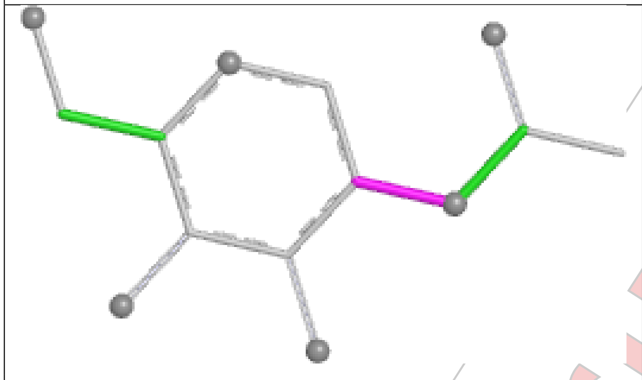

Torsions

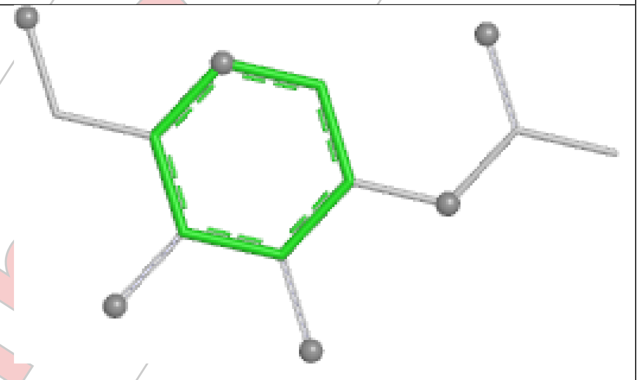

Rings

#### Ligand NAG A 301

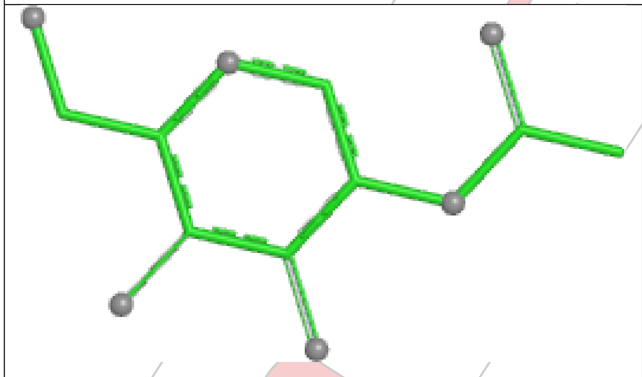

Bond lengths

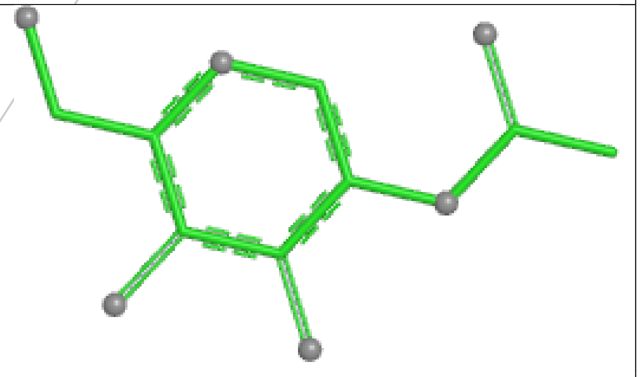

Bond angles

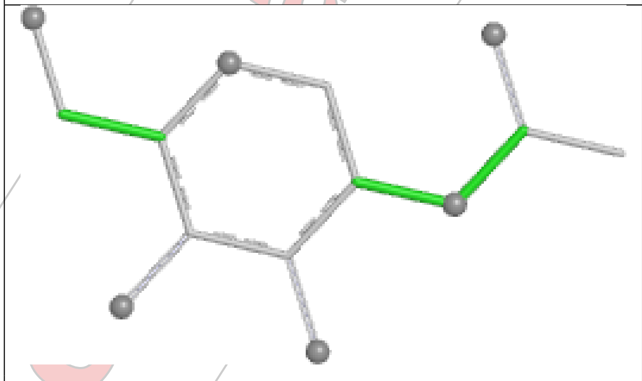

Torsions

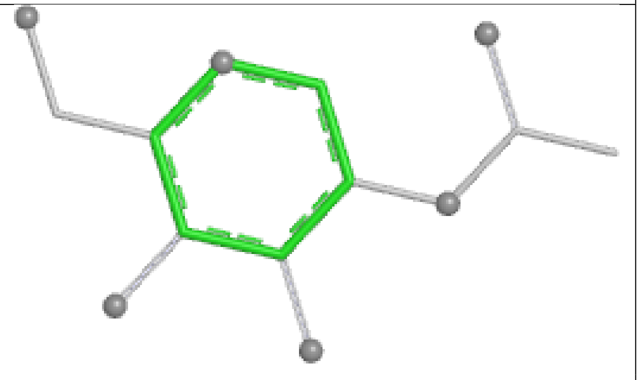

Rings

#### Ligand NAG B 301

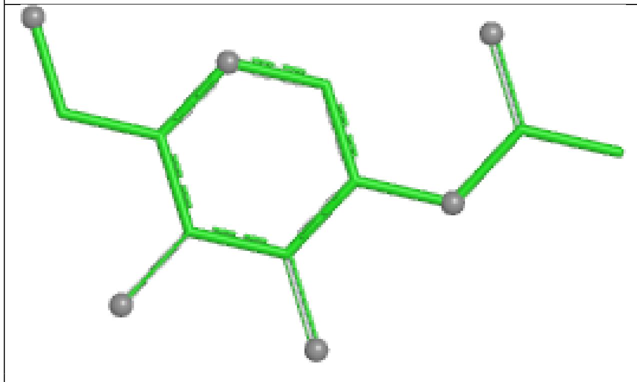

Bond lengths

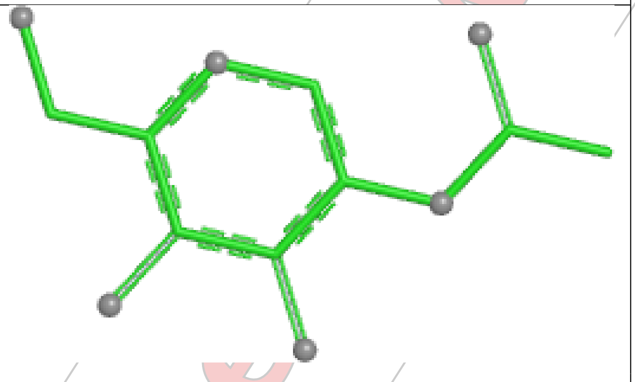

Bond angles

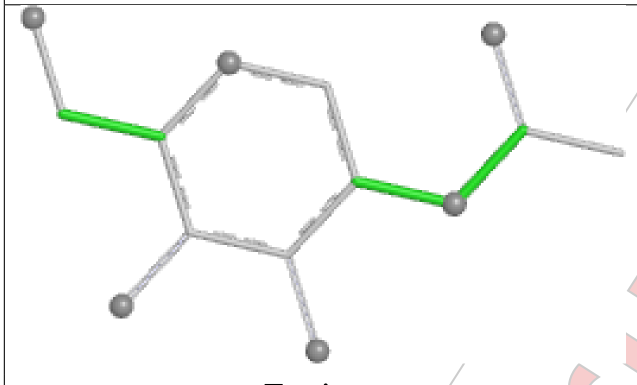

Torsions

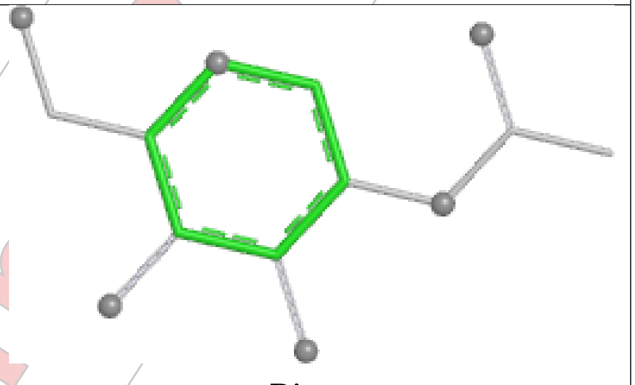

Rings

#### Ligand NAG C 301

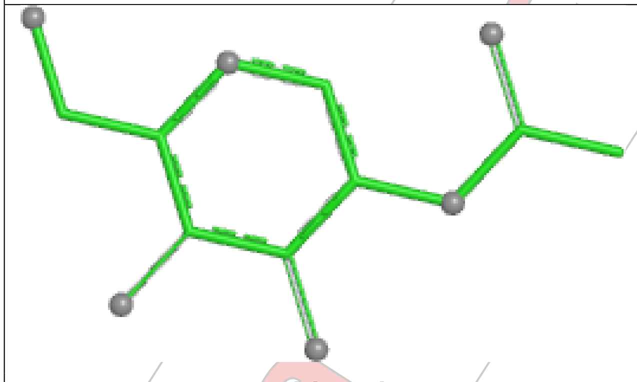

Bond lengths

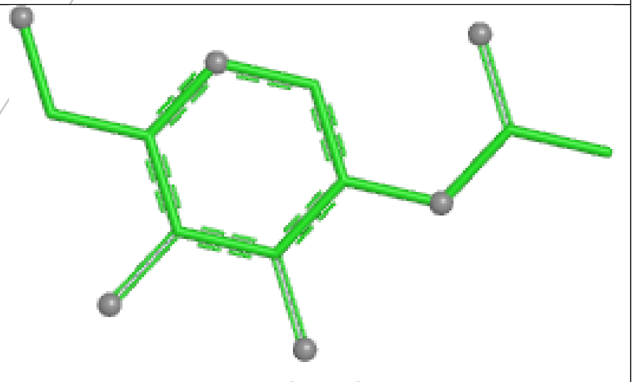

Bond angles

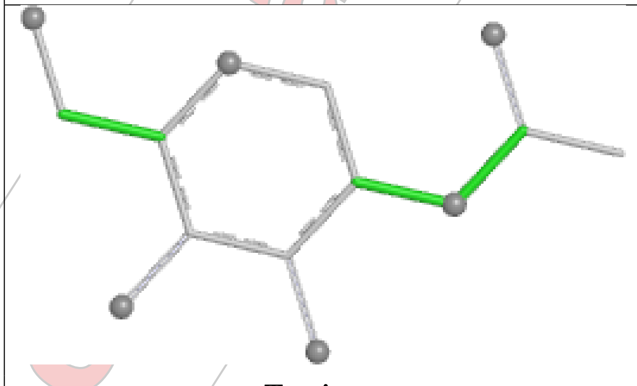

Torsions

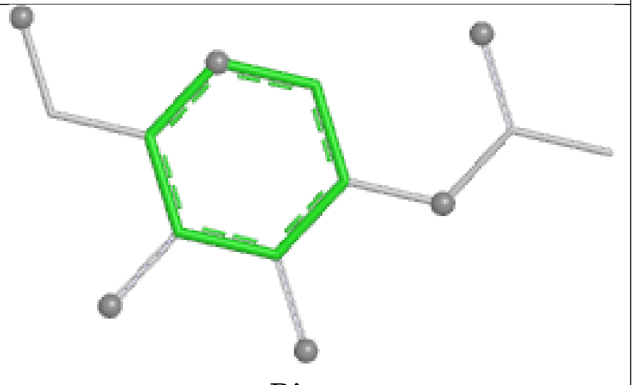

Rings

#### Ligand NAG A 302

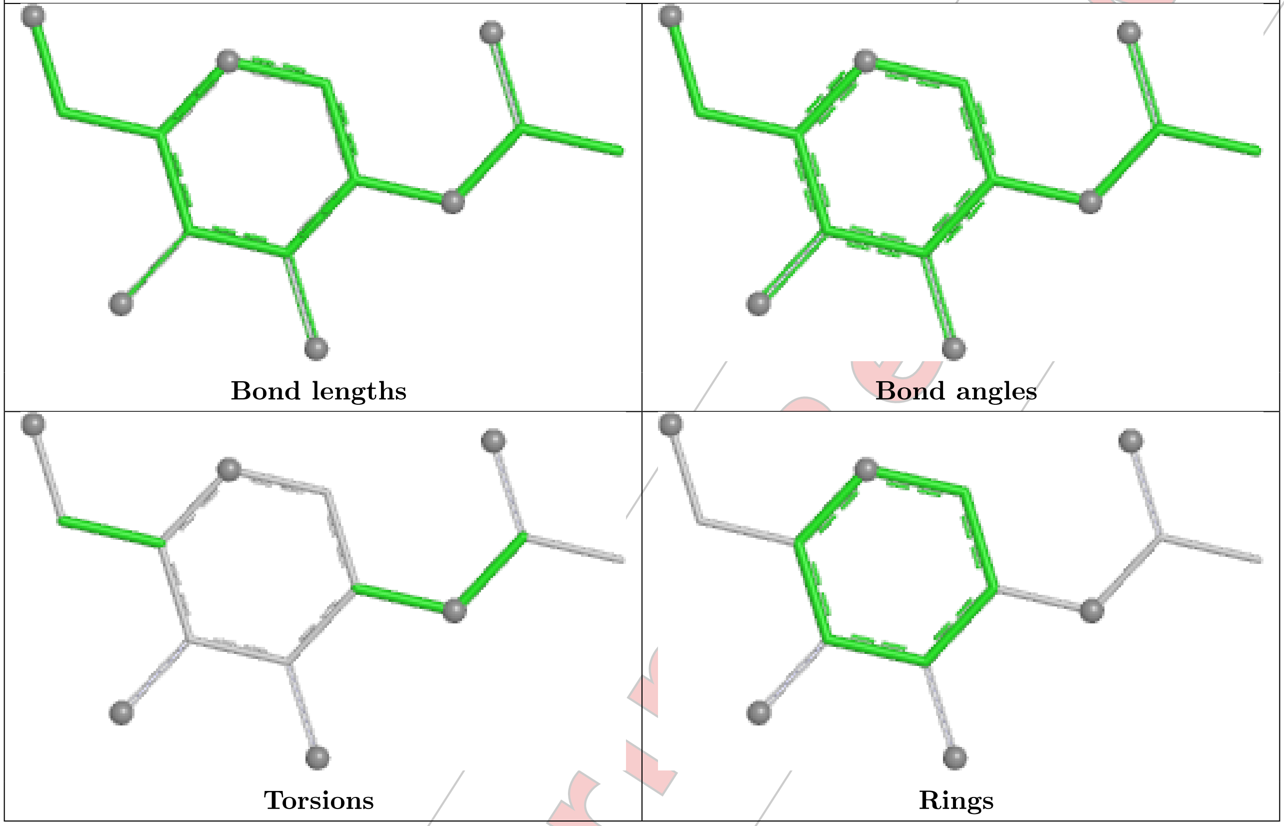


#### Ligand NAG B 304

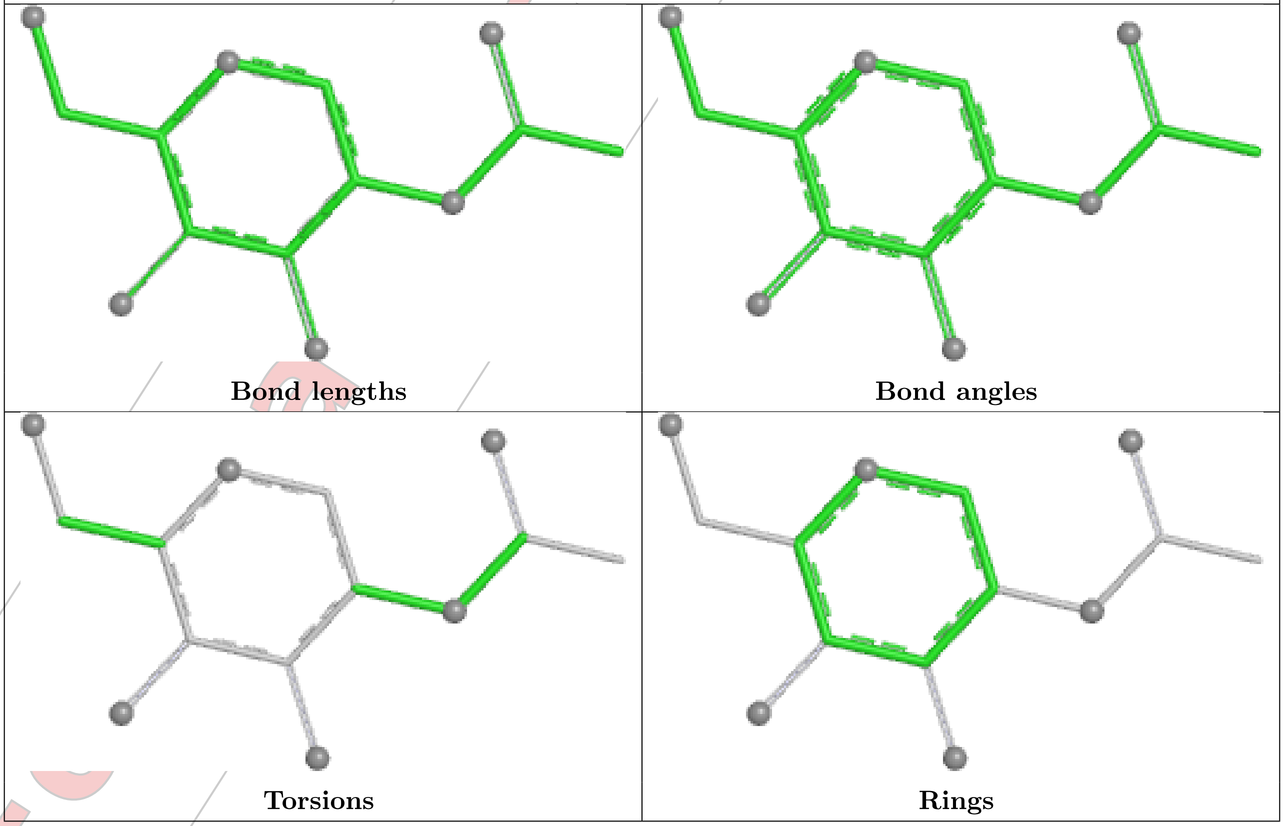

Ligand NAG A 303

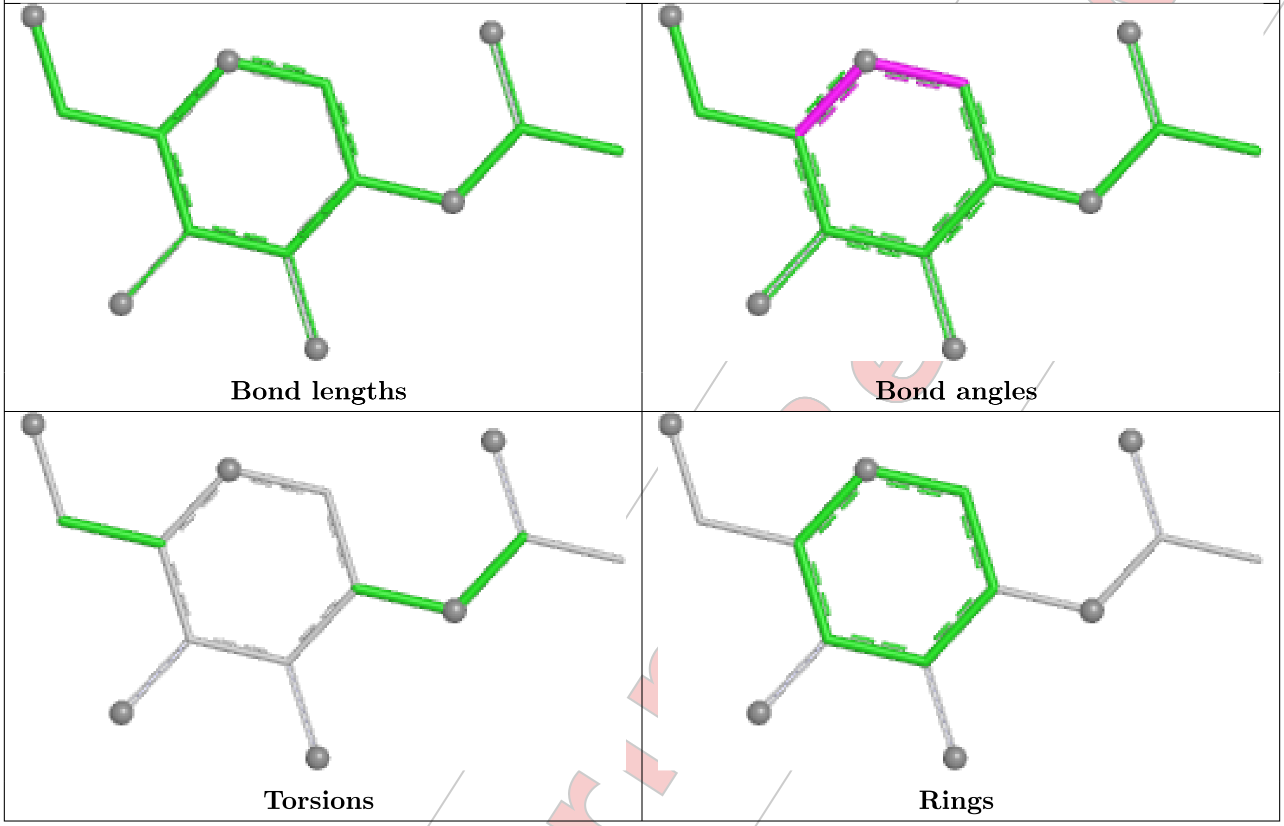

| Bond lengths | Bond angles |
| --- | --- |
| Torsions | Rings |

#### 5.7 Other polymers (i)

There are no such residues in this entry.

#### 5.8 Polymer linkage issues (i)

There are no chain breaks in this entry.

For Manuscript Review

### 6 Fit of model and data (i)

#### 6.1 Protein, DNA and RNA chains (i)

In the following table, the column labelled '#RSRZ> 2' contains the number (and percentage) of RSRZ outliers, followed by percent RSRZ outliers for the chain as percentile scores relative to all X-ray entries and entries of similar resolution. The OWAB column contains the minimum, median, $95^{th}$ percentile and maximum values of the occupancy-weighted average B-factor per residue. The column labelled 'Q< 0.9' lists the number of (and percentage) of residues with an average occupancy less than 0.9.

| Mol | Chain | Analysed | <RSRZ> | #RSRZ>2 | | | OWAB(Å$^2$) | Q<0.9 |
|---|---|---|---|---|---|---|---|---|
| 1 | A | 208/219 (94%) | 0.29 | 6 (2%) | 54 | 50 | 39, 59, 118, 144 | 0 |
| 1 | C | 194/219 (88%) | 0.85 | 19 (9%) | 14 | 13 | 47, 90, 141, 166 | 0 |
| 2 | B | 239/250 (95%) | -0.04 | 1 (0%) | 89 | 86 | 36, 52, 84, 150 | 0 |
| 2 | D | 239/250 (95%) | 0.67 | 23 (9%) | 15 | 14 | 43, 83, 138, 156 | 0 |
| All | All | 880/938 (93%) | 0.43 | 49 (5%) | 31 | 29 | 36, 68, 131, 166 | 0 |

All (49) RSRZ outliers are listed below:

| Mol | Chain | Res | Type | RSRZ |
|---|---|---|---|---|
| 1 | A | 2 | GLN | 4.5 |
| 1 | A | 104 | TYR | 4.3 |
| 1 | C | 104 | TYR | 3.9 |
| 2 | D | 92 | SER | 3.7 |
| 2 | D | 33 | TRP | 3.6 |
| 2 | D | 74 | LEU | 3.3 |
| 1 | A | 189 | PHE | 3.2 |
| 1 | C | 189 | PHE | 3.0 |
| 2 | D | 26 | LEU | 2.9 |
| 1 | C | 84 | ILE | 2.8 |
| 2 | D | 91 | ALA | 2.8 |
| 1 | C | 25 | TYR | 2.8 |
| 1 | C | 186 | GLN | 2.8 |
| 1 | C | 161 | PHE | 2.7 |
| 1 | C | 32 | TYR | 2.7 |
| 2 | D | 48 | TYR | 2.7 |
| 1 | C | 50 | ARG | 2.7 |
| 2 | D | 20 | LEU | 2.7 |
| 2 | D | 212 | LEU | 2.6 |
| 1 | A | 156 | MET | 2.6 |
| 1 | C | 147 | ASP | 2.6 |

*Continued on next page...*

For Manuscript Review

*Continued from previous page...*

| Mol | Chain | Res | Type | RSRZ |
|---|---|---|---|---|
| 2 | D | 28 | HIS | 2.5 |
| 1 | C | 155 | THR | 2.5 |
| 1 | C | 2 | GLN | 2.4 |
| 1 | C | 34 | LEU | 2.3 |
| 1 | C | 187 | THR | 2.3 |
| 2 | D | 58 | THR | 2.3 |
| 1 | C | 197 | GLU | 2.3 |
| 2 | D | 113 | LEU | 2.3 |
| 2 | D | 38 | ALA | 2.3 |
| 1 | C | 103 | ASN | 2.3 |
| 1 | A | 199 | ASN | 2.2 |
| 1 | C | 136 | GLN | 2.2 |
| 2 | D | 219 | PRO | 2.2 |
| 1 | C | 49 | ILE | 2.2 |
| 2 | D | 31 | MET | 2.2 |
| 2 | D | 98 | GLY | 2.2 |
| 2 | B | 220 | GLU | 2.2 |
| 2 | D | 69 | ASP | 2.2 |
| 2 | D | 100 | TYR | 2.1 |
| 1 | A | 155 | THR | 2.1 |
| 1 | C | 137 | ASP | 2.1 |
| 2 | D | 75 | LEU | 2.1 |
| 2 | D | 139 | LYS | 2.1 |
| 2 | D | 80 | VAL | 2.1 |
| 2 | D | 131 | LYS | 2.0 |
| 2 | D | 213 | SER | 2.0 |
| 2 | D | 241 | SER | 2.0 |
| 1 | C | 68 | PHE | 2.0 |

#### 6.2 Non-standard residues in protein, DNA, RNA chains (i)

There are no non-standard protein/DNA/RNA residues in this entry.

#### 6.3 Carbohydrates (i)

There are no monosaccharides in this entry.

#### 6.4 Ligands (i)

In the following table, the Atoms column lists the number of modelled atoms in the group and the number defined in the chemical component dictionary. The B-factors column lists the minimum,

For Manuscript Review

median, 95$^{th}$ percentile and maximum values of B factors of atoms in the group. The column labelled 'Q< 0.9' lists the number of atoms with occupancy less than 0.9.

| Mol | Type | Chain | Res | Atoms | RSCC | RSR | B-factors(Å$^2$) | Q<0.9 |
|---|---|---|---|---|---|---|---|---|
| 3 | NAG | D | 301 | 14/15 | -0.12 | 0.17 | 114,125,133,133 | 0 |
| 3 | NAG | D | 302 | 14/15 | 0.16 | 0.19 | 115,124,128,130 | 0 |
| 3 | NAG | C | 301 | 14/15 | 0.27 | 0.15 | 128,135,139,139 | 0 |
| 3 | NAG | A | 302 | 14/15 | 0.38 | 0.18 | 103,119,122,125 | 0 |
| 3 | NAG | B | 303 | 14/15 | 0.46 | 0.17 | 94,98,101,105 | 0 |
| 3 | NAG | D | 303 | 14/15 | 0.54 | 0.17 | 67,83,89,91 | 0 |
| 4 | GOL | C | 302 | 6/6 | 0.69 | 0.20 | 76,80,81,84 | 0 |
| 3 | NAG | A | 303 | 14/15 | 0.74 | 0.12 | 87,104,111,115 | 0 |
| 3 | NAG | B | 301 | 14/15 | 0.78 | 0.10 | 82,86,90,91 | 0 |
| 4 | GOL | A | 304 | 6/6 | 0.82 | 0.19 | 47,47,52,52 | 6 |
| 4 | GOL | B | 307 | 6/6 | 0.83 | 0.13 | 51,54,59,62 | 6 |
| 4 | GOL | B | 306 | 6/6 | 0.83 | 0.14 | 55,56,57,61 | 6 |
| 4 | GOL | B | 305 | 6/6 | 0.84 | 0.12 | 44,47,49,50 | 6 |
| 5 | CL | D | 305 | 1/1 | 0.84 | 0.09 | 101,101,101,101 | 0 |
| 3 | NAG | B | 304 | 14/15 | 0.87 | 0.10 | 52,58,67,74 | 0 |
| 5 | CL | D | 306 | 1/1 | 0.90 | 0.13 | 85,85,85,85 | 0 |
| 5 | CL | B | 312 | 1/1 | 0.91 | 0.12 | 77,77,77,77 | 0 |
| 3 | NAG | A | 301 | 14/15 | 0.92 | 0.10 | 64,68,71,83 | 0 |
| 3 | NAG | B | 302 | 14/15 | 0.92 | 0.10 | 33,40,48,51 | 0 |
| 5 | CL | B | 309 | 1/1 | 0.93 | 0.10 | 65,65,65,65 | 0 |
| 5 | CL | B | 308 | 1/1 | 0.93 | 0.14 | 62,62,62,62 | 0 |
| 5 | CL | B | 310 | 1/1 | 0.94 | 0.06 | 73,73,73,73 | 0 |
| 5 | CL | B | 311 | 1/1 | 0.96 | 0.06 | 68,68,68,68 | 0 |
| 5 | CL | D | 304 | 1/1 | 0.96 | 0.07 | 60,60,60,60 | 0 |

The following is a graphical depiction of the model fit to experimental electron density of all instances of the Ligand of Interest. In addition, ligands with molecular weight > 250 and outliers as shown on the geometry validation Tables will also be included. Each fit is shown from different orientation to approximate a three-dimensional view.

For Manuscript Review

**Electron density around NAG D 301:**

$2mF_o$-$DF_c$ (at 0.7 rmsd) in gray
$mF_o$-$DF_c$ (at 3 rmsd) in purple (negative)
and green (positive)

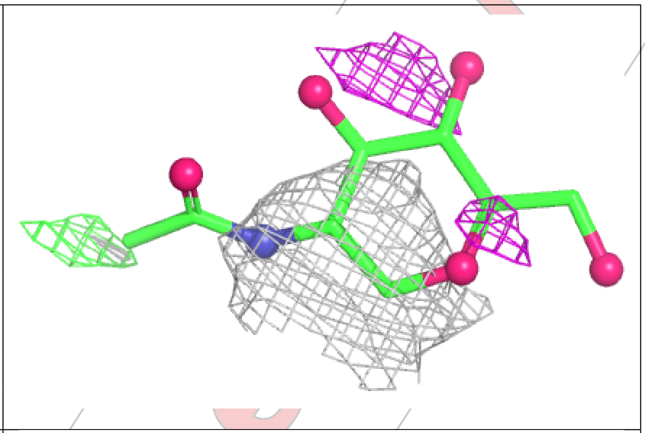

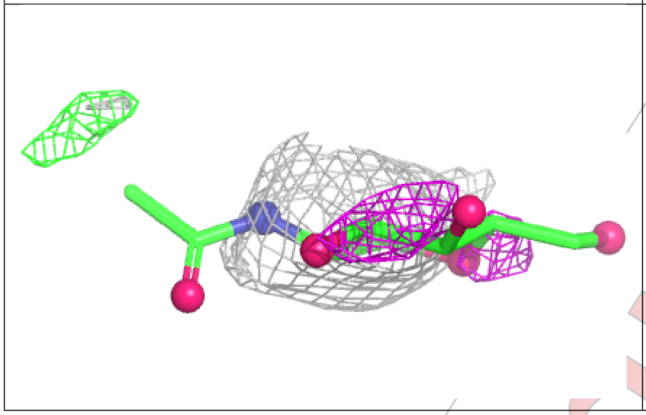

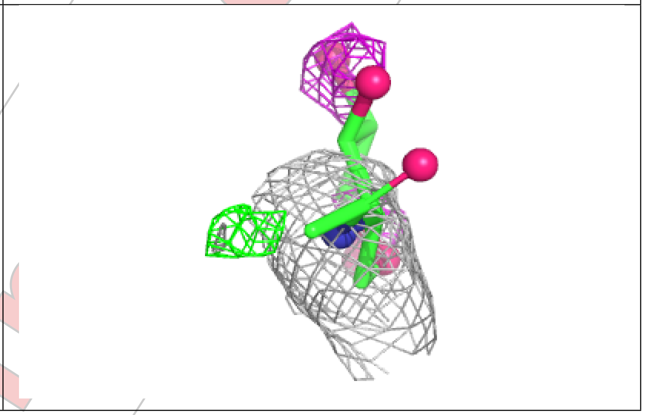

**Electron density around NAG D 302:**

$2mF_o$-$DF_c$ (at 0.7 rmsd) in gray
$mF_o$-$DF_c$ (at 3 rmsd) in purple (negative)
and green (positive)

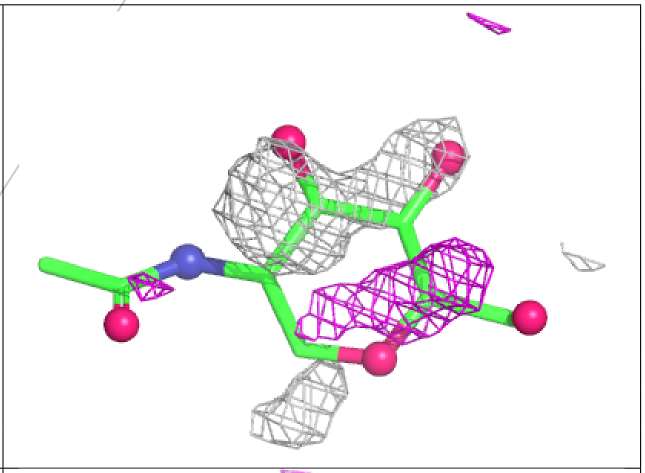

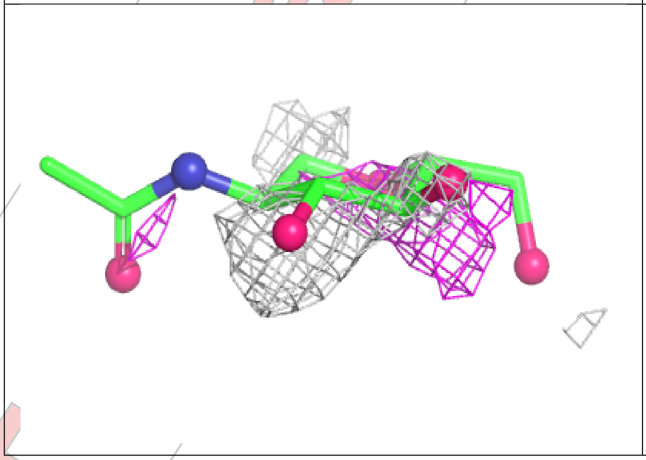

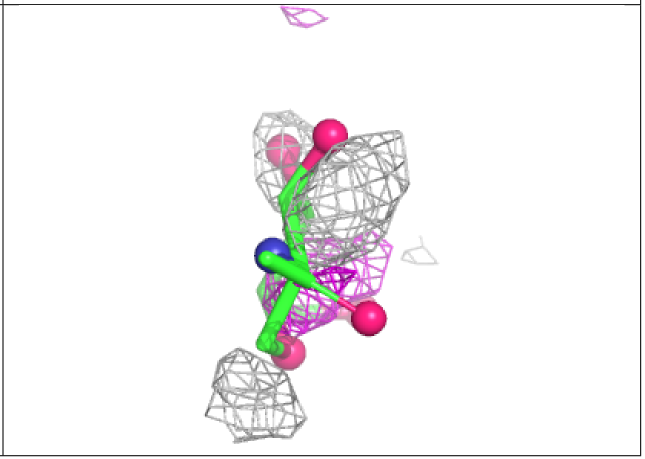

**Electron density around NAG C 301:**

$2mF_o\text{-}DF_c$ (at 0.7 rmsd) in gray
$mF_o\text{-}DF_c$ (at 3 rmsd) in purple (negative)
and green (positive)

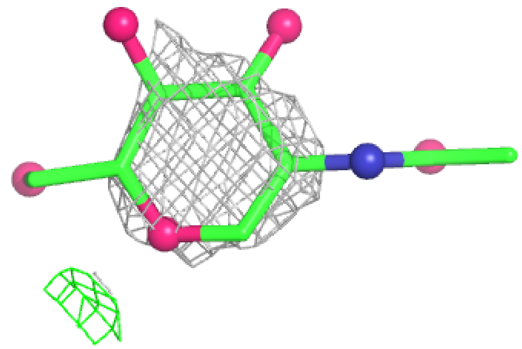

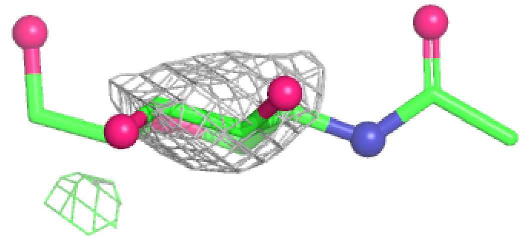

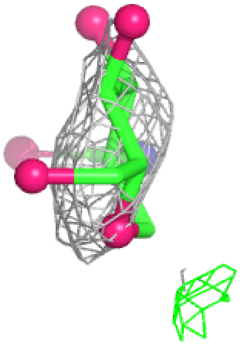

**Electron density around NAG A 302:**

$2mF_o\text{-}DF_c$ (at 0.7 rmsd) in gray
$mF_o\text{-}DF_c$ (at 3 rmsd) in purple (negative)
and green (positive)

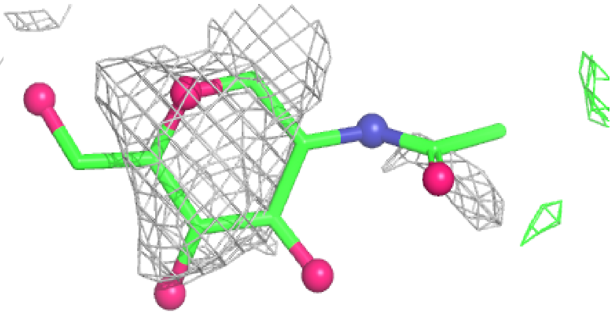

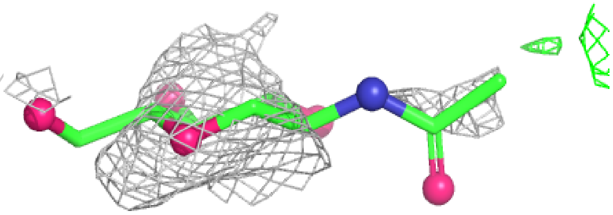

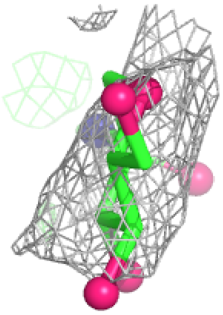

**Electron density around NAG B 303:**

2mF$_o$-DF$_c$ (at 0.7 rmsd) in gray
mF$_o$-DF$_c$ (at 3 rmsd) in purple (negative)
and green (positive)

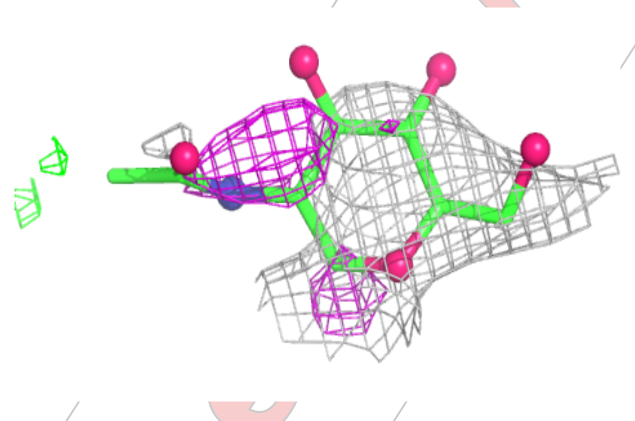

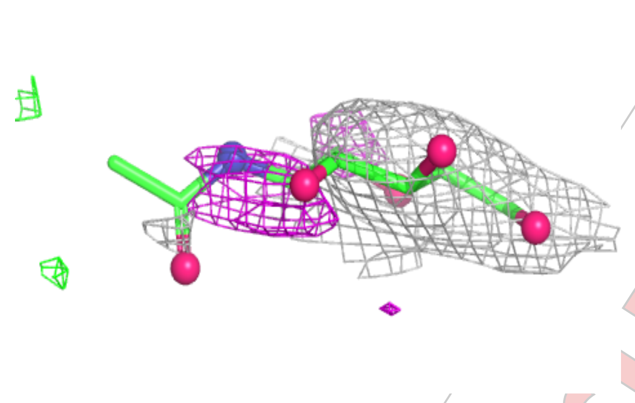

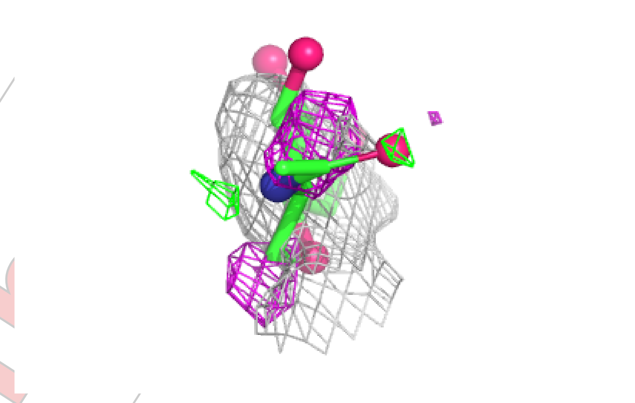

**Electron density around NAG D 303:**

2mF$_o$-DF$_c$ (at 0.7 rmsd) in gray
mF$_o$-DF$_c$ (at 3 rmsd) in purple (negative)
and green (positive)

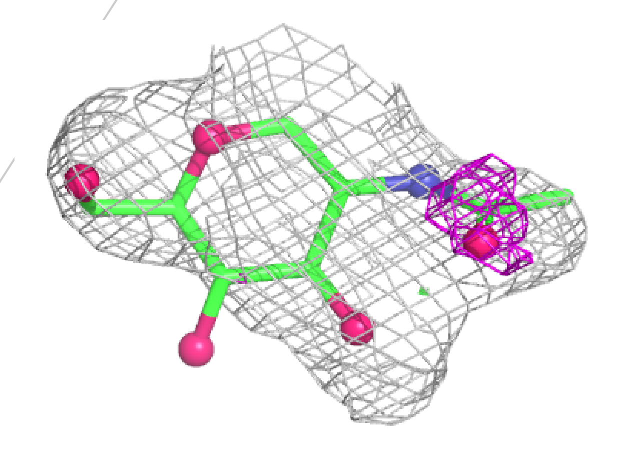

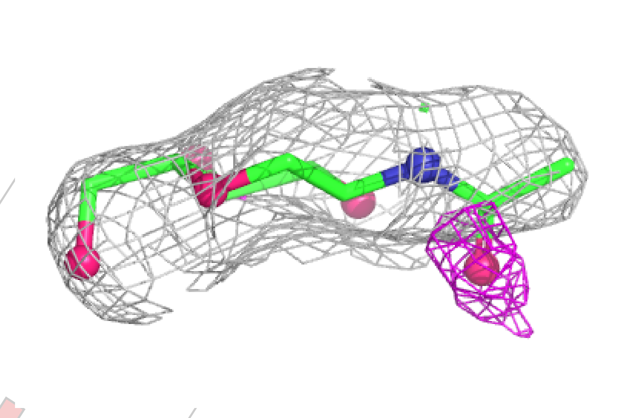

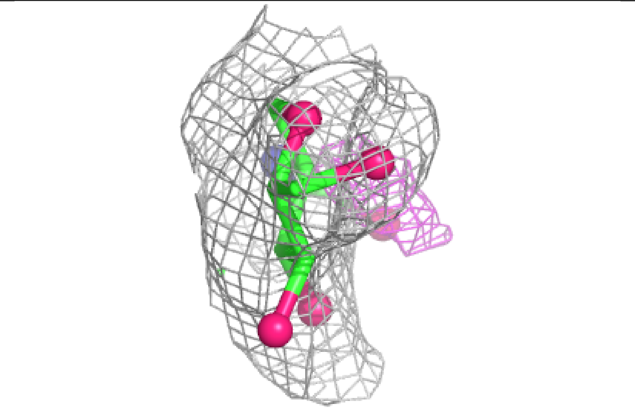

**Electron density around NAG A 303:**

$2mF_o$-$DF_c$ (at 0.7 rmsd) in gray
$mF_o$-$DF_c$ (at 3 rmsd) in purple (negative)
and green (positive)

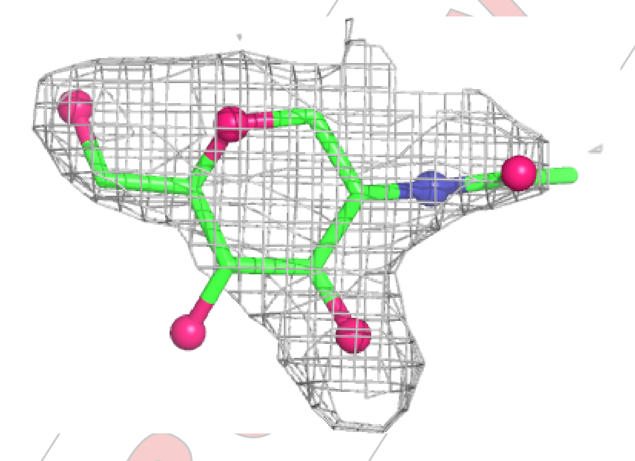

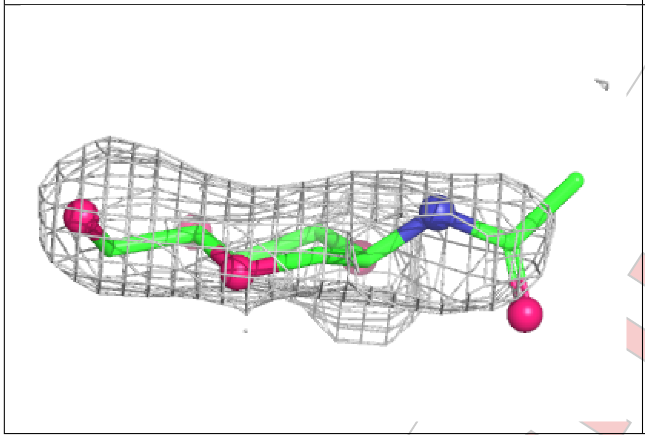

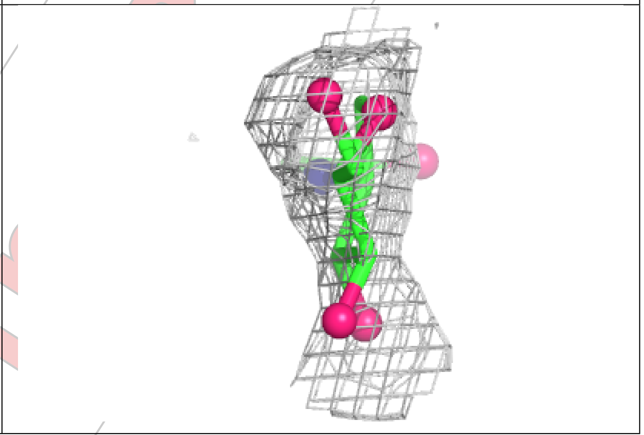

**Electron density around NAG B 301:**

$2mF_o$-$DF_c$ (at 0.7 rmsd) in gray
$mF_o$-$DF_c$ (at 3 rmsd) in purple (negative)
and green (positive)

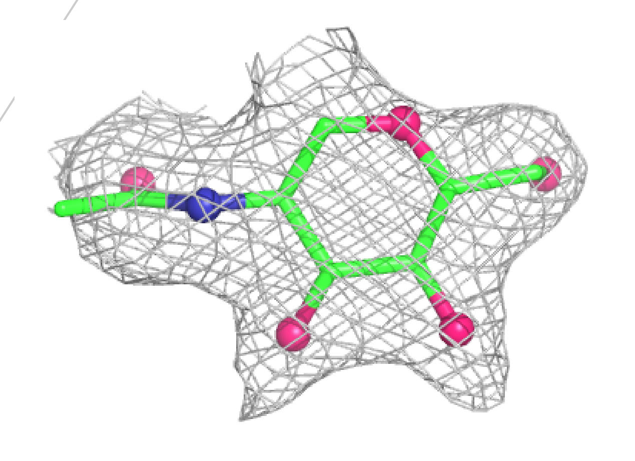

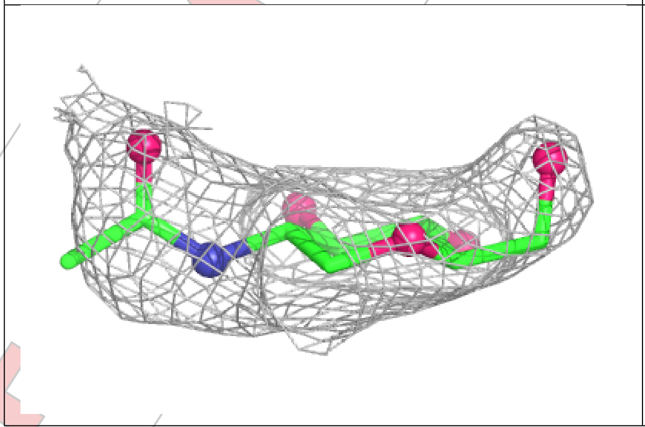

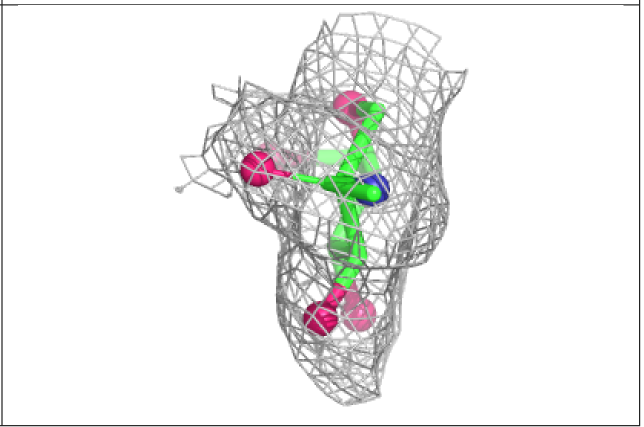

**Electron density around NAG B 304:**

$2mF_o$-$DF_c$ (at 0.7 rmsd) in gray
$mF_o$-$DF_c$ (at 3 rmsd) in purple (negative)
and green (positive)

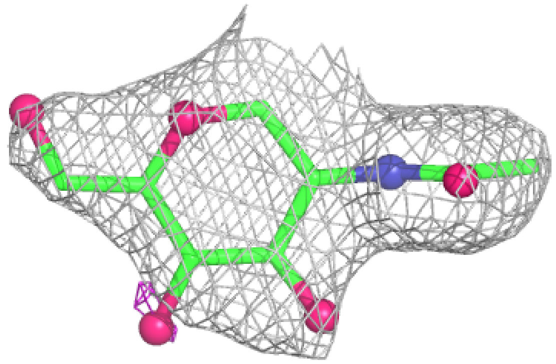

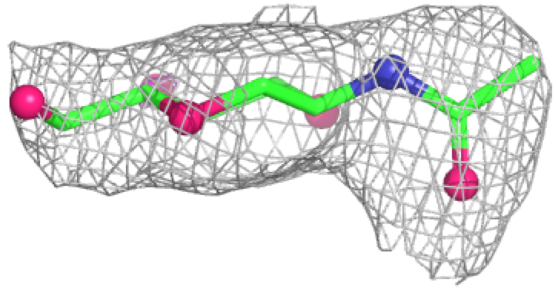

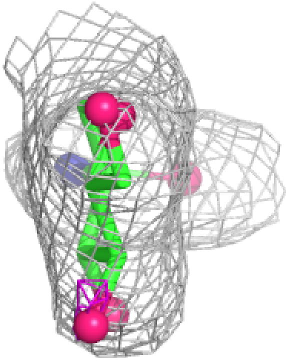

**Electron density around NAG A 301:**

$2mF_o$-$DF_c$ (at 0.7 rmsd) in gray
$mF_o$-$DF_c$ (at 3 rmsd) in purple (negative)
and green (positive)

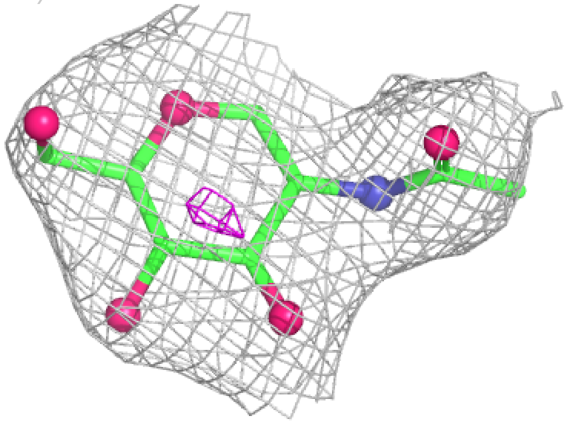

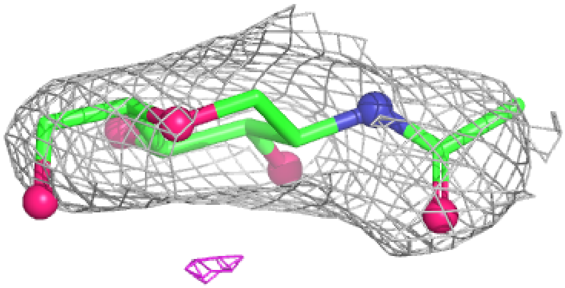

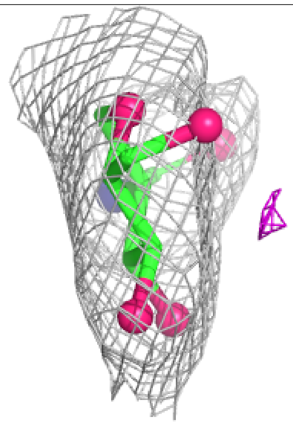

For Manuscript Review

| | |
|---|---|
| **Electron density around NAG B 302:**<br><br>2mF$_o$-DF$_c$ (at 0.7 rmsd) in gray<br>mF$_o$-DF$_c$ (at 3 rmsd) in purple (negative)<br>and green (positive) | 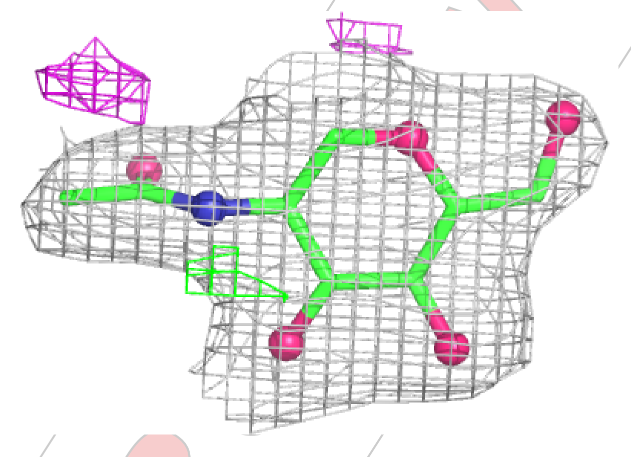 |
| 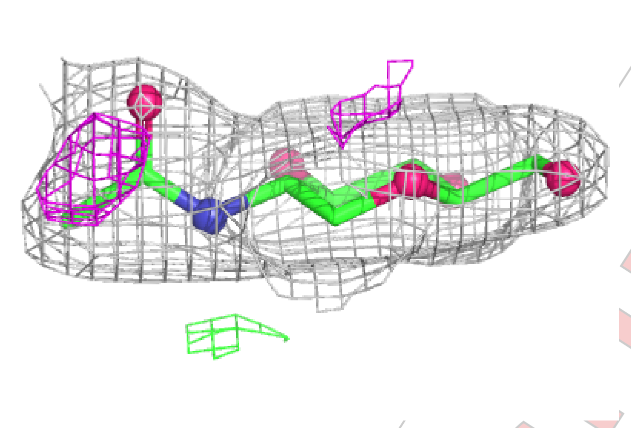 | 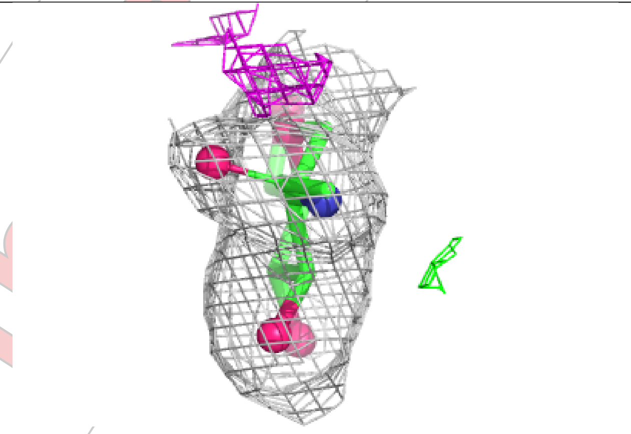 |

#### 6.5 Other polymers (i)

There are no such residues in this entry.

For Manuscript Review

### Full wwPDB X-ray Structure Validation Report (i)

Mar 21, 2025 – 02:09 PM EDT

| | | |
|---|---|---|
| PDB ID | : | 9NV2 |
| Title | : | Crystal structure of NP207 TCR in complex with NP366-H-2Db |
| Deposited on | : | 2025-03-20 |
| Resolution | : | 1.95 Å(reported) |

**This wwPDB validation report is for manuscript review**

This is a Full wwPDB X-ray Structure Validation Report.

This report is produced by the wwPDB biocuration pipeline after annotation of the structure.

We welcome your comments at **
A user guide is available at
https://www.wwpdb.org/validation/2017/XrayValidationReportHelp
with specific help available everywhere you see the (i) symbol.

The types of validation reports are described at
http://www.wwpdb.org/validation/2017/FAQs#types.

The following versions of software and data (see references (i)) were used in the production of this report:

| | | |
|---|---|---|
| MolProbity | : | 4.02b-467 |
| Mogul | : | 2022.3.0, CSD as543be (2022) |
| Xtriage (Phenix) | : | 1.21 |
| EDS | : | 3.0 |
| buster-report | : | 1.1.7 (2018) |
| Percentile statistics | : | 20231227.v01 (using entries in the PDB archive December 27th 2023) |
| CCP4 | : | 9.0.004 (Gargrove) |
| Density-Fitness | : | 1.0.11 |
| Ideal geometry (proteins) | : | Engh & Huber (2001) |
| Ideal geometry (DNA, RNA) | : | Parkinson et al. (1996) |

For Manuscript Review

### 1 Overall quality at a glance (i)

The following experimental techniques were used to determine the structure:
*X-RAY DIFFRACTION*

The reported resolution of this entry is 1.95 Å.

Percentile scores (ranging between 0-100) for global validation metrics of the entry are shown in the following graphic. The table shows the number of entries on which the scores are based.

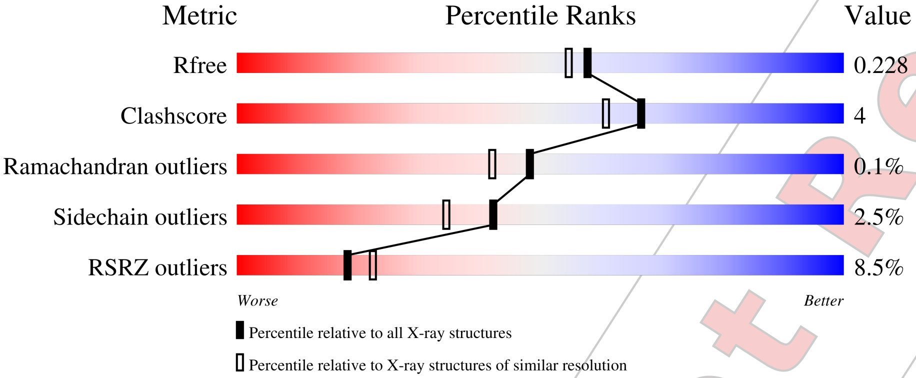


| Metric | Whole archive (#Entries) | Similar resolution (#Entries, resolution range(Å)) |
|---|---|---|
| $R_{free}$ | 164625 | 3187 (1.96-1.96) |
| Clashscore | 180529 | 3412 (1.96-1.96) |
| Ramachandran outliers | 177936 | 3390 (1.96-1.96) |
| Sidechain outliers | 177891 | 3390 (1.96-1.96) |
| RSRZ outliers | 164620 | 3186 (1.96-1.96) |

The table below summarises the geometric issues observed across the polymeric chains and their fit to the electron density. The red, orange, yellow and green segments of the lower bar indicate the fraction of residues that contain outliers for >=3, 2, 1 and 0 types of geometric quality criteria respectively. A grey segment represents the fraction of residues that are not modelled. The numeric value for each fraction is indicated below the corresponding segment, with a dot representing fractions <=5% The upper red bar (where present) indicates the fraction of residues that have poor fit to the electron density. The numeric value is given above the bar.

| Mol | Chain | Length | Quality of chain |
|---|---|---|---|
| 1 | A | 223 | 5% / 80% 11% 8% |
| 2 | B | 254 | 2% / 85% 10% • |
| 3 | C | 281 | 19% / 85% 12% • |
| 4 | D | 100 | 87% 11% • |

*Continued on next page...*



For Manuscript Review

*Continued from previous page...*

| Mol | Chain | Length | Quality of chain |
|---|---|---|---|
| 5 | P | 9 | 100% |
| 6 | E | 3 | 100% |

For Manuscript Review

#### 2 Entry composition (i)

There are 10 unique types of molecules in this entry. The entry contains 7027 atoms, of which 0 are hydrogens and 0 are deuteriums.

In the tables below, the ZeroOcc column contains the number of atoms modelled with zero occupancy, the AltConf column contains the number of residues with at least one atom in alternate conformation and the Trace column contains the number of residues modelled with at most 2 atoms.

- Molecule 1 is a protein called NP207 TCR alpha-chain.

| Mol | Chain | Residues | Atoms | | | | | ZeroOcc | AltConf | Trace |
|---|---|---|---|---|---|---|---|---|---|---|
| | | | Total | C | N | O | S | | | |
| 1 | A | 205 | 1599 | 1008 | 258 | 323 | 10 | 0 | 1 | 0 |

- Molecule 2 is a protein called NP207 TCR beta-chain.

| Mol | Chain | Residues | Atoms | | | | | ZeroOcc | AltConf | Trace |
|---|---|---|---|---|---|---|---|---|---|---|
| | | | Total | C | N | O | S | | | |
| 2 | B | 243 | 1959 | 1243 | 339 | 369 | 8 | 0 | 1 | 0 |

- Molecule 3 is a protein called H-2 class I histocompatibility antigen, D-B alpha chain.

| Mol | Chain | Residues | Atoms | | | | | ZeroOcc | AltConf | Trace |
|---|---|---|---|---|---|---|---|---|---|---|
| | | | Total | C | N | O | S | | | |
| 3 | C | 271 | 2198 | 1386 | 396 | 408 | 8 | 0 | 4 | 0 |

There is a discrepancy between the modelled and reference sequences:

| Chain | Residue | Modelled | Actual | Comment | Reference |
|---|---|---|---|---|---|
| C | 0 | MET | - | initiating methionine | UNP P01899 |

- Molecule 4 is a protein called Beta-2-microglobulin.

| Mol | Chain | Residues | Atoms | | | | | ZeroOcc | AltConf | Trace |
|---|---|---|---|---|---|---|---|---|---|---|
| | | | Total | C | N | O | S | | | |
| 4 | D | 98 | 813 | 518 | 137 | 151 | 7 | 0 | 0 | 0 |

There are 2 discrepancies between the modelled and reference sequences:

| Chain | Residue | Modelled | Actual | Comment | Reference |
|---|---|---|---|---|---|
| D | 0 | MET | - | initiating methionine | UNP P01887 |
| D | 85 | ASP | ALA | conflict | UNP P01887 |

For Manuscript Review

- Molecule 5 is a protein called Nucleoprotein peptide.

| Mol | Chain | Residues | Atoms | | | | | ZeroOcc | AltConf | Trace |
|---|---|---|---|---|---|---|---|---|---|---|
| | | | Total | C | N | O | S | | | |
| 5 | P | 9 | 69 | 38 | 11 | 18 | 2 | 0 | 0 | 0 |

- Molecule 6 is an oligosaccharide called beta-D-mannopyranose-(1-4)-2-acetamido-2-deoxy-beta-D-glucopyranose-(1-4)-2-acetamido-2-deoxy-beta-D-glucopyranose.

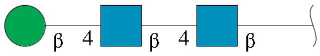


| Mol | Chain | Residues | Atoms | | | | ZeroOcc | AltConf | Trace |
|---|---|---|---|---|---|---|---|---|---|
| | | | Total | C | N | O | | | |
| 6 | E | 3 | 39 | 22 | 2 | 15 | 0 | 0 | 0 |

- Molecule 7 is 2-acetamido-2-deoxy-beta-D-glucopyranose (three-letter code: NAG) (formula: $C_8H_{15}NO_6$) (labeled as "Ligand of Interest" by depositor).

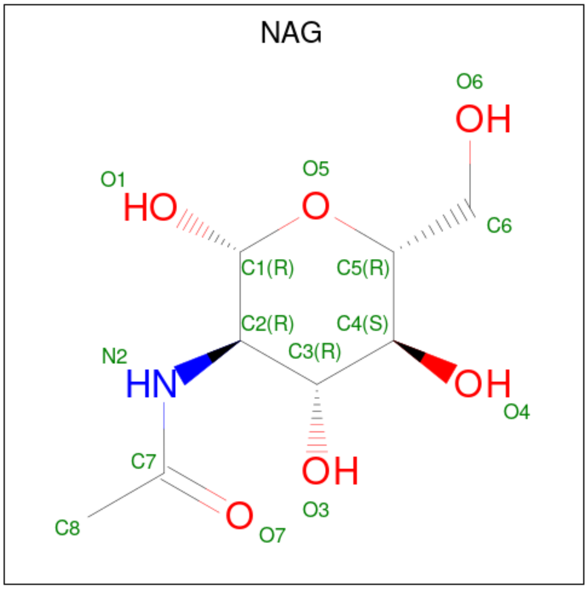


| Mol | Chain | Residues | Atoms | | | | ZeroOcc | AltConf |
|---|---|---|---|---|---|---|---|---|
| | | | Total | C | N | O | | |
| 7 | A | 1 | 14 | 8 | 1 | 5 | 0 | 0 |
| 7 | A | 1 | 14 | 8 | 1 | 5 | 0 | 0 |
| 7 | A | 1 | 14 | 8 | 1 | 5 | 0 | 0 |

*Continued on next page...*

For Manuscript Review

*Continued from previous page...*

| Mol | Chain | Residues | Atoms | | | | ZeroOcc | AltConf |
|---|---|---|---|---|---|---|---|---|
| | | | Total | C | N | O | | |
| 7 | B | 1 | 14 | 8 | 1 | 5 | 0 | 0 |

- Molecule 8 is 1,2-ETHANEDIOL (three-letter code: EDO) (formula: $C_2H_6O_2$).

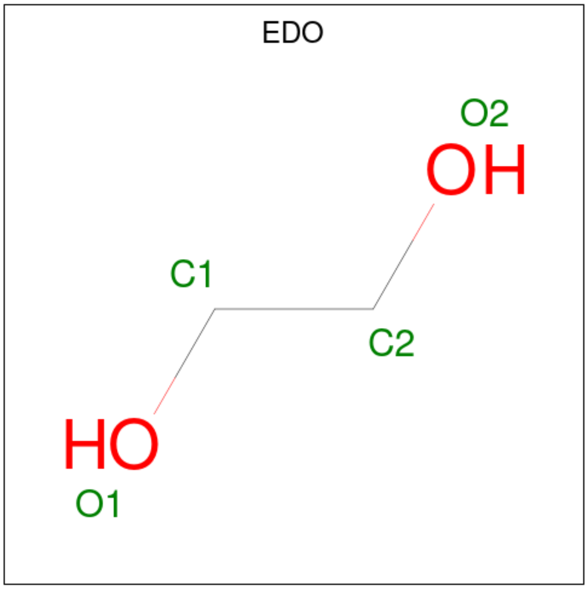


| Mol | Chain | Residues | Atoms | | | ZeroOcc | AltConf |
|---|---|---|---|---|---|---|---|
| | | | Total | C | O | | |
| 8 | A | 1 | 4 | 2 | 2 | 0 | 0 |
| 8 | A | 1 | 4 | 2 | 2 | 0 | 0 |
| 8 | A | 1 | 4 | 2 | 2 | 0 | 0 |
| 8 | B | 1 | 4 | 2 | 2 | 0 | 0 |
| 8 | C | 1 | 4 | 2 | 2 | 0 | 0 |
| 8 | C | 1 | 4 | 2 | 2 | 0 | 0 |
| 8 | C | 1 | 4 | 2 | 2 | 0 | 0 |
| 8 | C | 1 | 4 | 2 | 2 | 0 | 0 |
| 8 | D | 1 | 4 | 2 | 2 | 0 | 0 |
| 8 | D | 1 | 4 | 2 | 2 | 0 | 0 |

For Manuscript Review

- Molecule 9 is CHLORIDE ION (three-letter code: CL) (formula: Cl).

| Mol | Chain | Residues | Atoms | ZeroOcc | AltConf |
|---|---|---|---|---|---|
| 9 | A | 1 | Total Cl<br>1 1 | 0 | 0 |
| 9 | B | 1 | Total Cl<br>1 1 | 0 | 0 |
| 9 | C | 1 | Total Cl<br>1 1 | 0 | 0 |

- Molecule 10 is water.

| Mol | Chain | Residues | Atoms | ZeroOcc | AltConf |
|---|---|---|---|---|---|
| 10 | A | 62 | Total O<br>62 62 | 0 | 0 |
| 10 | B | 76 | Total O<br>76 76 | 0 | 0 |
| 10 | C | 76 | Total O<br>76 76 | 0 | 0 |
| 10 | D | 28 | Total O<br>28 28 | 0 | 0 |
| 10 | P | 9 | Total O<br>9 9 | 0 | 0 |

For Manuscript Review

### 3 Residue-property plots (i)

These plots are drawn for all protein, RNA, DNA and oligosaccharide chains in the entry. The first graphic for a chain summarises the proportions of the various outlier classes displayed in the second graphic. The second graphic shows the sequence view annotated by issues in geometry and electron density. Residues are color-coded according to the number of geometric quality criteria for which they contain at least one outlier: green = 0, yellow = 1, orange = 2 and red = 3 or more. A red dot above a residue indicates a poor fit to the electron density (RSRZ > 2). Stretches of 2 or more consecutive residues without any outlier are shown as a green connector. Residues present in the sample, but not in the model, are shown in grey.

- Molecule 1: NP207 TCR alpha-chain

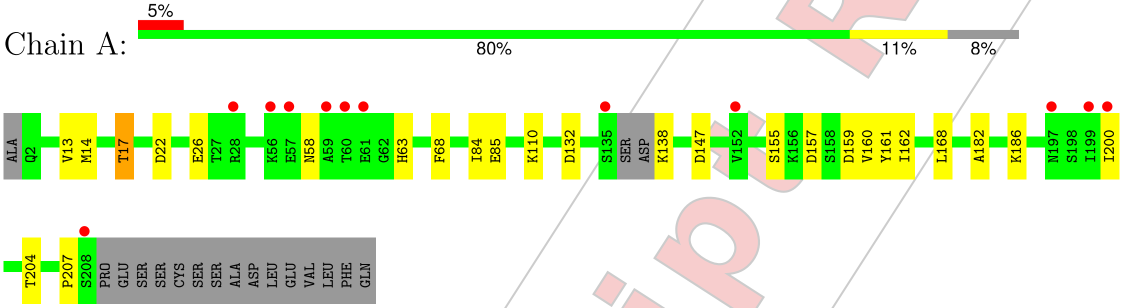


- Molecule 2: NP207 TCR beta-chain

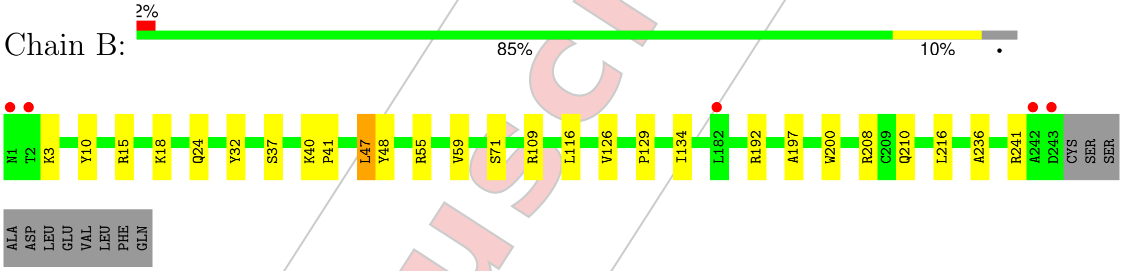


- Molecule 3: H-2 class I histocompatibility antigen, D-B alpha chain

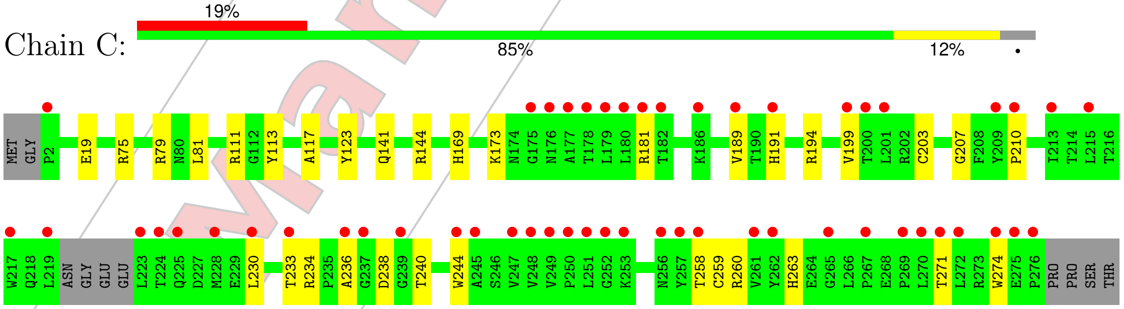


- Molecule 4: Beta-2-microglobulin

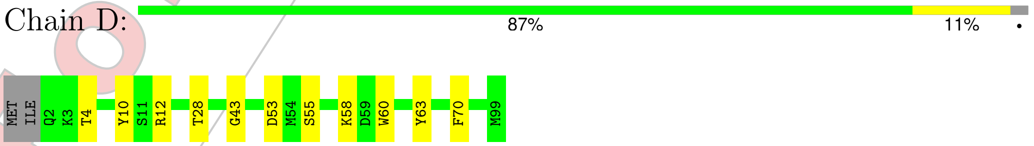


For Manuscript Review

• Molecule 5: Nucleoprotein peptide

Chain P: 100%

There are no outlier residues recorded for this chain.

• Molecule 6: beta-D-mannopyranose-(1-4)-2-acetamido-2-deoxy-beta-D-glucopyranose-(1-4)-2-acetamido-2-deoxy-beta-D-glucopyranose

Chain E: 100%

NAG1
NAG2
BMA3

For Manuscript Review

### 4 Data and refinement statistics (i)

| Property | Value | Source |
|---|---|---|
| Space group | P 1 | Depositor |
| Cell constants<br>a, b, c, $\alpha$, $\beta$, $\gamma$ | 57.53Å 68.02Å 74.68Å<br>72.83° 89.36° 67.21° | Depositor |
| Resolution (Å) | 44.87 – 1.95<br>44.87 – 1.95 | Depositor<br>EDS |
| % Data completeness<br>(in resolution range) | 94.4 (44.87-1.95)<br>95.0 (44.87-1.95) | Depositor<br>EDS |
| $R_{merge}$ | 0.05 | Depositor |
| $R_{sym}$ | (Not available) | Depositor |
| $< I/\sigma(I) >$ [1] | 1.36 (at 1.95Å) | Xtriage |
| Refinement program | PHENIX 1.21.2_5419 | Depositor |
| R, $R_{free}$ | 0.194 , 0.228<br>0.194 , 0.228 | Depositor<br>DCC |
| $R_{free}$ test set | 3513 reflections (4.86%) | wwPDB-VP |
| Wilson B-factor (Å$^2$) | 37.8 | Xtriage |
| Anisotropy | 0.363 | Xtriage |
| Bulk solvent $k_{sol}$(e/Å$^3$), $B_{sol}$(Å$^2$) | 0.32 , 40.9 | EDS |
| L-test for twinning[2] | $< \|L\| > = 0.50$, $< L^2 > = 0.33$ | Xtriage |
| Estimated twinning fraction | 0.011 for h,h-k,-l | Xtriage |
| $F_o$,$F_c$ correlation | 0.96 | EDS |
| Total number of atoms | 7027 | wwPDB-VP |
| Average B, all atoms (Å$^2$) | 58.0 | wwPDB-VP |

Xtriage's analysis on translational NCS is as follows: *The largest off-origin peak in the Patterson function is 5.84% of the height of the origin peak. No significant pseudotranslation is detected.*

[1]Intensities estimated from amplitudes.

[2]Theoretical values of $< |L| >$, $< L^2 >$ for acentric reflections are 0.5, 0.333 respectively for untwinned datasets, and 0.375, 0.2 for perfectly twinned datasets.

For Manuscript Review

### 5 Model quality (i)

#### 5.1 Standard geometry (i)

Bond lengths and bond angles in the following residue types are not validated in this section: NAG, CL, EDO, BMA

The Z score for a bond length (or angle) is the number of standard deviations the observed value is removed from the expected value. A bond length (or angle) with $|Z| > 5$ is considered an outlier worth inspection. RMSZ is the root-mean-square of all Z scores of the bond lengths (or angles).

| Mol | Chain | Bond lengths | | Bond angles | |
|---|---|---|---|---|---|
| | | RMSZ | #$|Z|$ >5 | RMSZ | #$|Z|$ >5 |
| 1 | A | 0.46 | 0/1630 | 0.61 | 1/2206 (0.0%) |
| 2 | B | 0.42 | 0/2014 | 0.61 | 0/2743 |
| 3 | C | 0.44 | 0/2269 | 0.63 | 0/3084 |
| 4 | D | 0.46 | 0/839 | 0.59 | 0/1137 |
| 5 | P | 0.61 | 0/68 | 0.69 | 0/88 |
| All | All | 0.45 | 0/6820 | 0.61 | 1/9258 (0.0%) |

There are no bond length outliers.

All (1) bond angle outliers are listed below:

| Mol | Chain | Res | Type | Atoms | Z | Observed($^o$) | Ideal($^o$) |
|---|---|---|---|---|---|---|---|
| 1 | A | 168 | LEU | CA-CB-CG | 5.43 | 127.79 | 115.30 |

There are no chirality outliers.

There are no planarity outliers.

#### 5.2 Too-close contacts (i)

In the following table, the Non-H and H(model) columns list the number of non-hydrogen atoms and hydrogen atoms in the chain respectively. The H(added) column lists the number of hydrogen atoms added and optimized by MolProbity. The Clashes column lists the number of clashes within the asymmetric unit, whereas Symm-Clashes lists symmetry-related clashes.

| Mol | Chain | Non-H | H(model) | H(added) | Clashes | Symm-Clashes |
|---|---|---|---|---|---|---|
| 1 | A | 1599 | 0 | 1518 | 13 | 0 |
| 2 | B | 1959 | 0 | 1881 | 13 | 0 |
| 3 | C | 2198 | 0 | 2014 | 18 | 0 |
| 4 | D | 813 | 0 | 782 | 10 | 0 |
| 5 | P | 69 | 0 | 61 | 0 | 0 |

*Continued on next page...*

For Manuscript Review

Continued from previous page...

| Mol | Chain | Non-H | H(model) | H(added) | Clashes | Symm-Clashes |
|---|---|---|---|---|---|---|
| 6 | E | 39 | 0 | 34 | 0 | 0 |
| 7 | A | 42 | 0 | 39 | 0 | 0 |
| 7 | B | 14 | 0 | 13 | 0 | 0 |
| 8 | A | 12 | 0 | 18 | 1 | 0 |
| 8 | B | 4 | 0 | 6 | 0 | 0 |
| 8 | C | 16 | 0 | 24 | 3 | 0 |
| 8 | D | 8 | 0 | 12 | 3 | 0 |
| 9 | A | 1 | 0 | 0 | 0 | 0 |
| 9 | B | 1 | 0 | 0 | 0 | 0 |
| 9 | C | 1 | 0 | 0 | 0 | 0 |
| 10 | A | 62 | 0 | 0 | 0 | 0 |
| 10 | B | 76 | 0 | 0 | 0 | 0 |
| 10 | C | 76 | 0 | 0 | 0 | 0 |
| 10 | D | 28 | 0 | 0 | 0 | 0 |
| 10 | P | 9 | 0 | 0 | 0 | 0 |
| All | All | 7027 | 0 | 6402 | 48 | 0 |

The all-atom clashscore is defined as the number of clashes found per 1000 atoms (including hydrogen atoms). The all-atom clashscore for this structure is 4.

All (48) close contacts within the same asymmetric unit are listed below, sorted by their clash magnitude.

| Atom-1 | Atom-2 | Interatomic distance (Å) | Clash overlap (Å) |
|---|---|---|---|
| 3:C:236:ALA:HB1 | 4:D:12:ARG:HG3 | 1.51 | 0.91 |
| 8:C:304:EDO:H22 | 4:D:58:LYS:HE2 | 1.62 | 0.81 |
| 1:A:159:ASP:HB2 | 1:A:186:LYS:HE2 | 1.75 | 0.67 |
| 1:A:200:ILE:HD11 | 1:A:204:THR:HG21 | 1.78 | 0.65 |
| 2:B:126:VAL:HG23 | 2:B:236:ALA:HB3 | 1.82 | 0.61 |
| 1:A:132:ASP:HB2 | 1:A:138:LYS:HB2 | 1.81 | 0.61 |
| 3:C:19:GLU:OE1 | 3:C:75[A]:ARG:HD2 | 2.05 | 0.57 |
| 2:B:116:LEU:HD12 | 2:B:216:LEU:HD21 | 1.87 | 0.57 |
| 3:C:169:HIS:O | 3:C:173:LYS:HD3 | 2.05 | 0.57 |
| 4:D:53:ASP:H | 8:D:102:EDO:H11 | 1.71 | 0.55 |
| 3:C:117:ALA:HB2 | 4:D:60:TRP:CE2 | 2.42 | 0.55 |
| 1:A:110:LYS:NZ | 2:B:41:PRO:HD3 | 2.22 | 0.54 |
| 1:A:157:ASP:HB3 | 1:A:160:VAL:HB | 1.88 | 0.54 |
| 3:C:189:VAL:HG22 | 3:C:203:CYS:HB3 | 1.89 | 0.54 |
| 2:B:10:TYR:CE1 | 2:B:109[A]:ARG:HD3 | 2.43 | 0.53 |
| 3:C:230:LEU:HD12 | 3:C:244:TRP:O | 2.08 | 0.53 |
| 3:C:194:ARG:H | 3:C:199:VAL:HA | 1.74 | 0.53 |
| 2:B:37:SER:HB2 | 2:B:40:LYS:HE2 | 1.91 | 0.52 |

Continued on next page...

For Manuscript Review

*Continued from previous page...*

| Atom-1 | Atom-2 | Interatomic distance (Å) | Clash overlap (Å) |
|---|---|---|---|
| 3:C:141:GLN:HE22 | 3:C:144:ARG:NH2 | 2.09 | 0.51 |
| 1:A:14:MET:O | 1:A:17:THR:HB | 2.12 | 0.49 |
| 1:A:84:ILE:HG13 | 1:A:85:GLU:OE2 | 2.11 | 0.49 |
| 3:C:207:GLY:HA2 | 3:C:240:THR:HB | 1.93 | 0.49 |
| 1:A:13:VAL:HB | 1:A:17:THR:HG21 | 1.94 | 0.49 |
| 3:C:191:HIS:NE2 | 3:C:199:VAL:HG21 | 2.28 | 0.49 |
| 2:B:134:ILE:HG23 | 2:B:197:ALA:HB1 | 1.95 | 0.48 |
| 4:D:43:GLY:H | 8:D:101:EDO:H22 | 1.79 | 0.47 |
| 3:C:75[A]:ARG:O | 3:C:79[A]:ARG:HG2 | 2.14 | 0.47 |
| 1:A:161:TYR:O | 1:A:182:ALA:HA | 2.15 | 0.47 |
| 3:C:210:PRO:HD2 | 3:C:263:HIS:CE1 | 2.50 | 0.47 |
| 3:C:238:ASP:HB3 | 4:D:12:ARG:NH1 | 2.30 | 0.46 |
| 1:A:22:ASP:HB3 | 8:A:305:EDO:H21 | 1.97 | 0.46 |
| 3:C:123:TYR:HD1 | 8:C:303:EDO:H11 | 1.79 | 0.46 |
| 2:B:129:PRO:HD2 | 2:B:200:TRP:CZ2 | 2.51 | 0.45 |
| 1:A:155:SER:HB3 | 1:A:162:ILE:HD12 | 1.97 | 0.45 |
| 4:D:43:GLY:HA2 | 8:D:101:EDO:H22 | 1.99 | 0.45 |
| 3:C:191:HIS:HB2 | 3:C:274:TRP:NE1 | 2.33 | 0.44 |
| 4:D:55:SER:HB2 | 4:D:63:TYR:CZ | 2.52 | 0.44 |
| 2:B:32:TYR:CE2 | 2:B:47:LEU:HD13 | 2.53 | 0.44 |
| 4:D:28:THR:HG22 | 4:D:63:TYR:HB2 | 2.00 | 0.43 |
| 1:A:58:ASN:OD1 | 1:A:68:PHE:HB3 | 2.19 | 0.43 |
| 1:A:110:LYS:NZ | 2:B:40:LYS:HA | 2.34 | 0.42 |
| 2:B:47:LEU:HD12 | 2:B:48:TYR:N | 2.34 | 0.42 |
| 2:B:208:ARG:NH1 | 2:B:210:GLN:HB2 | 2.35 | 0.41 |
| 3:C:260:ARG:HD2 | 3:C:271:THR:HG22 | 2.02 | 0.41 |
| 2:B:24:GLN:O | 2:B:71:SER:HB2 | 2.21 | 0.41 |
| 2:B:15:ARG:HH21 | 2:B:18:LYS:NZ | 2.18 | 0.41 |
| 3:C:234:ARG:HD2 | 4:D:10:TYR:CE1 | 2.55 | 0.41 |
| 3:C:113:TYR:CD2 | 8:C:304:EDO:H12 | 2.56 | 0.40 |

There are no symmetry-related clashes.

#### 5.3 Torsion angles (i)

##### 5.3.1 Protein backbone (i)

In the following table, the Percentiles column shows the percent Ramachandran outliers of the chain as a percentile score with respect to all X-ray entries followed by that with respect to entries of similar resolution.

For Manuscript Review

The Analysed column shows the number of residues for which the backbone conformation was analysed, and the total number of residues.

| Mol | Chain | Analysed | Favoured | Allowed | Outliers | Percentiles | |
|---|---|---|---|---|---|---|---|
| 1 | A | 202/223 (91%) | 194 (96%) | 7 (4%) | 1 (0%) | 25 | 16 |
| 2 | B | 242/254 (95%) | 235 (97%) | 7 (3%) | 0 | 100 | 100 |
| 3 | C | 271/281 (96%) | 258 (95%) | 13 (5%) | 0 | 100 | 100 |
| 4 | D | 96/100 (96%) | 94 (98%) | 2 (2%) | 0 | 100 | 100 |
| 5 | P | 7/9 (78%) | 6 (86%) | 1 (14%) | 0 | 100 | 100 |
| All | All | 818/867 (94%) | 787 (96%) | 30 (4%) | 1 (0%) | 48 | 42 |

All (1) Ramachandran outliers are listed below:

| Mol | Chain | Res | Type |
|---|---|---|---|
| 1 | A | 207 | PRO |

##### 5.3.2 Protein sidechains (i)

In the following table, the Percentiles column shows the percent sidechain outliers of the chain as a percentile score with respect to all X-ray entries followed by that with respect to entries of similar resolution.

The Analysed column shows the number of residues for which the sidechain conformation was analysed, and the total number of residues.

| Mol | Chain | Analysed | Rotameric | Outliers | Percentiles | |
|---|---|---|---|---|---|---|
| 1 | A | 180/201 (90%) | 176 (98%) | 4 (2%) | 47 | 41 |
| 2 | B | 215/225 (96%) | 209 (97%) | 6 (3%) | 38 | 29 |
| 3 | C | 216/239 (90%) | 210 (97%) | 6 (3%) | 38 | 29 |
| 4 | D | 93/95 (98%) | 91 (98%) | 2 (2%) | 47 | 41 |
| 5 | P | 8/8 (100%) | 8 (100%) | 0 | 100 | 100 |
| All | All | 712/768 (93%) | 694 (98%) | 18 (2%) | 42 | 34 |

All (18) residues with a non-rotameric sidechain are listed below:

| Mol | Chain | Res | Type |
|---|---|---|---|
| 1 | A | 17 | THR |
| 1 | A | 26 | GLU |
| 1 | A | 63 | HIS |
| 1 | A | 147 | ASP |

*Continued on next page...*

For Manuscript Review

*Continued from previous page...*

| Mol | Chain | Res | Type |
|---|---|---|---|
| 2 | B | 3 | LYS |
| 2 | B | 47 | LEU |
| 2 | B | 55 | ARG |
| 2 | B | 59 | VAL |
| 2 | B | 192 | ARG |
| 2 | B | 241 | ARG |
| 3 | C | 81 | LEU |
| 3 | C | 111 | ARG |
| 3 | C | 181 | ARG |
| 3 | C | 233 | THR |
| 3 | C | 258 | THR |
| 3 | C | 259 | CYS |
| 4 | D | 4 | THR |
| 4 | D | 70 | PHE |

Sometimes sidechains can be flipped to improve hydrogen bonding and reduce clashes. All (2) such sidechains are listed below:

| Mol | Chain | Res | Type |
|---|---|---|---|
| 3 | C | 141 | GLN |
| 3 | C | 174 | ASN |

##### 5.3.3 RNA ⓘ

There are no RNA molecules in this entry.

#### 5.4 Non-standard residues in protein, DNA, RNA chains ⓘ

There are no non-standard protein/DNA/RNA residues in this entry.

#### 5.5 Carbohydrates ⓘ

3 monosaccharides are modelled in this entry.

In the following table, the Counts columns list the number of bonds (or angles) for which Mogul statistics could be retrieved, the number of bonds (or angles) that are observed in the model and the number of bonds (or angles) that are defined in the Chemical Component Dictionary. The Link column lists molecule types, if any, to which the group is linked. The Z score for a bond length (or angle) is the number of standard deviations the observed value is removed from the expected value. A bond length (or angle) with $|Z| > 2$ is considered an outlier worth inspection. RMSZ is the root-mean-square of all Z scores of the bond lengths (or angles).

For Manuscript Review

| Mol | Type | Chain | Res | Link | Bond lengths | | | Bond angles | | |
|---|---|---|---|---|---|---|---|---|---|---|
| | | | | | Counts | RMSZ | #\|Z\| > 2 | Counts | RMSZ | #\|Z\| > 2 |
| 6 | NAG | E | 1 | 6,2 | 14,14,15 | 0.75 | 0 | 17,19,21 | 1.38 | 3 (17%) |
| 6 | NAG | E | 2 | 6 | 14,14,15 | 0.71 | 0 | 17,19,21 | 1.19 | 3 (17%) |
| 6 | BMA | E | 3 | 6 | 11,11,12 | 0.91 | 0 | 15,15,17 | 2.28 | 5 (33%) |

In the following table, the Chirals column lists the number of chiral outliers, the number of chiral centers analysed, the number of these observed in the model and the number defined in the Chemical Component Dictionary. Similar counts are reported in the Torsion and Rings columns. '-' means no outliers of that kind were identified.

| Mol | Type | Chain | Res | Link | Chirals | Torsions | Rings |
|---|---|---|---|---|---|---|---|
| 6 | NAG | E | 1 | 6,2 | - | 2/6/23/26 | 0/1/1/1 |
| 6 | NAG | E | 2 | 6 | - | 0/6/23/26 | 0/1/1/1 |
| 6 | BMA | E | 3 | 6 | - | 2/2/19/22 | 0/1/1/1 |

There are no bond length outliers.

All (11) bond angle outliers are listed below:

| Mol | Chain | Res | Type | Atoms | Z | Observed($^o$) | Ideal($^o$) |
|---|---|---|---|---|---|---|---|
| 6 | E | 3 | BMA | C1-O5-C5 | 5.34 | 119.35 | 112.19 |
| 6 | E | 3 | BMA | C3-C4-C5 | 3.88 | 117.27 | 110.23 |
| 6 | E | 3 | BMA | C2-C3-C4 | 3.20 | 116.49 | 110.86 |
| 6 | E | 2 | NAG | O5-C1-C2 | -2.98 | 106.69 | 111.29 |
| 6 | E | 3 | BMA | O4-C4-C3 | -2.60 | 104.24 | 110.38 |
| 6 | E | 1 | NAG | O5-C1-C2 | -2.30 | 107.73 | 111.29 |
| 6 | E | 3 | BMA | O5-C5-C4 | 2.24 | 116.28 | 110.83 |
| 6 | E | 1 | NAG | O4-C4-C3 | -2.18 | 105.23 | 110.38 |
| 6 | E | 2 | NAG | O4-C4-C3 | -2.12 | 105.38 | 110.38 |
| 6 | E | 1 | NAG | C3-C4-C5 | -2.09 | 106.44 | 110.23 |
| 6 | E | 2 | NAG | C4-C3-C2 | 2.06 | 114.04 | 111.02 |

There are no chirality outliers.

All (4) torsion outliers are listed below:

| Mol | Chain | Res | Type | Atoms |
|---|---|---|---|---|
| 6 | E | 3 | BMA | O5-C5-C6-O6 |
| 6 | E | 1 | NAG | C4-C5-C6-O6 |
| 6 | E | 3 | BMA | C4-C5-C6-O6 |
| 6 | E | 1 | NAG | O5-C5-C6-O6 |

There are no ring outliers.

For Manuscript Review

No monomer is involved in short contacts.

The following is a two-dimensional graphical depiction of Mogul quality analysis of bond lengths, bond angles, torsion angles, and ring geometry for oligosaccharide.

**Oligosaccharide Chain E**

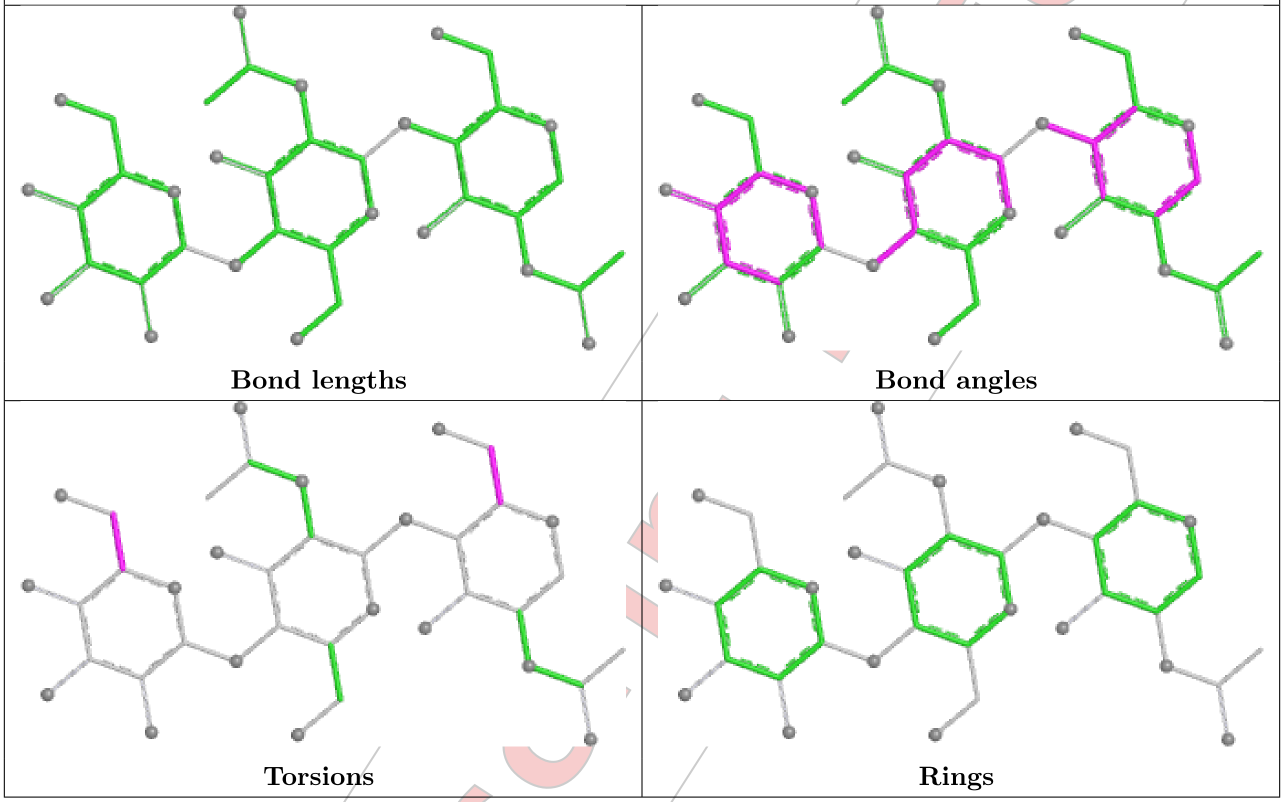


#### 5.6 Ligand geometry (i)

Of 17 ligands modelled in this entry, 3 are monoatomic - leaving 14 for Mogul analysis.

In the following table, the Counts columns list the number of bonds (or angles) for which Mogul statistics could be retrieved, the number of bonds (or angles) that are observed in the model and the number of bonds (or angles) that are defined in the Chemical Component Dictionary. The Link column lists molecule types, if any, to which the group is linked. The Z score for a bond length (or angle) is the number of standard deviations the observed value is removed from the expected value. A bond length (or angle) with $|Z| > 2$ is considered an outlier worth inspection. RMSZ is the root-mean-square of all Z scores of the bond lengths (or angles).

| Mol | Type | Chain | Res | Link | Bond lengths | | | Bond angles | | |
|---|---|---|---|---|---|---|---|---|---|---|
| | | | | | Counts | RMSZ | $\#\|Z\| > 2$ | Counts | RMSZ | $\#\|Z\| > 2$ |
| 8 | EDO | C | 301 | - | 3,3,3 | 0.27 | 0 | 2,2,2 | 0.18 | 0 |
| 8 | EDO | A | 305 | - | 3,3,3 | 0.26 | 0 | 2,2,2 | 0.38 | 0 |
| 8 | EDO | C | 304 | - | 3,3,3 | 0.25 | 0 | 2,2,2 | 0.25 | 0 |
| 8 | EDO | B | 302 | - | 3,3,3 | 0.26 | 0 | 2,2,2 | 0.22 | 0 |
| 7 | NAG | A | 301 | 1 | 14,14,15 | 0.70 | 0 | 17,19,21 | 1.16 | 2 (11%) |

For Manuscript Review

| Mol | Type | Chain | Res | Link | Bond lengths | | | Bond angles | | |
|---|---|---|---|---|---|---|---|---|---|---|
| | | | | | Counts | RMSZ | #\|Z\| > 2 | Counts | RMSZ | #\|Z\| > 2 |
| 8 | EDO | C | 302 | - | 3,3,3 | 0.26 | 0 | 2,2,2 | 0.41 | 0 |
| 8 | EDO | D | 102 | - | 3,3,3 | 0.29 | 0 | 2,2,2 | 0.20 | 0 |
| 8 | EDO | D | 101 | - | 3,3,3 | 0.28 | 0 | 2,2,2 | 0.51 | 0 |
| 8 | EDO | C | 303 | - | 3,3,3 | 0.31 | 0 | 2,2,2 | 0.43 | 0 |
| 7 | NAG | A | 303 | 1 | 14,14,15 | 0.70 | 0 | 17,19,21 | 0.92 | 1 (5%) |
| 8 | EDO | A | 306 | - | 3,3,3 | 0.21 | 0 | 2,2,2 | 0.43 | 0 |
| 8 | EDO | A | 304 | - | 3,3,3 | 0.25 | 0 | 2,2,2 | 0.39 | 0 |
| 7 | NAG | B | 301 | 2 | 14,14,15 | 0.79 | 0 | 17,19,21 | 0.93 | 0 |
| 7 | NAG | A | 302 | 1 | 14,14,15 | 1.04 | 1 (7%) | 17,19,21 | 1.40 | 1 (5%) |

In the following table, the Chirals column lists the number of chiral outliers, the number of chiral centers analysed, the number of these observed in the model and the number defined in the Chemical Component Dictionary. Similar counts are reported in the Torsion and Rings columns. '-' means no outliers of that kind were identified.

| Mol | Type | Chain | Res | Link | Chirals | Torsions | Rings |
|---|---|---|---|---|---|---|---|
| 8 | EDO | C | 301 | - | - | 0/1/1/1 | - |
| 8 | EDO | A | 305 | - | - | 1/1/1/1 | - |
| 8 | EDO | C | 304 | - | - | 0/1/1/1 | - |
| 8 | EDO | B | 302 | - | - | 1/1/1/1 | - |
| 7 | NAG | A | 301 | 1 | - | 0/6/23/26 | 0/1/1/1 |
| 8 | EDO | C | 302 | - | - | 1/1/1/1 | - |
| 8 | EDO | D | 102 | - | - | 1/1/1/1 | - |
| 8 | EDO | D | 101 | - | - | 0/1/1/1 | - |
| 8 | EDO | C | 303 | - | - | 1/1/1/1 | - |
| 7 | NAG | A | 303 | 1 | - | 2/6/23/26 | 0/1/1/1 |
| 8 | EDO | A | 306 | - | - | 1/1/1/1 | - |
| 8 | EDO | A | 304 | - | - | 0/1/1/1 | - |
| 7 | NAG | B | 301 | 2 | - | 1/6/23/26 | 0/1/1/1 |
| 7 | NAG | A | 302 | 1 | - | 0/6/23/26 | 0/1/1/1 |

All (1) bond length outliers are listed below:

| Mol | Chain | Res | Type | Atoms | Z | Observed(Å) | Ideal(Å) |
|---|---|---|---|---|---|---|---|
| 7 | A | 302 | NAG | C1-C2 | 3.23 | 1.56 | 1.52 |

All (4) bond angle outliers are listed below:

| Mol | Chain | Res | Type | Atoms | Z | Observed(°) | Ideal(°) |
|---|---|---|---|---|---|---|---|
| 7 | A | 302 | NAG | C2-N2-C7 | 4.17 | 128.49 | 122.90 |
| 7 | A | 301 | NAG | C2-N2-C7 | 2.71 | 126.53 | 122.90 |

*Continued on next page...*

For Manuscript Review

*Continued from previous page...*

| Mol | Chain | Res | Type | Atoms | Z | Observed(°) | Ideal(°) |
|---|---|---|---|---|---|---|---|
| 7 | A | 303 | NAG | C2-N2-C7 | 2.29 | 125.97 | 122.90 |
| 7 | A | 301 | NAG | O5-C1-C2 | -2.11 | 108.03 | 111.29 |

There are no chirality outliers.

All (9) torsion outliers are listed below:

| Mol | Chain | Res | Type | Atoms |
|---|---|---|---|---|
| 8 | C | 303 | EDO | O1-C1-C2-O2 |
| 8 | D | 102 | EDO | O1-C1-C2-O2 |
| 8 | A | 306 | EDO | O1-C1-C2-O2 |
| 8 | B | 302 | EDO | O1-C1-C2-O2 |
| 7 | A | 303 | NAG | C4-C5-C6-O6 |
| 8 | A | 305 | EDO | O1-C1-C2-O2 |
| 7 | A | 303 | NAG | O5-C5-C6-O6 |
| 8 | C | 302 | EDO | O1-C1-C2-O2 |
| 7 | B | 301 | NAG | C4-C5-C6-O6 |

There are no ring outliers.

5 monomers are involved in 7 short contacts:

| Mol | Chain | Res | Type | Clashes | Symm-Clashes |
|---|---|---|---|---|---|
| 8 | A | 305 | EDO | 1 | 0 |
| 8 | C | 304 | EDO | 2 | 0 |
| 8 | D | 102 | EDO | 1 | 0 |
| 8 | D | 101 | EDO | 2 | 0 |
| 8 | C | 303 | EDO | 1 | 0 |

The following is a two-dimensional graphical depiction of Mogul quality analysis of bond lengths, bond angles, torsion angles, and ring geometry for all instances of the Ligand of Interest. In addition, ligands with molecular weight > 250 and outliers as shown on the validation Tables will also be included. For torsion angles, if less then 5% of the Mogul distribution of torsion angles is within 10 degrees of the torsion angle in question, then that torsion angle is considered an outlier. Any bond that is central to one or more torsion angles identified as an outlier by Mogul will be highlighted in the graph. For rings, the root-mean-square deviation (RMSD) between the ring in question and similar rings identified by Mogul is calculated over all ring torsion angles. If the average RMSD is greater than 60 degrees and the minimal RMSD between the ring in question and any Mogul-identified rings is also greater than 60 degrees, then that ring is considered an outlier. The outliers are highlighted in purple. The color gray indicates Mogul did not find sufficient equivalents in the CSD to analyse the geometry.

For Manuscript Review

**Ligand NAG A 301**

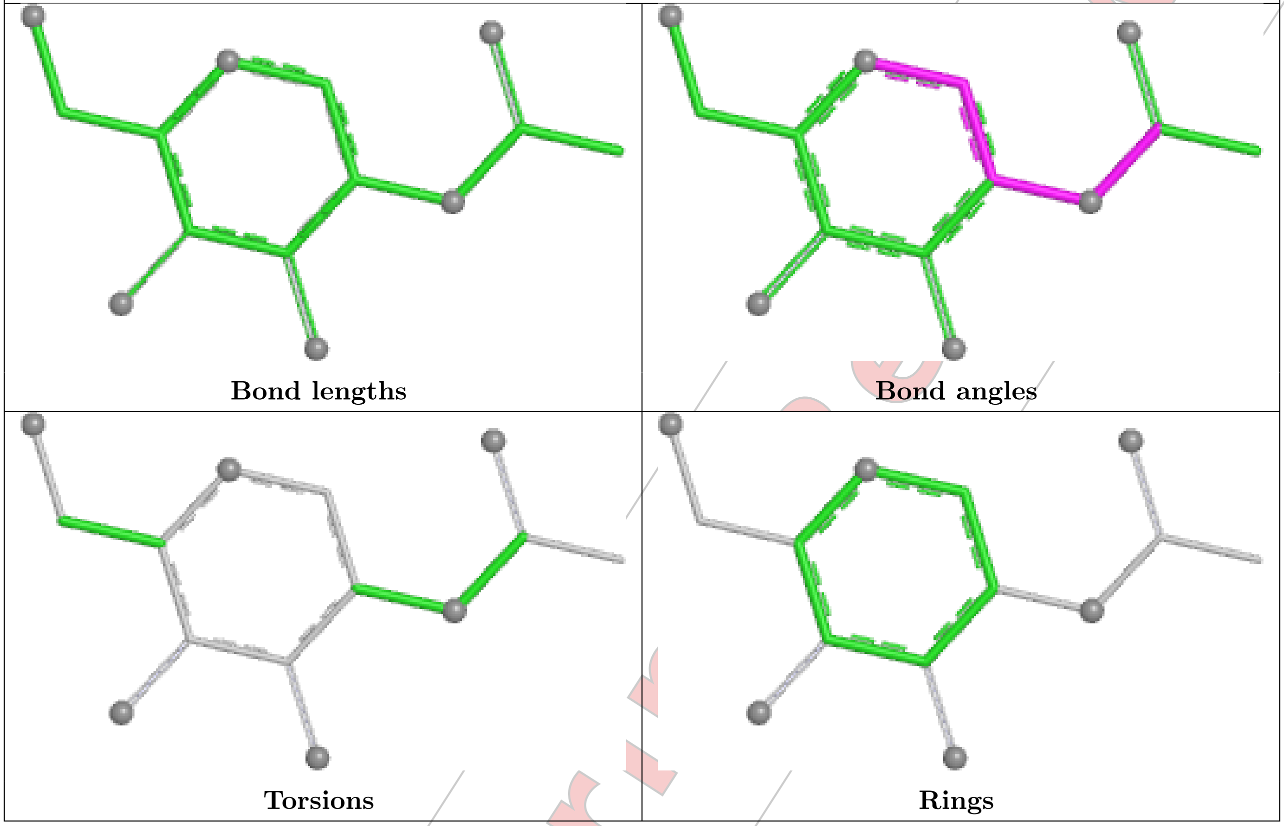


**Ligand NAG A 303**

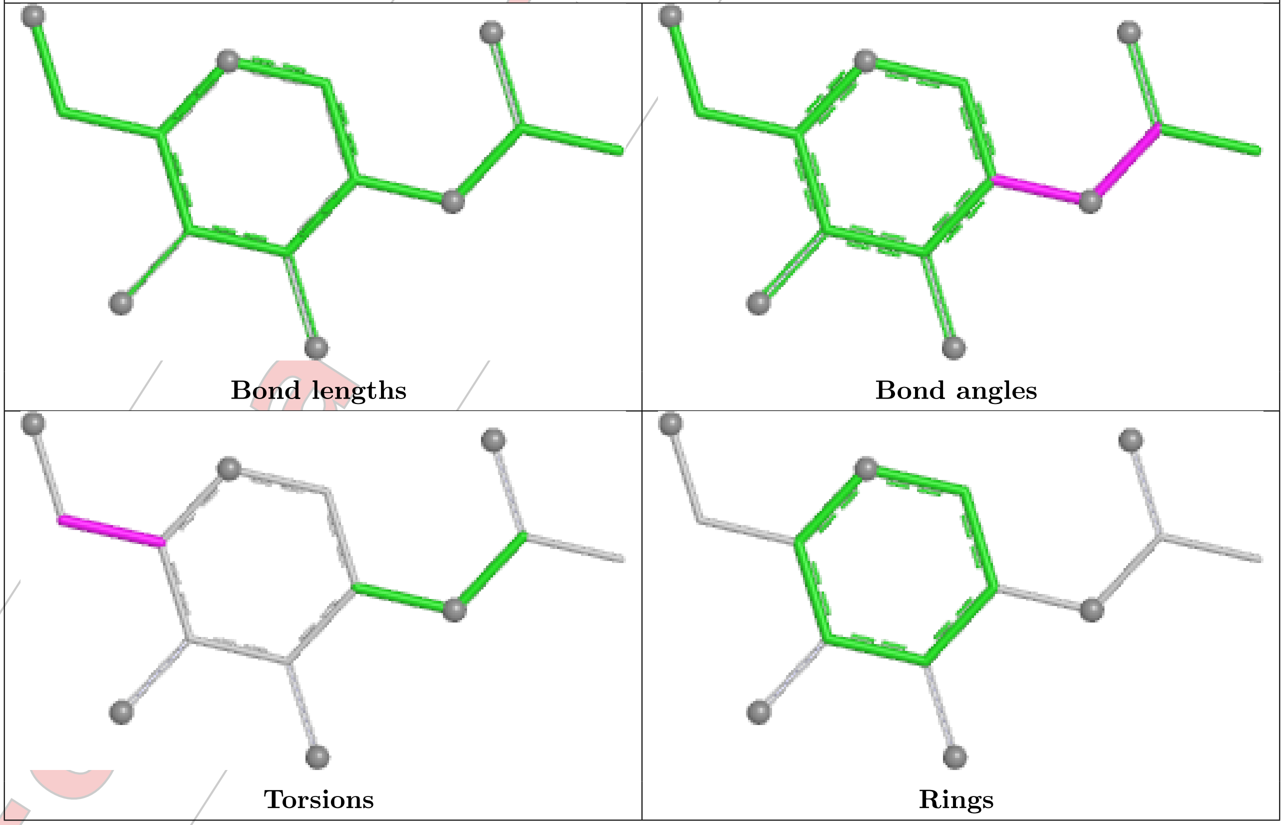

#### Ligand NAG B 301

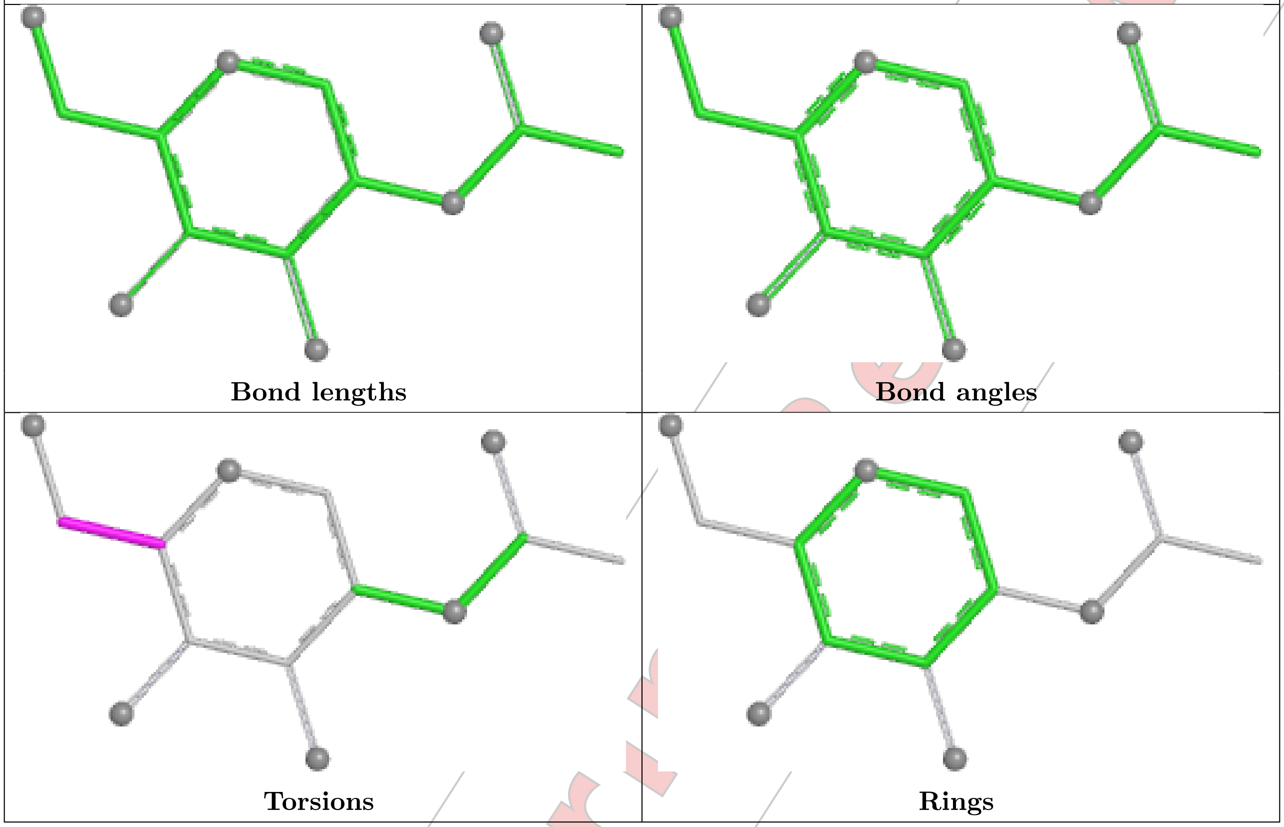


#### Ligand NAG A 302

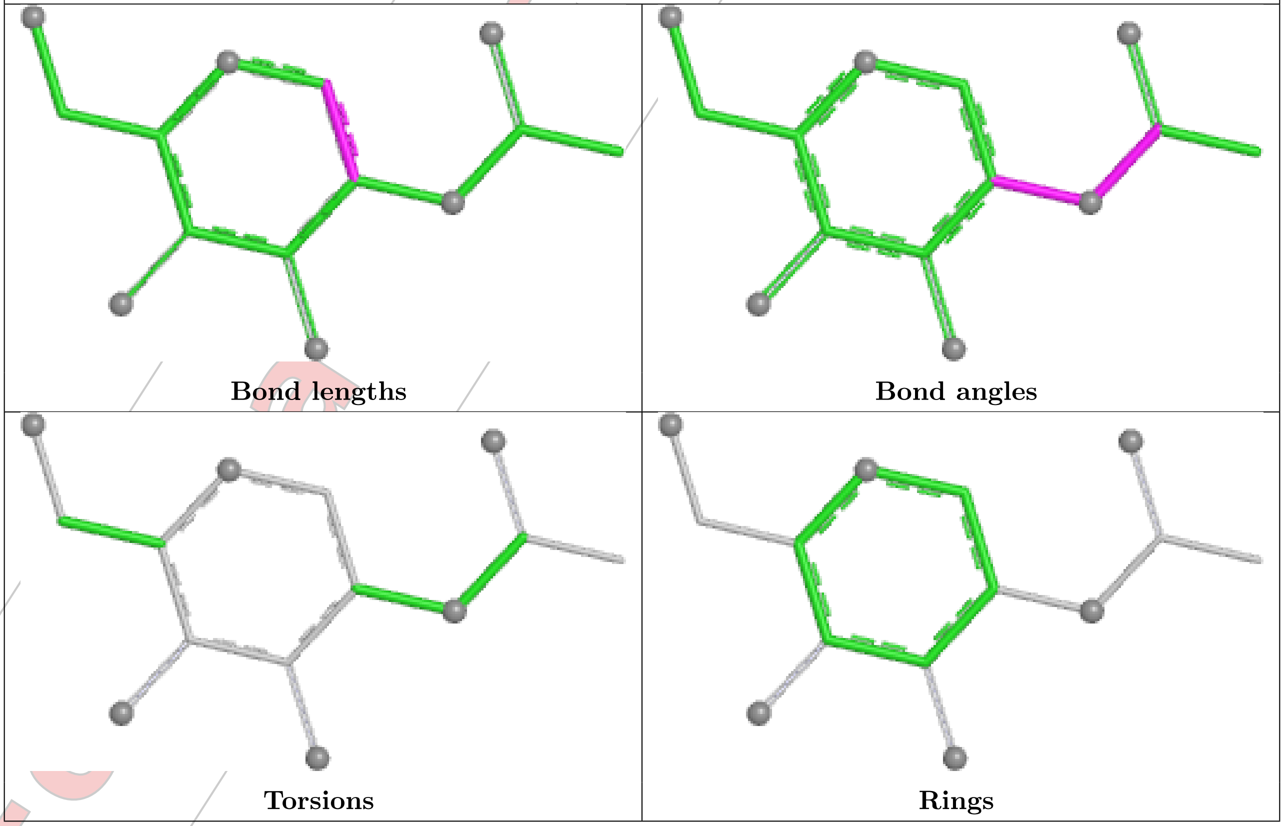

#### 5.7 Other polymers (i)

There are no such residues in this entry.

#### 5.8 Polymer linkage issues (i)

There are no chain breaks in this entry.

For Manuscript Review

### 6 Fit of model and data (i)

#### 6.1 Protein, DNA and RNA chains (i)

In the following table, the column labelled '#RSRZ> 2' contains the number (and percentage) of RSRZ outliers, followed by percent RSRZ outliers for the chain as percentile scores relative to all X-ray entries and entries of similar resolution. The OWAB column contains the minimum, median, $95^{th}$ percentile and maximum values of the occupancy-weighted average B-factor per residue. The column labelled 'Q< 0.9' lists the number of (and percentage) of residues with an average occupancy less than 0.9.

| Mol | Chain | Analysed | <RSRZ> | #RSRZ>2 | | | OWAB(Å$^2$) | Q<0.9 |
|---|---|---|---|---|---|---|---|---|
| 1 | A | 205/223 (91%) | 0.46 | 12 (5%) | 29 | 36 | 24, 51, 100, 122 | 1 (0%) |
| 2 | B | 243/254 (95%) | 0.19 | 5 (2%) | 63 | 69 | 21, 48, 80, 115 | 1 (0%) |
| 3 | C | 271/281 (96%) | 0.77 | 53 (19%) | 4 | 4 | 18, 50, 145, 174 | 4 (1%) |
| 4 | D | 98/100 (98%) | 0.32 | 0 | 100 | 100 | 37, 51, 82, 91 | 0 |
| 5 | P | 9/9 (100%) | -0.64 | 0 | 100 | 100 | 30, 31, 39, 41 | 0 |
| All | All | 826/867 (95%) | 0.45 | 70 (8%) | 18 | 22 | 18, 50, 129, 174 | 6 (0%) |

All (70) RSRZ outliers are listed below:

| Mol | Chain | Res | Type | RSRZ |
|---|---|---|---|---|
| 3 | C | 215 | LEU | 4.8 |
| 3 | C | 248 | VAL | 4.6 |
| 3 | C | 261 | VAL | 4.6 |
| 3 | C | 251 | LEU | 4.5 |
| 1 | A | 59 | ALA | 4.4 |
| 3 | C | 237 | GLY | 4.2 |
| 3 | C | 180 | LEU | 4.1 |
| 1 | A | 135 | SER | 4.1 |
| 3 | C | 217 | TRP | 4.1 |
| 3 | C | 219 | LEU | 4.0 |
| 3 | C | 223 | LEU | 3.9 |
| 3 | C | 175 | GLY | 3.9 |
| 3 | C | 179 | LEU | 3.7 |
| 3 | C | 276 | PRO | 3.5 |
| 3 | C | 178 | THR | 3.4 |
| 3 | C | 177 | ALA | 3.4 |
| 3 | C | 272 | LEU | 3.4 |
| 3 | C | 230 | LEU | 3.3 |
| 3 | C | 274 | TRP | 3.3 |
| 3 | C | 236 | ALA | 3.2 |

*Continued on next page...*

For Manuscript Review

Continued from previous page...

| Mol | Chain | Res | Type | RSRZ |
|---|---|---|---|---|
| 3 | C | 182 | THR | 3.2 |
| 3 | C | 265 | GLY | 3.2 |
| 2 | B | 243 | ASP | 3.2 |
| 3 | C | 189 | VAL | 3.2 |
| 3 | C | 233 | THR | 3.1 |
| 3 | C | 213 | ILE | 3.1 |
| 3 | C | 209 | TYR | 3.1 |
| 3 | C | 247 | VAL | 3.1 |
| 3 | C | 270 | LEU | 3.0 |
| 1 | A | 28 | ARG | 3.0 |
| 2 | B | 2 | THR | 3.0 |
| 1 | A | 60 | THR | 2.9 |
| 3 | C | 262 | TYR | 2.9 |
| 3 | C | 176 | ASN | 2.9 |
| 3 | C | 201 | LEU | 2.8 |
| 3 | C | 2 | PRO | 2.8 |
| 1 | A | 197 | ASN | 2.8 |
| 3 | C | 271 | THR | 2.8 |
| 1 | A | 208 | SER | 2.8 |
| 3 | C | 224 | THR | 2.7 |
| 2 | B | 182 | LEU | 2.7 |
| 3 | C | 257 | TYR | 2.7 |
| 2 | B | 1 | ASN | 2.6 |
| 1 | A | 61 | GLU | 2.6 |
| 3 | C | 269 | PRO | 2.6 |
| 3 | C | 210 | PRO | 2.5 |
| 3 | C | 256 | ASN | 2.5 |
| 3 | C | 252 | GLY | 2.5 |
| 1 | A | 56 | LYS | 2.5 |
| 3 | C | 245 | ALA | 2.5 |
| 1 | A | 199 | ILE | 2.5 |
| 1 | A | 200 | ILE | 2.4 |
| 3 | C | 225 | GLN | 2.3 |
| 3 | C | 249 | VAL | 2.3 |
| 1 | A | 152 | VAL | 2.3 |
| 3 | C | 228 | MET | 2.3 |
| 3 | C | 239 | GLY | 2.2 |
| 3 | C | 253 | LYS | 2.2 |
| 3 | C | 267 | PRO | 2.2 |
| 2 | B | 242 | ALA | 2.2 |
| 3 | C | 186 | LYS | 2.2 |
| 3 | C | 250 | PRO | 2.2 |

Continued on next page...

For Manuscript Review

*Continued from previous page...*

| Mol | Chain | Res | Type | RSRZ |
|---|---|---|---|---|
| 3 | C | 200 | THR | 2.2 |
| 3 | C | 258 | THR | 2.2 |
| 3 | C | 244 | TRP | 2.2 |
| 3 | C | 275 | GLU | 2.2 |
| 3 | C | 181 | ARG | 2.2 |
| 1 | A | 57 | GLU | 2.1 |
| 3 | C | 191 | HIS | 2.1 |
| 3 | C | 199 | VAL | 2.0 |

#### 6.2 Non-standard residues in protein, DNA, RNA chains (i)

There are no non-standard protein/DNA/RNA residues in this entry.

#### 6.3 Carbohydrates (i)

SUGAR-RSR INFOmissingINFO

#### 6.4 Ligands (i)

In the following table, the Atoms column lists the number of modelled atoms in the group and the number defined in the chemical component dictionary. The B-factors column lists the minimum, median, $95^{th}$ percentile and maximum values of B factors of atoms in the group. The column labelled 'Q< 0.9' lists the number of atoms with occupancy less than 0.9.

| Mol | Type | Chain | Res | Atoms | RSCC | RSR | B-factors(Å$^2$) | Q<0.9 |
|---|---|---|---|---|---|---|---|---|
| 7 | NAG | B | 301 | 14/15 | 0.53 | 0.19 | 76,85,89,89 | 0 |
| 7 | NAG | A | 303 | 14/15 | 0.55 | 0.17 | 91,95,98,100 | 14 |
| 7 | NAG | A | 302 | 14/15 | 0.67 | 0.15 | 64,66,68,69 | 0 |
| 8 | EDO | D | 102 | 4/4 | 0.75 | 0.15 | 50,53,55,62 | 0 |
| 7 | NAG | A | 301 | 14/15 | 0.77 | 0.11 | 71,79,81,81 | 0 |
| 8 | EDO | C | 304 | 4/4 | 0.79 | 0.26 | 54,56,56,57 | 0 |
| 8 | EDO | A | 304 | 4/4 | 0.80 | 0.13 | 68,69,70,70 | 0 |
| 8 | EDO | C | 302 | 4/4 | 0.86 | 0.15 | 51,53,54,59 | 0 |
| 8 | EDO | A | 306 | 4/4 | 0.87 | 0.16 | 45,46,54,56 | 0 |
| 8 | EDO | C | 301 | 4/4 | 0.89 | 0.13 | 44,49,51,52 | 0 |
| 8 | EDO | A | 305 | 4/4 | 0.91 | 0.11 | 54,55,57,58 | 0 |
| 8 | EDO | C | 303 | 4/4 | 0.91 | 0.20 | 45,47,47,53 | 0 |
| 8 | EDO | D | 101 | 4/4 | 0.92 | 0.13 | 38,42,45,52 | 0 |
| 8 | EDO | B | 302 | 4/4 | 0.93 | 0.11 | 44,44,47,52 | 0 |
| 9 | CL | C | 305 | 1/1 | 0.93 | 0.10 | 40,40,40,40 | 1 |

*Continued on next page...*

For Manuscript Review

*Continued from previous page...*

| Mol | Type | Chain | Res | Atoms | RSCC | RSR | B-factors($Å^2$) | Q<0.9 |
|---|---|---|---|---|---|---|---|---|
| 9 | CL | B | 303 | 1/1 | 0.95 | 0.11 | 58,58,58,58 | 0 |
| 9 | CL | A | 307 | 1/1 | 0.97 | 0.05 | 37,37,37,37 | 1 |

The following is a graphical depiction of the model fit to experimental electron density of all instances of the Ligand of Interest. In addition, ligands with molecular weight > 250 and outliers as shown on the geometry validation Tables will also be included. Each fit is shown from different orientation to approximate a three-dimensional view.

**Electron density around NAG B 301:**

2mF$_o$-DF$_c$ (at 0.7 rmsd) in gray
mF$_o$-DF$_c$ (at 3 rmsd) in purple (negative) and green (positive)

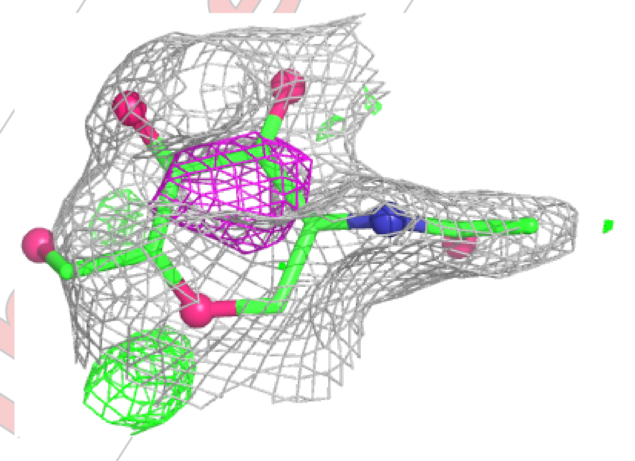

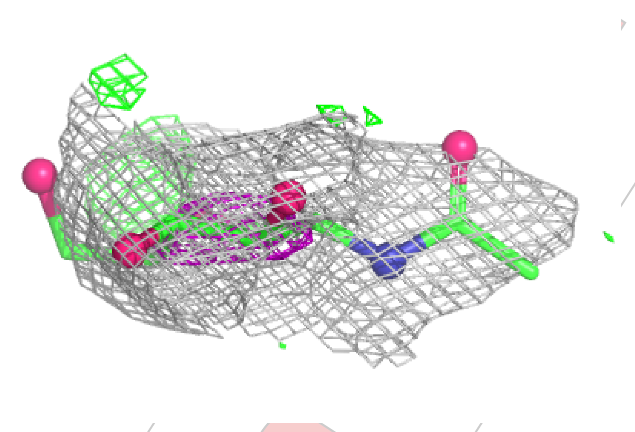

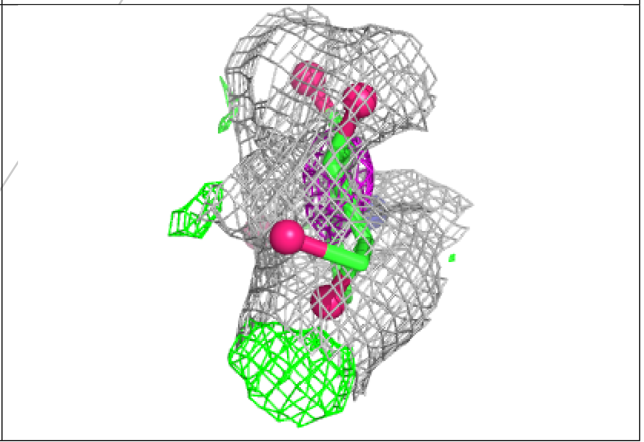

For Manuscript Review

**Electron density around NAG A 303:**

$2mF_o$-$DF_c$ (at 0.7 rmsd) in gray
$mF_o$-$DF_c$ (at 3 rmsd) in purple (negative)
and green (positive)

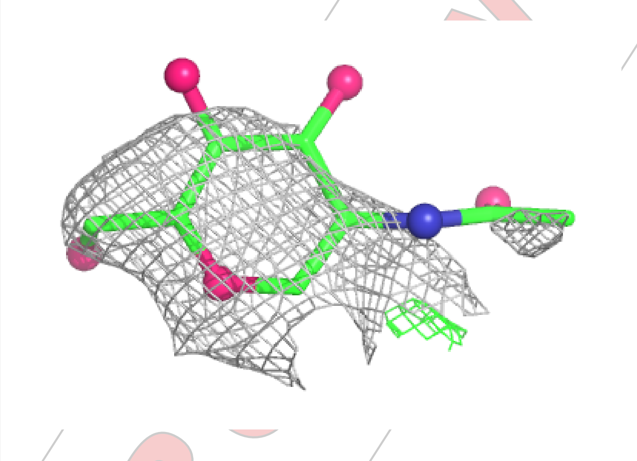

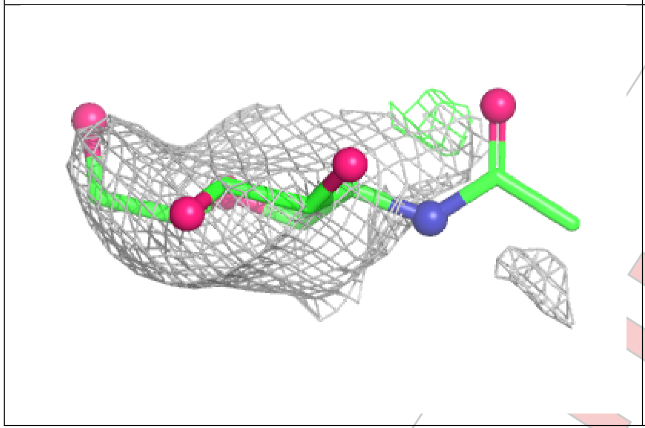

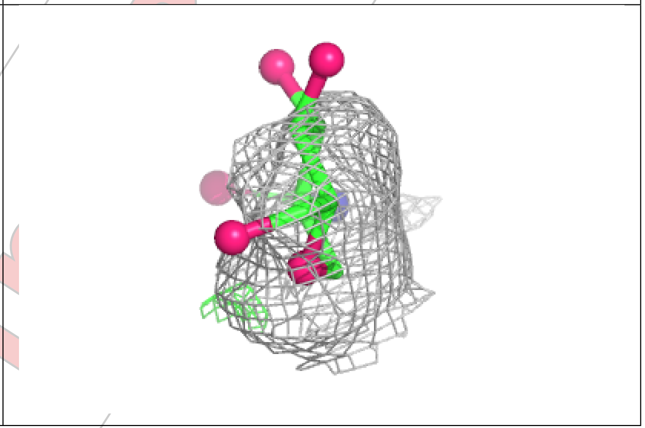

**Electron density around NAG A 302:**

$2mF_o$-$DF_c$ (at 0.7 rmsd) in gray
$mF_o$-$DF_c$ (at 3 rmsd) in purple (negative)
and green (positive)

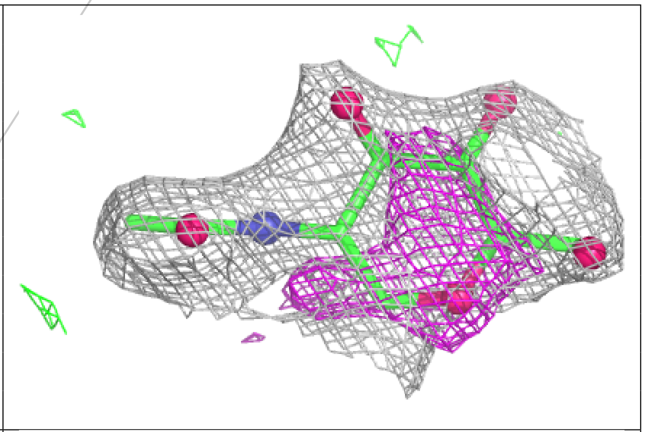

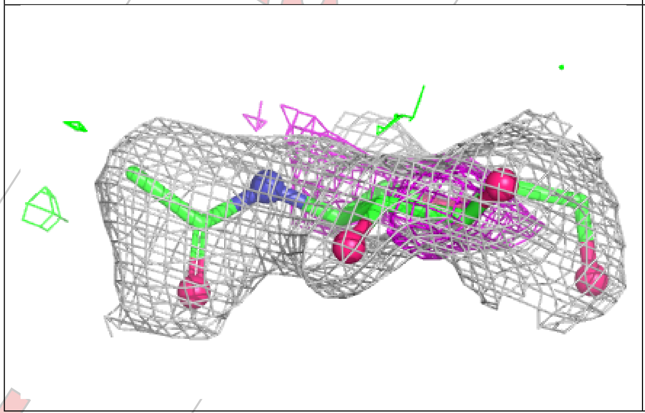

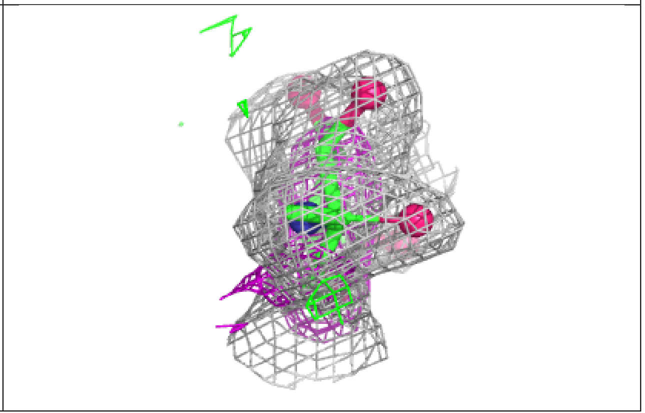

**Electron density around NAG A 301:**

$2mF_o$-$DF_c$ (at 0.7 rmsd) in gray
$mF_o$-$DF_c$ (at 3 rmsd) in purple (negative)
and green (positive)

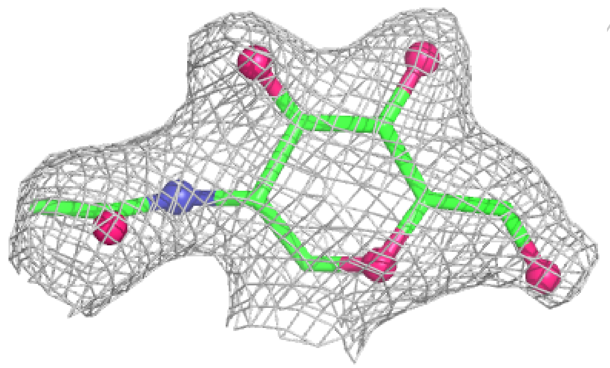

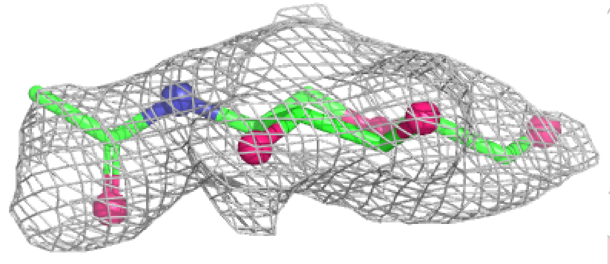

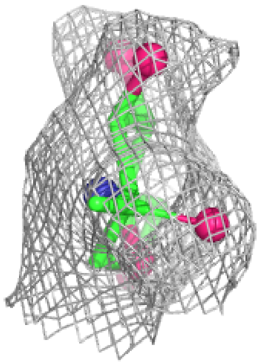

#### 6.5 Other polymers (i)

There are no such residues in this entry.

For Manuscript Review

### Full wwPDB X-ray Structure Validation Report (i)

Mar 21, 2025 – 02:40 PM EDT

| | | |
|---|---|---|
| PDB ID | : | 9NV6 |
| Title | : | Crystal structure of NP101 TCR in complex with NP366-H-2Db |
| Deposited on | : | 2025-03-20 |
| Resolution | : | 2.18 Å(reported) |

**This wwPDB validation report is for manuscript review**

This is a Full wwPDB X-ray Structure Validation Report.

This report is produced by the wwPDB biocuration pipeline after annotation of the structure.

We welcome your comments at **
A user guide is available at
https://www.wwpdb.org/validation/2017/XrayValidationReportHelp
with specific help available everywhere you see the (i) symbol.

The types of validation reports are described at
http://www.wwpdb.org/validation/2017/FAQs#types.

The following versions of software and data (see references (i)) were used in the production of this report:

| | | |
|---|---|---|
| MolProbity | : | 4.02b-467 |
| Mogul | : | 2022.3.0, CSD as543be (2022) |
| Xtriage (Phenix) | : | 1.21 |
| EDS | : | 3.0 |
| buster-report | : | 1.1.7 (2018) |
| Percentile statistics | : | 20231227.v01 (using entries in the PDB archive December 27th 2023) |
| CCP4 | : | 9.0.004 (Gargrove) |
| Density-Fitness | : | 1.0.11 |
| Ideal geometry (proteins) | : | Engh & Huber (2001) |
| Ideal geometry (DNA, RNA) | : | Parkinson et al. (1996) |

For Manuscript Review

### 1 Overall quality at a glance (i)

The following experimental techniques were used to determine the structure:
*X-RAY DIFFRACTION*

The reported resolution of this entry is 2.18 Å.

Percentile scores (ranging between 0-100) for global validation metrics of the entry are shown in the following graphic. The table shows the number of entries on which the scores are based.

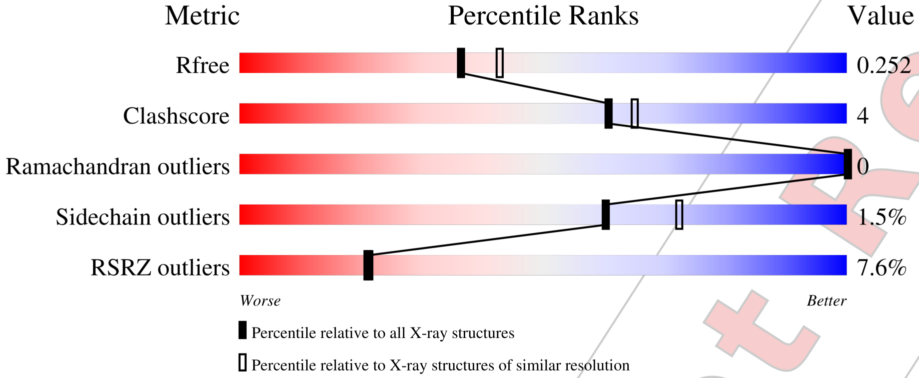


| Metric | Whole archive (#Entries) | Similar resolution (#Entries, resolution range(Å)) |
|---|---|---|
| $R_{free}$ | 164625 | 8336 (2.20-2.16) |
| Clashscore | 180529 | 9404 (2.20-2.16) |
| Ramachandran outliers | 177936 | 9297 (2.20-2.16) |
| Sidechain outliers | 177891 | 9297 (2.20-2.16) |
| RSRZ outliers | 164620 | 8337 (2.20-2.16) |

The table below summarises the geometric issues observed across the polymeric chains and their fit to the electron density. The red, orange, yellow and green segments of the lower bar indicate the fraction of residues that contain outliers for >=3, 2, 1 and 0 types of geometric quality criteria respectively. A grey segment represents the fraction of residues that are not modelled. The numeric value for each fraction is indicated below the corresponding segment, with a dot representing fractions <=5% The upper red bar (where present) indicates the fraction of residues that have poor fit to the electron density. The numeric value is given above the bar.

| Mol | Chain | Length | Quality of chain |
|---|---|---|---|
| 1 | A | 223 | 13% / 82% 11% 7% |
| 2 | B | 254 | 10% / 79% 15% • 5% |
| 3 | C | 281 | 2% / 93% 5% • |
| 4 | D | 100 | 3% / 85% 13% • |

*Continued on next page...*



For Manuscript Review

*Continued from previous page...*

| Mol | Chain | Length | Quality of chain |
|---|---|---|---|
| 5 | P | 9 | 89% 11% |

For Manuscript Review

#### 2 Entry composition (i)

There are 8 unique types of molecules in this entry. The entry contains 6782 atoms, of which 0 are hydrogens and 0 are deuteriums.

In the tables below, the ZeroOcc column contains the number of atoms modelled with zero occupancy, the AltConf column contains the number of residues with at least one atom in alternate conformation and the Trace column contains the number of residues modelled with at most 2 atoms.

- Molecule 1 is a protein called NP101 TCR alpha-chain.

| Mol | Chain | Residues | Atoms | | | | | ZeroOcc | AltConf | Trace |
|---|---|---|---|---|---|---|---|---|---|---|
| | | | Total | C | N | O | S | | | |
| 1 | A | 208 | 1595 | 1006 | 256 | 323 | 10 | 0 | 0 | 0 |

- Molecule 2 is a protein called NP101 TCR beta-chain.

| Mol | Chain | Residues | Atoms | | | | | ZeroOcc | AltConf | Trace |
|---|---|---|---|---|---|---|---|---|---|---|
| | | | Total | C | N | O | S | | | |
| 2 | B | 241 | 1899 | 1213 | 329 | 349 | 8 | 0 | 0 | 0 |

- Molecule 3 is a protein called H-2 class I histocompatibility antigen, D-B alpha chain.

| Mol | Chain | Residues | Atoms | | | | | ZeroOcc | AltConf | Trace |
|---|---|---|---|---|---|---|---|---|---|---|
| | | | Total | C | N | O | S | | | |
| 3 | C | 276 | 2264 | 1430 | 400 | 425 | 9 | 0 | 0 | 0 |

There is a discrepancy between the modelled and reference sequences:

| Chain | Residue | Modelled | Actual | Comment | Reference |
|---|---|---|---|---|---|
| C | 0 | MET | - | initiating methionine | UNP P01899 |

- Molecule 4 is a protein called Beta-2-microglobulin.

| Mol | Chain | Residues | Atoms | | | | | ZeroOcc | AltConf | Trace |
|---|---|---|---|---|---|---|---|---|---|---|
| | | | Total | C | N | O | S | | | |
| 4 | D | 98 | 809 | 515 | 136 | 151 | 7 | 0 | 0 | 0 |

There are 2 discrepancies between the modelled and reference sequences:

| Chain | Residue | Modelled | Actual | Comment | Reference |
|---|---|---|---|---|---|
| D | 0 | MET | - | initiating methionine | UNP P01887 |
| D | 85 | ASP | ALA | conflict | UNP P01887 |

For Manuscript Review

- Molecule 5 is a protein called Nucleoprotein peptide.

| Mol | Chain | Residues | Atoms | ZeroOcc | AltConf | Trace |
|---|---|---|---|---|---|---|
| 5 | P | 9 | Total C N O S<br>69 38 11 18 2 | 0 | 0 | 0 |

- Molecule 6 is 2-acetamido-2-deoxy-beta-D-glucopyranose (three-letter code: NAG) (formula: $C_8H_{15}NO_6$) (labeled as "Ligand of Interest" by depositor).

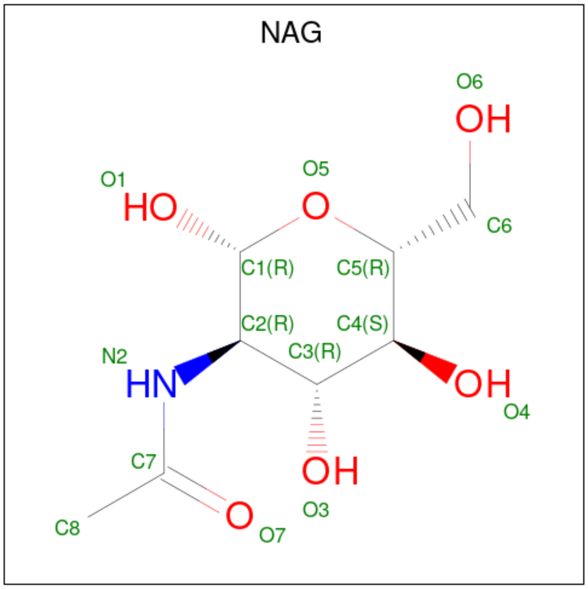


| Mol | Chain | Residues | Atoms | ZeroOcc | AltConf |
|---|---|---|---|---|---|
| 6 | A | 1 | Total C N O<br>14 8 1 5 | 0 | 0 |
| 6 | B | 1 | Total C N O<br>14 8 1 5 | 0 | 0 |
| 6 | B | 1 | Total C N O<br>14 8 1 5 | 0 | 0 |

- Molecule 7 is IODIDE ION (three-letter code: IOD) (formula: I).

| Mol | Chain | Residues | Atoms | ZeroOcc | AltConf |
|---|---|---|---|---|---|
| 7 | A | 3 | Total I<br>3 3 | 0 | 0 |
| 7 | B | 2 | Total I<br>2 2 | 0 | 0 |
| 7 | C | 7 | Total I<br>7 7 | 0 | 0 |
| 7 | D | 4 | Total I<br>4 4 | 0 | 0 |

For Manuscript Review

- Molecule 8 is water.

| Mol | Chain | Residues | Atoms | ZeroOcc | AltConf |
|---|---|---|---|---|---|
| 8 | A | 19 | Total O<br>19 19 | 0 | 0 |
| 8 | B | 10 | Total O<br>10 10 | 0 | 0 |
| 8 | C | 40 | Total O<br>40 40 | 0 | 0 |
| 8 | D | 14 | Total O<br>14 14 | 0 | 0 |
| 8 | P | 5 | Total O<br>5 5 | 0 | 0 |

For Manuscript Review

### 3 Residue-property plots ⓘ

These plots are drawn for all protein, RNA, DNA and oligosaccharide chains in the entry. The first graphic for a chain summarises the proportions of the various outlier classes displayed in the second graphic. The second graphic shows the sequence view annotated by issues in geometry and electron density. Residues are color-coded according to the number of geometric quality criteria for which they contain at least one outlier: green = 0, yellow = 1, orange = 2 and red = 3 or more. A red dot above a residue indicates a poor fit to the electron density (RSRZ > 2). Stretches of 2 or more consecutive residues without any outlier are shown as a green connector. Residues present in the sample, but not in the model, are shown in grey.

- Molecule 1: NP101 TCR alpha-chain

Chain A:

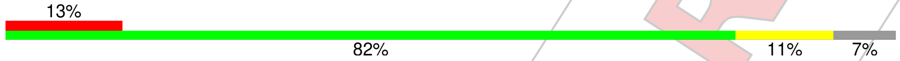


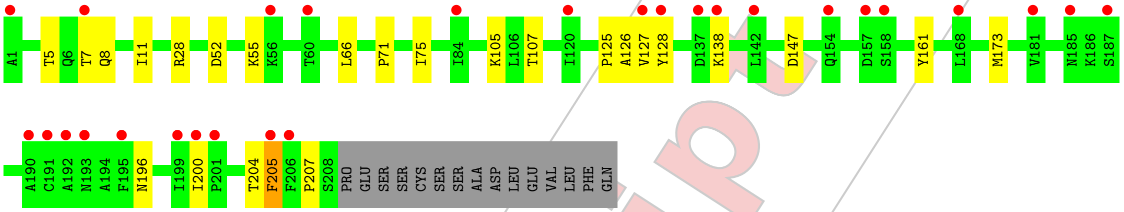


- Molecule 2: NP101 TCR beta-chain

Chain B:

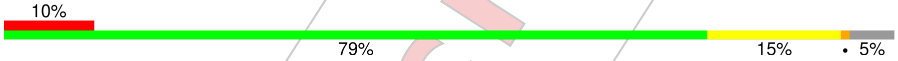


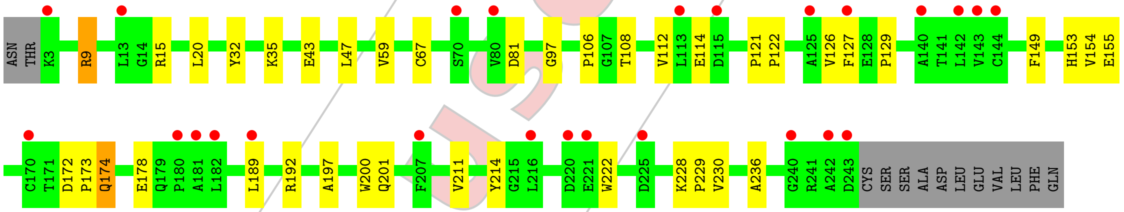


- Molecule 3: H-2 class I histocompatibility antigen, D-B alpha chain

Chain C:

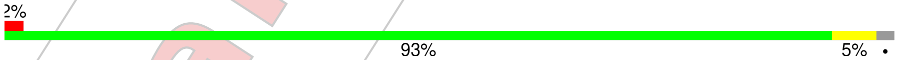


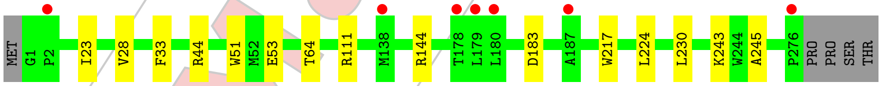


- Molecule 4: Beta-2-microglobulin

Chain D:

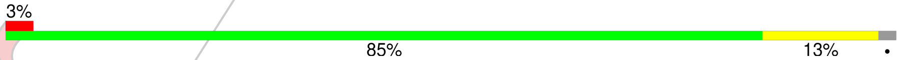


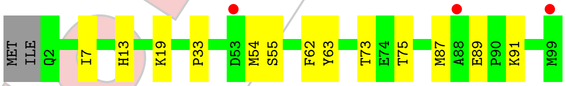


- Molecule 5: Nucleoprotein peptide

For Manuscript Review

Chain P: 89% 11%

A1 M6 M9

For Manuscript Review

### 4 Data and refinement statistics (i)

| Property | Value | Source |
|---|---|---|
| Space group | P 1 | Depositor |
| Cell constants<br>a, b, c, $\alpha$, $\beta$, $\gamma$ | 51.10Å 53.59Å 101.49Å<br>79.28° 88.37° 65.64° | Depositor |
| Resolution (Å) | 47.91 – 2.18<br>47.91 – 2.18 | Depositor<br>EDS |
| % Data completeness<br>(in resolution range) | 95.5 (47.91-2.18)<br>95.5 (47.91-2.18) | Depositor<br>EDS |
| $R_{merge}$ | 0.06 | Depositor |
| $R_{sym}$ | (Not available) | Depositor |
| $< I/\sigma(I) >$ [1] | 1.32 (at 2.18Å) | Xtriage |
| Refinement program | PHENIX 1.21.2_5419 | Depositor |
| R, $R_{free}$ | 0.199 , 0.251<br>0.199 , 0.252 | Depositor<br>DCC |
| $R_{free}$ test set | 2302 reflections (4.56%) | wwPDB-VP |
| Wilson B-factor (Å$^2$) | 48.7 | Xtriage |
| Anisotropy | 0.162 | Xtriage |
| Bulk solvent $k_{sol}$(e/Å$^3$), $B_{sol}$(Å$^2$) | 0.29 , 45.2 | EDS |
| L-test for twinning[2] | $< \|L\| > = 0.49$, $< L^2 > = 0.32$ | Xtriage |
| Estimated twinning fraction | No twinning to report. | Xtriage |
| $F_o$,$F_c$ correlation | 0.95 | EDS |
| Total number of atoms | 6782 | wwPDB-VP |
| Average B, all atoms (Å$^2$) | 74.0 | wwPDB-VP |

Xtriage's analysis on translational NCS is as follows: *The largest off-origin peak in the Patterson function is 5.76% of the height of the origin peak. No significant pseudotranslation is detected.*

[1] Intensities estimated from amplitudes.

[2] Theoretical values of $< |L| >$, $< L^2 >$ for acentric reflections are 0.5, 0.333 respectively for untwinned datasets, and 0.375, 0.2 for perfectly twinned datasets.

For Manuscript Review

### 5 Model quality (i)

#### 5.1 Standard geometry (i)

Bond lengths and bond angles in the following residue types are not validated in this section: NAG, IOD

The Z score for a bond length (or angle) is the number of standard deviations the observed value is removed from the expected value. A bond length (or angle) with $|Z| > 5$ is considered an outlier worth inspection. RMSZ is the root-mean-square of all Z scores of the bond lengths (or angles).

| Mol | Chain | Bond lengths | | Bond angles | |
|---|---|---|---|---|---|
| | | RMSZ | #$\|Z\|$ >5 | RMSZ | #$\|Z\|$ >5 |
| 1 | A | 0.28 | 0/1627 | 0.49 | 0/2207 |
| 2 | B | 0.30 | 0/1954 | 0.52 | 0/2664 |
| 3 | C | 0.28 | 0/2331 | 0.53 | 0/3166 |
| 4 | D | 0.30 | 0/835 | 0.51 | 0/1133 |
| 5 | P | 0.39 | 0/68 | 0.58 | 0/88 |
| All | All | 0.29 | 0/6815 | 0.52 | 0/9258 |

There are no bond length outliers.

There are no bond angle outliers.

There are no chirality outliers.

There are no planarity outliers.

#### 5.2 Too-close contacts (i)

In the following table, the Non-H and H(model) columns list the number of non-hydrogen atoms and hydrogen atoms in the chain respectively. The H(added) column lists the number of hydrogen atoms added and optimized by MolProbity. The Clashes column lists the number of clashes within the asymmetric unit, whereas Symm-Clashes lists symmetry-related clashes.

| Mol | Chain | Non-H | H(model) | H(added) | Clashes | Symm-Clashes |
|---|---|---|---|---|---|---|
| 1 | A | 1595 | 0 | 1502 | 15 | 0 |
| 2 | B | 1899 | 0 | 1812 | 22 | 0 |
| 3 | C | 2264 | 0 | 2136 | 9 | 0 |
| 4 | D | 809 | 0 | 771 | 9 | 0 |
| 5 | P | 69 | 0 | 61 | 1 | 0 |
| 6 | A | 14 | 0 | 13 | 0 | 0 |
| 6 | B | 28 | 0 | 26 | 0 | 0 |
| 7 | A | 3 | 0 | 0 | 1 | 0 |

*Continued on next page...*

For Manuscript Review

*Continued from previous page...*

| Mol | Chain | Non-H | H(model) | H(added) | Clashes | Symm-Clashes |
|---|---|---|---|---|---|---|
| 7 | B | 2 | 0 | 0 | 0 | 0 |
| 7 | C | 7 | 0 | 0 | 2 | 0 |
| 7 | D | 4 | 0 | 0 | 0 | 0 |
| 8 | A | 19 | 0 | 0 | 0 | 0 |
| 8 | B | 10 | 0 | 0 | 0 | 0 |
| 8 | C | 40 | 0 | 0 | 1 | 0 |
| 8 | D | 14 | 0 | 0 | 0 | 0 |
| 8 | P | 5 | 0 | 0 | 0 | 0 |
| All | All | 6782 | 0 | 6321 | 53 | 0 |

The all-atom clashscore is defined as the number of clashes found per 1000 atoms (including hydrogen atoms). The all-atom clashscore for this structure is 4.

All (53) close contacts within the same asymmetric unit are listed below, sorted by their clash magnitude.

| Atom-1 | Atom-2 | Interatomic distance (Å) | Clash overlap (Å) |
|---|---|---|---|
| 1:A:71:PRO:HG2 | 7:A:302:IOD:I | 2.56 | 0.74 |
| 1:A:196:ASN:HA | 1:A:200:ILE:HD11 | 1.71 | 0.71 |
| 4:D:7:ILE:HD12 | 4:D:91:LYS:HD2 | 1.76 | 0.67 |
| 2:B:153:HIS:HB3 | 2:B:214:TYR:HB2 | 1.77 | 0.66 |
| 7:C:304:IOD:I | 8:C:425:HOH:O | 2.84 | 0.66 |
| 3:C:51:TRP:HB3 | 7:C:306:IOD:I | 2.66 | 0.65 |
| 2:B:121:PRO:HD3 | 2:B:229:PRO:HB3 | 1.81 | 0.62 |
| 3:C:230:LEU:HD11 | 3:C:243:LYS:HE3 | 1.83 | 0.60 |
| 3:C:230:LEU:HD13 | 3:C:245:ALA:HB2 | 1.89 | 0.54 |
| 1:A:52:ASP:HB2 | 1:A:55:LYS:HG3 | 1.89 | 0.53 |
| 1:A:8:GLN:HE21 | 1:A:11:ILE:HG13 | 1.73 | 0.53 |
| 1:A:126:ALA:HB1 | 1:A:128:TYR:HE2 | 1.73 | 0.53 |
| 4:D:73:THR:HG22 | 4:D:75:THR:H | 1.72 | 0.52 |
| 2:B:9:ARG:CZ | 2:B:106:PRO:HB2 | 2.39 | 0.52 |
| 2:B:20:LEU:HD22 | 2:B:108:THR:HG21 | 1.91 | 0.52 |
| 2:B:126:VAL:HG23 | 2:B:236:ALA:HB3 | 1.92 | 0.52 |
| 1:A:5:THR:HG22 | 1:A:7:THR:HG23 | 1.93 | 0.51 |
| 4:D:13:HIS:HB2 | 4:D:19:LYS:NZ | 2.26 | 0.51 |
| 2:B:173:PRO:HB2 | 2:B:174:GLN:OE1 | 2.11 | 0.50 |
| 2:B:228:LYS:HD3 | 2:B:230:VAL:HG13 | 1.92 | 0.50 |
| 3:C:217:TRP:HB2 | 3:C:224:LEU:HD11 | 1.93 | 0.50 |
| 4:D:87:MET:C | 4:D:89:GLU:H | 2.16 | 0.49 |
| 2:B:222:TRP:HB2 | 2:B:228:LYS:HE3 | 1.94 | 0.49 |
| 3:C:144:ARG:HH11 | 3:C:144:ARG:HG2 | 1.78 | 0.48 |
| 1:A:105:LYS:HE3 | 1:A:107:THR:OG1 | 2.12 | 0.48 |

*Continued on next page...*

For Manuscript Review

*Continued from previous page...*

| Atom-1 | Atom-2 | Interatomic distance (Å) | Clash overlap (Å) |
|---|---|---|---|
| 2:B:197:ALA:O | 2:B:201:GLN:HG2 | 2.13 | 0.48 |
| 2:B:114:GLU:H | 2:B:114:GLU:CD | 2.17 | 0.48 |
| 1:A:66:LEU:HD13 | 1:A:75:ILE:HD11 | 1.95 | 0.48 |
| 1:A:126:ALA:HB1 | 1:A:128:TYR:CE2 | 2.48 | 0.48 |
| 2:B:35:LYS:O | 2:B:43:GLU:HG2 | 2.14 | 0.47 |
| 2:B:15:ARG:HH11 | 2:B:15:ARG:HA | 1.79 | 0.46 |
| 2:B:122:PRO:HB3 | 2:B:149:PHE:HB3 | 1.97 | 0.46 |
| 4:D:33:PRO:HB3 | 4:D:62:PHE:CE2 | 2.50 | 0.46 |
| 4:D:89:GLU:HG2 | 4:D:89:GLU:O | 2.16 | 0.46 |
| 2:B:172:ASP:HB2 | 2:B:189:LEU:CD1 | 2.47 | 0.45 |
| 3:C:23:ILE:HD12 | 4:D:54:MET:SD | 2.58 | 0.44 |
| 2:B:129:PRO:HD2 | 2:B:200:TRP:CZ2 | 2.52 | 0.44 |
| 1:A:161:TYR:CE2 | 2:B:178:GLU:HA | 2.52 | 0.44 |
| 2:B:155:GLU:HG3 | 2:B:214:TYR:HE2 | 1.83 | 0.44 |
| 3:C:53:GLU:H | 3:C:53:GLU:HG2 | 1.61 | 0.44 |
| 2:B:32:TYR:CE2 | 2:B:47:LEU:HD13 | 2.53 | 0.43 |
| 1:A:138:LYS:HA | 1:A:138:LYS:HD2 | 1.79 | 0.43 |
| 2:B:81:ASP:O | 2:B:112:VAL:HG21 | 2.19 | 0.43 |
| 1:A:127:VAL:O | 1:A:207:PRO:HD2 | 2.18 | 0.43 |
| 1:A:138:LYS:HB3 | 2:B:127:PHE:HE2 | 1.84 | 0.42 |
| 2:B:154:VAL:HG13 | 2:B:211:VAL:HG13 | 2.01 | 0.42 |
| 3:C:28:VAL:HG23 | 3:C:33:PHE:CE1 | 2.55 | 0.41 |
| 1:A:126:ALA:CB | 1:A:205:PHE:HB3 | 2.50 | 0.41 |
| 2:B:97:GLY:HA3 | 5:P:6:MET:HG3 | 2.03 | 0.41 |
| 1:A:125:PRO:O | 1:A:204:THR:HA | 2.21 | 0.41 |
| 4:D:33:PRO:HB3 | 4:D:62:PHE:CD2 | 2.55 | 0.41 |
| 4:D:55:SER:HB2 | 4:D:63:TYR:CZ | 2.55 | 0.41 |
| 3:C:44:ARG:HG3 | 3:C:64:THR:OG1 | 2.22 | 0.40 |

There are no symmetry-related clashes.

#### 5.3 Torsion angles (i)

##### 5.3.1 Protein backbone (i)

In the following table, the Percentiles column shows the percent Ramachandran outliers of the chain as a percentile score with respect to all X-ray entries followed by that with respect to entries of similar resolution.

The Analysed column shows the number of residues for which the backbone conformation was analysed, and the total number of residues.

For Manuscript Review

| Mol | Chain | Analysed | Favoured | Allowed | Outliers | Percentiles | |
|---|---|---|---|---|---|---|---|
| 1 | A | 206/223 (92%) | 196 (95%) | 10 (5%) | 0 | 100 | 100 |
| 2 | B | 239/254 (94%) | 230 (96%) | 9 (4%) | 0 | 100 | 100 |
| 3 | C | 274/281 (98%) | 268 (98%) | 6 (2%) | 0 | 100 | 100 |
| 4 | D | 96/100 (96%) | 93 (97%) | 3 (3%) | 0 | 100 | 100 |
| 5 | P | 7/9 (78%) | 6 (86%) | 1 (14%) | 0 | 100 | 100 |
| All | All | 822/867 (95%) | 793 (96%) | 29 (4%) | 0 | 100 | 100 |

There are no Ramachandran outliers to report.

##### 5.3.2 Protein sidechains ⓘ

In the following table, the Percentiles column shows the percent sidechain outliers of the chain as a percentile score with respect to all X-ray entries followed by that with respect to entries of similar resolution.

The Analysed column shows the number of residues for which the sidechain conformation was analysed, and the total number of residues.

| Mol | Chain | Analysed | Rotameric | Outliers | Percentiles | |
|---|---|---|---|---|---|---|
| 1 | A | 176/201 (88%) | 172 (98%) | 4 (2%) | 45 | 56 |
| 2 | B | 202/225 (90%) | 197 (98%) | 5 (2%) | 42 | 53 |
| 3 | C | 234/239 (98%) | 232 (99%) | 2 (1%) | 75 | 85 |
| 4 | D | 92/95 (97%) | 92 (100%) | 0 | 100 | 100 |
| 5 | P | 8/8 (100%) | 8 (100%) | 0 | 100 | 100 |
| All | All | 712/768 (93%) | 701 (98%) | 11 (2%) | 60 | 72 |

All (11) residues with a non-rotameric sidechain are listed below:

| Mol | Chain | Res | Type |
|---|---|---|---|
| 1 | A | 28 | ARG |
| 1 | A | 147 | ASP |
| 1 | A | 173 | MET |
| 1 | A | 205 | PHE |
| 2 | B | 9 | ARG |
| 2 | B | 59 | VAL |
| 2 | B | 67 | CYS |
| 2 | B | 174 | GLN |
| 2 | B | 192 | ARG |
| 3 | C | 111 | ARG |

*Continued on next page...*

For Manuscript Review

*Continued from previous page...*

| Mol | Chain | Res | Type |
|---|---|---|---|
| 3 | C | 183 | ASP |

Sometimes sidechains can be flipped to improve hydrogen bonding and reduce clashes. All (6) such sidechains are listed below:

| Mol | Chain | Res | Type |
|---|---|---|---|
| 1 | A | 63 | HIS |
| 2 | B | 76 | HIS |
| 2 | B | 212 | GLN |
| 3 | C | 86 | ASN |
| 3 | C | 115 | GLN |
| 4 | D | 13 | HIS |

##### 5.3.3 RNA (i)

There are no RNA molecules in this entry.

#### 5.4 Non-standard residues in protein, DNA, RNA chains (i)

There are no non-standard protein/DNA/RNA residues in this entry.

#### 5.5 Carbohydrates (i)

There are no oligosaccharides in this entry.

#### 5.6 Ligand geometry (i)

Of 19 ligands modelled in this entry, 16 are monoatomic - leaving 3 for Mogul analysis.

In the following table, the Counts columns list the number of bonds (or angles) for which Mogul statistics could be retrieved, the number of bonds (or angles) that are observed in the model and the number of bonds (or angles) that are defined in the Chemical Component Dictionary. The Link column lists molecule types, if any, to which the group is linked. The Z score for a bond length (or angle) is the number of standard deviations the observed value is removed from the expected value. A bond length (or angle) with $|Z| > 2$ is considered an outlier worth inspection. RMSZ is the root-mean-square of all Z scores of the bond lengths (or angles).

| Mol | Type | Chain | Res | Link | Bond lengths | | | Bond angles | | |
|---|---|---|---|---|---|---|---|---|---|---|
| | | | | | Counts | RMSZ | $\#\|Z\| > 2$ | Counts | RMSZ | $\#\|Z\| > 2$ |
| 6 | NAG | B | 301 | 2 | 14,14,15 | 0.73 | 0 | 17,19,21 | 0.87 | 0 |
| 6 | NAG | A | 301 | 1 | 14,14,15 | 0.74 | 0 | 17,19,21 | 0.72 | 0 |

For Manuscript Review

| Mol | Type | Chain | Res | Link | Bond lengths | | | Bond angles | | |
|---|---|---|---|---|---|---|---|---|---|---|
| | | | | | Counts | RMSZ | #\|Z\| > 2 | Counts | RMSZ | #\|Z\| > 2 |
| 6 | NAG | B | 302 | 2 | 14,14,15 | 0.79 | 0 | 17,19,21 | 1.81 | 2 (11%) |

In the following table, the Chirals column lists the number of chiral outliers, the number of chiral centers analysed, the number of these observed in the model and the number defined in the Chemical Component Dictionary. Similar counts are reported in the Torsion and Rings columns. '-' means no outliers of that kind were identified.

| Mol | Type | Chain | Res | Link | Chirals | Torsions | Rings |
|---|---|---|---|---|---|---|---|
| 6 | NAG | B | 301 | 2 | - | 2/6/23/26 | 0/1/1/1 |
| 6 | NAG | A | 301 | 1 | - | 0/6/23/26 | 0/1/1/1 |
| 6 | NAG | B | 302 | 2 | - | 1/6/23/26 | 0/1/1/1 |

There are no bond length outliers.

All (2) bond angle outliers are listed below:

| Mol | Chain | Res | Type | Atoms | Z | Observed(°) | Ideal(°) |
|---|---|---|---|---|---|---|---|
| 6 | B | 302 | NAG | C2-N2-C7 | 5.93 | 130.85 | 122.90 |
| 6 | B | 302 | NAG | O7-C7-N2 | 2.47 | 126.34 | 121.98 |

There are no chirality outliers.

All (3) torsion outliers are listed below:

| Mol | Chain | Res | Type | Atoms |
|---|---|---|---|---|
| 6 | B | 302 | NAG | C3-C2-N2-C7 |
| 6 | B | 301 | NAG | C8-C7-N2-C2 |
| 6 | B | 301 | NAG | O7-C7-N2-C2 |

There are no ring outliers.

No monomer is involved in short contacts.

The following is a two-dimensional graphical depiction of Mogul quality analysis of bond lengths, bond angles, torsion angles, and ring geometry for all instances of the Ligand of Interest. In addition, ligands with molecular weight > 250 and outliers as shown on the validation Tables will also be included. For torsion angles, if less then 5% of the Mogul distribution of torsion angles is within 10 degrees of the torsion angle in question, then that torsion angle is considered an outlier. Any bond that is central to one or more torsion angles identified as an outlier by Mogul will be highlighted in the graph. For rings, the root-mean-square deviation (RMSD) between the ring in question and similar rings identified by Mogul is calculated over all ring torsion angles. If the average RMSD is greater than 60 degrees and the minimal RMSD between the ring in question and any Mogul-identified rings is also greater than 60 degrees, then that ring is considered an outlier. The outliers are highlighted in purple. The color gray indicates Mogul did not find sufficient

For Manuscript Review

equivalents in the CSD to analyse the geometry.

**Ligand NAG B 301**

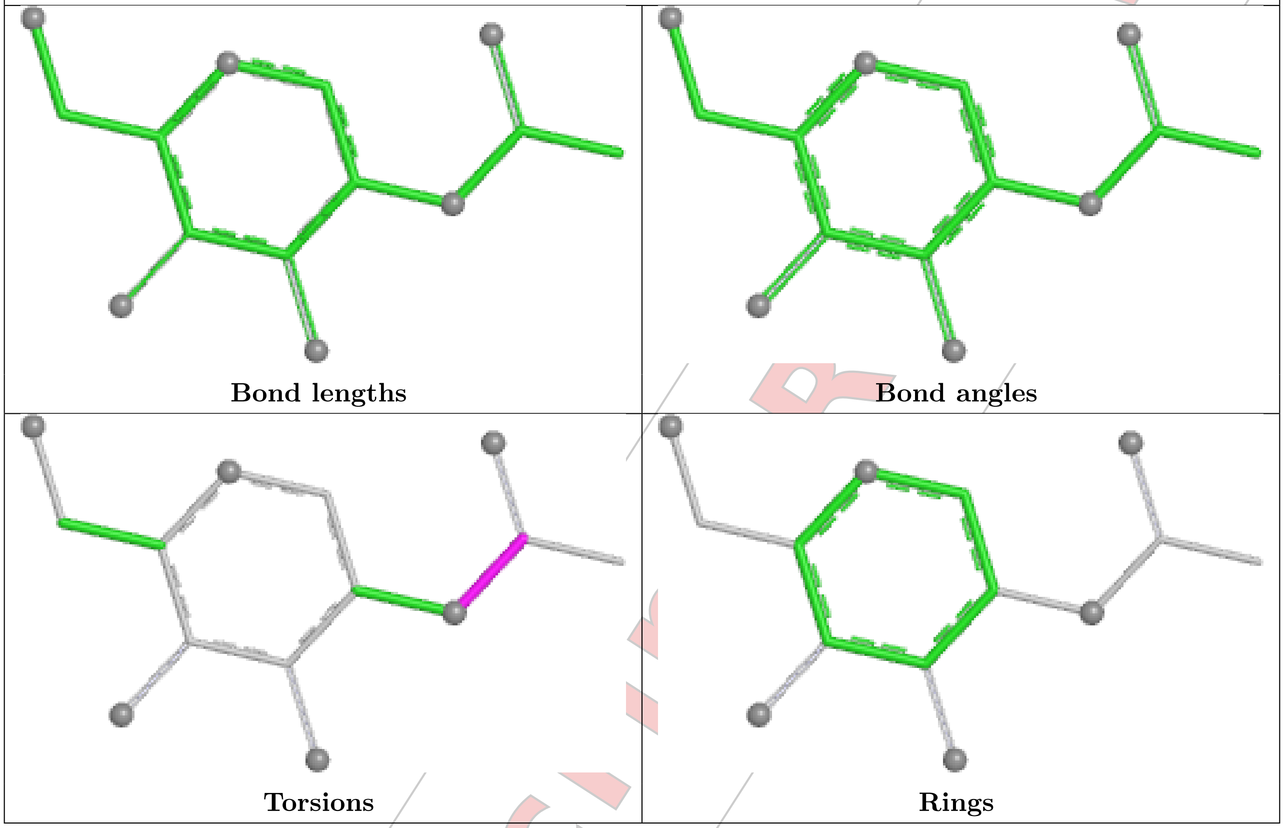


**Ligand NAG A 301**

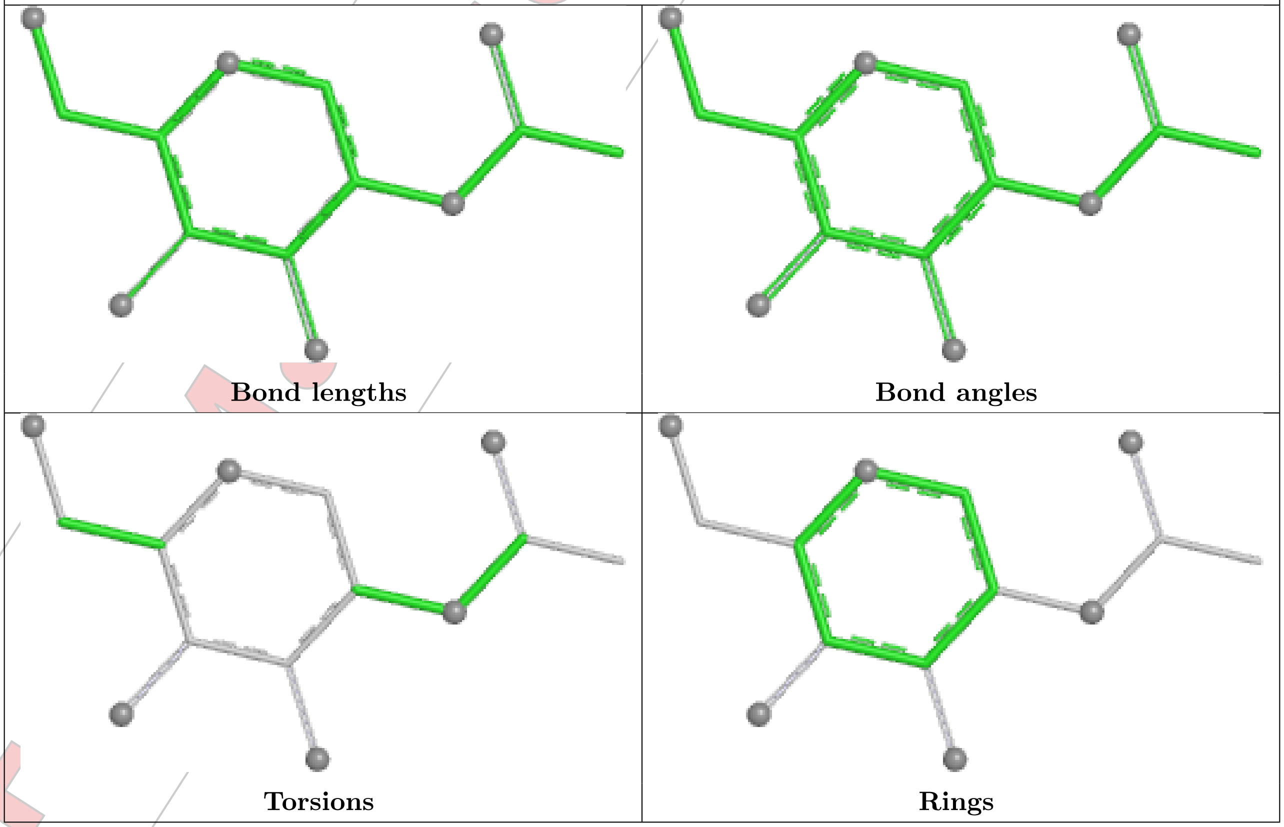

**Ligand NAG B 302**

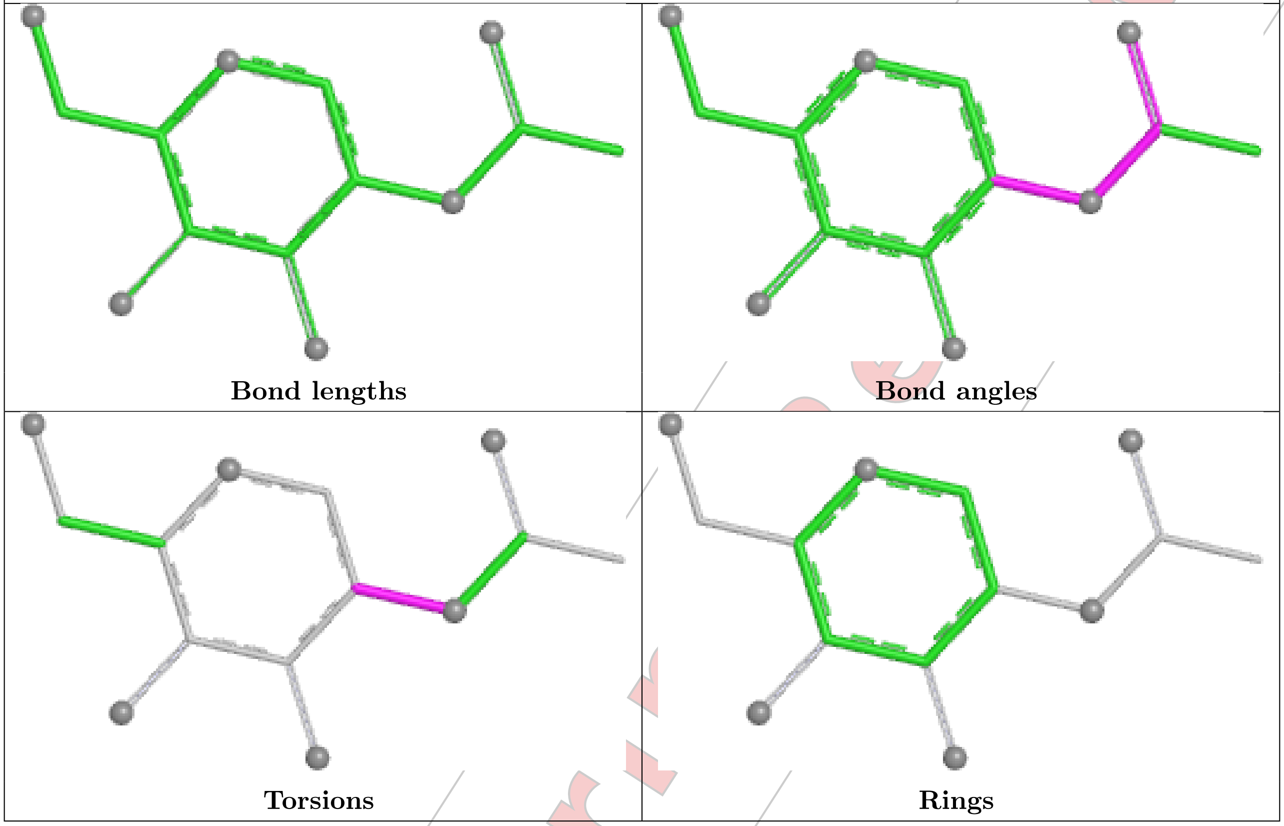


#### 5.7 Other polymers (i)

There are no such residues in this entry.

#### 5.8 Polymer linkage issues (i)

There are no chain breaks in this entry.

For Manuscript Review

### 6 Fit of model and data (i)

#### 6.1 Protein, DNA and RNA chains (i)

In the following table, the column labelled '#RSRZ> 2' contains the number (and percentage) of RSRZ outliers, followed by percent RSRZ outliers for the chain as percentile scores relative to all X-ray entries and entries of similar resolution. The OWAB column contains the minimum, median, $95^{th}$ percentile and maximum values of the occupancy-weighted average B-factor per residue. The column labelled 'Q< 0.9' lists the number of (and percentage) of residues with an average occupancy less than 0.9.

| Mol | Chain | Analysed | <RSRZ> | #RSRZ>2 | | | OWAB(Å$^2$) | Q<0.9 |
|---|---|---|---|---|---|---|---|---|
| 1 | A | 208/223 (93%) | 0.82 | 28 (13%) | 8 | 7 | 36, 80, 153, 165 | 0 |
| 2 | B | 241/254 (94%) | 0.86 | 25 (10%) | 13 | 12 | 40, 88, 143, 179 | 0 |
| 3 | C | 276/281 (98%) | 0.35 | 7 (2%) | 58 | 57 | 33, 58, 96, 123 | 0 |
| 4 | D | 98/100 (98%) | 0.53 | 3 (3%) | 51 | 51 | 42, 62, 98, 124 | 0 |
| 5 | P | 9/9 (100%) | -0.28 | 0 | 100 | 100 | 38, 44, 53, 56 | 0 |
| All | All | 832/867 (95%) | 0.63 | 63 (7%) | 21 | 21 | 33, 68, 138, 179 | 0 |

All (63) RSRZ outliers are listed below:

| Mol | Chain | Res | Type | RSRZ |
|---|---|---|---|---|
| 2 | B | 140 | ALA | 3.8 |
| 4 | D | 88 | ALA | 3.8 |
| 2 | B | 142 | LEU | 3.7 |
| 2 | B | 125 | ALA | 3.7 |
| 1 | A | 127 | VAL | 3.5 |
| 3 | C | 2 | PRO | 3.4 |
| 3 | C | 179 | LEU | 3.3 |
| 1 | A | 1 | ALA | 3.2 |
| 1 | A | 193 | ASN | 3.1 |
| 1 | A | 142 | LEU | 3.1 |
| 2 | B | 225 | ASP | 3.0 |
| 3 | C | 178 | THR | 2.9 |
| 2 | B | 181 | ALA | 2.8 |
| 2 | B | 240 | GLY | 2.8 |
| 1 | A | 199 | ILE | 2.8 |
| 4 | D | 99 | MET | 2.8 |
| 2 | B | 127 | PHE | 2.7 |
| 1 | A | 181 | VAL | 2.7 |
| 1 | A | 187 | SER | 2.7 |
| 1 | A | 192 | ALA | 2.7 |

*Continued on next page...*

For Manuscript Review

*Continued from previous page...*

| Mol | Chain | Res | Type | RSRZ |
|---|---|---|---|---|
| 1 | A | 157 | ASP | 2.7 |
| 2 | B | 220 | ASP | 2.7 |
| 2 | B | 143 | VAL | 2.7 |
| 2 | B | 3 | LYS | 2.6 |
| 1 | A | 120 | ILE | 2.6 |
| 1 | A | 185 | ASN | 2.6 |
| 1 | A | 7 | THR | 2.6 |
| 3 | C | 276 | PRO | 2.6 |
| 1 | A | 205 | PHE | 2.5 |
| 2 | B | 243 | ASP | 2.5 |
| 4 | D | 53 | ASP | 2.5 |
| 1 | A | 154 | GLN | 2.5 |
| 1 | A | 200 | ILE | 2.5 |
| 2 | B | 115 | ASP | 2.5 |
| 1 | A | 201 | PRO | 2.4 |
| 2 | B | 216 | LEU | 2.4 |
| 2 | B | 144 | CYS | 2.4 |
| 1 | A | 60 | THR | 2.4 |
| 1 | A | 128 | TYR | 2.4 |
| 2 | B | 182 | LEU | 2.4 |
| 3 | C | 180 | LEU | 2.4 |
| 1 | A | 191 | CYS | 2.3 |
| 2 | B | 242 | ALA | 2.3 |
| 1 | A | 158 | SER | 2.3 |
| 1 | A | 195 | PHE | 2.3 |
| 2 | B | 189 | LEU | 2.3 |
| 1 | A | 206 | PHE | 2.3 |
| 1 | A | 56 | LYS | 2.2 |
| 2 | B | 113 | LEU | 2.2 |
| 3 | C | 187 | ALA | 2.2 |
| 2 | B | 221 | GLU | 2.2 |
| 2 | B | 207 | PHE | 2.1 |
| 1 | A | 190 | ALA | 2.1 |
| 2 | B | 180 | PRO | 2.1 |
| 3 | C | 138 | MET | 2.1 |
| 1 | A | 137 | ASP | 2.1 |
| 2 | B | 70 | SER | 2.1 |
| 1 | A | 168 | LEU | 2.1 |
| 2 | B | 80 | VAL | 2.0 |
| 2 | B | 13 | LEU | 2.0 |
| 1 | A | 84 | ILE | 2.0 |
| 1 | A | 138 | LYS | 2.0 |

*Continued on next page...*

For Manuscript Review

*Continued from previous page...*

| Mol | Chain | Res | Type | RSRZ |
|---|---|---|---|---|
| 2 | B | 170 | CYS | 2.0 |

#### 6.2 Non-standard residues in protein, DNA, RNA chains (i)

There are no non-standard protein/DNA/RNA residues in this entry.

#### 6.3 Carbohydrates (i)

There are no monosaccharides in this entry.

#### 6.4 Ligands (i)

In the following table, the Atoms column lists the number of modelled atoms in the group and the number defined in the chemical component dictionary. The B-factors column lists the minimum, median, $95^{th}$ percentile and maximum values of B factors of atoms in the group. The column labelled 'Q< 0.9' lists the number of atoms with occupancy less than 0.9.

| Mol | Type | Chain | Res | Atoms | RSCC | RSR | B-factors($Å^2$) | Q<0.9 |
|---|---|---|---|---|---|---|---|---|
| 7 | IOD | D | 102 | 1/1 | 0.40 | 0.34 | 146,146,146,146 | 1 |
| 6 | NAG | B | 302 | 14/15 | 0.64 | 0.16 | 107,115,128,128 | 0 |
| 7 | IOD | B | 304 | 1/1 | 0.72 | 0.12 | 131,131,131,131 | 1 |
| 7 | IOD | C | 304 | 1/1 | 0.75 | 0.28 | 83,83,83,83 | 1 |
| 6 | NAG | B | 301 | 14/15 | 0.83 | 0.12 | 61,69,76,79 | 0 |
| 7 | IOD | D | 103 | 1/1 | 0.84 | 0.11 | 125,125,125,125 | 1 |
| 6 | NAG | A | 301 | 14/15 | 0.90 | 0.09 | 60,72,77,78 | 0 |
| 7 | IOD | C | 307 | 1/1 | 0.92 | 0.10 | 74,74,74,74 | 1 |
| 7 | IOD | C | 306 | 1/1 | 0.94 | 0.07 | 82,82,82,82 | 1 |
| 7 | IOD | A | 304 | 1/1 | 0.94 | 0.08 | 87,87,87,87 | 1 |
| 7 | IOD | C | 301 | 1/1 | 0.96 | 0.07 | 68,68,68,68 | 1 |
| 7 | IOD | A | 302 | 1/1 | 0.96 | 0.05 | 69,69,69,69 | 1 |
| 7 | IOD | D | 104 | 1/1 | 0.96 | 0.08 | 66,66,66,66 | 1 |
| 7 | IOD | B | 303 | 1/1 | 0.97 | 0.05 | 56,56,56,56 | 1 |
| 7 | IOD | C | 305 | 1/1 | 0.98 | 0.04 | 59,59,59,59 | 1 |
| 7 | IOD | C | 302 | 1/1 | 0.98 | 0.04 | 47,47,47,47 | 1 |
| 7 | IOD | A | 303 | 1/1 | 0.98 | 0.04 | 55,55,55,55 | 1 |
| 7 | IOD | D | 101 | 1/1 | 0.99 | 0.02 | 47,47,47,47 | 1 |
| 7 | IOD | C | 303 | 1/1 | 0.99 | 0.05 | 43,43,43,43 | 1 |

The following is a graphical depiction of the model fit to experimental electron density of all instances of the Ligand of Interest. In addition, ligands with molecular weight > 250 and outliers

For Manuscript Review

as shown on the geometry validation Tables will also be included. Each fit is shown from different orientation to approximate a three-dimensional view.

**Electron density around NAG B 302:**

2mF$_o$-DF$_c$ (at 0.7 rmsd) in gray
mF$_o$-DF$_c$ (at 3 rmsd) in purple (negative)
and green (positive)

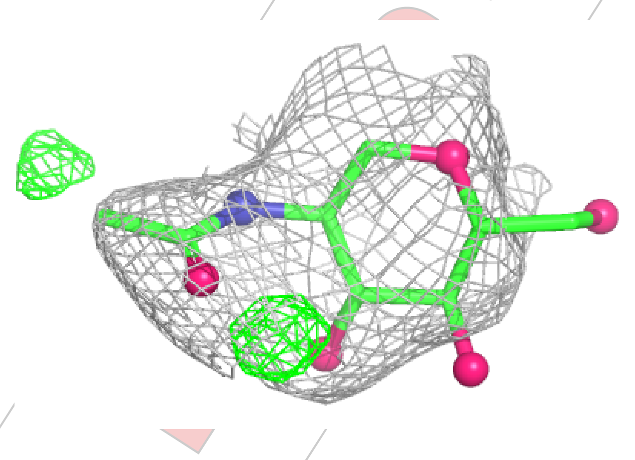

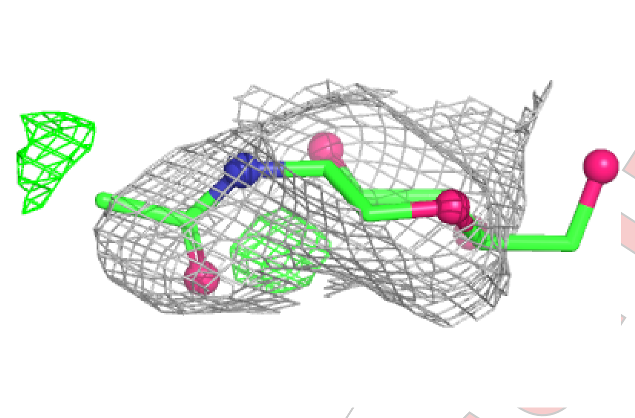

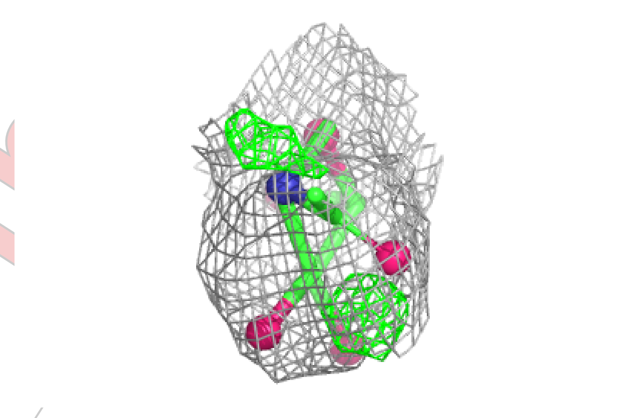

For Manuscript Review

**Electron density around NAG B 301:**

$2mF_o$-$DF_c$ (at 0.7 rmsd) in gray
$mF_o$-$DF_c$ (at 3 rmsd) in purple (negative)
and green (positive)

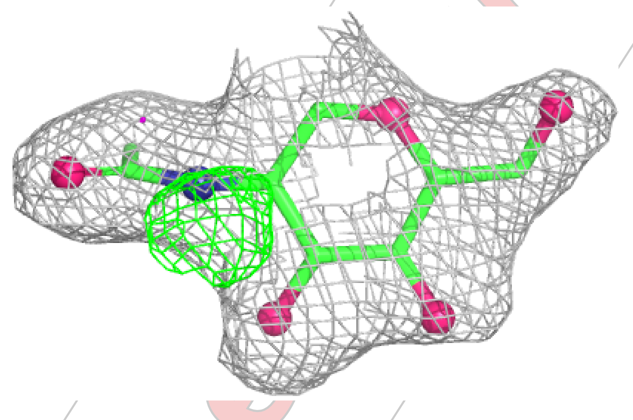

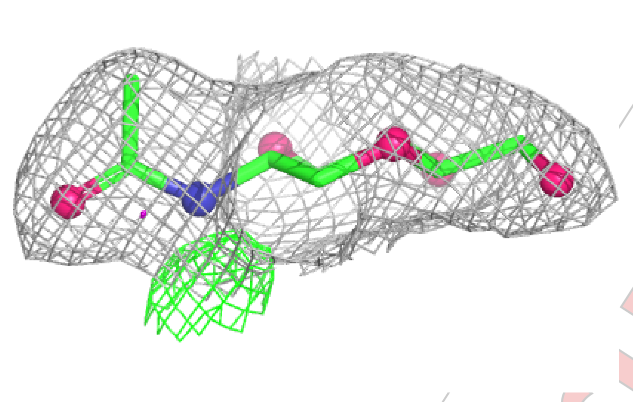

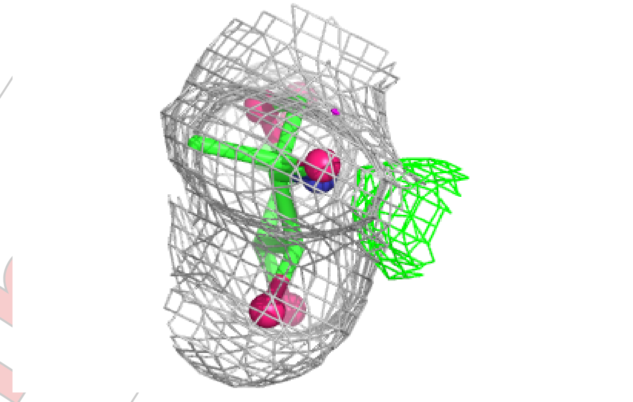

**Electron density around NAG A 301:**

$2mF_o$-$DF_c$ (at 0.7 rmsd) in gray
$mF_o$-$DF_c$ (at 3 rmsd) in purple (negative)
and green (positive)

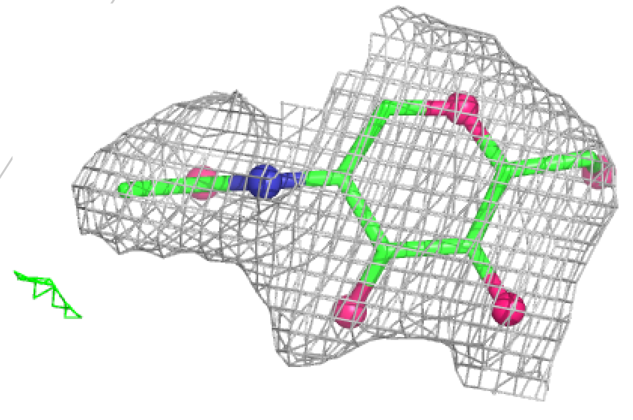

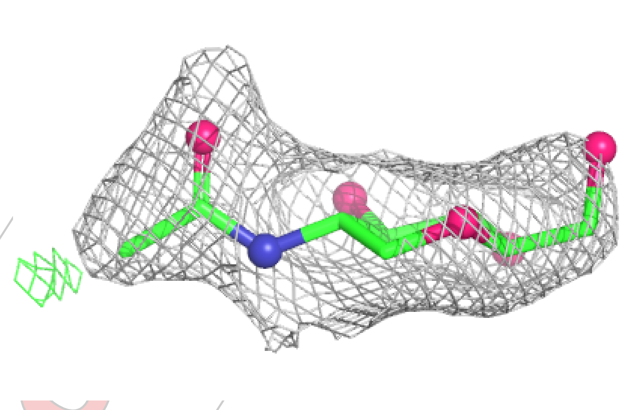

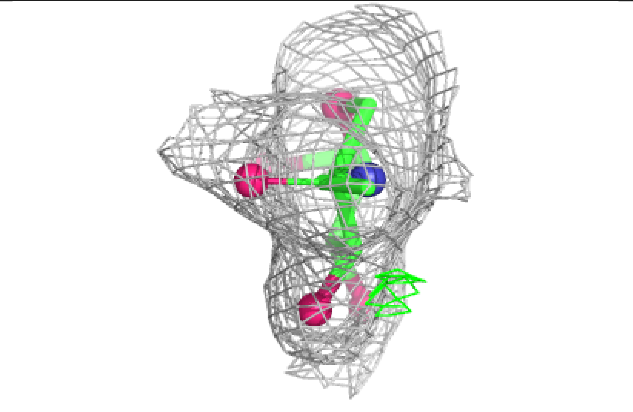

For Manuscript Review

#### 6.5 Other polymers (i)

There are no such residues in this entry.

For Manuscript Review

Full wwPDB X-ray Structure Validation Report (i)

Mar 21, 2025 – 02:39 PM EDT

| | | |
|---|---|---|
| PDB ID | : | 9NVA |
| Title | : | Crystal structure of NP105 TCR in complex with NP366-H-2Db |
| Deposited on | : | 2025-03-20 |
| Resolution | : | 2.78 Å(reported) |

**This wwPDB validation report is for manuscript review**

For Manuscript Review

This is a Full wwPDB X-ray Structure Validation Report.

This report is produced by the wwPDB biocuration pipeline after annotation of the structure.

We welcome your comments at **
A user guide is available at
https://www.wwpdb.org/validation/2017/XrayValidationReportHelp
with specific help available everywhere you see the (i) symbol.

The types of validation reports are described at
http://www.wwpdb.org/validation/2017/FAQs#types.

---

The following versions of software and data (see references (i)) were used in the production of this report:

| | | |
|---|---|---|
| MolProbity | : | 4.02b-467 |
| Mogul | : | 2022.3.0, CSD as543be (2022) |
| Xtriage (Phenix) | : | 1.21 |
| EDS | : | 3.0 |
| buster-report | : | 1.1.7 (2018) |
| Percentile statistics | : | 20231227.v01 (using entries in the PDB archive December 27th 2023) |
| CCP4 | : | 9.0.004 (Gargrove) |
| Density-Fitness | : | 1.0.11 |
| Ideal geometry (proteins) | : | Engh & Huber (2001) |
| Ideal geometry (DNA, RNA) | : | Parkinson et al. (1996) |

### 1 Overall quality at a glance (i)

The following experimental techniques were used to determine the structure:
*X-RAY DIFFRACTION*

The reported resolution of this entry is 2.78 Å.

Percentile scores (ranging between 0-100) for global validation metrics of the entry are shown in the following graphic. The table shows the number of entries on which the scores are based.

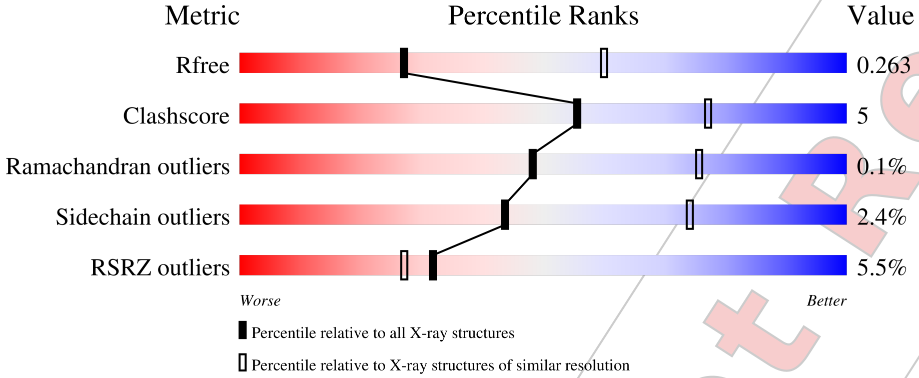


| Metric | Whole archive (#Entries) | Similar resolution (#Entries, resolution range(Å)) |
|---|---|---|
| $R_{free}$ | 164625 | 4924 (2.80-2.76) |
| Clashscore | 180529 | 5458 (2.80-2.76) |
| Ramachandran outliers | 177936 | 5386 (2.80-2.76) |
| Sidechain outliers | 177891 | 5388 (2.80-2.76) |
| RSRZ outliers | 164620 | 4926 (2.80-2.76) |

The table below summarises the geometric issues observed across the polymeric chains and their fit to the electron density. The red, orange, yellow and green segments of the lower bar indicate the fraction of residues that contain outliers for >=3, 2, 1 and 0 types of geometric quality criteria respectively. A grey segment represents the fraction of residues that are not modelled. The numeric value for each fraction is indicated below the corresponding segment, with a dot representing fractions <=5% The upper red bar (where present) indicates the fraction of residues that have poor fit to the electron density. The numeric value is given above the bar.

| Mol | Chain | Length | Quality of chain |
|---|---|---|---|
| 1 | A | 216 | 9% / 81% 13% 6% |
| 2 | B | 257 | 3% / 82% 12% • 5% |
| 3 | C | 281 | 6% / 86% 12% • |
| 4 | D | 100 | 3% / 86% 14% |

*Continued on next page...*



For Manuscript Review

*Continued from previous page...*

| Mol | Chain | Length | Quality of chain |
|---|---|---|---|
| 5 | P | 9 | 100% |

For Manuscript Review

#### 2 Entry composition (i)

There are 9 unique types of molecules in this entry. The entry contains 6834 atoms, of which 0 are hydrogens and 0 are deuteriums.

In the tables below, the ZeroOcc column contains the number of atoms modelled with zero occupancy, the AltConf column contains the number of residues with at least one atom in alternate conformation and the Trace column contains the number of residues modelled with at most 2 atoms.

- Molecule 1 is a protein called NP105 TCR alpha-chain.

| Mol | Chain | Residues | Atoms | | | | | ZeroOcc | AltConf | Trace |
|---|---|---|---|---|---|---|---|---|---|---|
| | | | Total | C | N | O | S | | | |
| 1 | A | 203 | 1581 | 987 | 269 | 315 | 10 | 0 | 1 | 0 |

- Molecule 2 is a protein called NP105 TCR beta-chain.

| Mol | Chain | Residues | Atoms | | | | | ZeroOcc | AltConf | Trace |
|---|---|---|---|---|---|---|---|---|---|---|
| | | | Total | C | N | O | S | | | |
| 2 | B | 245 | 1998 | 1265 | 346 | 379 | 8 | 0 | 3 | 0 |

- Molecule 3 is a protein called H-2 class I histocompatibility antigen, D-B alpha chain.

| Mol | Chain | Residues | Atoms | | | | | ZeroOcc | AltConf | Trace |
|---|---|---|---|---|---|---|---|---|---|---|
| | | | Total | C | N | O | S | | | |
| 3 | C | 277 | 2265 | 1430 | 400 | 426 | 9 | 0 | 2 | 0 |

There is a discrepancy between the modelled and reference sequences:

| Chain | Residue | Modelled | Actual | Comment | Reference |
|---|---|---|---|---|---|
| C | 0 | MET | - | initiating methionine | UNP P01899 |

- Molecule 4 is a protein called Beta-2-microglobulin.

| Mol | Chain | Residues | Atoms | | | | | ZeroOcc | AltConf | Trace |
|---|---|---|---|---|---|---|---|---|---|---|
| | | | Total | C | N | O | S | | | |
| 4 | D | 100 | 838 | 534 | 142 | 154 | 8 | 0 | 1 | 0 |

There are 2 discrepancies between the modelled and reference sequences:

| Chain | Residue | Modelled | Actual | Comment | Reference |
|---|---|---|---|---|---|
| D | 0 | MET | - | initiating methionine | UNP P01887 |
| D | 85 | ASP | ALA | conflict | UNP P01887 |

For Manuscript Review

- Molecule 5 is a protein called Nucleoprotein peptide.

| Mol | Chain | Residues | Atoms | | | | | ZeroOcc | AltConf | Trace |
|---|---|---|---|---|---|---|---|---|---|---|
| | | | Total | C | N | O | S | | | |
| 5 | P | 9 | 69 | 38 | 11 | 18 | 2 | 0 | 0 | 0 |

- Molecule 6 is 2-acetamido-2-deoxy-beta-D-glucopyranose (three-letter code: NAG) (formula: $C_8H_{15}NO_6$) (labeled as "Ligand of Interest" by depositor).

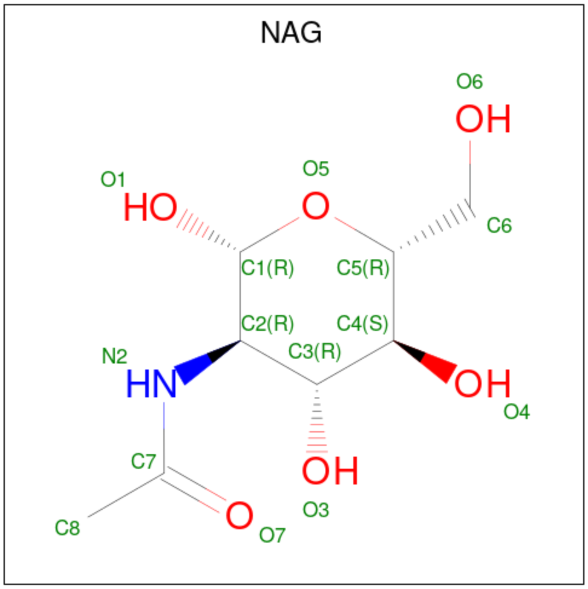


| Mol | Chain | Residues | Atoms | | | | ZeroOcc | AltConf |
|---|---|---|---|---|---|---|---|---|
| | | | Total | C | N | O | | |
| 6 | A | 1 | 14 | 8 | 1 | 5 | 0 | 0 |
| 6 | B | 1 | 14 | 8 | 1 | 5 | 0 | 0 |

- Molecule 7 is GLYCEROL (three-letter code: GOL) (formula: $C_3H_8O_3$).

For Manuscript Review

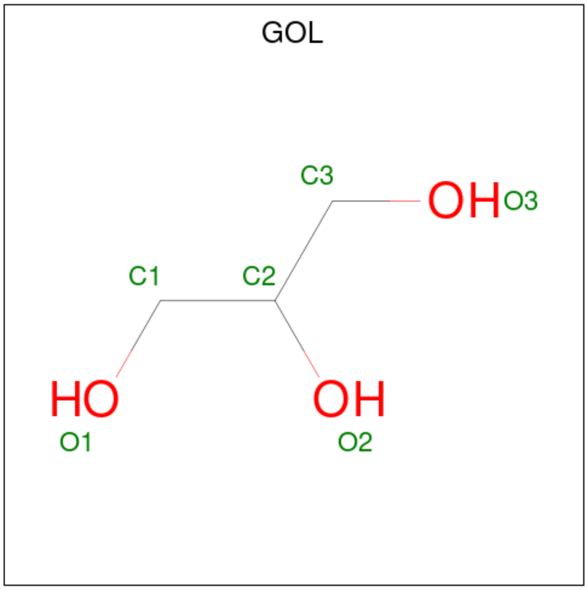


| Mol | Chain | Residues | Atoms | ZeroOcc | AltConf |
|---|---|---|---|---|---|
| 7 | A | 1 | Total C O<br>6 3 3 | 0 | 0 |

- Molecule 8 is CHLORIDE ION (three-letter code: CL) (formula: Cl).

| Mol | Chain | Residues | Atoms | ZeroOcc | AltConf |
|---|---|---|---|---|---|
| 8 | B | 3 | Total Cl<br>3 3 | 0 | 0 |
| 8 | C | 1 | Total Cl<br>1 1 | 0 | 0 |
| 8 | D | 1 | Total Cl<br>1 1 | 0 | 0 |

- Molecule 9 is water.

| Mol | Chain | Residues | Atoms | ZeroOcc | AltConf |
|---|---|---|---|---|---|
| 9 | A | 9 | Total O<br>9 9 | 0 | 0 |
| 9 | B | 14 | Total O<br>14 14 | 0 | 0 |
| 9 | C | 12 | Total O<br>12 12 | 0 | 0 |
| 9 | D | 8 | Total O<br>8 8 | 0 | 0 |
| 9 | P | 1 | Total O<br>1 1 | 0 | 0 |

For Manuscript Review

### 3 Residue-property plots (i)

These plots are drawn for all protein, RNA, DNA and oligosaccharide chains in the entry. The first graphic for a chain summarises the proportions of the various outlier classes displayed in the second graphic. The second graphic shows the sequence view annotated by issues in geometry and electron density. Residues are color-coded according to the number of geometric quality criteria for which they contain at least one outlier: green = 0, yellow = 1, orange = 2 and red = 3 or more. A red dot above a residue indicates a poor fit to the electron density (RSRZ > 2). Stretches of 2 or more consecutive residues without any outlier are shown as a green connector. Residues present in the sample, but not in the model, are shown in grey.

• Molecule 1: NP105 TCR alpha-chain

Chain A:

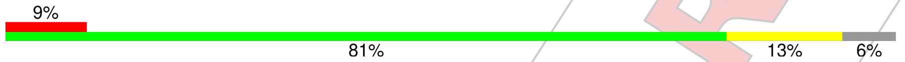


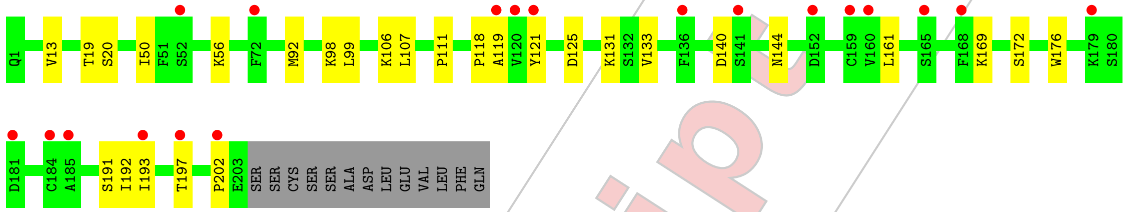


• Molecule 2: NP105 TCR beta-chain

Chain B:

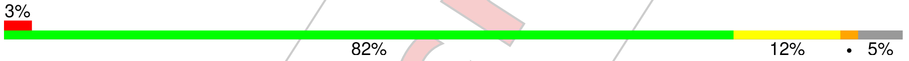


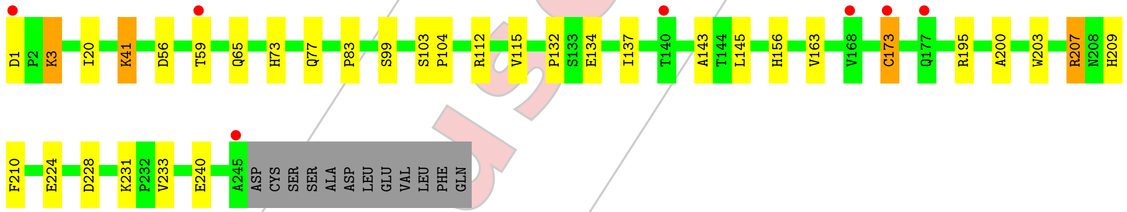


• Molecule 3: H-2 class I histocompatibility antigen, D-B alpha chain

Chain C:

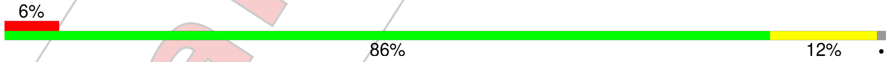


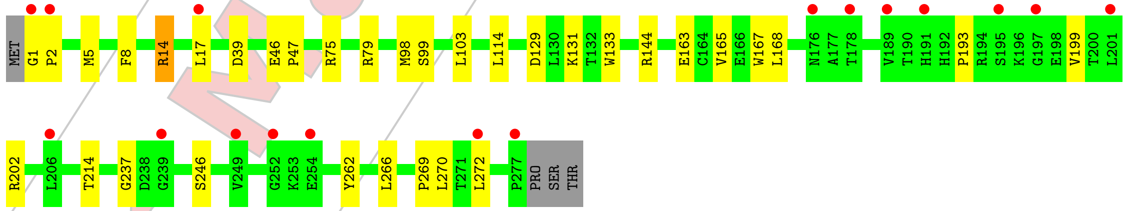


• Molecule 4: Beta-2-microglobulin

Chain D:

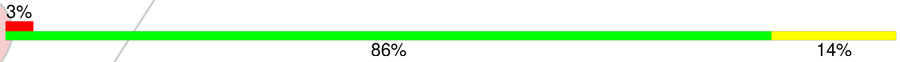


For Manuscript Review

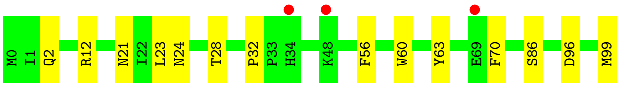


• Molecule 5: Nucleoprotein peptide

Chain P:

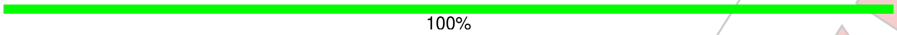


There are no outlier residues recorded for this chain.

For Manuscript Review

### 4 Data and refinement statistics (i)

| Property | Value | Source |
|---|---|---|
| Space group | P 21 21 21 | Depositor |
| Cell constants<br>a, b, c, $\alpha$, $\beta$, $\gamma$ | 58.17Å 131.70Å 141.94Å<br>90.00° 90.00° 90.00° | Depositor |
| Resolution (Å) | 49.83 – 2.78<br>49.83 – 2.78 | Depositor<br>EDS |
| % Data completeness<br>(in resolution range) | 95.1 (49.83-2.78)<br>95.1 (49.83-2.78) | Depositor<br>EDS |
| $R_{merge}$ | 0.16 | Depositor |
| $R_{sym}$ | (Not available) | Depositor |
| $< I/\sigma(I) >$ [1] | 1.00 (at 2.77Å) | Xtriage |
| Refinement program | PHENIX 1.21.2_5419 | Depositor |
| R, $R_{free}$ | 0.215 , 0.263<br>0.215 , 0.263 | Depositor<br>DCC |
| $R_{free}$ test set | 1380 reflections (4.84%) | wwPDB-VP |
| Wilson B-factor (Å$^2$) | 54.9 | Xtriage |
| Anisotropy | 0.415 | Xtriage |
| Bulk solvent $k_{sol}$(e/Å$^3$), $B_{sol}$(Å$^2$) | 0.33 , 53.9 | EDS |
| L-test for twinning[2] | $< \|L\| >= 0.49$, $< L^2 > = 0.32$ | Xtriage |
| Estimated twinning fraction | No twinning to report. | Xtriage |
| $F_o$,$F_c$ correlation | 0.93 | EDS |
| Total number of atoms | 6834 | wwPDB-VP |
| Average B, all atoms (Å$^2$) | 70.0 | wwPDB-VP |

Xtriage's analysis on translational NCS is as follows: *The largest off-origin peak in the Patterson function is 3.88% of the height of the origin peak. No significant pseudotranslation is detected.*

[1]Intensities estimated from amplitudes.

[2]Theoretical values of $< |L| >$, $< L^2 >$ for acentric reflections are 0.5, 0.333 respectively for untwinned datasets, and 0.375, 0.2 for perfectly twinned datasets.

For Manuscript Review

### 5 Model quality (i)

#### 5.1 Standard geometry (i)

Bond lengths and bond angles in the following residue types are not validated in this section: GOL, CL, NAG

The Z score for a bond length (or angle) is the number of standard deviations the observed value is removed from the expected value. A bond length (or angle) with $|Z| > 5$ is considered an outlier worth inspection. RMSZ is the root-mean-square of all Z scores of the bond lengths (or angles).

| Mol | Chain | Bond lengths | | Bond angles | |
|---|---|---|---|---|---|
| | | RMSZ | #$\|Z\|$ >5 | RMSZ | #$\|Z\|$ >5 |
| 1 | A | 0.26 | 0/1616 | 0.46 | 0/2187 |
| 2 | B | 0.25 | 0/2053 | 0.47 | 0/2792 |
| 3 | C | 0.25 | 0/2333 | 0.49 | 0/3175 |
| 4 | D | 0.25 | 0/865 | 0.46 | 0/1172 |
| 5 | P | 0.24 | 0/68 | 0.40 | 0/88 |
| All | All | 0.25 | 0/6935 | 0.48 | 0/9414 |

There are no bond length outliers.

There are no bond angle outliers.

There are no chirality outliers.

There are no planarity outliers.

#### 5.2 Too-close contacts (i)

In the following table, the Non-H and H(model) columns list the number of non-hydrogen atoms and hydrogen atoms in the chain respectively. The H(added) column lists the number of hydrogen atoms added and optimized by MolProbity. The Clashes column lists the number of clashes within the asymmetric unit, whereas Symm-Clashes lists symmetry-related clashes.

| Mol | Chain | Non-H | H(model) | H(added) | Clashes | Symm-Clashes |
|---|---|---|---|---|---|---|
| 1 | A | 1581 | 0 | 1500 | 14 | 0 |
| 2 | B | 1998 | 0 | 1912 | 20 | 0 |
| 3 | C | 2265 | 0 | 2106 | 22 | 0 |
| 4 | D | 838 | 0 | 806 | 11 | 0 |
| 5 | P | 69 | 0 | 61 | 0 | 0 |
| 6 | A | 14 | 0 | 13 | 1 | 0 |
| 6 | B | 14 | 0 | 13 | 1 | 0 |
| 7 | A | 6 | 0 | 8 | 1 | 0 |

*Continued on next page...*

For Manuscript Review

*Continued from previous page...*

| Mol | Chain | Non-H | H(model) | H(added) | Clashes | Symm-Clashes |
|---|---|---|---|---|---|---|
| 8 | B | 3 | 0 | 0 | 0 | 0 |
| 8 | C | 1 | 0 | 0 | 0 | 0 |
| 8 | D | 1 | 0 | 0 | 0 | 0 |
| 9 | A | 9 | 0 | 0 | 0 | 0 |
| 9 | B | 14 | 0 | 0 | 0 | 0 |
| 9 | C | 12 | 0 | 0 | 0 | 0 |
| 9 | D | 8 | 0 | 0 | 0 | 0 |
| 9 | P | 1 | 0 | 0 | 0 | 0 |
| All | All | 6834 | 0 | 6419 | 61 | 0 |

The all-atom clashscore is defined as the number of clashes found per 1000 atoms (including hydrogen atoms). The all-atom clashscore for this structure is 5.

All (61) close contacts within the same asymmetric unit are listed below, sorted by their clash magnitude.

| Atom-1 | Atom-2 | Interatomic distance (Å) | Clash overlap (Å) |
|---|---|---|---|
| 1:A:20:SER:HB2 | 6:A:301:NAG:H62 | 1.86 | 0.57 |
| 2:B:134:GLU:HA | 2:B:137:ILE:HD12 | 1.86 | 0.57 |
| 2:B:112:ARG:HD3 | 2:B:156:HIS:CD2 | 2.39 | 0.57 |
| 1:A:98:LYS:HE3 | 2:B:56:ASP:HB3 | 1.87 | 0.57 |
| 3:C:5:MET:HB2 | 3:C:168:LEU:HD13 | 1.86 | 0.56 |
| 3:C:193:PRO:HA | 3:C:199:VAL:HG23 | 1.87 | 0.56 |
| 1:A:133:VAL:HG12 | 1:A:176:TRP:HB3 | 1.87 | 0.56 |
| 3:C:237:GLY:HA3 | 4:D:12:ARG:HH21 | 1.72 | 0.54 |
| 2:B:20:ILE:HG12 | 2:B:77:GLN:HG3 | 1.90 | 0.52 |
| 2:B:203:TRP:HE3 | 2:B:210:PHE:HE2 | 1.56 | 0.52 |
| 1:A:193:ILE:HG22 | 1:A:197:THR:HG21 | 1.91 | 0.52 |
| 2:B:173:CYS:HB3 | 2:B:195:ARG:HD2 | 1.92 | 0.51 |
| 4:D:2:GLN:HG3 | 4:D:32:PRO:HD3 | 1.93 | 0.51 |
| 2:B:137:ILE:HG23 | 2:B:200:ALA:HB1 | 1.93 | 0.50 |
| 2:B:163:VAL:HG22 | 2:B:210:PHE:HD1 | 1.76 | 0.50 |
| 7:A:302:GOL:H31 | 2:B:104:PRO:HG3 | 1.94 | 0.50 |
| 3:C:103:LEU:HD13 | 3:C:168:LEU:HD23 | 1.94 | 0.49 |
| 3:C:202:ARG:HE | 3:C:246:SER:HB3 | 1.78 | 0.49 |
| 3:C:237:GLY:HA3 | 4:D:12:ARG:NH2 | 2.27 | 0.49 |
| 3:C:133:TRP:HB2 | 3:C:144:ARG:HG3 | 1.95 | 0.49 |
| 2:B:1:ASP:HA | 2:B:3:LYS:HE2 | 1.96 | 0.48 |
| 3:C:163:GLU:HG2 | 3:C:167:TRP:CD1 | 2.49 | 0.47 |
| 3:C:214:THR:HB | 3:C:262:TYR:HB2 | 1.96 | 0.47 |
| 2:B:41[A]:LYS:H | 2:B:41[A]:LYS:HG2 | 1.56 | 0.46 |
| 1:A:125:ASP:HB2 | 1:A:131:LYS:HB2 | 1.98 | 0.46 |

*Continued on next page...*

For Manuscript Review

*Continued from previous page...*

| Atom-1 | Atom-2 | Interatomic distance (Å) | Clash overlap (Å) |
|---|---|---|---|
| 3:C:46:GLU:HG3 | 3:C:47:PRO:HD2 | 1.98 | 0.46 |
| 3:C:98:MET:HG3 | 4:D:60:TRP:CZ3 | 2.50 | 0.46 |
| 3:C:103:LEU:HD11 | 3:C:165:VAL:HG13 | 1.97 | 0.45 |
| 4:D:24:ASN:N | 4:D:24:ASN:HD22 | 2.14 | 0.45 |
| 1:A:50:ILE:HG13 | 1:A:56:LYS:NZ | 2.32 | 0.44 |
| 4:D:28:THR:HG22 | 4:D:63:TYR:HB2 | 1.99 | 0.44 |
| 1:A:92:MET:HB2 | 1:A:99:LEU:HD13 | 1.98 | 0.44 |
| 3:C:163:GLU:HG2 | 3:C:167:TRP:HD1 | 1.83 | 0.44 |
| 4:D:21:ASN:HB3 | 4:D:70:PHE:CE1 | 2.53 | 0.43 |
| 4:D:23:LEU:HD13 | 4:D:70:PHE:CZ | 2.54 | 0.43 |
| 2:B:99:SER:HB3 | 2:B:103:SER:HB3 | 1.99 | 0.43 |
| 1:A:13:VAL:HG11 | 1:A:19:THR:HG22 | 2.01 | 0.42 |
| 2:B:83:PRO:HA | 2:B:115:VAL:HB | 2.01 | 0.42 |
| 4:D:2:GLN:HG2 | 4:D:86:SER:HB3 | 2.00 | 0.42 |
| 3:C:79:ARG:HE | 3:C:79:ARG:HB2 | 1.56 | 0.42 |
| 1:A:19:THR:HG21 | 1:A:107:LEU:HD21 | 2.01 | 0.42 |
| 1:A:111:PRO:HB3 | 1:A:169:LYS:HD2 | 2.02 | 0.42 |
| 3:C:1:GLY:HA2 | 3:C:2:PRO:HD3 | 1.91 | 0.42 |
| 3:C:266:LEU:HD13 | 3:C:270:LEU:HG | 2.02 | 0.42 |
| 2:B:41[B]:LYS:H | 2:B:41[B]:LYS:HG2 | 1.63 | 0.42 |
| 2:B:132:PRO:HG2 | 2:B:143:ALA:HB1 | 2.01 | 0.42 |
| 3:C:262:TYR:CE1 | 3:C:269:PRO:HB3 | 2.54 | 0.42 |
| 2:B:207:ARG:HD3 | 2:B:207:ARG:HA | 1.68 | 0.41 |
| 3:C:8:PHE:HB3 | 4:D:56:PHE:CE1 | 2.55 | 0.41 |
| 1:A:144:ASN:H | 1:A:192:ILE:HD11 | 1.85 | 0.41 |
| 2:B:65:GLN:NE2 | 6:B:301:NAG:H62 | 2.34 | 0.41 |
| 1:A:172:SER:HB2 | 2:B:195:ARG:HD3 | 2.02 | 0.41 |
| 2:B:231:LYS:HG2 | 2:B:233:VAL:HG13 | 2.02 | 0.41 |
| 3:C:262:TYR:CD1 | 3:C:269:PRO:HB3 | 2.55 | 0.41 |
| 1:A:119:ALA:HB3 | 1:A:121:TYR:HE1 | 1.86 | 0.41 |
| 1:A:118:PRO:O | 1:A:197:THR:HA | 2.21 | 0.41 |
| 3:C:129:ASP:O | 3:C:131:LYS:HG3 | 2.21 | 0.40 |
| 2:B:209:HIS:CE1 | 2:B:240:GLU:HB2 | 2.56 | 0.40 |
| 3:C:99:SER:HB3 | 3:C:114:LEU:HD12 | 2.02 | 0.40 |
| 4:D:96:ASP:HB3 | 4:D:99:MET:HB2 | 2.03 | 0.40 |
| 3:C:14[B]:ARG:HB3 | 3:C:17:LEU:HD23 | 2.03 | 0.40 |

There are no symmetry-related clashes.

For Manuscript Review

#### 5.3 Torsion angles (i)

##### 5.3.1 Protein backbone (i)

In the following table, the Percentiles column shows the percent Ramachandran outliers of the chain as a percentile score with respect to all X-ray entries followed by that with respect to entries of similar resolution.

The Analysed column shows the number of residues for which the backbone conformation was analysed, and the total number of residues.

| Mol | Chain | Analysed | Favoured | Allowed | Outliers | Percentiles | |
|---|---|---|---|---|---|---|---|
| 1 | A | 202/216 (94%) | 188 (93%) | 13 (6%) | 1 (0%) | 25 | 52 |
| 2 | B | 246/257 (96%) | 235 (96%) | 11 (4%) | 0 | 100 | 100 |
| 3 | C | 277/281 (99%) | 263 (95%) | 14 (5%) | 0 | 100 | 100 |
| 4 | D | 99/100 (99%) | 95 (96%) | 4 (4%) | 0 | 100 | 100 |
| 5 | P | 7/9 (78%) | 6 (86%) | 1 (14%) | 0 | 100 | 100 |
| All | All | 831/863 (96%) | 787 (95%) | 43 (5%) | 1 (0%) | 48 | 76 |

All (1) Ramachandran outliers are listed below:

| Mol | Chain | Res | Type |
|---|---|---|---|
| 1 | A | 202 | PRO |

##### 5.3.2 Protein sidechains (i)

In the following table, the Percentiles column shows the percent sidechain outliers of the chain as a percentile score with respect to all X-ray entries followed by that with respect to entries of similar resolution.

The Analysed column shows the number of residues for which the sidechain conformation was analysed, and the total number of residues.

| Mol | Chain | Analysed | Rotameric | Outliers | Percentiles | |
|---|---|---|---|---|---|---|
| 1 | A | 179/194 (92%) | 175 (98%) | 4 (2%) | 47 | 76 |
| 2 | B | 218/228 (96%) | 208 (95%) | 10 (5%) | 23 | 52 |
| 3 | C | 230/239 (96%) | 225 (98%) | 5 (2%) | 47 | 76 |
| 4 | D | 95/95 (100%) | 95 (100%) | 0 | 100 | 100 |
| 5 | P | 8/8 (100%) | 8 (100%) | 0 | 100 | 100 |
| All | All | 730/764 (96%) | 711 (97%) | 19 (3%) | 44 | 72 |

For Manuscript Review

All (19) residues with a non-rotameric sidechain are listed below:

| Mol | Chain | Res | Type |
|---|---|---|---|
| 1 | A | 106 | LYS |
| 1 | A | 140 | ASP |
| 1 | A | 161 | LEU |
| 1 | A | 191 | SER |
| 2 | B | 3 | LYS |
| 2 | B | 41[A] | LYS |
| 2 | B | 41[B] | LYS |
| 2 | B | 59 | THR |
| 2 | B | 73 | HIS |
| 2 | B | 145 | LEU |
| 2 | B | 173 | CYS |
| 2 | B | 207 | ARG |
| 2 | B | 224 | GLU |
| 2 | B | 228 | ASP |
| 3 | C | 14[A] | ARG |
| 3 | C | 14[B] | ARG |
| 3 | C | 39 | ASP |
| 3 | C | 75 | ARG |
| 3 | C | 272 | LEU |

Sometimes sidechains can be flipped to improve hydrogen bonding and reduce clashes. All (5) such sidechains are listed below:

| Mol | Chain | Res | Type |
|---|---|---|---|
| 1 | A | 1 | GLN |
| 1 | A | 64 | HIS |
| 2 | B | 29 | HIS |
| 3 | C | 174 | ASN |
| 4 | D | 24 | ASN |

##### 5.3.3 RNA (i)

There are no RNA molecules in this entry.

#### 5.4 Non-standard residues in protein, DNA, RNA chains (i)

There are no non-standard protein/DNA/RNA residues in this entry.

For Manuscript Review

#### 5.5 Carbohydrates (i)

There are no oligosaccharides in this entry.

#### 5.6 Ligand geometry (i)

Of 8 ligands modelled in this entry, 5 are monoatomic - leaving 3 for Mogul analysis.

In the following table, the Counts columns list the number of bonds (or angles) for which Mogul statistics could be retrieved, the number of bonds (or angles) that are observed in the model and the number of bonds (or angles) that are defined in the Chemical Component Dictionary. The Link column lists molecule types, if any, to which the group is linked. The Z score for a bond length (or angle) is the number of standard deviations the observed value is removed from the expected value. A bond length (or angle) with $|Z| > 2$ is considered an outlier worth inspection. RMSZ is the root-mean-square of all Z scores of the bond lengths (or angles).

| Mol | Type | Chain | Res | Link | Bond lengths | | | Bond angles | | |
|---|---|---|---|---|---|---|---|---|---|---|
| | | | | | Counts | RMSZ | $\#\|Z\| > 2$ | Counts | RMSZ | $\#\|Z\| > 2$ |
| 6 | NAG | A | 301 | 1 | 14,14,15 | 0.74 | 0 | 17,19,21 | 0.81 | 0 |
| 6 | NAG | B | 301 | 2 | 14,14,15 | 0.83 | 1 (7%) | 17,19,21 | 1.02 | 1 (5%) |
| 7 | GOL | A | 302 | - | 5,5,5 | 0.34 | 0 | 5,5,5 | 0.42 | 0 |

In the following table, the Chirals column lists the number of chiral outliers, the number of chiral centers analysed, the number of these observed in the model and the number defined in the Chemical Component Dictionary. Similar counts are reported in the Torsion and Rings columns. '-' means no outliers of that kind were identified.

| Mol | Type | Chain | Res | Link | Chirals | Torsions | Rings |
|---|---|---|---|---|---|---|---|
| 6 | NAG | A | 301 | 1 | - | 1/6/23/26 | 0/1/1/1 |
| 6 | NAG | B | 301 | 2 | - | 1/6/23/26 | 0/1/1/1 |
| 7 | GOL | A | 302 | - | - | 0/4/4/4 | - |

All (1) bond length outliers are listed below:

| Mol | Chain | Res | Type | Atoms | Z | Observed(Å) | Ideal(Å) |
|---|---|---|---|---|---|---|---|
| 6 | B | 301 | NAG | O5-C1 | -2.03 | 1.40 | 1.43 |

All (1) bond angle outliers are listed below:

| Mol | Chain | Res | Type | Atoms | Z | Observed($^o$) | Ideal($^o$) |
|---|---|---|---|---|---|---|---|
| 6 | B | 301 | NAG | O5-C1-C2 | -2.67 | 107.17 | 111.29 |

There are no chirality outliers.

For Manuscript Review

All (2) torsion outliers are listed below:

| Mol | Chain | Res | Type | Atoms |
|---|---|---|---|---|
| 6 | A | 301 | NAG | O5-C5-C6-O6 |
| 6 | B | 301 | NAG | O5-C5-C6-O6 |

There are no ring outliers.

3 monomers are involved in 3 short contacts:

| Mol | Chain | Res | Type | Clashes | Symm-Clashes |
|---|---|---|---|---|---|
| 6 | A | 301 | NAG | 1 | 0 |
| 6 | B | 301 | NAG | 1 | 0 |
| 7 | A | 302 | GOL | 1 | 0 |

The following is a two-dimensional graphical depiction of Mogul quality analysis of bond lengths, bond angles, torsion angles, and ring geometry for all instances of the Ligand of Interest. In addition, ligands with molecular weight > 250 and outliers as shown on the validation Tables will also be included. For torsion angles, if less then 5% of the Mogul distribution of torsion angles is within 10 degrees of the torsion angle in question, then that torsion angle is considered an outlier. Any bond that is central to one or more torsion angles identified as an outlier by Mogul will be highlighted in the graph. For rings, the root-mean-square deviation (RMSD) between the ring in question and similar rings identified by Mogul is calculated over all ring torsion angles. If the average RMSD is greater than 60 degrees and the minimal RMSD between the ring in question and any Mogul-identified rings is also greater than 60 degrees, then that ring is considered an outlier. The outliers are highlighted in purple. The color gray indicates Mogul did not find sufficient equivalents in the CSD to analyse the geometry.

For Manuscript Review

#### Ligand NAG A 301

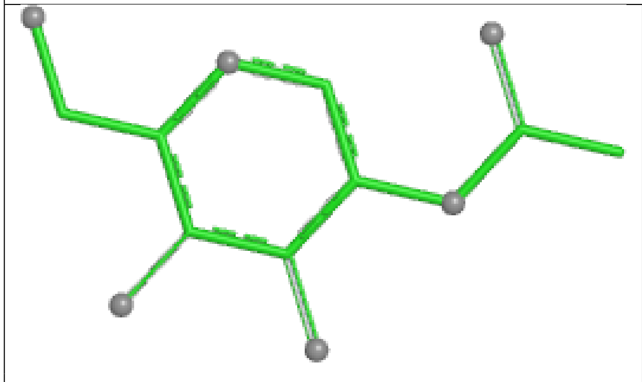

Bond lengths

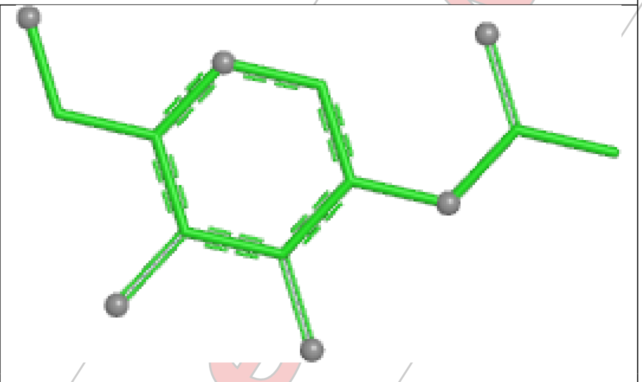

Bond angles

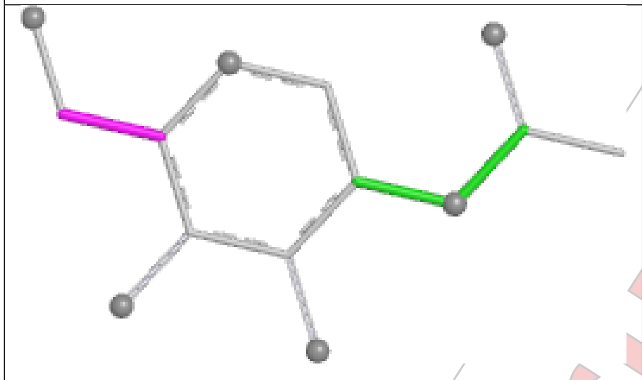

Torsions

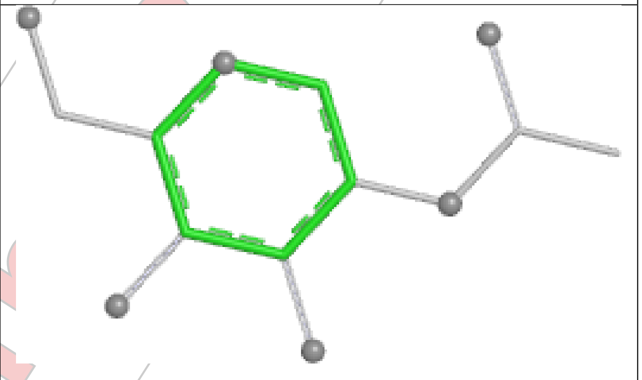

Rings

#### Ligand NAG B 301

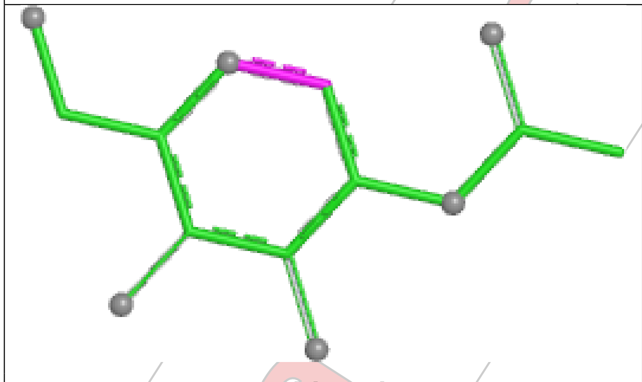

Bond lengths

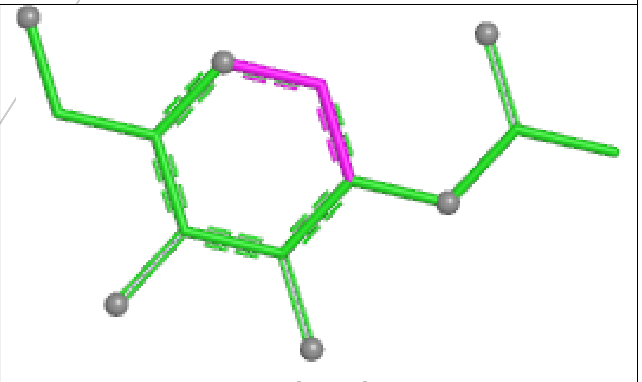

Bond angles

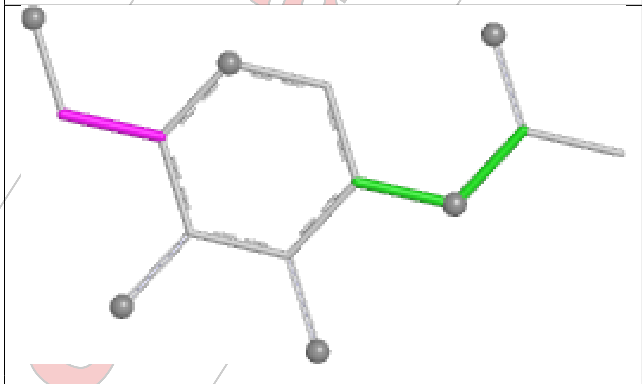

Torsions

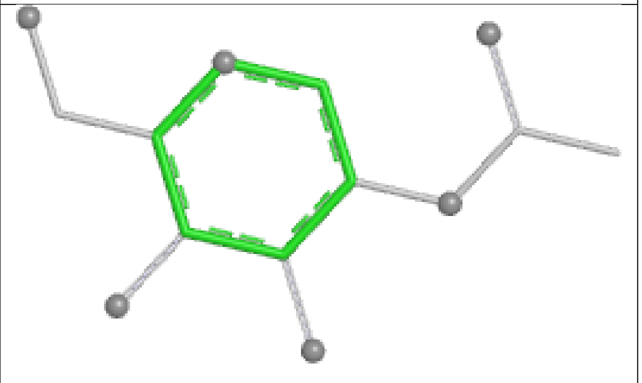

Rings

#### 5.7 Other polymers (i)

There are no such residues in this entry.

#### 5.8 Polymer linkage issues (i)

There are no chain breaks in this entry.

For Manuscript Review

### 6 Fit of model and data (i)

#### 6.1 Protein, DNA and RNA chains (i)

In the following table, the column labelled '#RSRZ> 2' contains the number (and percentage) of RSRZ outliers, followed by percent RSRZ outliers for the chain as percentile scores relative to all X-ray entries and entries of similar resolution. The OWAB column contains the minimum, median, $95^{th}$ percentile and maximum values of the occupancy-weighted average B-factor per residue. The column labelled 'Q< 0.9' lists the number of (and percentage) of residues with an average occupancy less than 0.9.

| Mol | Chain | Analysed | <RSRZ> | #RSRZ>2 | | | OWAB(Å$^2$) | Q<0.9 |
|---|---|---|---|---|---|---|---|---|
| 1 | A | 203/216 (93%) | 0.68 | 19 (9%) | 15 | 13 | 26, 67, 129, 167 | 1 (0%) |
| 2 | B | 245/257 (95%) | 0.22 | 7 (2%) | 54 | 48 | 27, 61, 106, 147 | 3 (1%) |
| 3 | C | 277/281 (98%) | 0.34 | 17 (6%) | 28 | 24 | 31, 65, 114, 144 | 2 (0%) |
| 4 | D | 100/100 (100%) | 0.30 | 3 (3%) | 52 | 47 | 26, 71, 104, 126 | 1 (1%) |
| 5 | P | 9/9 (100%) | -0.27 | 0 | 100 | 100 | 40, 45, 49, 54 | 0 |
| All | All | 834/863 (96%) | 0.38 | 46 (5%) | 32 | 27 | 26, 63, 119, 167 | 7 (0%) |

All (46) RSRZ outliers are listed below:

| Mol | Chain | Res | Type | RSRZ |
|---|---|---|---|---|
| 1 | A | 141 | SER | 4.2 |
| 3 | C | 195 | SER | 4.0 |
| 2 | B | 245 | ALA | 3.7 |
| 3 | C | 197 | GLY | 3.4 |
| 3 | C | 206 | LEU | 3.2 |
| 3 | C | 254 | GLU | 3.2 |
| 3 | C | 1 | GLY | 3.2 |
| 2 | B | 59 | THR | 3.1 |
| 4 | D | 34[A] | HIS | 3.1 |
| 1 | A | 72 | PHE | 3.0 |
| 4 | D | 69 | GLU | 2.9 |
| 1 | A | 120 | VAL | 2.8 |
| 2 | B | 1 | ASP | 2.7 |
| 3 | C | 189 | VAL | 2.6 |
| 1 | A | 119 | ALA | 2.6 |
| 3 | C | 277 | PRO | 2.6 |
| 1 | A | 152 | ASP | 2.6 |
| 1 | A | 185 | ALA | 2.6 |
| 1 | A | 159 | CYS | 2.6 |
| 1 | A | 165 | SER | 2.6 |

*Continued on next page...*

For Manuscript Review

*Continued from previous page...*

| Mol | Chain | Res | Type | RSRZ |
|---|---|---|---|---|
| 1 | A | 179 | LYS | 2.5 |
| 2 | B | 168 | VAL | 2.5 |
| 3 | C | 178 | THR | 2.5 |
| 3 | C | 191 | HIS | 2.5 |
| 1 | A | 181 | ASP | 2.5 |
| 3 | C | 2 | PRO | 2.4 |
| 3 | C | 249 | VAL | 2.4 |
| 1 | A | 184 | CYS | 2.3 |
| 1 | A | 202 | PRO | 2.3 |
| 1 | A | 193 | ILE | 2.3 |
| 3 | C | 176 | ASN | 2.2 |
| 2 | B | 173 | CYS | 2.2 |
| 3 | C | 252 | GLY | 2.2 |
| 1 | A | 197 | THR | 2.2 |
| 1 | A | 160 | VAL | 2.2 |
| 3 | C | 239 | GLY | 2.2 |
| 2 | B | 177 | GLN | 2.2 |
| 1 | A | 52 | SER | 2.1 |
| 3 | C | 17 | LEU | 2.1 |
| 3 | C | 272 | LEU | 2.1 |
| 4 | D | 48 | LYS | 2.1 |
| 1 | A | 121 | TYR | 2.1 |
| 3 | C | 201 | LEU | 2.1 |
| 2 | B | 140 | THR | 2.0 |
| 1 | A | 136 | PHE | 2.0 |
| 1 | A | 168 | PHE | 2.0 |

#### 6.2 Non-standard residues in protein, DNA, RNA chains (i)

There are no non-standard protein/DNA/RNA residues in this entry.

#### 6.3 Carbohydrates (i)

There are no monosaccharides in this entry.

#### 6.4 Ligands (i)

In the following table, the Atoms column lists the number of modelled atoms in the group and the number defined in the chemical component dictionary. The B-factors column lists the minimum, median, $95^{th}$ percentile and maximum values of B factors of atoms in the group. The column labelled 'Q< 0.9' lists the number of atoms with occupancy less than 0.9.

For Manuscript Review

| Mol | Type | Chain | Res | Atoms | RSCC | RSR | B-factors($Å^2$) | Q<0.9 |
|---|---|---|---|---|---|---|---|---|
| 6 | NAG | A | 301 | 14/15 | 0.63 | 0.19 | 71,77,85,87 | 0 |
| 8 | CL | B | 303 | 1/1 | 0.73 | 0.15 | 88,88,88,88 | 0 |
| 6 | NAG | B | 301 | 14/15 | 0.76 | 0.18 | 68,80,86,96 | 0 |
| 8 | CL | D | 101 | 1/1 | 0.86 | 0.12 | 78,78,78,78 | 0 |
| 8 | CL | B | 304 | 1/1 | 0.87 | 0.15 | 64,64,64,64 | 0 |
| 8 | CL | C | 301 | 1/1 | 0.87 | 0.12 | 90,90,90,90 | 0 |
| 8 | CL | B | 302 | 1/1 | 0.87 | 0.11 | 62,62,62,62 | 0 |
| 7 | GOL | A | 302 | 6/6 | 0.88 | 0.13 | 49,58,75,75 | 0 |

The following is a graphical depiction of the model fit to experimental electron density of all instances of the Ligand of Interest. In addition, ligands with molecular weight > 250 and outliers as shown on the geometry validation Tables will also be included. Each fit is shown from different orientation to approximate a three-dimensional view.

**Electron density around NAG A 301:**

2mF$_o$-DF$_c$ (at 0.7 rmsd) in gray
mF$_o$-DF$_c$ (at 3 rmsd) in purple (negative) and green (positive)

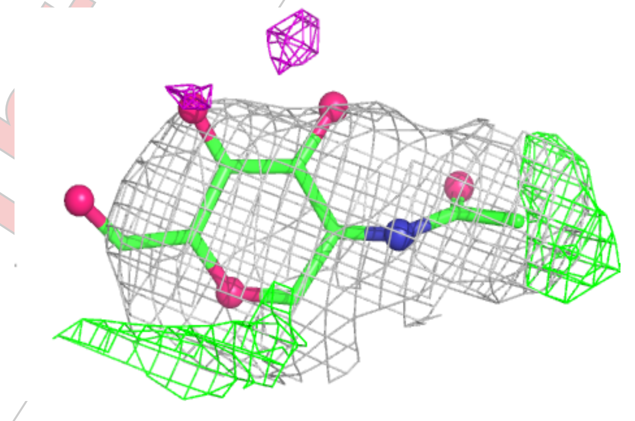

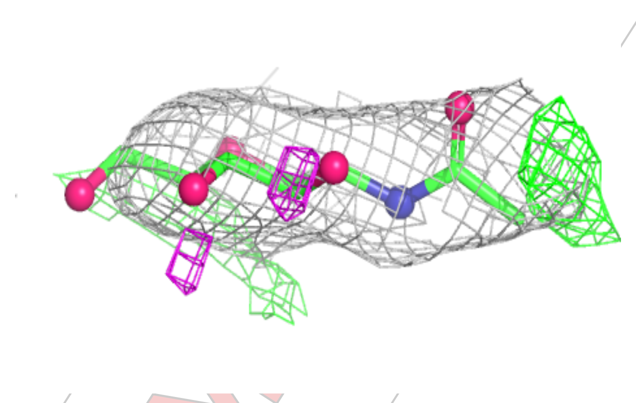

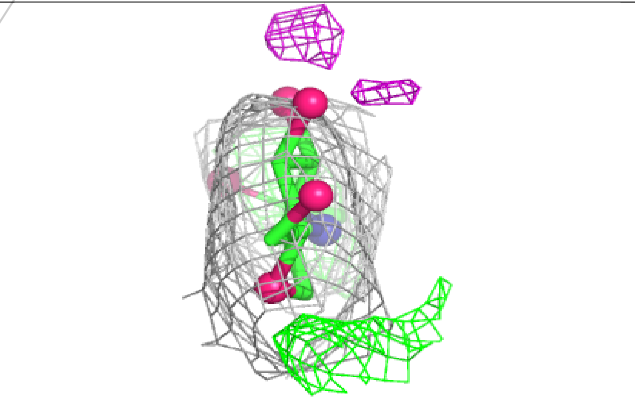

For Manuscript Review

**Electron density around NAG B 301:**

$2mF_o$-$DF_c$ (at 0.7 rmsd) in gray
$mF_o$-$DF_c$ (at 3 rmsd) in purple (negative)
and green (positive)

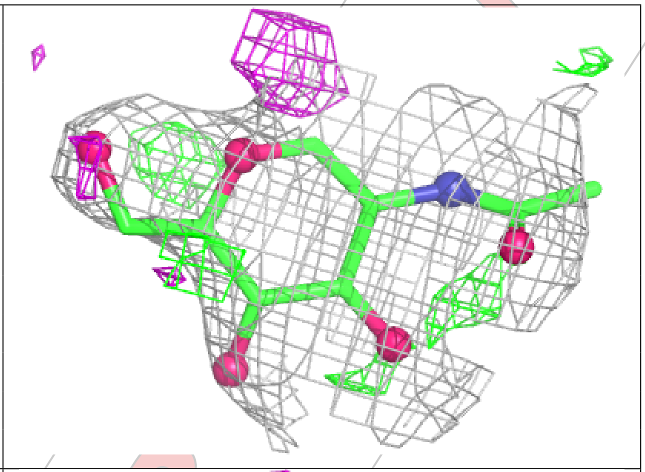

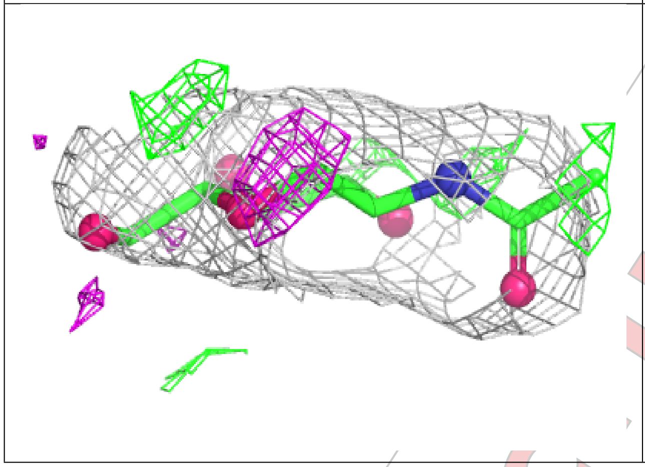

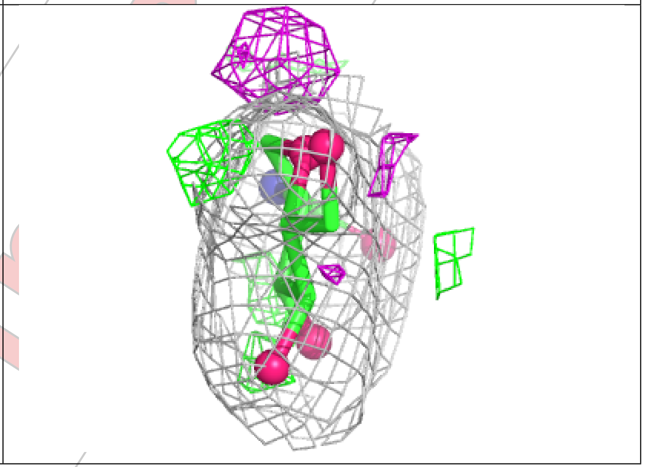

#### 6.5 Other polymers ⓘ

There are no such residues in this entry.

For Manuscript Review

WORLDWIDE PDB PROTEIN DATA BANK

### Full wwPDB X-ray Structure Validation Report (i)

Mar 21, 2025 – 03:44 PM EDT

| | | |
|---|---|---|
| PDB ID | : | 9NVE |
| Title | : | Crystal structure of NP203 TCR in complex with NP366-H-2Db |
| Deposited on | : | 2025-03-20 |
| Resolution | : | 2.10 Å(reported) |

**This wwPDB validation report is for manuscript review**

This is a Full wwPDB X-ray Structure Validation Report.

This report is produced by the wwPDB biocuration pipeline after annotation of the structure.

We welcome your comments at **
A user guide is available at
https://www.wwpdb.org/validation/2017/XrayValidationReportHelp
with specific help available everywhere you see the (i) symbol.

The types of validation reports are described at
http://www.wwpdb.org/validation/2017/FAQs#types.

---

The following versions of software and data (see references (i)) were used in the production of this report:

| | | |
|---|---|---|
| MolProbity | : | 4.02b-467 |
| Mogul | : | 2022.3.0, CSD as543be (2022) |
| Xtriage (Phenix) | : | 1.21 |
| EDS | : | 3.0 |
| buster-report | : | 1.1.7 (2018) |
| Percentile statistics | : | 20231227.v01 (using entries in the PDB archive December 27th 2023) |
| CCP4 | : | 9.0.004 (Gargrove) |
| Density-Fitness | : | 1.0.11 |
| Ideal geometry (proteins) | : | Engh & Huber (2001) |
| Ideal geometry (DNA, RNA) | : | Parkinson et al. (1996) |

### 1 Overall quality at a glance (i)

The following experimental techniques were used to determine the structure:
*X-RAY DIFFRACTION*

The reported resolution of this entry is 2.10 Å.

Percentile scores (ranging between 0-100) for global validation metrics of the entry are shown in the following graphic. The table shows the number of entries on which the scores are based.

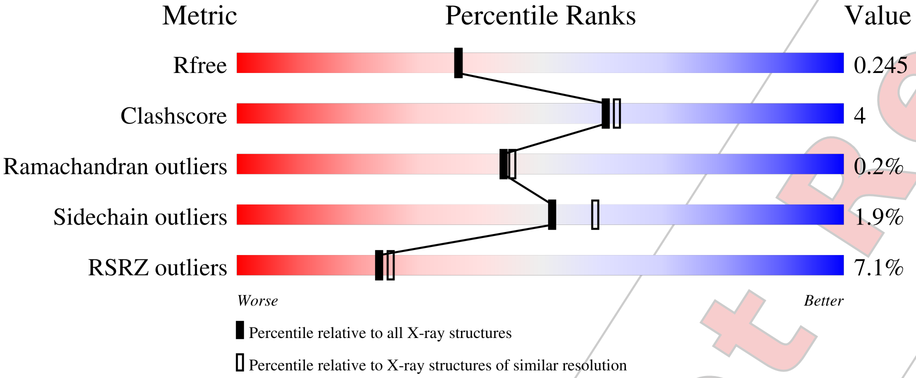


| Metric | Whole archive (#Entries) | Similar resolution (#Entries, resolution range(Å)) |
|---|---|---|
| $R_{free}$ | 164625 | 6234 (2.10-2.10) |
| Clashscore | 180529 | 6893 (2.10-2.10) |
| Ramachandran outliers | 177936 | 6839 (2.10-2.10) |
| Sidechain outliers | 177891 | 6840 (2.10-2.10) |
| RSRZ outliers | 164620 | 6234 (2.10-2.10) |

The table below summarises the geometric issues observed across the polymeric chains and their fit to the electron density. The red, orange, yellow and green segments of the lower bar indicate the fraction of residues that contain outliers for >=3, 2, 1 and 0 types of geometric quality criteria respectively. A grey segment represents the fraction of residues that are not modelled. The numeric value for each fraction is indicated below the corresponding segment, with a dot representing fractions <=5% The upper red bar (where present) indicates the fraction of residues that have poor fit to the electron density. The numeric value is given above the bar.

| Mol | Chain | Length | Quality of chain |
|---|---|---|---|
| 1 | A | 223 | 6% / 85% 10% 5% |
| 2 | B | 254 | 2% / 85% 11% 5% |
| 3 | C | 281 | 13% / 85% 12% •• |
| 4 | D | 100 | 5% / 90% 10% |

*Continued on next page...*



For Manuscript Review

*Continued from previous page...*

| Mol | Chain | Length | Quality of chain |
|---|---|---|---|
| 5 | P | 9 | 11% / 89% / 11% |
| 6 | E | 2 | 50% / 50% |

For Manuscript Review

### 2 Entry composition (i)

There are 10 unique types of molecules in this entry. The entry contains 6963 atoms, of which 0 are hydrogens and 0 are deuteriums.

In the tables below, the ZeroOcc column contains the number of atoms modelled with zero occupancy, the AltConf column contains the number of residues with at least one atom in alternate conformation and the Trace column contains the number of residues modelled with at most 2 atoms.

- Molecule 1 is a protein called NP203 TCR alpha-chain.

| Mol | Chain | Residues | Atoms | | | | | ZeroOcc | AltConf | Trace |
|---|---|---|---|---|---|---|---|---|---|---|
| | | | Total | C | N | O | S | | | |
| 1 | A | 212 | 1656 | 1038 | 270 | 338 | 10 | 0 | 0 | 0 |

- Molecule 2 is a protein called NP203 TCR beta-chain.

| Mol | Chain | Residues | Atoms | | | | | ZeroOcc | AltConf | Trace |
|---|---|---|---|---|---|---|---|---|---|---|
| | | | Total | C | N | O | S | | | |
| 2 | B | 242 | 1944 | 1235 | 333 | 368 | 8 | 0 | 0 | 0 |

- Molecule 3 is a protein called H-2 class I histocompatibility antigen, D-B alpha chain.

| Mol | Chain | Residues | Atoms | | | | | ZeroOcc | AltConf | Trace |
|---|---|---|---|---|---|---|---|---|---|---|
| | | | Total | C | N | O | S | | | |
| 3 | C | 277 | 2250 | 1423 | 399 | 418 | 10 | 0 | 0 | 0 |

There is a discrepancy between the modelled and reference sequences:

| Chain | Residue | Modelled | Actual | Comment | Reference |
|---|---|---|---|---|---|
| C | 0 | MET | - | initiating methionine | UNP P01899 |

- Molecule 4 is a protein called Beta-2-microglobulin.

| Mol | Chain | Residues | Atoms | | | | | ZeroOcc | AltConf | Trace |
|---|---|---|---|---|---|---|---|---|---|---|
| | | | Total | C | N | O | S | | | |
| 4 | D | 100 | 826 | 527 | 139 | 153 | 7 | 0 | 0 | 0 |

There are 2 discrepancies between the modelled and reference sequences:

| Chain | Residue | Modelled | Actual | Comment | Reference |
|---|---|---|---|---|---|
| D | 0 | MET | - | initiating methionine | UNP P01887 |
| D | 85 | ASP | ALA | conflict | UNP P01887 |

For Manuscript Review

- Molecule 5 is a protein called Nucleoprotein peptide.

| Mol | Chain | Residues | Atoms | ZeroOcc | AltConf | Trace |
|---|---|---|---|---|---|---|
| 5 | P | 9 | Total C N O S<br>69 38 11 18 2 | 0 | 0 | 0 |

- Molecule 6 is an oligosaccharide called 2-acetamido-2-deoxy-beta-D-glucopyranose-(1-4)-2-acetamido-2-deoxy-beta-D-glucopyranose.

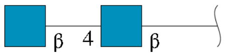


| Mol | Chain | Residues | Atoms | ZeroOcc | AltConf | Trace |
|---|---|---|---|---|---|---|
| 6 | E | 2 | Total C N O<br>28 16 2 10 | 0 | 0 | 0 |

- Molecule 7 is 2-acetamido-2-deoxy-beta-D-glucopyranose (three-letter code: NAG) (formula: $C_8H_{15}NO_6$) (labeled as "Ligand of Interest" by depositor).

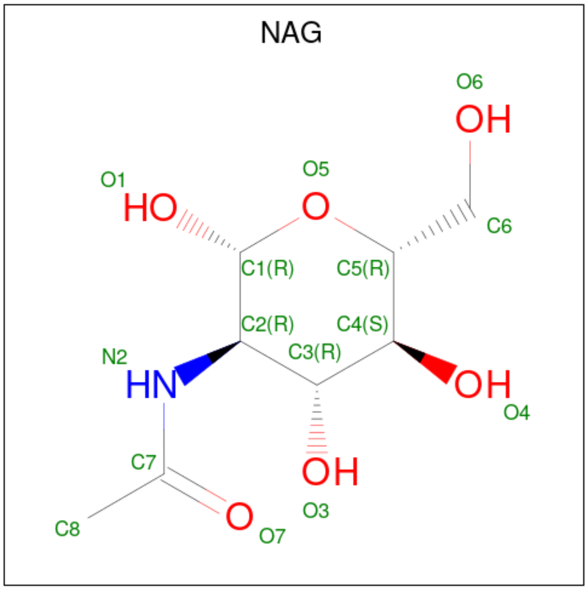


| Mol | Chain | Residues | Atoms | ZeroOcc | AltConf |
|---|---|---|---|---|---|
| 7 | A | 1 | Total C N O<br>14 8 1 5 | 0 | 0 |
| 7 | B | 1 | Total C N O<br>14 8 1 5 | 0 | 0 |

- Molecule 8 is 1,2-ETHANEDIOL (three-letter code: EDO) (formula: $C_2H_6O_2$).

For Manuscript Review

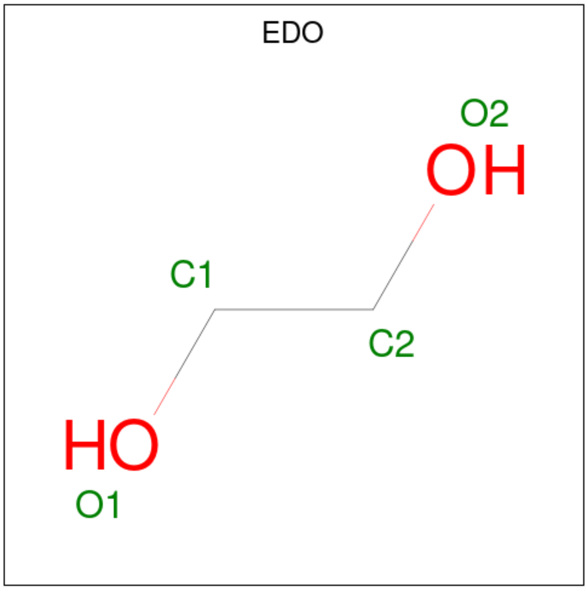


| Mol | Chain | Residues | Atoms | ZeroOcc | AltConf |
|---|---|---|---|---|---|
| 8 | A | 1 | Total C O<br>4 2 2 | 0 | 0 |
| 8 | B | 1 | Total C O<br>4 2 2 | 0 | 0 |
| 8 | B | 1 | Total C O<br>4 2 2 | 0 | 0 |
| 8 | B | 1 | Total C O<br>4 2 2 | 0 | 0 |

- Molecule 9 is CHLORIDE ION (three-letter code: CL) (formula: Cl).

| Mol | Chain | Residues | Atoms | ZeroOcc | AltConf |
|---|---|---|---|---|---|
| 9 | A | 1 | Total Cl<br>1 1 | 0 | 0 |
| 9 | B | 1 | Total Cl<br>1 1 | 0 | 0 |
| 9 | D | 1 | Total Cl<br>1 1 | 0 | 0 |

- Molecule 10 is water.

| Mol | Chain | Residues | Atoms | ZeroOcc | AltConf |
|---|---|---|---|---|---|
| 10 | A | 21 | Total O<br>21 21 | 0 | 0 |
| 10 | B | 60 | Total O<br>60 60 | 0 | 0 |

*Continued on next page...*

For Manuscript Review

*Continued from previous page...*

| Mol | Chain | Residues | Atoms | ZeroOcc | AltConf |
|---|---|---|---|---|---|
| 10 | C | 48 | Total O<br>48 48 | 0 | 0 |
| 10 | D | 8 | Total O<br>8 8 | 0 | 0 |
| 10 | P | 6 | Total O<br>6 6 | 0 | 0 |

For Manuscript Review

### 3 Residue-property plots (i)

These plots are drawn for all protein, RNA, DNA and oligosaccharide chains in the entry. The first graphic for a chain summarises the proportions of the various outlier classes displayed in the second graphic. The second graphic shows the sequence view annotated by issues in geometry and electron density. Residues are color-coded according to the number of geometric quality criteria for which they contain at least one outlier: green = 0, yellow = 1, orange = 2 and red = 3 or more. A red dot above a residue indicates a poor fit to the electron density (RSRZ > 2). Stretches of 2 or more consecutive residues without any outlier are shown as a green connector. Residues present in the sample, but not in the model, are shown in grey.

• Molecule 1: NP203 TCR alpha-chain

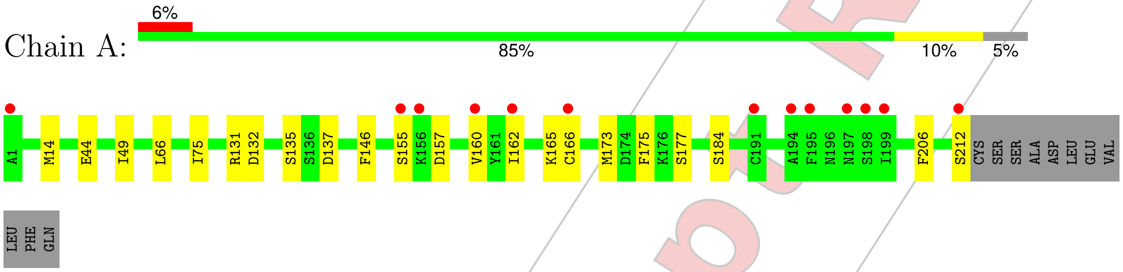


• Molecule 2: NP203 TCR beta-chain

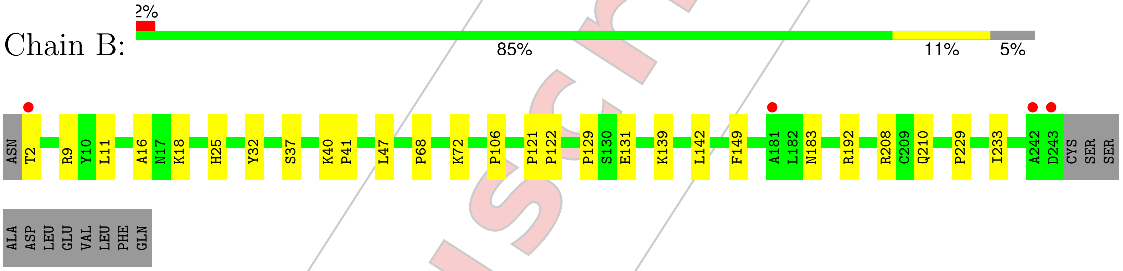


• Molecule 3: H-2 class I histocompatibility antigen, D-B alpha chain

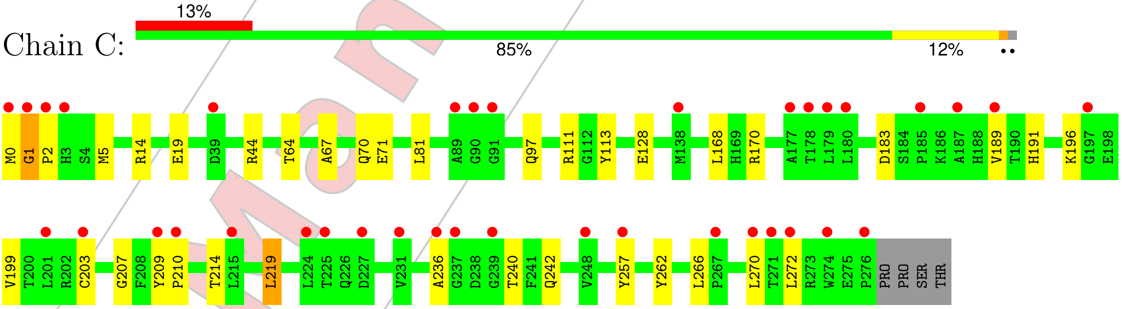


• Molecule 4: Beta-2-microglobulin

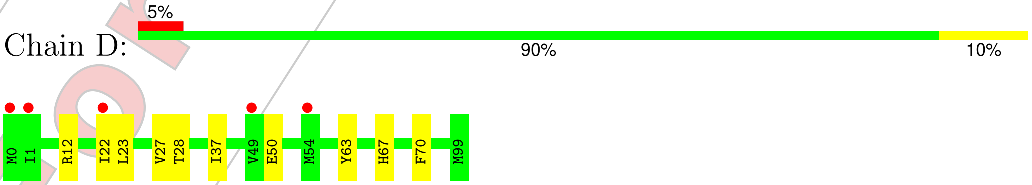


• Molecule 5: Nucleoprotein peptide

For Manuscript Review

Chain P:

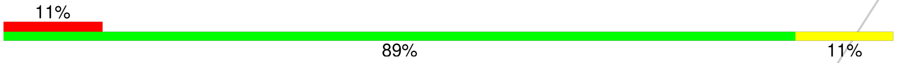


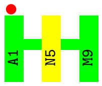


• Molecule 6: 2-acetamido-2-deoxy-beta-D-glucopyranose-(1-4)-2-acetamido-2-deoxy-beta-D-glucopyranose

Chain E:

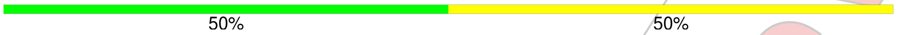


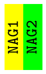


For Manuscript Review

### 4 Data and refinement statistics (i)

| Property | Value | Source |
|---|---|---|
| Space group | P 1 | Depositor |
| Cell constants<br>a, b, c, $\alpha$, $\beta$, $\gamma$ | 56.78Å 66.62Å 76.78Å<br>68.78° 89.27° 66.42° | Depositor |
| Resolution (Å) | 49.94 – 2.10<br>49.94 – 2.10 | Depositor<br>EDS |
| % Data completeness<br>(in resolution range) | 95.8 (49.94-2.10)<br>95.8 (49.94-2.10) | Depositor<br>EDS |
| $R_{merge}$ | 0.04 | Depositor |
| $R_{sym}$ | (Not available) | Depositor |
| $< I/\sigma(I) >$ [1] | 1.49 (at 2.10Å) | Xtriage |
| Refinement program | PHENIX 1.21.2_5419 | Depositor |
| R, $R_{free}$ | 0.211 , 0.245<br>0.211 , 0.245 | Depositor<br>DCC |
| $R_{free}$ test set | 2770 reflections (4.96%) | wwPDB-VP |
| Wilson B-factor (Å$^2$) | 39.5 | Xtriage |
| Anisotropy | 0.343 | Xtriage |
| Bulk solvent $k_{sol}$(e/Å$^3$), $B_{sol}$(Å$^2$) | 0.31 , 42.4 | EDS |
| L-test for twinning[2] | $< \|L\| >$ = 0.49, $< L^2 >$ = 0.32 | Xtriage |
| Estimated twinning fraction | 0.013 for h,h-k,-l | Xtriage |
| $F_o$,$F_c$ correlation | 0.95 | EDS |
| Total number of atoms | 6963 | wwPDB-VP |
| Average B, all atoms (Å$^2$) | 65.0 | wwPDB-VP |

Xtriage's analysis on translational NCS is as follows: *The largest off-origin peak in the Patterson function is 5.92% of the height of the origin peak. No significant pseudotranslation is detected.*

[1] Intensities estimated from amplitudes.

[2] Theoretical values of $< |L| >$, $< L^2 >$ for acentric reflections are 0.5, 0.333 respectively for untwinned datasets, and 0.375, 0.2 for perfectly twinned datasets.

For Manuscript Review

### 5 Model quality (i)

#### 5.1 Standard geometry (i)

Bond lengths and bond angles in the following residue types are not validated in this section: EDO, CL, NAG

The Z score for a bond length (or angle) is the number of standard deviations the observed value is removed from the expected value. A bond length (or angle) with $|Z| > 5$ is considered an outlier worth inspection. RMSZ is the root-mean-square of all Z scores of the bond lengths (or angles).

| Mol | Chain | Bond lengths | | Bond angles | |
|---|---|---|---|---|---|
| | | RMSZ | #$\|Z\|$ >5 | RMSZ | #$\|Z\|$ >5 |
| 1 | A | 0.25 | 0/1689 | 0.48 | 0/2286 |
| 2 | B | 0.25 | 0/1999 | 0.50 | 0/2723 |
| 3 | C | 0.24 | 0/2317 | 0.50 | 0/3148 |
| 4 | D | 0.25 | 0/852 | 0.51 | 0/1155 |
| 5 | P | 0.26 | 0/68 | 0.41 | 0/88 |
| All | All | 0.25 | 0/6925 | 0.49 | 0/9400 |

There are no bond length outliers.

There are no bond angle outliers.

There are no chirality outliers.

There are no planarity outliers.

#### 5.2 Too-close contacts (i)

In the following table, the Non-H and H(model) columns list the number of non-hydrogen atoms and hydrogen atoms in the chain respectively. The H(added) column lists the number of hydrogen atoms added and optimized by MolProbity. The Clashes column lists the number of clashes within the asymmetric unit, whereas Symm-Clashes lists symmetry-related clashes.

| Mol | Chain | Non-H | H(model) | H(added) | Clashes | Symm-Clashes |
|---|---|---|---|---|---|---|
| 1 | A | 1656 | 0 | 1584 | 11 | 0 |
| 2 | B | 1944 | 0 | 1864 | 18 | 0 |
| 3 | C | 2250 | 0 | 2114 | 22 | 0 |
| 4 | D | 826 | 0 | 798 | 8 | 0 |
| 5 | P | 69 | 0 | 61 | 4 | 0 |
| 6 | E | 28 | 0 | 25 | 0 | 0 |
| 7 | A | 14 | 0 | 13 | 0 | 0 |
| 7 | B | 14 | 0 | 13 | 1 | 0 |

*Continued on next page...*

For Manuscript Review

*Continued from previous page...*

| Mol | Chain | Non-H | H(model) | H(added) | Clashes | Symm-Clashes |
|---|---|---|---|---|---|---|
| 8 | A | 4 | 0 | 6 | 0 | 0 |
| 8 | B | 12 | 0 | 18 | 2 | 0 |
| 9 | A | 1 | 0 | 0 | 1 | 0 |
| 9 | B | 1 | 0 | 0 | 0 | 0 |
| 9 | D | 1 | 0 | 0 | 0 | 0 |
| 10 | A | 21 | 0 | 0 | 0 | 0 |
| 10 | B | 60 | 0 | 0 | 0 | 0 |
| 10 | C | 48 | 0 | 0 | 0 | 0 |
| 10 | D | 8 | 0 | 0 | 0 | 0 |
| 10 | P | 6 | 0 | 0 | 0 | 0 |
| All | All | 6963 | 0 | 6496 | 55 | 0 |

The all-atom clashscore is defined as the number of clashes found per 1000 atoms (including hydrogen atoms). The all-atom clashscore for this structure is 4.

All (55) close contacts within the same asymmetric unit are listed below, sorted by their clash magnitude.

| Atom-1 | Atom-2 | Interatomic distance (Å) | Clash overlap (Å) |
|---|---|---|---|
| 3:C:97:GLN:HE22 | 5:P:5:ASN:HD21 | 1.22 | 0.85 |
| 3:C:236:ALA:HB1 | 4:D:12:ARG:HG3 | 1.66 | 0.76 |
| 2:B:37:SER:H | 2:B:40:LYS:HE2 | 1.58 | 0.69 |
| 2:B:2:THR:HG21 | 2:B:25:HIS:HB2 | 1.74 | 0.69 |
| 2:B:16:ALA:HB3 | 7:B:301:NAG:H82 | 1.75 | 0.69 |
| 1:A:157:ASP:HB3 | 1:A:160:VAL:HG22 | 1.77 | 0.65 |
| 1:A:155:SER:HB3 | 1:A:162:ILE:HD13 | 1.80 | 0.63 |
| 2:B:11:LEU:HD11 | 2:B:18:LYS:HD3 | 1.83 | 0.61 |
| 2:B:40:LYS:HE3 | 2:B:41:PRO:O | 2.01 | 0.60 |
| 3:C:214:THR:HB | 3:C:262:TYR:HB2 | 1.86 | 0.58 |
| 3:C:266:LEU:HD13 | 3:C:270:LEU:HD13 | 1.87 | 0.57 |
| 2:B:40:LYS:HG2 | 2:B:41:PRO:HD2 | 1.86 | 0.56 |
| 3:C:209:TYR:HB3 | 3:C:210:PRO:HD3 | 1.87 | 0.56 |
| 2:B:106:PRO:HB2 | 8:B:303:EDO:H11 | 1.88 | 0.56 |
| 3:C:266:LEU:HD22 | 3:C:270:LEU:HD11 | 1.89 | 0.54 |
| 2:B:121:PRO:HD3 | 2:B:229:PRO:HB3 | 1.92 | 0.51 |
| 3:C:5:MET:HB2 | 3:C:168:LEU:HD13 | 1.93 | 0.51 |
| 1:A:177:SER:HA | 9:A:303:CL:CL | 2.47 | 0.51 |
| 2:B:32:TYR:CE2 | 2:B:47:LEU:HD13 | 2.47 | 0.50 |
| 4:D:12:ARG:HB2 | 4:D:22:ILE:HD11 | 1.95 | 0.48 |
| 3:C:70:GLN:NE2 | 5:P:5:ASN:H | 2.11 | 0.48 |
| 2:B:210:GLN:HG3 | 2:B:233:ILE:HG23 | 1.95 | 0.48 |
| 3:C:70:GLN:HE22 | 5:P:5:ASN:H | 1.62 | 0.48 |

*Continued on next page...*

For Manuscript Review

*Continued from previous page...*

| Atom-1 | Atom-2 | Interatomic distance (Å) | Clash overlap (Å) |
|---|---|---|---|
| 3:C:111:ARG:HG2 | 3:C:113:TYR:CZ | 2.50 | 0.47 |
| 1:A:146:PHE:HE2 | 1:A:165:LYS:HD3 | 1.80 | 0.46 |
| 4:D:23:LEU:O | 4:D:67:HIS:HA | 2.15 | 0.46 |
| 2:B:129:PRO:HD3 | 2:B:142:LEU:HG | 1.98 | 0.46 |
| 1:A:49:ILE:HD13 | 1:A:66:LEU:HG | 1.98 | 0.45 |
| 3:C:44:ARG:HG3 | 3:C:64:THR:OG1 | 2.16 | 0.45 |
| 1:A:212:SER:HB2 | 2:B:131:GLU:HG2 | 1.99 | 0.45 |
| 4:D:50:GLU:HB2 | 4:D:67:HIS:CE1 | 2.52 | 0.44 |
| 1:A:66:LEU:HD22 | 1:A:75:ILE:HD11 | 2.00 | 0.43 |
| 3:C:189:VAL:HG22 | 3:C:203:CYS:HB3 | 2.00 | 0.43 |
| 3:C:207:GLY:HA2 | 3:C:240:THR:HB | 2.00 | 0.43 |
| 3:C:242:GLN:HE22 | 4:D:12:ARG:HA | 1.83 | 0.43 |
| 4:D:28:THR:HG22 | 4:D:63:TYR:HB2 | 2.00 | 0.43 |
| 3:C:14:ARG:HH21 | 3:C:19:GLU:HB3 | 1.84 | 0.43 |
| 3:C:67:ALA:O | 3:C:71:GLU:HG2 | 2.17 | 0.43 |
| 3:C:97:GLN:NE2 | 5:P:5:ASN:HD21 | 2.03 | 0.43 |
| 3:C:0:MET:HG3 | 3:C:1:GLY:H | 1.84 | 0.43 |
| 3:C:14:ARG:NH2 | 3:C:19:GLU:HB3 | 2.34 | 0.43 |
| 3:C:219:LEU:HB3 | 3:C:257:TYR:CD2 | 2.53 | 0.43 |
| 2:B:9:ARG:HG3 | 8:B:303:EDO:H12 | 2.01 | 0.43 |
| 1:A:175:PHE:CD2 | 2:B:139:LYS:HE2 | 2.54 | 0.42 |
| 1:A:14:MET:HE2 | 1:A:14:MET:HB3 | 1.95 | 0.42 |
| 4:D:27:VAL:HG21 | 4:D:37:ILE:HD13 | 2.01 | 0.42 |
| 2:B:208:ARG:NH2 | 2:B:210:GLN:HB2 | 2.33 | 0.42 |
| 3:C:191:HIS:NE2 | 3:C:199:VAL:HG11 | 2.35 | 0.42 |
| 2:B:40:LYS:HE2 | 2:B:40:LYS:HB3 | 1.76 | 0.42 |
| 2:B:68:PRO:HG2 | 2:B:72:LYS:HB3 | 2.02 | 0.41 |
| 4:D:22:ILE:O | 4:D:70:PHE:HE1 | 2.03 | 0.41 |
| 2:B:122:PRO:HB3 | 2:B:149:PHE:HB3 | 2.02 | 0.41 |
| 1:A:132:ASP:HB3 | 1:A:135:SER:O | 2.21 | 0.41 |
| 1:A:160:VAL:HG12 | 1:A:184:SER:HB2 | 2.03 | 0.41 |
| 3:C:111:ARG:CZ | 3:C:128:GLU:HG3 | 2.51 | 0.41 |

There are no symmetry-related clashes.

#### 5.3 Torsion angles (i)

##### 5.3.1 Protein backbone (i)

In the following table, the Percentiles column shows the percent Ramachandran outliers of the chain as a percentile score with respect to all X-ray entries followed by that with respect to entries

For Manuscript Review

of similar resolution.

The Analysed column shows the number of residues for which the backbone conformation was analysed, and the total number of residues.

| Mol | Chain | Analysed | Favoured | Allowed | Outliers | Percentiles | |
|---|---|---|---|---|---|---|---|
| 1 | A | 210/223 (94%) | 199 (95%) | 11 (5%) | 0 | 100 | 100 |
| 2 | B | 240/254 (94%) | 236 (98%) | 4 (2%) | 0 | 100 | 100 |
| 3 | C | 275/281 (98%) | 258 (94%) | 15 (6%) | 2 (1%) | 19 | 16 |
| 4 | D | 98/100 (98%) | 95 (97%) | 3 (3%) | 0 | 100 | 100 |
| 5 | P | 7/9 (78%) | 6 (86%) | 1 (14%) | 0 | 100 | 100 |
| All | All | 830/867 (96%) | 794 (96%) | 34 (4%) | 2 (0%) | 44 | 45 |

All (2) Ramachandran outliers are listed below:

| Mol | Chain | Res | Type |
|---|---|---|---|
| 3 | C | 2 | PRO |
| 3 | C | 1 | GLY |

##### 5.3.2 Protein sidechains ⓘ

In the following table, the Percentiles column shows the percent sidechain outliers of the chain as a percentile score with respect to all X-ray entries followed by that with respect to entries of similar resolution.

The Analysed column shows the number of residues for which the sidechain conformation was analysed, and the total number of residues.

| Mol | Chain | Analysed | Rotameric | Outliers | Percentiles | |
|---|---|---|---|---|---|---|
| 1 | A | 189/201 (94%) | 183 (97%) | 6 (3%) | 34 | 37 |
| 2 | B | 214/225 (95%) | 212 (99%) | 2 (1%) | 75 | 82 |
| 3 | C | 229/239 (96%) | 223 (97%) | 6 (3%) | 41 | 46 |
| 4 | D | 94/95 (99%) | 94 (100%) | 0 | 100 | 100 |
| 5 | P | 8/8 (100%) | 8 (100%) | 0 | 100 | 100 |
| All | All | 734/768 (96%) | 720 (98%) | 14 (2%) | 52 | 59 |

All (14) residues with a non-rotameric sidechain are listed below:

| Mol | Chain | Res | Type |
|---|---|---|---|
| 1 | A | 44 | GLU |
| 1 | A | 131 | ARG |

*Continued on next page...*

For Manuscript Review

*Continued from previous page...*

| Mol | Chain | Res | Type |
|---|---|---|---|
| 1 | A | 137 | ASP |
| 1 | A | 166 | CYS |
| 1 | A | 173 | MET |
| 1 | A | 206 | PHE |
| 2 | B | 183 | ASN |
| 2 | B | 192 | ARG |
| 3 | C | 81 | LEU |
| 3 | C | 170 | ARG |
| 3 | C | 183 | ASP |
| 3 | C | 196 | LYS |
| 3 | C | 219 | LEU |
| 3 | C | 272 | LEU |

Sometimes sidechains can be flipped to improve hydrogen bonding and reduce clashes. All (5) such sidechains are listed below:

| Mol | Chain | Res | Type |
|---|---|---|---|
| 2 | B | 183 | ASN |
| 3 | C | 70 | GLN |
| 3 | C | 86 | ASN |
| 3 | C | 97 | GLN |
| 3 | C | 188 | HIS |

##### 5.3.3 RNA (i)

There are no RNA molecules in this entry.

#### 5.4 Non-standard residues in protein, DNA, RNA chains (i)

There are no non-standard protein/DNA/RNA residues in this entry.

#### 5.5 Carbohydrates (i)

2 monosaccharides are modelled in this entry.

In the following table, the Counts columns list the number of bonds (or angles) for which Mogul statistics could be retrieved, the number of bonds (or angles) that are observed in the model and the number of bonds (or angles) that are defined in the Chemical Component Dictionary. The Link column lists molecule types, if any, to which the group is linked. The Z score for a bond length (or angle) is the number of standard deviations the observed value is removed from the

For Manuscript Review

expected value. A bond length (or angle) with $|Z| > 2$ is considered an outlier worth inspection. RMSZ is the root-mean-square of all Z scores of the bond lengths (or angles).

| Mol | Type | Chain | Res | Link | Bond lengths | | | Bond angles | | |
|---|---|---|---|---|---|---|---|---|---|---|
| | | | | | Counts | RMSZ | $\#\|Z\| > 2$ | Counts | RMSZ | $\#\|Z\| > 2$ |
| 6 | NAG | E | 1 | 2,6 | 14,14,15 | 0.72 | 0 | 17,19,21 | 1.01 | 1 (5%) |
| 6 | NAG | E | 2 | 6 | 14,14,15 | 0.72 | 0 | 17,19,21 | 0.78 | 0 |

In the following table, the Chirals column lists the number of chiral outliers, the number of chiral centers analysed, the number of these observed in the model and the number defined in the Chemical Component Dictionary. Similar counts are reported in the Torsion and Rings columns. '-' means no outliers of that kind were identified.

| Mol | Type | Chain | Res | Link | Chirals | Torsions | Rings |
|---|---|---|---|---|---|---|---|
| 6 | NAG | E | 1 | 2,6 | - | 0/6/23/26 | 0/1/1/1 |
| 6 | NAG | E | 2 | 6 | - | 0/6/23/26 | 0/1/1/1 |

There are no bond length outliers.

All (1) bond angle outliers are listed below:

| Mol | Chain | Res | Type | Atoms | Z | Observed($^o$) | Ideal($^o$) |
|---|---|---|---|---|---|---|---|
| 6 | E | 1 | NAG | C1-O5-C5 | 2.17 | 115.09 | 112.19 |

There are no chirality outliers.

There are no torsion outliers.

There are no ring outliers.

No monomer is involved in short contacts.

The following is a two-dimensional graphical depiction of Mogul quality analysis of bond lengths, bond angles, torsion angles, and ring geometry for oligosaccharide.

For Manuscript Review

**Oligosaccharide Chain E**

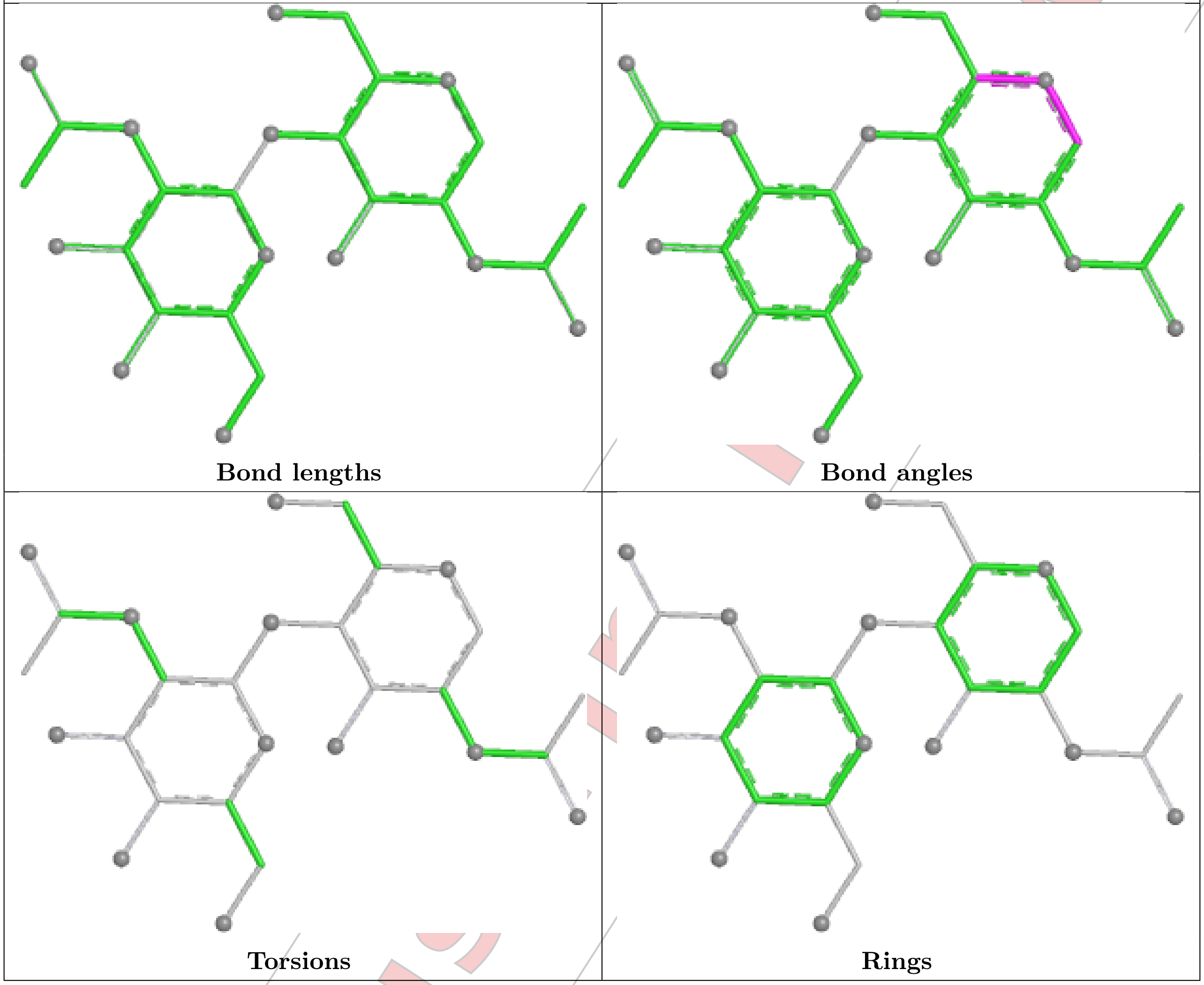


#### 5.6 Ligand geometry (i)

Of 9 ligands modelled in this entry, 3 are monoatomic - leaving 6 for Mogul analysis.

In the following table, the Counts columns list the number of bonds (or angles) for which Mogul statistics could be retrieved, the number of bonds (or angles) that are observed in the model and the number of bonds (or angles) that are defined in the Chemical Component Dictionary. The Link column lists molecule types, if any, to which the group is linked. The Z score for a bond length (or angle) is the number of standard deviations the observed value is removed from the expected value. A bond length (or angle) with $|Z| > 2$ is considered an outlier worth inspection. RMSZ is the root-mean-square of all Z scores of the bond lengths (or angles).

| Mol | Type | Chain | Res | Link | Bond lengths | | | Bond angles | | |
|---|---|---|---|---|---|---|---|---|---|---|
| | | | | | Counts | RMSZ | $\#\|Z\| > 2$ | Counts | RMSZ | $\#\|Z\| > 2$ |
| 8 | EDO | B | 302 | - | 3,3,3 | 0.25 | 0 | 2,2,2 | 0.33 | 0 |
| 7 | NAG | B | 301 | 2 | 14,14,15 | 0.74 | 0 | 17,19,21 | 0.95 | 1 (5%) |
| 8 | EDO | A | 302 | - | 3,3,3 | 0.25 | 0 | 2,2,2 | 0.34 | 0 |

For Manuscript Review

| Mol | Type | Chain | Res | Link | Bond lengths | | | Bond angles | | |
|---|---|---|---|---|---|---|---|---|---|---|
| | | | | | Counts | RMSZ | #\|Z\| > 2 | Counts | RMSZ | #\|Z\| > 2 |
| 8 | EDO | B | 303 | - | 3,3,3 | 0.26 | 0 | 2,2,2 | 0.29 | 0 |
| 7 | NAG | A | 301 | 1 | 14,14,15 | 0.66 | 0 | 17,19,21 | 0.95 | 0 |
| 8 | EDO | B | 304 | - | 3,3,3 | 0.25 | 0 | 2,2,2 | 0.37 | 0 |

In the following table, the Chirals column lists the number of chiral outliers, the number of chiral centers analysed, the number of these observed in the model and the number defined in the Chemical Component Dictionary. Similar counts are reported in the Torsion and Rings columns. '-' means no outliers of that kind were identified.

| Mol | Type | Chain | Res | Link | Chirals | Torsions | Rings |
|---|---|---|---|---|---|---|---|
| 8 | EDO | B | 302 | - | - | 0/1/1/1 | - |
| 7 | NAG | B | 301 | 2 | - | 1/6/23/26 | 0/1/1/1 |
| 8 | EDO | A | 302 | - | - | 0/1/1/1 | - |
| 8 | EDO | B | 303 | - | - | 1/1/1/1 | - |
| 7 | NAG | A | 301 | 1 | - | 0/6/23/26 | 0/1/1/1 |
| 8 | EDO | B | 304 | - | - | 0/1/1/1 | - |

There are no bond length outliers.

All (1) bond angle outliers are listed below:

| Mol | Chain | Res | Type | Atoms | Z | Observed($^o$) | Ideal($^o$) |
|---|---|---|---|---|---|---|---|
| 7 | B | 301 | NAG | C1-O5-C5 | 2.31 | 115.28 | 112.19 |

There are no chirality outliers.

All (2) torsion outliers are listed below:

| Mol | Chain | Res | Type | Atoms |
|---|---|---|---|---|
| 7 | B | 301 | NAG | O5-C5-C6-O6 |
| 8 | B | 303 | EDO | O1-C1-C2-O2 |

There are no ring outliers.

2 monomers are involved in 3 short contacts:

| Mol | Chain | Res | Type | Clashes | Symm-Clashes |
|---|---|---|---|---|---|
| 7 | B | 301 | NAG | 1 | 0 |
| 8 | B | 303 | EDO | 2 | 0 |

The following is a two-dimensional graphical depiction of Mogul quality analysis of bond lengths, bond angles, torsion angles, and ring geometry for all instances of the Ligand of Interest. In addition, ligands with molecular weight > 250 and outliers as shown on the validation Tables will also be included. For torsion angles, if less then 5% of the Mogul distribution of torsion angles is

For Manuscript Review

within 10 degrees of the torsion angle in question, then that torsion angle is considered an outlier. Any bond that is central to one or more torsion angles identified as an outlier by Mogul will be highlighted in the graph. For rings, the root-mean-square deviation (RMSD) between the ring in question and similar rings identified by Mogul is calculated over all ring torsion angles. If the average RMSD is greater than 60 degrees and the minimal RMSD between the ring in question and any Mogul-identified rings is also greater than 60 degrees, then that ring is considered an outlier. The outliers are highlighted in purple. The color gray indicates Mogul did not find sufficient equivalents in the CSD to analyse the geometry.

**Ligand NAG B 301**

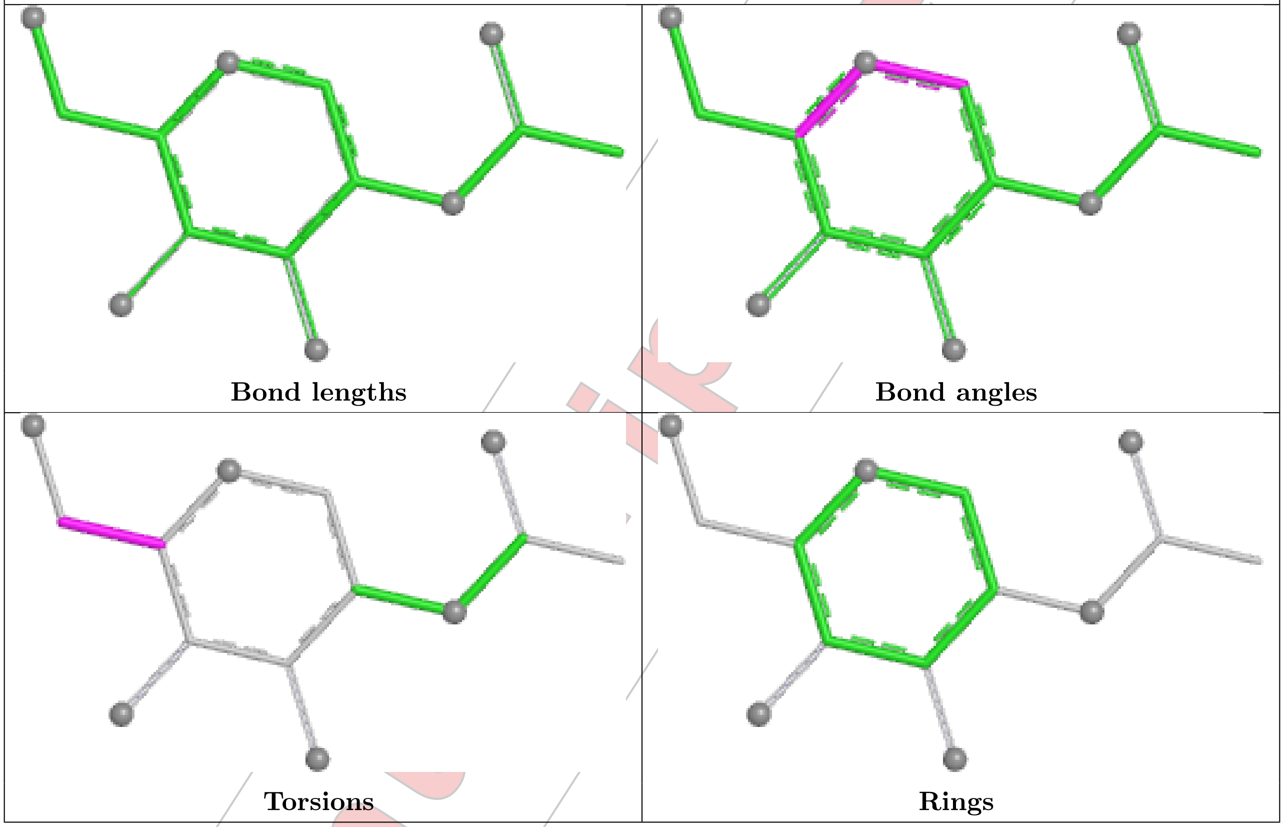


For Manuscript Review

**Ligand NAG A 301**

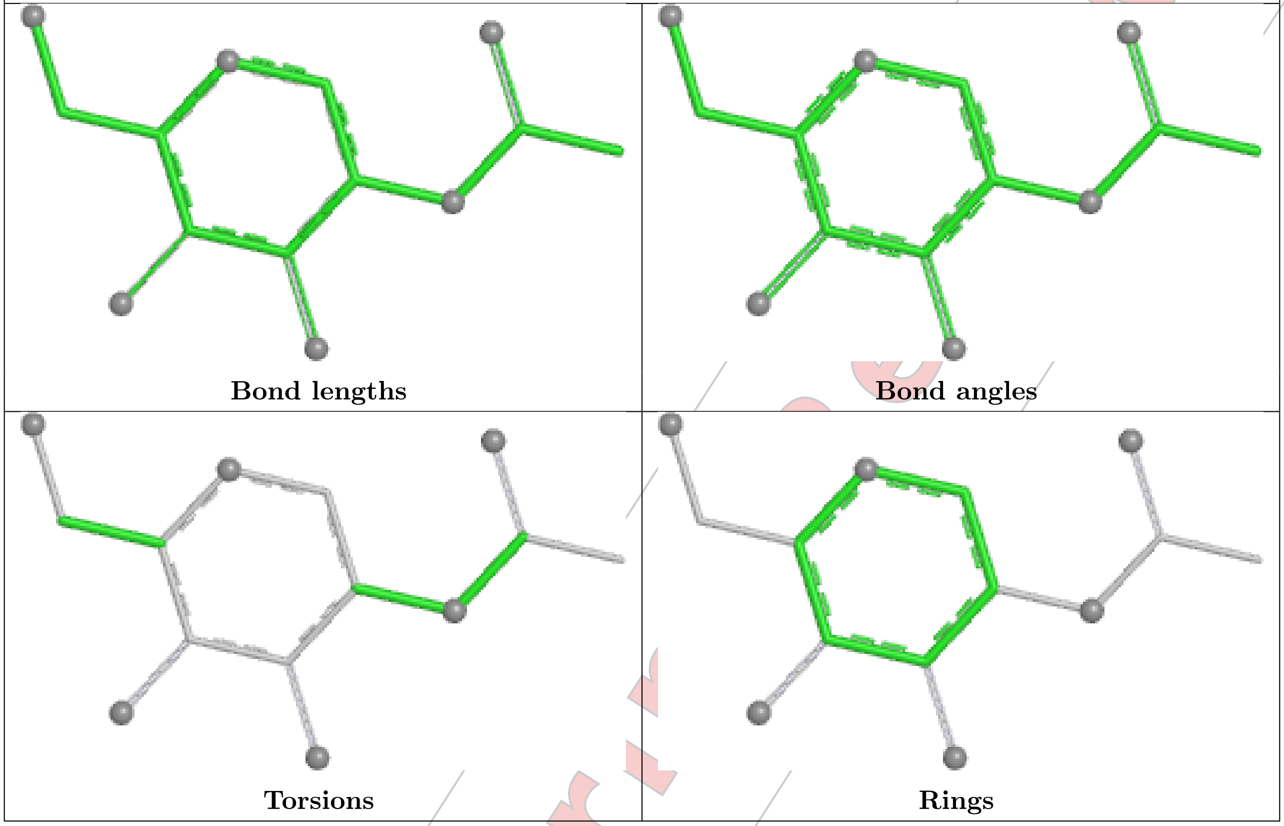


#### 5.7 Other polymers (i)

There are no such residues in this entry.

#### 5.8 Polymer linkage issues (i)

There are no chain breaks in this entry.

For Manuscript Review

### 6 Fit of model and data (i)

#### 6.1 Protein, DNA and RNA chains (i)

In the following table, the column labelled '#RSRZ> 2' contains the number (and percentage) of RSRZ outliers, followed by percent RSRZ outliers for the chain as percentile scores relative to all X-ray entries and entries of similar resolution. The OWAB column contains the minimum, median, $95^{th}$ percentile and maximum values of the occupancy-weighted average B-factor per residue. The column labelled 'Q< 0.9' lists the number of (and percentage) of residues with an average occupancy less than 0.9.

| Mol | Chain | Analysed | <RSRZ> | #RSRZ>2 | | | OWAB(Å$^2$) | Q<0.9 |
|---|---|---|---|---|---|---|---|---|
| 1 | A | 212/223 (95%) | 0.61 | 13 (6%) | 28 | 30 | 30, 65, 125, 148 | 0 |
| 2 | B | 242/254 (95%) | 0.13 | 4 (1%) | 69 | 70 | 29, 50, 77, 123 | 0 |
| 3 | C | 277/281 (98%) | 0.61 | 37 (13%) | 8 | 9 | 30, 58, 152, 165 | 0 |
| 4 | D | 100/100 (100%) | 0.67 | 5 (5%) | 35 | 37 | 49, 64, 104, 106 | 0 |
| 5 | P | 9/9 (100%) | -0.04 | 1 (11%) | 12 | 12 | 30, 33, 45, 49 | 0 |
| All | All | 840/867 (96%) | 0.47 | 60 (7%) | 23 | 25 | 29, 56, 134, 165 | 0 |

All (60) RSRZ outliers are listed below:

| Mol | Chain | Res | Type | RSRZ |
|---|---|---|---|---|
| 3 | C | 276 | PRO | 4.4 |
| 1 | A | 194 | ALA | 4.1 |
| 3 | C | 2 | PRO | 4.1 |
| 3 | C | 209 | TYR | 3.9 |
| 1 | A | 195 | PHE | 3.9 |
| 1 | A | 199 | ILE | 3.9 |
| 3 | C | 180 | LEU | 3.9 |
| 1 | A | 160 | VAL | 3.7 |
| 2 | B | 243 | ASP | 3.7 |
| 3 | C | 215 | LEU | 3.3 |
| 2 | B | 2 | THR | 3.3 |
| 1 | A | 212 | SER | 3.2 |
| 1 | A | 1 | ALA | 3.2 |
| 3 | C | 89 | ALA | 3.0 |
| 3 | C | 248 | VAL | 2.9 |
| 3 | C | 271 | THR | 2.9 |
| 3 | C | 210 | PRO | 2.8 |
| 1 | A | 162 | ILE | 2.8 |
| 4 | D | 1 | ILE | 2.8 |
| 3 | C | 0 | MET | 2.8 |

*Continued on next page...*

For Manuscript Review

*Continued from previous page...*

| Mol | Chain | Res | Type | RSRZ |
|---|---|---|---|---|
| 3 | C | 236 | ALA | 2.8 |
| 3 | C | 203 | CYS | 2.7 |
| 4 | D | 49 | VAL | 2.7 |
| 3 | C | 177 | ALA | 2.7 |
| 3 | C | 225 | THR | 2.7 |
| 3 | C | 185 | PRO | 2.7 |
| 3 | C | 274 | TRP | 2.7 |
| 1 | A | 198 | SER | 2.6 |
| 3 | C | 91 | GLY | 2.6 |
| 4 | D | 22 | ILE | 2.6 |
| 3 | C | 179 | LEU | 2.6 |
| 3 | C | 227 | ASP | 2.5 |
| 2 | B | 242 | ALA | 2.5 |
| 4 | D | 54 | MET | 2.5 |
| 3 | C | 201 | LEU | 2.5 |
| 1 | A | 191 | CYS | 2.5 |
| 3 | C | 1 | GLY | 2.4 |
| 3 | C | 138 | MET | 2.4 |
| 3 | C | 224 | LEU | 2.4 |
| 3 | C | 270 | LEU | 2.4 |
| 3 | C | 267 | PRO | 2.4 |
| 3 | C | 257 | TYR | 2.4 |
| 3 | C | 90 | GLY | 2.4 |
| 3 | C | 189 | VAL | 2.3 |
| 4 | D | 0 | MET | 2.3 |
| 2 | B | 181 | ALA | 2.3 |
| 3 | C | 178 | THR | 2.2 |
| 3 | C | 239 | GLY | 2.2 |
| 3 | C | 231 | VAL | 2.2 |
| 3 | C | 197 | GLY | 2.2 |
| 3 | C | 237 | GLY | 2.2 |
| 3 | C | 187 | ALA | 2.1 |
| 1 | A | 166 | CYS | 2.1 |
| 3 | C | 272 | LEU | 2.1 |
| 1 | A | 156 | LYS | 2.1 |
| 1 | A | 155 | SER | 2.1 |
| 5 | P | 1 | ALA | 2.0 |
| 1 | A | 197 | ASN | 2.0 |
| 3 | C | 39 | ASP | 2.0 |
| 3 | C | 3 | HIS | 2.0 |

For Manuscript Review

#### 6.2 Non-standard residues in protein, DNA, RNA chains (i)

There are no non-standard protein/DNA/RNA residues in this entry.

#### 6.3 Carbohydrates (i)

SUGAR-RSR INFOmissingINFO

#### 6.4 Ligands (i)

In the following table, the Atoms column lists the number of modelled atoms in the group and the number defined in the chemical component dictionary. The B-factors column lists the minimum, median, $95^{th}$ percentile and maximum values of B factors of atoms in the group. The column labelled 'Q< 0.9' lists the number of atoms with occupancy less than 0.9.

| Mol | Type | Chain | Res | Atoms | RSCC | RSR | B-factors($Å^2$) | Q<0.9 |
|---|---|---|---|---|---|---|---|---|
| 7 | NAG | B | 301 | 14/15 | 0.71 | 0.13 | 76,83,90,91 | 0 |
| 8 | EDO | B | 302 | 4/4 | 0.76 | 0.16 | 80,81,84,86 | 0 |
| 8 | EDO | B | 304 | 4/4 | 0.77 | 0.16 | 51,53,55,57 | 0 |
| 8 | EDO | A | 302 | 4/4 | 0.80 | 0.20 | 53,54,56,58 | 0 |
| 8 | EDO | B | 303 | 4/4 | 0.86 | 0.17 | 41,46,46,49 | 0 |
| 7 | NAG | A | 301 | 14/15 | 0.86 | 0.10 | 41,45,48,53 | 0 |
| 9 | CL | B | 305 | 1/1 | 0.87 | 0.12 | 74,74,74,74 | 0 |
| 9 | CL | A | 303 | 1/1 | 0.91 | 0.09 | 76,76,76,76 | 0 |
| 9 | CL | D | 101 | 1/1 | 0.92 | 0.11 | 78,78,78,78 | 0 |

The following is a graphical depiction of the model fit to experimental electron density of all instances of the Ligand of Interest. In addition, ligands with molecular weight > 250 and outliers as shown on the geometry validation Tables will also be included. Each fit is shown from different orientation to approximate a three-dimensional view.

For Manuscript Review

**Electron density around NAG B 301:**

$2mF_o$-$DF_c$ (at 0.7 rmsd) in gray
$mF_o$-$DF_c$ (at 3 rmsd) in purple (negative)
and green (positive)

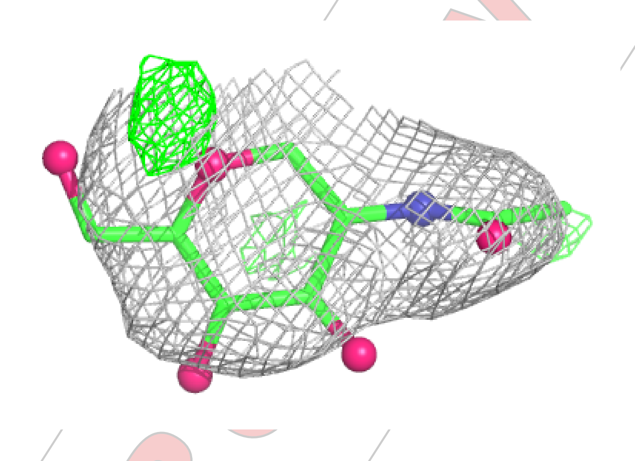

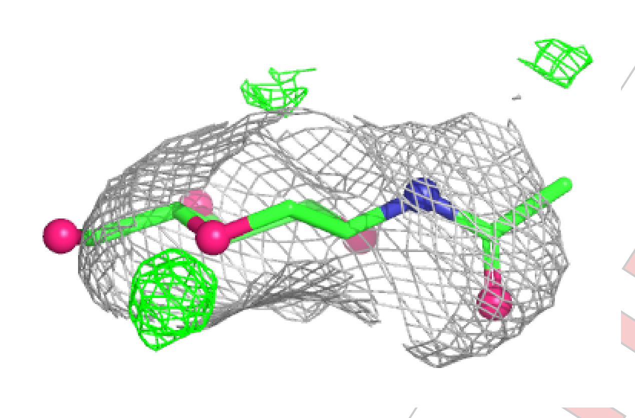

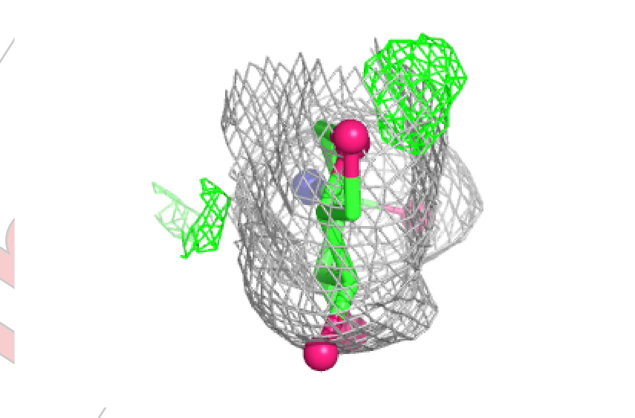

For Manuscript Review

**Electron density around NAG A 301:**

$2mF_o$-$DF_c$ (at 0.7 rmsd) in gray
$mF_o$-$DF_c$ (at 3 rmsd) in purple (negative)
and green (positive)

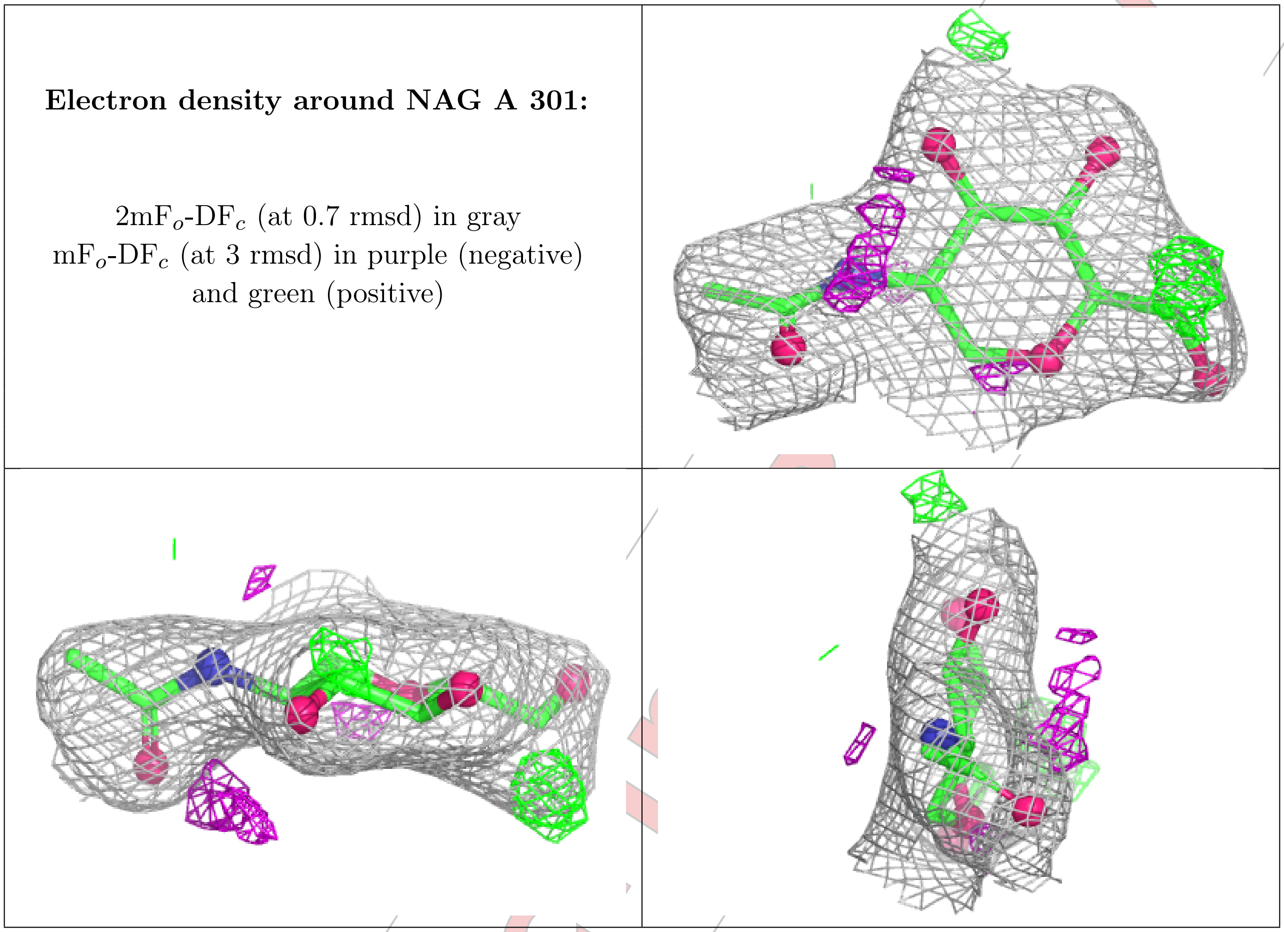

#### 6.5 Other polymers ⓘ

There are no such residues in this entry.

For Manuscript Review

### Full wwPDB X-ray Structure Validation Report (i)

Mar 24, 2025 – 02:37 PM EDT

| | | |
|---|---|---|
| PDB ID | : | 9NW2 |
| Title | : | Crystal structure of NP204 TCR in complex with NP366-H-2Db |
| Deposited on | : | 2025-03-21 |
| Resolution | : | 4.44 Å(reported) |

**This wwPDB validation report is for manuscript review**

This is a Full wwPDB X-ray Structure Validation Report.

This report is produced by the wwPDB biocuration pipeline after annotation of the structure.

We welcome your comments at **
A user guide is available at
https://www.wwpdb.org/validation/2017/XrayValidationReportHelp
with specific help available everywhere you see the (i) symbol.

The types of validation reports are described at
http://www.wwpdb.org/validation/2017/FAQs#types.

The following versions of software and data (see references (i)) were used in the production of this report:

| | | |
|---|---|---|
| MolProbity | : | 4.02b-467 |
| Xtriage (Phenix) | : | 1.21 |
| EDS | : | 3.0 |
| Percentile statistics | : | 20231227.v01 (using entries in the PDB archive December 27th 2023) |
| CCP4 | : | 9.0.004 (Gargrove) |
| Density-Fitness | : | 1.0.11 |
| Ideal geometry (proteins) | : | Engh & Huber (2001) |
| Ideal geometry (DNA, RNA) | : | Parkinson et al. (1996) |
| Validation Pipeline (wwPDB-VP) | : | 2.41.5 |

For Manuscript Review

### 1 Overall quality at a glance (i)

The following experimental techniques were used to determine the structure:
*X-RAY DIFFRACTION*

The reported resolution of this entry is 4.44 Å.

Percentile scores (ranging between 0-100) for global validation metrics of the entry are shown in the following graphic. The table shows the number of entries on which the scores are based.

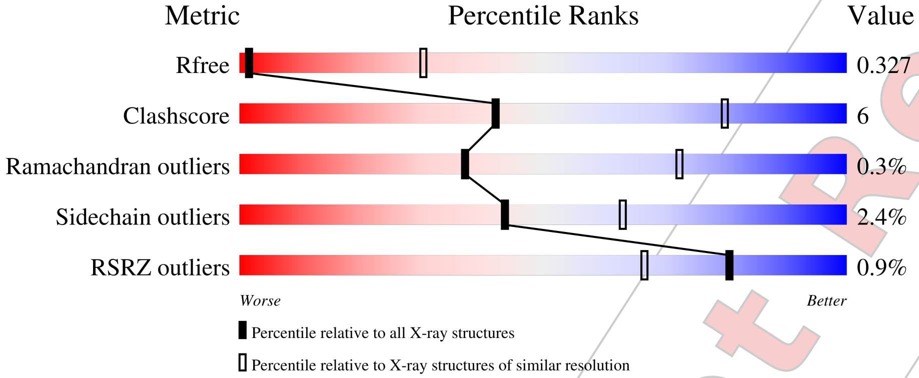


| Metric | Whole archive (#Entries) | Similar resolution (#Entries, resolution range(Å)) |
|---|---|---|
| $R_{free}$ | 164625 | 1018 (4.98-3.90) |
| Clashscore | 180529 | 1071 (4.98-3.90) |
| Ramachandran outliers | 177936 | 1017 (5.00-3.88) |
| Sidechain outliers | 177891 | 1001 (5.00-3.88) |
| RSRZ outliers | 164620 | 1015 (4.98-3.90) |

The table below summarises the geometric issues observed across the polymeric chains and their fit to the electron density. The red, orange, yellow and green segments of the lower bar indicate the fraction of residues that contain outliers for >=3, 2, 1 and 0 types of geometric quality criteria respectively. A grey segment represents the fraction of residues that are not modelled. The numeric value for each fraction is indicated below the corresponding segment, with a dot representing fractions <=5% The upper red bar (where present) indicates the fraction of residues that have poor fit to the electron density. The numeric value is given above the bar.

| Mol | Chain | Length | Quality of chain |
|---|---|---|---|
| 1 | A | 214 | 45% 8% 47% |
| 1 | F | 214 | 44% 8% 48% |
| 2 | B | 253 | % / 68% 19% • 10% |
| 2 | G | 253 | % / 69% 18% • 11% |
| 3 | C | 281 | % / 91% 7% • |

*Continued on next page...*

For Manuscript Review

*Continued from previous page...*

| Mol | Chain | Length | Quality of chain |
|---|---|---|---|
| 3 | H | 281 | 90% 8% • |
| 4 | D | 100 | 2% 81% 17% • |
| 4 | I | 100 | % 82% 15% • • |
| 5 | E | 9 | 67% 33% |
| 5 | J | 9 | 78% 22% |

For Manuscript Review

### 2 Entry composition (i)

There are 5 unique types of molecules in this entry. The entry contains 10901 atoms, of which 0 are hydrogens and 0 are deuteriums.

In the tables below, the ZeroOcc column contains the number of atoms modelled with zero occupancy, the AltConf column contains the number of residues with at least one atom in alternate conformation and the Trace column contains the number of residues modelled with at most 2 atoms.

- Molecule 1 is a protein called NP204 TCR alpha-chain.

| Mol | Chain | Residues | Atoms | ZeroOcc | AltConf | Trace |
|---|---|---|---|---|---|---|
| 1 | A | 114 | Total 822, C 509, N 144, O 166, S 3 | 0 | 0 | 0 |
| 1 | F | 112 | Total 827, C 515, N 145, O 164, S 3 | 0 | 0 | 0 |

- Molecule 2 is a protein called NP204 TCR beta-chain.

| Mol | Chain | Residues | Atoms | ZeroOcc | AltConf | Trace |
|---|---|---|---|---|---|---|
| 2 | B | 227 | Total 1760, C 1101, N 326, O 325, S 8 | 0 | 0 | 0 |
| 2 | G | 224 | Total 1734, C 1083, N 317, O 327, S 7 | 0 | 0 | 0 |

- Molecule 3 is a protein called H-2 class I histocompatibility antigen, D-B alpha chain.

| Mol | Chain | Residues | Atoms | ZeroOcc | AltConf | Trace |
|---|---|---|---|---|---|---|
| 3 | C | 278 | Total 2035, C 1278, N 356, O 392, S 9 | 0 | 0 | 0 |
| 3 | H | 277 | Total 2043, C 1283, N 361, O 390, S 9 | 0 | 0 | 0 |

There are 2 discrepancies between the modelled and reference sequences:

| Chain | Residue | Modelled | Actual | Comment | Reference |
|---|---|---|---|---|---|
| C | 0 | MET | - | initiating methionine | UNP P01899 |
| H | 0 | MET | - | initiating methionine | UNP P01899 |

- Molecule 4 is a protein called Beta-2-microglobulin.

| Mol | Chain | Residues | Atoms | ZeroOcc | AltConf | Trace |
|---|---|---|---|---|---|---|
| 4 | D | 98 | Total 769, C 493, N 124, O 146, S 6 | 0 | 0 | 0 |

*Continued on next page...*

For Manuscript Review

*Continued from previous page...*

| Mol | Chain | Residues | Atoms | | | | | ZeroOcc | AltConf | Trace |
|---|---|---|---|---|---|---|---|---|---|---|
| | | | Total | C | N | O | S | | | |
| 4 | I | 98 | 773 | 496 | 125 | 146 | 6 | 0 | 0 | 0 |

There are 4 discrepancies between the modelled and reference sequences:

| Chain | Residue | Modelled | Actual | Comment | Reference |
|---|---|---|---|---|---|
| D | 0 | MET | - | initiating methionine | UNP P01887 |
| D | 85 | ASP | ALA | conflict | UNP P01887 |
| I | 0 | MET | - | initiating methionine | UNP P01887 |
| I | 85 | ASP | ALA | conflict | UNP P01887 |

- Molecule 5 is a protein called Nucleoprotein peptide.

| Mol | Chain | Residues | Atoms | | | | | ZeroOcc | AltConf | Trace |
|---|---|---|---|---|---|---|---|---|---|---|
| | | | Total | C | N | O | S | | | |
| 5 | E | 9 | 69 | 38 | 11 | 18 | 2 | 0 | 0 | 0 |
| 5 | J | 9 | 69 | 38 | 11 | 18 | 2 | 0 | 0 | 0 |

For Manuscript Review

### 3 Residue-property plots (i)

These plots are drawn for all protein, RNA, DNA and oligosaccharide chains in the entry. The first graphic for a chain summarises the proportions of the various outlier classes displayed in the second graphic. The second graphic shows the sequence view annotated by issues in geometry and electron density. Residues are color-coded according to the number of geometric quality criteria for which they contain at least one outlier: green = 0, yellow = 1, orange = 2 and red = 3 or more. A red dot above a residue indicates a poor fit to the electron density (RSRZ > 2). Stretches of 2 or more consecutive residues without any outlier are shown as a green connector. Residues present in the sample, but not in the model, are shown in grey.

- Molecule 1: NP204 TCR alpha-chain

Chain A:

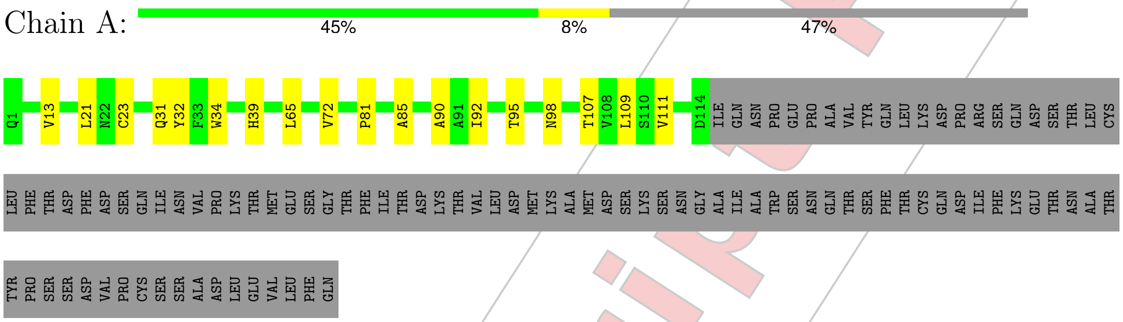


- Molecule 1: NP204 TCR alpha-chain

Chain F:

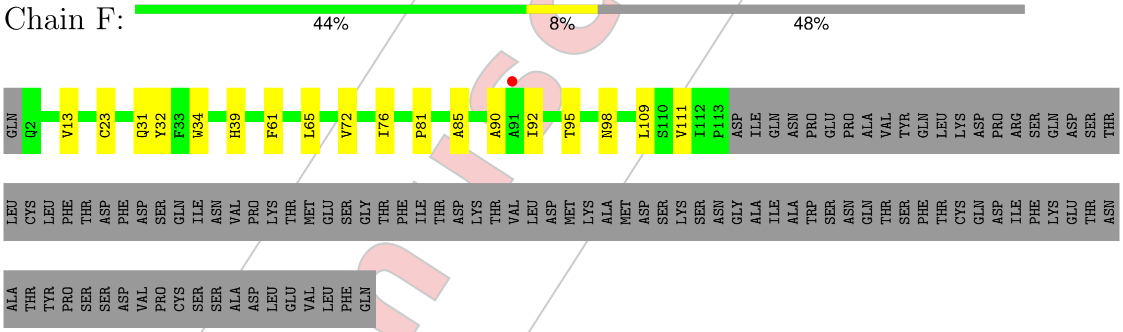


- Molecule 2: NP204 TCR beta-chain

Chain B:

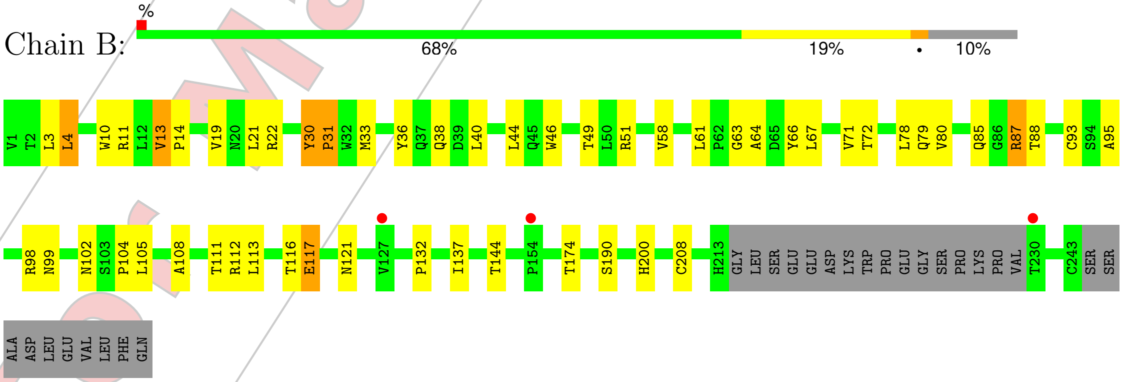


- Molecule 2: NP204 TCR beta-chain

For Manuscript Review

Chain G:

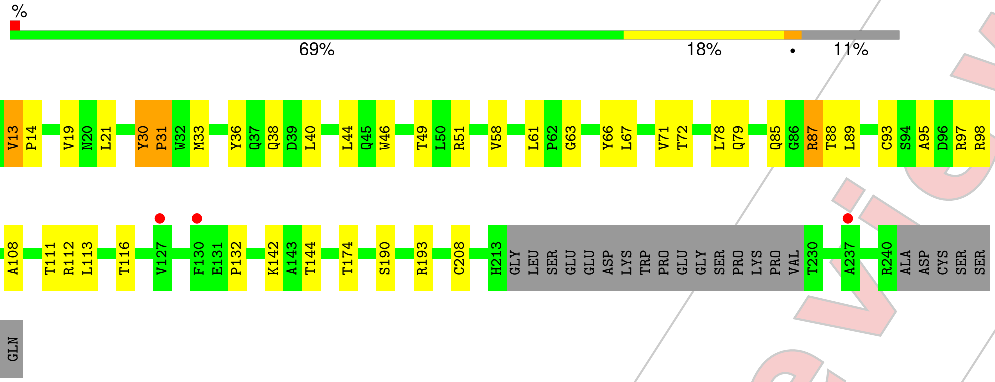


• Molecule 3: H-2 class I histocompatibility antigen, D-B alpha chain

Chain C:

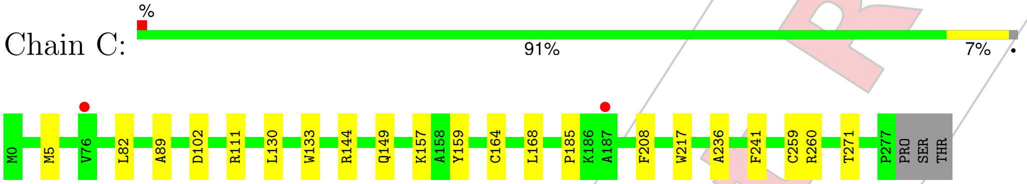


• Molecule 3: H-2 class I histocompatibility antigen, D-B alpha chain

Chain H:

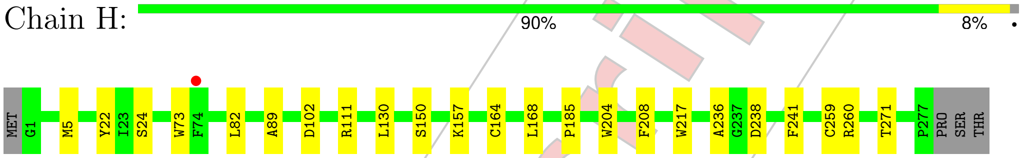


• Molecule 4: Beta-2-microglobulin

Chain D:

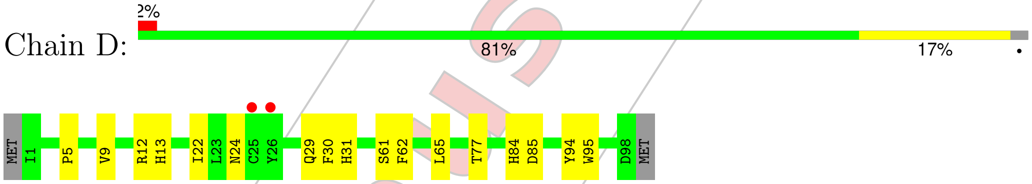


• Molecule 4: Beta-2-microglobulin

Chain I:

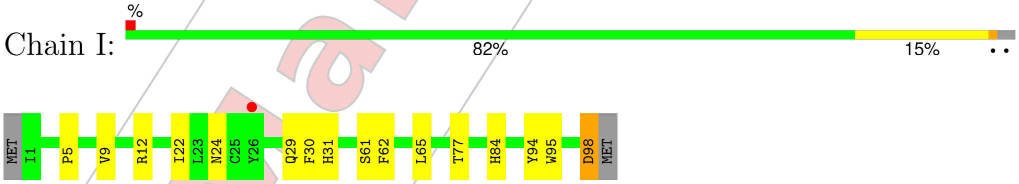


• Molecule 5: Nucleoprotein peptide

Chain E:

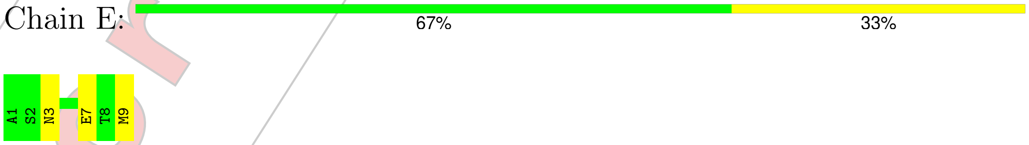


• Molecule 5: Nucleoprotein peptide

For Manuscript Review

Chain J:

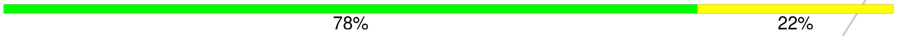


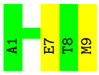


For Manuscript Review

### 4 Data and refinement statistics (i)

| Property | Value | Source |
|---|---|---|
| Space group | P 21 21 21 | Depositor |
| Cell constants<br>a, b, c, $\alpha$, $\beta$, $\gamma$ | 91.09Å 146.42Å 153.98Å<br>90.00° 90.00° 90.00° | Depositor |
| Resolution (Å) | 43.67 – 4.44<br>43.67 – 4.44 | Depositor<br>EDS |
| % Data completeness<br>(in resolution range) | 97.5 (43.67-4.44)<br>97.4 (43.67-4.44) | Depositor<br>EDS |
| $R_{merge}$ | 0.17 | Depositor |
| $R_{sym}$ | (Not available) | Depositor |
| $< I/\sigma(I) >$ [1] | 1.57 (at 4.45Å) | Xtriage |
| Refinement program | PHENIX 1.21.2_5419 | Depositor |
| R, $R_{free}$ | 0.294 , 0.328<br>0.295 , 0.327 | Depositor<br>DCC |
| $R_{free}$ test set | 572 reflections (4.33%) | wwPDB-VP |
| Wilson B-factor (Å$^2$) | 171.3 | Xtriage |
| Anisotropy | 0.329 | Xtriage |
| Bulk solvent $k_{sol}$(e/Å$^3$), $B_{sol}$(Å$^2$) | 0.23 , 230.4 | EDS |
| L-test for twinning[2] | $< \|L\| > = 0.41$, $< L^2 > = 0.24$ | Xtriage |
| Estimated twinning fraction | 0.000 for -h,l,k | Xtriage |
| $F_o$,$F_c$ correlation | 0.89 | EDS |
| Total number of atoms | 10901 | wwPDB-VP |
| Average B, all atoms (Å$^2$) | 240.0 | wwPDB-VP |

Xtriage's analysis on translational NCS is as follows: *The analyses of the Patterson function reveals a significant off-origin peak that is 46.05 % of the origin peak, indicating pseudo-translational symmetry. The chance of finding a peak of this or larger height randomly in a structure without pseudo-translational symmetry is equal to 1.2100e-04. The detected translational NCS is most likely also responsible for the elevated intensity ratio.*

[1] Intensities estimated from amplitudes.

[2] Theoretical values of $< |L| >$, $< L^2 >$ for acentric reflections are 0.5, 0.333 respectively for untwinned datasets, and 0.375, 0.2 for perfectly twinned datasets.

For Manuscript Review

### 5 Model quality (i)

#### 5.1 Standard geometry (i)

The Z score for a bond length (or angle) is the number of standard deviations the observed value is removed from the expected value. A bond length (or angle) with $|Z| > 5$ is considered an outlier worth inspection. RMSZ is the root-mean-square of all Z scores of the bond lengths (or angles).

| Mol | Chain | Bond lengths | | Bond angles | |
|---|---|---|---|---|---|
| | | RMSZ | #\|Z\| >5 | RMSZ | #\|Z\| >5 |
| 1 | A | 0.24 | 0/841 | 0.45 | 0/1146 |
| 1 | F | 0.24 | 0/847 | 0.45 | 0/1150 |
| 2 | B | 0.24 | 0/1804 | 0.51 | 0/2459 |
| 2 | G | 0.24 | 0/1778 | 0.50 | 0/2426 |
| 3 | C | 0.24 | 0/2090 | 0.46 | 0/2859 |
| 3 | H | 0.24 | 0/2098 | 0.46 | 0/2867 |
| 4 | D | 0.24 | 0/795 | 0.44 | 0/1091 |
| 4 | I | 0.24 | 0/799 | 0.45 | 0/1095 |
| 5 | E | 0.24 | 0/68 | 0.49 | 0/88 |
| 5 | J | 0.25 | 0/68 | 0.46 | 0/88 |
| All | All | 0.24 | 0/11188 | 0.47 | 0/15269 |

There are no bond length outliers.

There are no bond angle outliers.

There are no chirality outliers.

There are no planarity outliers.

#### 5.2 Too-close contacts (i)

In the following table, the Non-H and H(model) columns list the number of non-hydrogen atoms and hydrogen atoms in the chain respectively. The H(added) column lists the number of hydrogen atoms added and optimized by MolProbity. The Clashes column lists the number of clashes within the asymmetric unit, whereas Symm-Clashes lists symmetry-related clashes.

| Mol | Chain | Non-H | H(model) | H(added) | Clashes | Symm-Clashes |
|---|---|---|---|---|---|---|
| 1 | A | 822 | 0 | 733 | 11 | 0 |
| 1 | F | 827 | 0 | 751 | 10 | 0 |
| 2 | B | 1760 | 0 | 1639 | 32 | 0 |
| 2 | G | 1734 | 0 | 1599 | 34 | 0 |
| 3 | C | 2035 | 0 | 1690 | 12 | 0 |
| 3 | H | 2043 | 0 | 1714 | 15 | 0 |

*Continued on next page...*

For Manuscript Review

*Continued from previous page...*

| Mol | Chain | Non-H | H(model) | H(added) | Clashes | Symm-Clashes |
|---|---|---|---|---|---|---|
| 4 | D | 769 | 0 | 687 | 10 | 0 |
| 4 | I | 773 | 0 | 698 | 11 | 0 |
| 5 | E | 69 | 0 | 61 | 1 | 0 |
| 5 | J | 69 | 0 | 61 | 0 | 0 |
| All | All | 10901 | 0 | 9633 | 127 | 0 |

The all-atom clashscore is defined as the number of clashes found per 1000 atoms (including hydrogen atoms). The all-atom clashscore for this structure is 6.

All (127) close contacts within the same asymmetric unit are listed below, sorted by their clash magnitude.

| Atom-1 | Atom-2 | Interatomic distance (Å) | Clash overlap (Å) |
|---|---|---|---|
| 4:D:12:ARG:HH21 | 4:D:22:ILE:HG21 | 1.49 | 0.78 |
| 1:A:32:TYR:H | 1:A:92:ILE:HG12 | 1.52 | 0.75 |
| 4:I:12:ARG:HH21 | 4:I:22:ILE:HG21 | 1.54 | 0.73 |
| 4:D:24:ASN:HB3 | 4:D:65:LEU:HD11 | 1.72 | 0.71 |
| 4:I:29:GLN:HA | 4:I:61:SER:HB2 | 1.73 | 0.70 |
| 1:F:32:TYR:H | 1:F:92:ILE:HG12 | 1.57 | 0.70 |
| 4:I:24:ASN:HB3 | 4:I:65:LEU:HD11 | 1.72 | 0.69 |
| 2:G:95:ALA:H | 2:G:99:ASN:HD21 | 1.40 | 0.67 |
| 2:B:95:ALA:H | 2:B:99:ASN:HD21 | 1.40 | 0.67 |
| 4:D:29:GLN:HA | 4:D:61:SER:HB2 | 1.74 | 0.67 |
| 2:G:132:PRO:HG3 | 2:G:144:THR:H | 1.60 | 0.67 |
| 2:G:67:LEU:HB3 | 2:G:79:GLN:HB2 | 1.77 | 0.66 |
| 2:B:67:LEU:HB3 | 2:B:79:GLN:HB2 | 1.76 | 0.65 |
| 1:A:13:VAL:HG22 | 1:A:109:LEU:HD11 | 1.79 | 0.65 |
| 2:B:38:GLN:HB3 | 2:B:88:THR:HG23 | 1.79 | 0.65 |
| 2:B:132:PRO:HG3 | 2:B:144:THR:H | 1.61 | 0.65 |
| 2:G:142:LYS:HG2 | 2:G:193:ARG:HG2 | 1.81 | 0.62 |
| 1:F:13:VAL:HG22 | 1:F:109:LEU:HD11 | 1.82 | 0.61 |
| 2:B:61:LEU:HD23 | 2:B:63:GLY:H | 1.66 | 0.61 |
| 2:G:61:LEU:HD23 | 2:G:63:GLY:H | 1.67 | 0.59 |
| 2:G:36:TYR:HB3 | 2:G:44:LEU:HD11 | 1.86 | 0.58 |
| 3:H:5:MET:HB2 | 3:H:168:LEU:HD13 | 1.86 | 0.58 |
| 2:G:46:TRP:HZ2 | 2:G:49:THR:HG22 | 1.70 | 0.57 |
| 2:B:46:TRP:HZ2 | 2:B:49:THR:HG22 | 1.70 | 0.56 |
| 2:G:97:ARG:HH21 | 3:H:73:TRP:HB2 | 1.70 | 0.56 |
| 2:B:66:TYR:HB3 | 2:B:78:LEU:HD11 | 1.88 | 0.56 |
| 2:G:38:GLN:HB3 | 2:G:88:THR:HG23 | 1.87 | 0.56 |
| 1:A:21:LEU:HD22 | 1:A:107:THR:HG21 | 1.87 | 0.55 |
| 4:D:5:PRO:HB3 | 4:D:30:PHE:HB3 | 1.89 | 0.55 |

*Continued on next page...*

For Manuscript Review

Continued from previous page...

| Atom-1 | Atom-2 | Interatomic distance (Å) | Clash overlap (Å) |
|---|---|---|---|
| 2:B:11:ARG:HG2 | 2:B:19:VAL:HG21 | 1.89 | 0.54 |
| 4:D:30:PHE:HB2 | 4:D:84:HIS:CE1 | 2.41 | 0.54 |
| 4:I:30:PHE:HB2 | 4:I:84:HIS:CE1 | 2.43 | 0.54 |
| 2:B:174:THR:HG23 | 2:B:190:SER:HB2 | 1.89 | 0.54 |
| 2:G:66:TYR:HB3 | 2:G:78:LEU:HD11 | 1.88 | 0.54 |
| 2:B:36:TYR:HB3 | 2:B:44:LEU:HD11 | 1.89 | 0.54 |
| 3:H:82:LEU:HD11 | 3:H:89:ALA:HA | 1.90 | 0.53 |
| 3:C:5:MET:HB2 | 3:C:168:LEU:HD13 | 1.89 | 0.53 |
| 3:C:82:LEU:HD11 | 3:C:89:ALA:HA | 1.91 | 0.53 |
| 3:C:133:TRP:HB2 | 3:C:144:ARG:HG3 | 1.91 | 0.53 |
| 2:G:40:LEU:HD11 | 2:G:85:GLN:HB3 | 1.90 | 0.52 |
| 2:B:40:LEU:HD11 | 2:B:85:GLN:HB3 | 1.90 | 0.52 |
| 3:H:260:ARG:HA | 3:H:271:THR:HA | 1.91 | 0.52 |
| 4:I:5:PRO:HB3 | 4:I:30:PHE:HB3 | 1.91 | 0.52 |
| 1:A:23:CYS:HB3 | 1:A:72:VAL:HG13 | 1.91 | 0.52 |
| 2:G:63:GLY:HA3 | 2:G:87:ARG:NH2 | 2.24 | 0.52 |
| 3:H:102:ASP:HB2 | 3:H:111:ARG:HB3 | 1.92 | 0.52 |
| 4:D:31:HIS:HD1 | 4:D:62:PHE:HE2 | 1.58 | 0.52 |
| 3:C:102:ASP:HB2 | 3:C:111:ARG:HB3 | 1.91 | 0.51 |
| 2:G:11:ARG:HG2 | 2:G:19:VAL:HG21 | 1.92 | 0.51 |
| 1:A:95:THR:HG23 | 1:A:98:ASN:H | 1.74 | 0.51 |
| 2:B:3:LEU:HD12 | 2:B:4:LEU:HB2 | 1.93 | 0.50 |
| 2:B:21:LEU:HD13 | 2:B:111:THR:HG21 | 1.93 | 0.50 |
| 2:G:98:ARG:HG2 | 2:G:99:ASN:H | 1.76 | 0.50 |
| 2:B:71:VAL:HG12 | 2:B:72:THR:HG23 | 1.93 | 0.50 |
| 1:F:23:CYS:HB3 | 1:F:72:VAL:HG13 | 1.94 | 0.50 |
| 2:B:58:VAL:HG12 | 2:B:67:LEU:HD13 | 1.93 | 0.49 |
| 2:G:3:LEU:HD12 | 2:G:4:LEU:HB2 | 1.94 | 0.49 |
| 3:C:260:ARG:HA | 3:C:271:THR:HA | 1.93 | 0.49 |
| 2:B:117:GLU:O | 2:B:121:ASN:HA | 2.12 | 0.49 |
| 2:B:63:GLY:HA3 | 2:B:87:ARG:NH2 | 2.28 | 0.49 |
| 3:H:238:ASP:N | 4:I:12:ARG:HD3 | 2.29 | 0.48 |
| 2:G:30:TYR:CB | 2:G:95:ALA:HA | 2.44 | 0.48 |
| 2:G:4:LEU:HD12 | 2:G:108:ALA:HB2 | 1.96 | 0.47 |
| 2:G:71:VAL:HG12 | 2:G:72:THR:HG23 | 1.95 | 0.47 |
| 2:B:46:TRP:CZ2 | 2:B:49:THR:HG22 | 2.49 | 0.47 |
| 2:G:46:TRP:CZ2 | 2:G:49:THR:HG22 | 2.48 | 0.47 |
| 1:F:65:LEU:HD13 | 1:F:72:VAL:HB | 1.95 | 0.47 |
| 2:G:40:LEU:HD12 | 2:G:40:LEU:H | 1.80 | 0.47 |
| 4:I:31:HIS:HD1 | 4:I:62:PHE:HE2 | 1.63 | 0.47 |
| 2:B:102:ASN:HB3 | 3:C:149:GLN:O | 2.15 | 0.46 |

Continued on next page...

For Manuscript Review

*Continued from previous page...*

| Atom-1 | Atom-2 | Interatomic distance (Å) | Clash overlap (Å) |
|---|---|---|---|
| 2:G:174:THR:HG23 | 2:G:190:SER:HB2 | 1.96 | 0.46 |
| 2:G:97:ARG:NH2 | 3:H:73:TRP:HB2 | 2.30 | 0.46 |
| 2:B:31:PRO:HG2 | 2:B:51:ARG:HE | 1.80 | 0.46 |
| 2:G:95:ALA:H | 2:G:99:ASN:ND2 | 2.10 | 0.46 |
| 1:F:81:PRO:HA | 1:F:111:VAL:HB | 1.97 | 0.46 |
| 2:B:30:TYR:CB | 2:B:95:ALA:HA | 2.45 | 0.46 |
| 4:I:77:THR:HB | 4:I:94:TYR:HE2 | 1.81 | 0.46 |
| 2:B:33:MET:HA | 2:B:93:CYS:HA | 1.97 | 0.46 |
| 2:G:58:VAL:HG12 | 2:G:67:LEU:HD13 | 1.98 | 0.46 |
| 3:H:130:LEU:HB2 | 3:H:157:LYS:HE2 | 1.98 | 0.46 |
| 2:B:98:ARG:HG2 | 2:B:99:ASN:H | 1.81 | 0.45 |
| 2:B:137:ILE:HG21 | 2:B:200:HIS:CE1 | 2.52 | 0.45 |
| 2:G:101:TYR:HB3 | 3:H:150:SER:O | 2.16 | 0.45 |
| 1:A:65:LEU:HD13 | 1:A:72:VAL:HB | 1.97 | 0.45 |
| 2:B:95:ALA:H | 2:B:99:ASN:ND2 | 2.11 | 0.45 |
| 3:H:208:PHE:CE1 | 3:H:241:PHE:HB2 | 2.52 | 0.45 |
| 2:B:40:LEU:H | 2:B:40:LEU:HD12 | 1.81 | 0.44 |
| 3:C:208:PHE:CE1 | 3:C:241:PHE:HB2 | 2.52 | 0.44 |
| 1:F:95:THR:HG23 | 1:F:98:ASN:H | 1.82 | 0.44 |
| 3:H:217:TRP:CD2 | 3:H:259:CYS:HB2 | 2.52 | 0.44 |
| 1:A:81:PRO:HA | 1:A:111:VAL:HB | 1.99 | 0.44 |
| 2:B:4:LEU:HD12 | 2:B:108:ALA:HB2 | 2.00 | 0.44 |
| 2:G:21:LEU:HD13 | 2:G:111:THR:HG21 | 1.99 | 0.44 |
| 4:D:77:THR:HB | 4:D:94:TYR:HE2 | 1.82 | 0.44 |
| 2:G:31:PRO:HG2 | 2:G:51:ARG:HE | 1.82 | 0.44 |
| 1:F:34:TRP:HB2 | 1:F:90:ALA:HB3 | 1.99 | 0.44 |
| 1:F:98:ASN:O | 2:G:98:ARG:HG3 | 2.17 | 0.43 |
| 3:H:236:ALA:H | 3:H:241:PHE:HA | 1.83 | 0.43 |
| 2:G:13:VAL:HG22 | 2:G:14:PRO:HD2 | 1.99 | 0.43 |
| 2:B:13:VAL:HG22 | 2:B:14:PRO:HD2 | 2.01 | 0.43 |
| 3:C:185:PRO:HB3 | 3:C:208:PHE:HB3 | 2.01 | 0.43 |
| 2:G:11:ARG:O | 2:G:113:LEU:HA | 2.19 | 0.43 |
| 1:F:61:PHE:HD2 | 1:F:76:ILE:HG12 | 1.83 | 0.43 |
| 3:C:217:TRP:CD2 | 3:C:259:CYS:HB2 | 2.54 | 0.43 |
| 1:F:39:HIS:CD2 | 1:F:85:ALA:HB2 | 2.54 | 0.43 |
| 3:H:185:PRO:HB3 | 3:H:208:PHE:HB3 | 2.00 | 0.43 |
| 1:A:34:TRP:HB2 | 1:A:90:ALA:HB3 | 2.01 | 0.43 |
| 3:H:22:TYR:CZ | 3:H:24:SER:HB2 | 2.54 | 0.43 |
| 4:I:9:VAL:HG12 | 4:I:95:TRP:HD1 | 1.84 | 0.42 |
| 4:D:9:VAL:HG12 | 4:D:95:TRP:HD1 | 1.85 | 0.42 |
| 3:C:236:ALA:H | 3:C:241:PHE:HA | 1.84 | 0.42 |

*Continued on next page...*

For Manuscript Review

*Continued from previous page...*

| Atom-1 | Atom-2 | Interatomic distance (Å) | Clash overlap (Å) |
|---|---|---|---|
| 3:H:204:TRP:CH2 | 4:I:98:ASP:HB2 | 2.55 | 0.42 |
| 1:A:92:ILE:HB | 2:B:98:ARG:CZ | 2.50 | 0.41 |
| 2:G:30:TYR:HB2 | 2:G:95:ALA:HA | 2.02 | 0.41 |
| 2:G:33:MET:HA | 2:G:93:CYS:HA | 2.02 | 0.41 |
| 3:C:159:TYR:CD1 | 5:E:3:ASN:HB2 | 2.56 | 0.41 |
| 1:A:39:HIS:CD2 | 1:A:85:ALA:HB2 | 2.56 | 0.41 |
| 2:B:64:ALA:HB1 | 2:B:80:VAL:HG23 | 2.03 | 0.41 |
| 3:C:130:LEU:HB2 | 3:C:157:LYS:HE2 | 2.02 | 0.41 |
| 2:B:10:TRP:HE3 | 2:B:112:ARG:HB3 | 1.85 | 0.41 |
| 4:D:84:HIS:CD2 | 4:D:85:ASP:H | 2.39 | 0.41 |
| 4:D:12:ARG:HG2 | 4:D:13:HIS:ND1 | 2.36 | 0.41 |
| 2:G:66:TYR:CE1 | 2:G:89:LEU:HD21 | 2.56 | 0.40 |
| 2:B:11:ARG:O | 2:B:113:LEU:HA | 2.21 | 0.40 |
| 1:A:13:VAL:O | 1:A:111:VAL:HA | 2.21 | 0.40 |
| 2:G:88:THR:HA | 2:G:112:ARG:HA | 2.03 | 0.40 |
| 4:I:9:VAL:HG12 | 4:I:95:TRP:CD1 | 2.57 | 0.40 |

There are no symmetry-related clashes.

#### 5.3 Torsion angles (i)

##### 5.3.1 Protein backbone (i)

In the following table, the Percentiles column shows the percent Ramachandran outliers of the chain as a percentile score with respect to all X-ray entries followed by that with respect to entries of similar resolution.

The Analysed column shows the number of residues for which the backbone conformation was analysed, and the total number of residues.

| Mol | Chain | Analysed | Favoured | Allowed | Outliers | Percentiles | |
|---|---|---|---|---|---|---|---|
| 1 | A | 112/214 (52%) | 106 (95%) | 6 (5%) | 0 | 100 | 100 |
| 1 | F | 110/214 (51%) | 106 (96%) | 4 (4%) | 0 | 100 | 100 |
| 2 | B | 223/253 (88%) | 198 (89%) | 23 (10%) | 2 (1%) | 14 | 51 |
| 2 | G | 220/253 (87%) | 197 (90%) | 21 (10%) | 2 (1%) | 14 | 51 |
| 3 | C | 276/281 (98%) | 253 (92%) | 23 (8%) | 0 | 100 | 100 |
| 3 | H | 275/281 (98%) | 252 (92%) | 23 (8%) | 0 | 100 | 100 |
| 4 | D | 96/100 (96%) | 88 (92%) | 8 (8%) | 0 | 100 | 100 |

*Continued on next page...*

For Manuscript Review

*Continued from previous page...*

| Mol | Chain | Analysed | Favoured | Allowed | Outliers | Percentiles | |
|---|---|---|---|---|---|---|---|
| 4 | I | 96/100 (96%) | 88 (92%) | 8 (8%) | 0 | 100 | 100 |
| 5 | E | 7/9 (78%) | 6 (86%) | 1 (14%) | 0 | 100 | 100 |
| 5 | J | 7/9 (78%) | 6 (86%) | 1 (14%) | 0 | 100 | 100 |
| All | All | 1422/1714 (83%) | 1300 (91%) | 118 (8%) | 4 (0%) | 37 | 72 |

All (4) Ramachandran outliers are listed below:

| Mol | Chain | Res | Type |
|---|---|---|---|
| 2 | B | 31 | PRO |
| 2 | G | 31 | PRO |
| 2 | B | 104 | PRO |
| 2 | G | 104 | PRO |

##### 5.3.2 Protein sidechains (i)

In the following table, the Percentiles column shows the percent sidechain outliers of the chain as a percentile score with respect to all X-ray entries followed by that with respect to entries of similar resolution.

The Analysed column shows the number of residues for which the sidechain conformation was analysed, and the total number of residues.

| Mol | Chain | Analysed | Rotameric | Outliers | Percentiles | |
|---|---|---|---|---|---|---|
| 1 | A | 83/188 (44%) | 82 (99%) | 1 (1%) | 67 | 79 |
| 1 | F | 85/188 (45%) | 84 (99%) | 1 (1%) | 67 | 79 |
| 2 | B | 180/225 (80%) | 171 (95%) | 9 (5%) | 20 | 42 |
| 2 | G | 178/225 (79%) | 171 (96%) | 7 (4%) | 27 | 49 |
| 3 | C | 173/239 (72%) | 172 (99%) | 1 (1%) | 84 | 88 |
| 3 | H | 175/239 (73%) | 174 (99%) | 1 (1%) | 84 | 88 |
| 4 | D | 81/95 (85%) | 81 (100%) | 0 | 100 | 100 |
| 4 | I | 82/95 (86%) | 81 (99%) | 1 (1%) | 67 | 79 |
| 5 | E | 8/8 (100%) | 6 (75%) | 2 (25%) | 0 | 3 |
| 5 | J | 8/8 (100%) | 6 (75%) | 2 (25%) | 0 | 3 |
| All | All | 1053/1510 (70%) | 1028 (98%) | 25 (2%) | 44 | 63 |

All (25) residues with a non-rotameric sidechain are listed below:

For Manuscript Review

| Mol | Chain | Res | Type |
|---|---|---|---|
| 1 | A | 31 | GLN |
| 2 | B | 4 | LEU |
| 2 | B | 13 | VAL |
| 2 | B | 22 | ARG |
| 2 | B | 30 | TYR |
| 2 | B | 87 | ARG |
| 2 | B | 105 | LEU |
| 2 | B | 116 | THR |
| 2 | B | 117 | GLU |
| 2 | B | 208 | CYS |
| 3 | C | 164 | CYS |
| 5 | E | 7 | GLU |
| 5 | E | 9 | MET |
| 1 | F | 31 | GLN |
| 2 | G | 4 | LEU |
| 2 | G | 13 | VAL |
| 2 | G | 30 | TYR |
| 2 | G | 87 | ARG |
| 2 | G | 105 | LEU |
| 2 | G | 116 | THR |
| 2 | G | 208 | CYS |
| 3 | H | 164 | CYS |
| 4 | I | 98 | ASP |
| 5 | J | 7 | GLU |
| 5 | J | 9 | MET |

Sometimes sidechains can be flipped to improve hydrogen bonding and reduce clashes. All (1) such sidechains are listed below:

| Mol | Chain | Res | Type |
|---|---|---|---|
| 4 | D | 6 | GLN |

##### 5.3.3 RNA (i)

There are no RNA molecules in this entry.

#### 5.4 Non-standard residues in protein, DNA, RNA chains (i)

There are no non-standard protein/DNA/RNA residues in this entry.

For Manuscript Review

#### 5.5 Carbohydrates (i)

There are no oligosaccharides in this entry.

#### 5.6 Ligand geometry (i)

There are no ligands in this entry.

#### 5.7 Other polymers (i)

There are no such residues in this entry.

#### 5.8 Polymer linkage issues (i)

There are no chain breaks in this entry.

For Manuscript Review

### 6 Fit of model and data (i)

#### 6.1 Protein, DNA and RNA chains (i)

In the following table, the column labelled '#RSRZ> 2' contains the number (and percentage) of RSRZ outliers, followed by percent RSRZ outliers for the chain as percentile scores relative to all X-ray entries and entries of similar resolution. The OWAB column contains the minimum, median, $95^{th}$ percentile and maximum values of the occupancy-weighted average B-factor per residue. The column labelled 'Q< 0.9' lists the number of (and percentage) of residues with an average occupancy less than 0.9.

| Mol | Chain | Analysed | <RSRZ> | #RSRZ>2 | | | OWAB(Å$^2$) | Q<0.9 |
|---|---|---|---|---|---|---|---|---|
| 1 | A | 114/214 (53%) | -0.13 | 0 | 100 | 100 | 183, 250, 301, 326 | 0 |
| 1 | F | 112/214 (52%) | 0.01 | 1 (0%) | 81 | 67 | 182, 267, 340, 351 | 0 |
| 2 | B | 227/253 (89%) | -0.08 | 3 (1%) | 74 | 59 | 179, 250, 360, 440 | 0 |
| 2 | G | 224/253 (88%) | -0.01 | 3 (1%) | 74 | 59 | 186, 248, 351, 395 | 0 |
| 3 | C | 278/281 (98%) | -0.09 | 2 (0%) | 84 | 71 | 158, 221, 279, 321 | 0 |
| 3 | H | 277/281 (98%) | -0.11 | 1 (0%) | 89 | 79 | 167, 227, 281, 296 | 0 |
| 4 | D | 98/100 (98%) | 0.05 | 2 (2%) | 64 | 49 | 173, 210, 283, 326 | 0 |
| 4 | I | 98/100 (98%) | -0.06 | 1 (1%) | 79 | 64 | 169, 216, 301, 339 | 0 |
| 5 | E | 9/9 (100%) | 0.21 | 0 | 100 | 100 | 186, 221, 268, 290 | 0 |
| 5 | J | 9/9 (100%) | 0.35 | 0 | 100 | 100 | 193, 222, 262, 298 | 0 |
| All | All | 1446/1714 (84%) | -0.06 | 13 (0%) | 81 | 67 | 158, 236, 330, 440 | 0 |

All (13) RSRZ outliers are listed below:

| Mol | Chain | Res | Type | RSRZ |
|---|---|---|---|---|
| 3 | C | 187 | ALA | 4.1 |
| 2 | B | 127 | VAL | 3.5 |
| 4 | D | 25 | CYS | 3.0 |
| 2 | G | 130 | PHE | 2.9 |
| 2 | B | 230 | THR | 2.9 |
| 2 | B | 154 | PRO | 2.9 |
| 4 | I | 26 | TYR | 2.7 |
| 3 | H | 74 | PHE | 2.3 |
| 2 | G | 237 | ALA | 2.3 |
| 2 | G | 127 | VAL | 2.2 |
| 3 | C | 76 | VAL | 2.1 |
| 1 | F | 91 | ALA | 2.1 |
| 4 | D | 26 | TYR | 2.1 |

For Manuscript Review

#### 6.2 Non-standard residues in protein, DNA, RNA chains (i)

There are no non-standard protein/DNA/RNA residues in this entry.

#### 6.3 Carbohydrates (i)

There are no monosaccharides in this entry.

#### 6.4 Ligands (i)

There are no ligands in this entry.

#### 6.5 Other polymers (i)

There are no such residues in this entry.

For Manuscript Review
